# Self-reported sleep relates to hippocampal atrophy across the adult lifespan – results from the Lifebrain consortium

**DOI:** 10.1101/737858

**Authors:** Anders M. Fjell, Øystein Sørensen, Inge K. Amlien, David Bartrés-Faz, Didac Maciá Bros, Ilja Demuth, Christian A Drevon, Sandra Düzel, Klaus P. Ebmeier, Ane-Victoria Idland, Tim C. Kietzmann, Rogier Kievit, Simone Kühn, Ulman Lindenberger, Athanasia M Mowinckel, Lars Nyberg, Darren Price, Claire E. Sexton, Cristina Solé-Padullés, Sara Pudas, Donatas Sederevicius, Sana Suri, Gerd Wagner, Leiv Otto Watne, René Westerhausen, Enikő Zsoldos, Kristine B. Walhovd

## Abstract

**Background:** Poor sleep is associated with multiple age-related neurodegenerative and neuropsychiatric conditions. The hippocampus plays a special role in sleep and sleep-dependent cognition, and accelerated hippocampal atrophy is typically seen with higher age. Hence, it is critical to establish how the relationship between sleep and hippocampal volume loss unfolds across the adult lifespan.

**Methods:** Self-reported sleep measures and MRI-derived hippocampal volumes were obtained from 3105 cognitively normal participants (18-90 years) from major European brain studies in the Lifebrain consortium. Hippocampal volume change was estimated from 5116 MRIs from 1299 participants, covering up to 11 years. Cross-sectional analyses were repeated in a sample of 21390 participants from the UK Biobank.

**Results:** The relationship between self-reported sleep and age differed across sleep items. Sleep duration, efficiency, problems, and use of medication worsened monotonously with age, whereas subjective sleep quality, sleep latency, and daytime tiredness improved. Women reported worse sleep in general than men, but the relationship to age was similar. No cross-sectional sleep – hippocampal volume relationships was found. However, worse sleep quality, efficiency, problems, and daytime tiredness were related to greater hippocampal volume loss over time, with high scorers showing on average 0.22% greater annual loss than low scorers. Simulations showed that longitudinal effects were too small to be detected as age-interactions in cross-sectional analyses.

**Conclusions:** Worse self-reported sleep is associated with higher rates of hippocampal decline across the adult lifespan. This suggests that sleep is relevant to understand individual differences in hippocampal atrophy, but limited effect sizes call for cautious interpretation.

## Introduction

Disturbed sleep is associated with normal aging [1–3] and several age-related neurological and psychiatric conditions, including dementia [4–9]. The hippocampus plays a special role in sleep and sleep-dependent cognition [10, 11], and hippocampal atrophy increases in normal aging [12] and Alzheimer’s disease (AD) [13, 14]. Rodent research has shown that sleep deprivation can reduce spine density and attenuate synaptic efficacy in the hippocampus [15], possibly through changes in neural plasticity, reduction of hippocampal cell proliferation and neurogenesis [16]. Thus, it has been suggested that the hippocampus may be especially sensitive to sleep deprivation [16] over extended time periods [17].

Inspired by the mechanistic relationship between sleep and hippocampal morphology established in rodents, studies have tested the association between sleep and hippocampal volume in humans (see Table 1). Most have compared patients with different sleep-related conditions and normal controls. Many report smaller hippocampi in patients [18–22], or an inverse relationship between hippocampal volume and poor sleep [23, 24]. However, other studies found no relationship [25–27] or even larger volumes in patients [28]. Of three studies testing the relationship between self-reported sleep and hippocampal volume in healthy older adults, two reported that worse sleep or fatigue was associated with lower hippocampal volume [29, 30] whereas one found no significant relationship [31]. Two studies tested the association between self-reported sleep and longitudinal changes in hippocampal volume, and neither found significant effects [32, 33]. Given the sparsity of longitudinal studies, it is important to assess the relationship between self-reported sleep and hippocampal volume changes with high statistical power [34], which was the main purpose of the present study. We took advantage of the large-scale, European multi-site longitudinal Lifebrain consortium (http://www.lifebrain.uio.no/), to address the following main question: Is self-reported sleep related to hippocampal volume and volume change? We investigated this using an adult lifespan approach, testing if and how sleep, hippocampal volume and their relationship change with age.

**Table 1.**
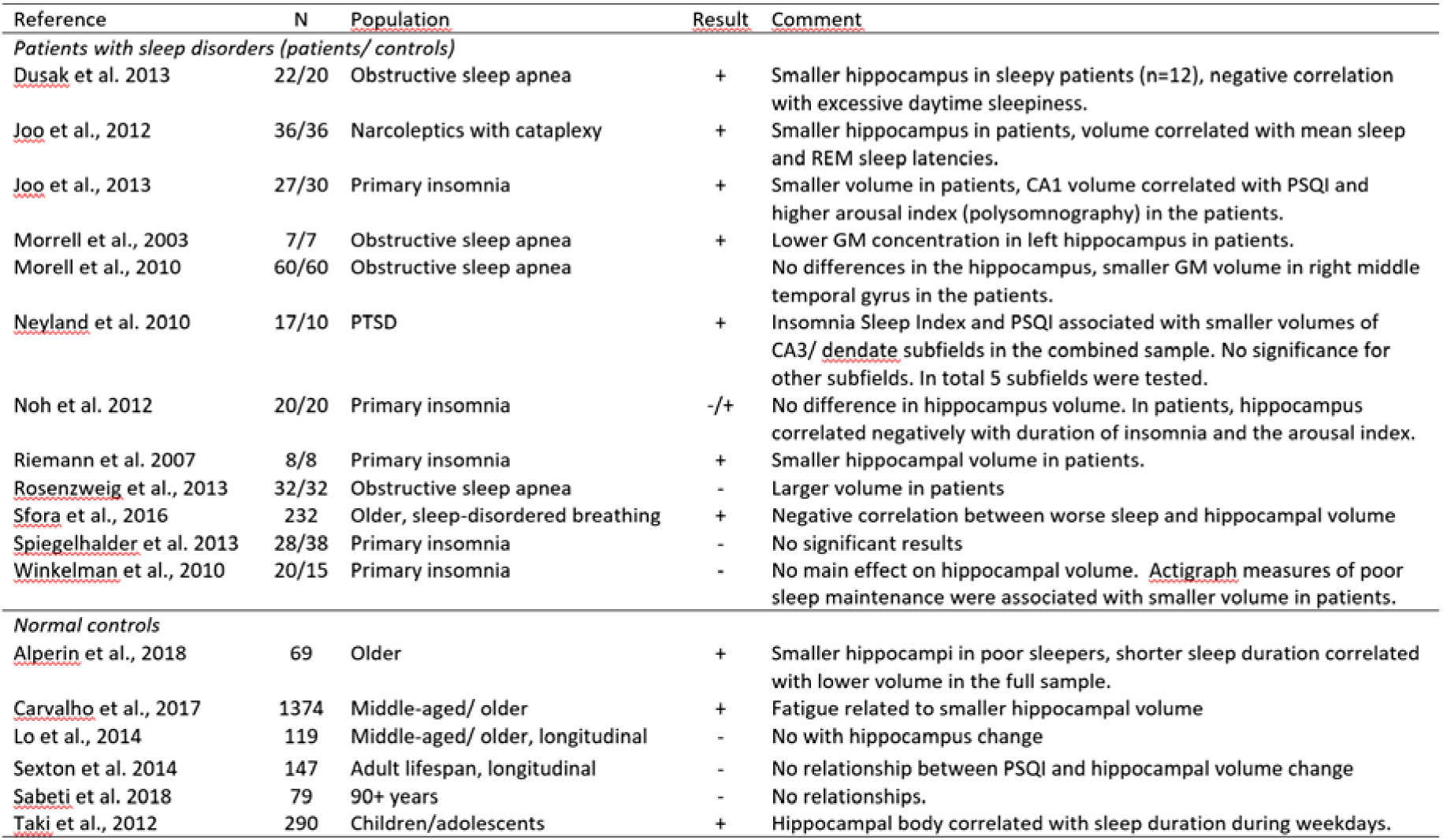
Representative studies on self-reported sleep, sleep disturbances and hippocampal volume. Result: “-“ indicates no relationship between hippocampus volume and sleep or an inverse relationship (e.g. higher volume in patients). “+” indicates the expected relationship between hippocampus volume and sleep, e.g. smaller volume in patients or a negative correlation between sleep problems and volume. Results for other brain structures than the hippocampus are not reported.

To ensure that the results were not specific to the self-report instrument (Pittsburgh Sleep Quality Inventory – PSQI) [35]) and hippocampus segmentation (FreeSurfer [36]) used in Lifebrain, cross-sectional replication analyses were performed using UK Biobank (UKB) data with different self-report measures of sleep and a different hippocampal segmentation approach.

## Methods and Materials

### Lifebrain sample

The sample was derived from the European Lifebrain project (http://www.lifebrain.uio.no/) [37], including participants from major European brain studies: Berlin Study of Aging-II (BASE-II) [38, 39], the BETULA project ([40], the Cambridge Centre for Ageing and Neuroscience study (Cam-CAN) [41], Center for Lifebrain Changes in Brain and Cognition longitudinal studies (LCBC) [42, 43], Whitehall-II (WH-II) [44], and University of Barcelona brain studies [45–47]. In total, self-reported sleep and hippocampal volume data from 3105 participants (18-90 years) were included. Longitudinal information on hippocampal volume was available for 1298 participants, yielding a total of 5116 observations. Mean interval from first to last examination was 3.5 years (range 0.2-11.0 years). Participants were screened to be cognitively healthy and in general not suffer from conditions known to affect brain function, such as dementia, major stroke, multiple sclerosis etc. Exact screening criteria were not identical across sub-samples. Sample characteristics are presented in Table 2, and detailed information about each sub-sample is presented in SI.

**Table 2.** Sample characteristics. Follow-up interval refers to time between MRI examinations

| Study | Unique participants | Observations | Mean follow-up interval (sd) | Max follow-up interval (range) | Age (range) | Sex (female/male) |
| --- | --- | --- | --- | --- | --- | --- |
| Barcelona | 145 | 222 | 3.1 (1.2) | 4.3 (3.7 - 4.9) | 69 (48 - 90) | 149/73 |
| BASE-II | 315 | 628 | 1.9 (0.7) | 1.9 (0.6 - 3.1) | 62 (24 - 81) | 223/405 |
| Betula | 311 | 500 | 4.0 (0.3) | 4.0 (3 - 5) | 61 (25 - 81) | 251/249 |
| Cam-CAN | 647 | 910 | 1.4 (0.7) | 1.4 (0.2 - 3.5) | 55 (18 - 89) | 464/446 |
| LCBC | 914 | 2083 | 2.9 (2.7) | 4.5 (0.2 - 11) | 52 (19 - 89) | 1308/775 |
| Whitehall-II Imaging | 773 | 773 | NA | NA | 70 (60 - 85) | 150/623 |
| Total Lifebrain | 3105 | 5116 | 2.6 (2.3) | 3.5 (0.2 - 11) | 58 (18 - 90) | 2545/2571 |
| Replication (UKB) | 21390 | 21390 | NA | NA | 63 (45 - 81) | 11237/ 10153 |

### Self-reported sleep assessment

Sleep was assessed using the Pittsburgh Sleep Quality Inventory (PSQI) [35], yielding seven domains (sleep quality, latency, duration, efficiency, problems, medication and daytime tiredness) and a global score, over a 1-month time interval. Each domain is scored 0-3 and the global 0-21. High scores indicate worse sleep, e.g. high score on the sleep duration scale means shorter sleep time. The results for the sleep medication question must be treated with caution, since most samples were screened for use of medications possibly affecting CNS function. The Karolinska Sleep Questionnaire (KSQ) [48, 49] was used for Betula. The items in KSQ cover almost perfectly the items in PSQI, and the KSQ was therefore transformed to PSQI scales (see SI for details). Since longitudinal information on sleep was lacking for most of the sample, and sleep quality tends to be stable across intervals up to years [50], sleep was treated as a trait variable. If multiple observations of sleep were available, these were averaged, and the mean value used in the analyses. We have previously found high stability of PSQI score across 3 years (r = .81 between examinations 3 years apart, see https://www.biorxiv.org/content/10.1101/335612v1).

### Magnetic resonance imaging acquisition and analysis

Lifebrain MRI data originated from ten different scanners (Table 3), mainly processed with FreeSurfer 6.0 (https://surfer.nmr.mgh.harvard.edu/) [36, 51–53] (FreeSurfer 5.3 was used for Whitehall-II), generating hippocampal and intracranial volume (ICV) estimates. Because FreeSurfer is almost fully automated, to avoid introducing possible site-specific biases, gross quality control measures were imposed and no manual editing was done. To assess the influence of scanner on hippocampal volume, seven participants were scanned on seven of the scanners (see SI for details). There was a significant main effect of scanner on hippocampal volume (F = 4.13, p = .046) in the Travelling Brains sample. However, the between-participant rank order was almost perfectly retained between scanners, yielding a mean between-scanner Pearson correlation for bilateral hippocampal volume of r = .98 (range .94-1.00). Thus, including site as a random effect covariate in the analyses of hippocampal volume is likely sufficient to remove the influence of scanner differences. Detailed results are found in SI.

**Table 3.**
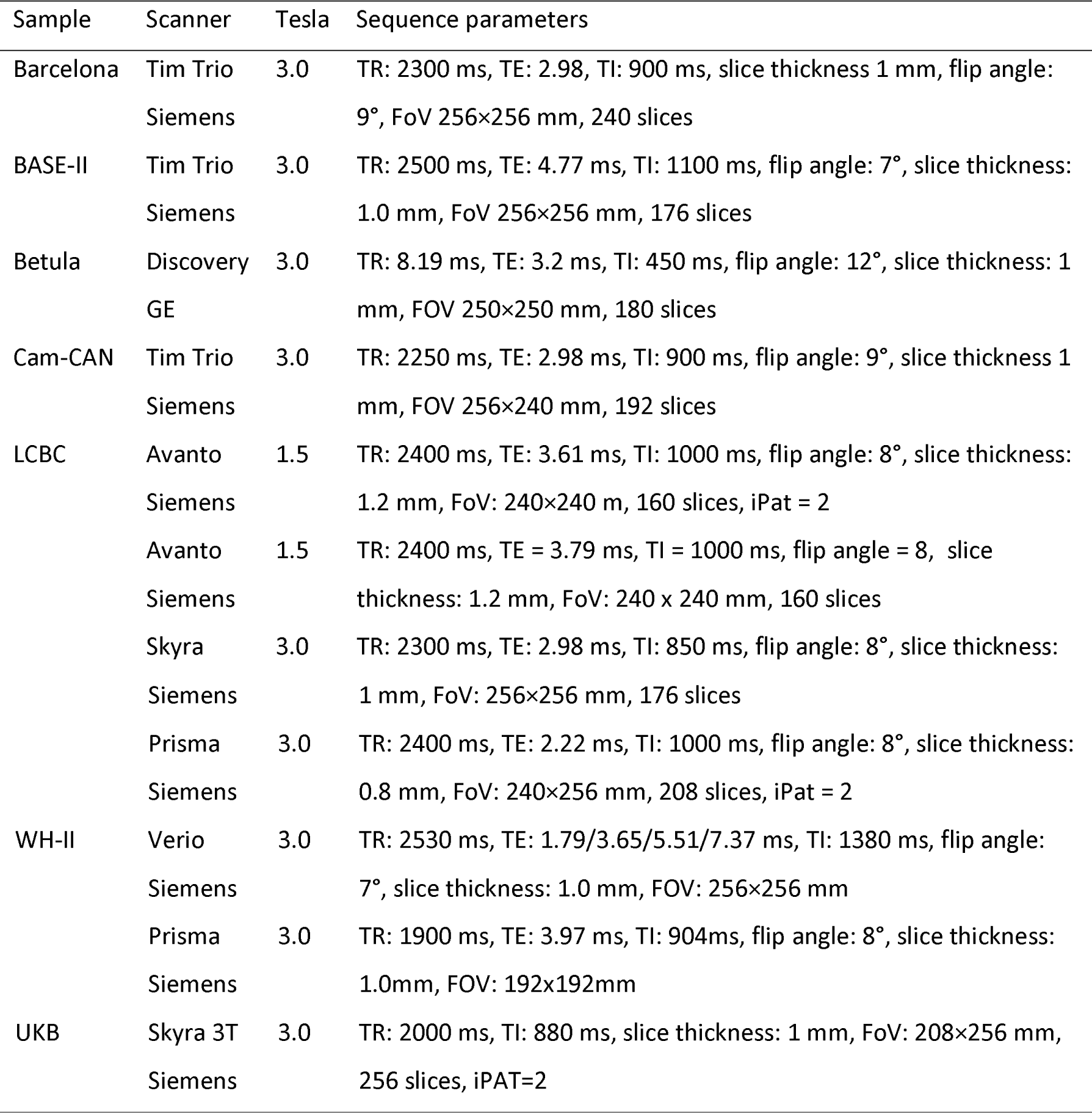
MR acquisition parameters. TR: Repetition time, TE: Echo time, TI: Inversion time, FoV: Field of View, iPat: in-plane acceleration

### Replication sample – UK Biobank

Cross-sectional analyses were repeated using 21390 participants from UKB (https://imaging.ukbiobank.ac.uk/) [54] (Table 2), with the sample size varying somewhat with number of missing data for each variable of interest (range 19782 – 21390). UKB does not contain PSQI, but includes several questions related to sleep (i.e. sleep duration, sleeplessness, daytime dozing/sleeping, daytime napping, problems getting up in the morning, snoring), allowing us to evaluate whether the Lifebrain results were specific to the PSQI scales. Hippocampal volume [55] and the volumetric scaling from T1 head image to standard space as proxy for ICV were used in the analyses, generated using publicly available tools, primarily based on FSL (FMRIB Software library, https://fsl.fmrib.ox.ac.uk/fsl/fslwiki). Details of the imaging protocol (http://biobank.ctsu.ox.ac.uk/crystal/refer.cgi?id=2367) and structural image processing are provided on the UK biobank website (http://biobank.ctsu.ox.ac.uk/crystal/refer.cgi?id=1977) (see Table 3). Sleep and MRI data were retrieved for the same participant examination. For detailed description of how the UK Biobank data were retrieved and analyzed, see SI.

### Statistical analyses

Analyses were run in R version 3.4.4 [56]. Generalized Additive Models (GAM) and Generalized Additive Mixed Models (GAMM) using the package “mgcv” version 1.8-28 [57] were used to derive age-functions for the different sleep variables and hippocampal volume, and to test the relationship between sleep, hippocampal volume and volume change. We used smooth terms for age and sleep, random effects for subject and study, and sex as covariates. For analyses including hippocampal volume, estimated intracranial volume (ICV) was an additional covariate. Interactions between age and sex were tested in separate models. Longitudinal models were tested by including time since baseline and sleep × time as predictors. Additional models were run controlling for symptoms of depression and body mass index (BMI). The R-code and full description of the procedures and results are given in SI.

## Results

### Relationships between self-reported sleep and age

The association between age and sleep was significant for all PSQI sub-scales as well as PSQI global (Table 4 and Figure 1). The relationships varied across scales and deviated from linearity. Scores for sleep duration, efficiency, sleep problems and use of medication increased monotonously with higher age, i.e. worse sleep with higher age. In contrast, sleep quality improved almost linearly, whereas problems with sleep latency were gradually reduced until about 50 years of age, before a slight increase was seen towards the end of the age range. Problems with daytime tiredness were decreased until about 70 years, before increasing in the last part of the lifespan. The global score was stable until mid-life, followed by a modest increase. These results suggest that the relationship between self-reported sleep and age is not uniform across different aspects of sleep.

**Figure 1.**
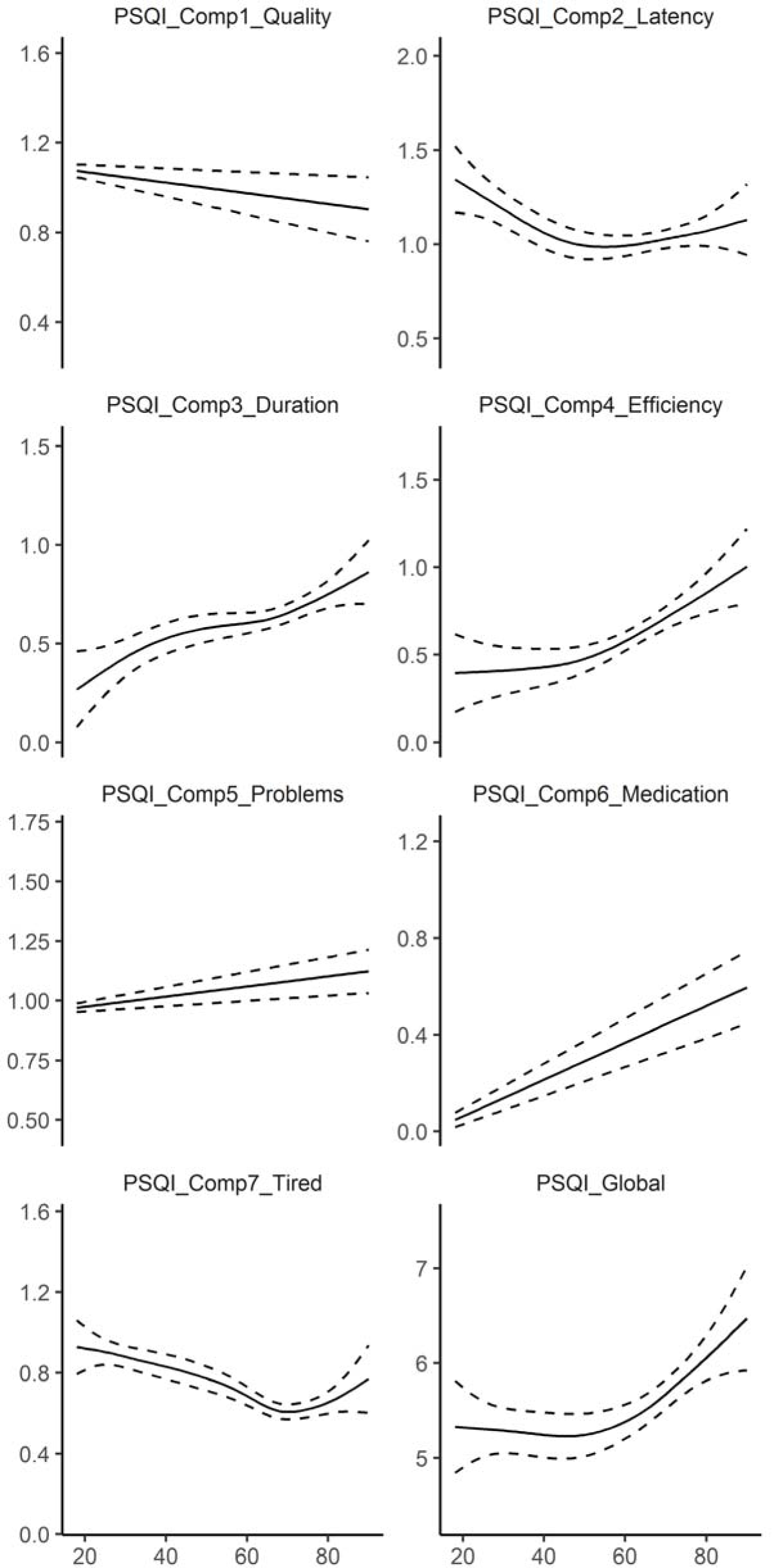
Relationships between age and self-reported sleep in Lifebrain. Generalized Additive Models (GAMs) were used to obtain age-curves for each sleep variable. Higher scores indicate worse sleep. Sex was included as covariate in the analyses. Dotted lines represent 95% CI.

**Table 4.** Associations between self-reported sleep and age in Lifebrain. GAMs are presented for each sleep variable, testing a smooth function of age and a linear function of sex. Study was included as a random effect term of no interest. Edf (effective degrees of freedom) and F-values are provided for age, whereas the linear estimate and the t-values are provided for sex. Negative estimates/ t-values indicate lower scores for men, i.e. less sleep problems. Only cross-sectional data were included in these analyses.

| Sleep scale | Variables | Edf/ estimate | F/t | p |
| --- | --- | --- | --- | --- |
| Quality | Age | 2.23 | 3.96 | .008 |
|  | Sex | -0.07 | -2.65 | .008 |
| Latency | Age | 2.87 | 4.15 | .004 |
|  | Sex | -0.26 | -7.71 | 1.8e<sup>-14</sup> |
| Duration | Age | 4.15 | 9.00 | 1.1e<sup>-08</sup> |
|  | Sex | -0.06 | -2.24 | .025 |
| Efficiency | Age | 2.67 | 27.67 | <2e<sup>-16</sup> |
|  | Sex | -0.17 | -4.89 | 1.1e<sup>-06</sup> |
| Problems | Age | 1.0 | 17.59 | 2.8e<sup>-05</sup> |
|  | Sex | -0.04 | -2.58 | .01 |
| Medication | Age | 1.02 | 79.78 | <2e<sup>-16</sup> |
|  | Sex | -0.11 | -3.85 | .0001 |
| Tiredness | Age | 4.13 | 13.99 | 9.4e<sup>-14</sup> |
|  | Sex | -0.01 | -0.37 | .71 |
| Global | Age | 2.59 | 7.74 | 2.6e<sup>-05</sup> |
|  | Sex | -0.76 | -6.63 | 4.0e<sup>-11</sup> |

For all sleep variables except daytime tiredness, women reported significantly worse sleep than men. The effect sizes were generally small, however, with the effect of being female generally being less than 0.2 PSQI sub-scale points. The largest main effect of sex was on latency, where women reported 0.26 points more on the 0-3 points PSQI scale. For the global score, women reported 0.76 points more than men, which is equal to 13.7% of the intercept of 5.55. We also tested age × sex interactions. These were not significant for any scale, suggesting that the association between self-reported sleep and age is similar for men and women.

### Self-reported sleep and hippocampal volume

Using both the cross-sectional and longitudinal MRI data to map the age-trajectory of hippocampal volume (Figure 2), we observed the expected non-linear decline which is more pronounced from about 60 years of age (edf = 7.3, F = 374.5, p < 2e^−16^). This was confirmed by a significant effect of age on longitudinal change over time (edf = 13.0, F = 162.5, p < 2e^−16^).

**Figure 2.**
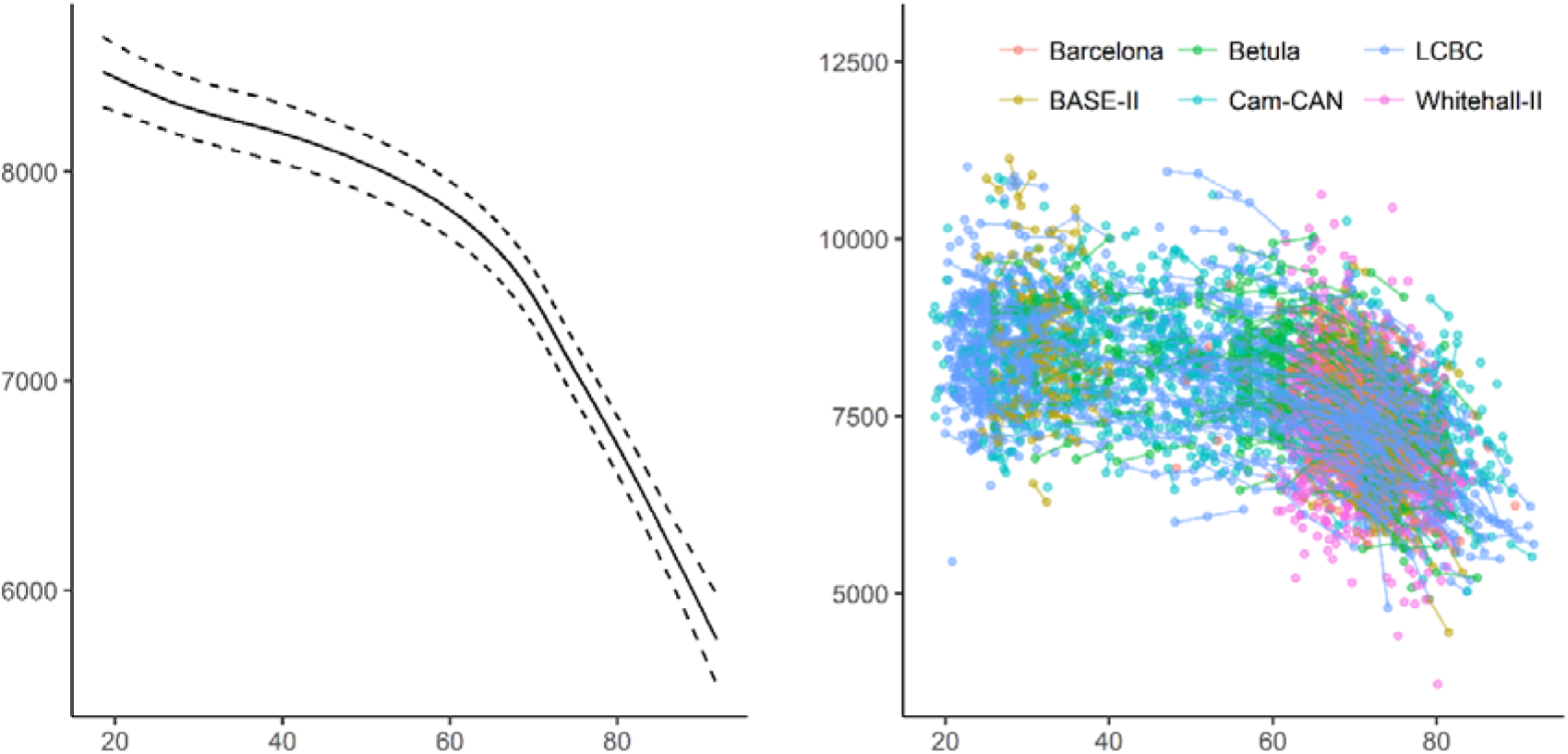
Relationships between age and hippocampal volume in Lifebrain. Left panel: GAMM was used to obtain the age-curve for hippocampal volume, using both cross-sectional and longitudinal information, covarying for sex, ICV and study (random effect). Dotted lines represent 95% CI. Right panel: Spaghetti plot of hippocampal volume and volume change for all participants, color-coded by sample. X-axis denotes age in years, y-axis hippocampal volume in mm^3^.

We tested for main effects of sleep on cross-sectional hippocampal volume. Study and Subject ID were random effects, age, sex and ICV were included as covariates of no interest (see SI for details). p was not below the Bonferroni-corrected threshold of .00625 (.05/8) for any of the relationships. We also tested for interaction with sex and age. No interactions with sex were found, whereas age and latency showed a very weak but significant interaction (edf = 1.08, F = 8.05, p = .0038), reflecting slightly higher hippocampal offset volumes and slightly greater age effects for those reporting better sleep. In sum, the cross-sectional relationships between self-reported sleep and hippocampal volume were very modest or non-existing (Figure 3).

**Figure 3.**
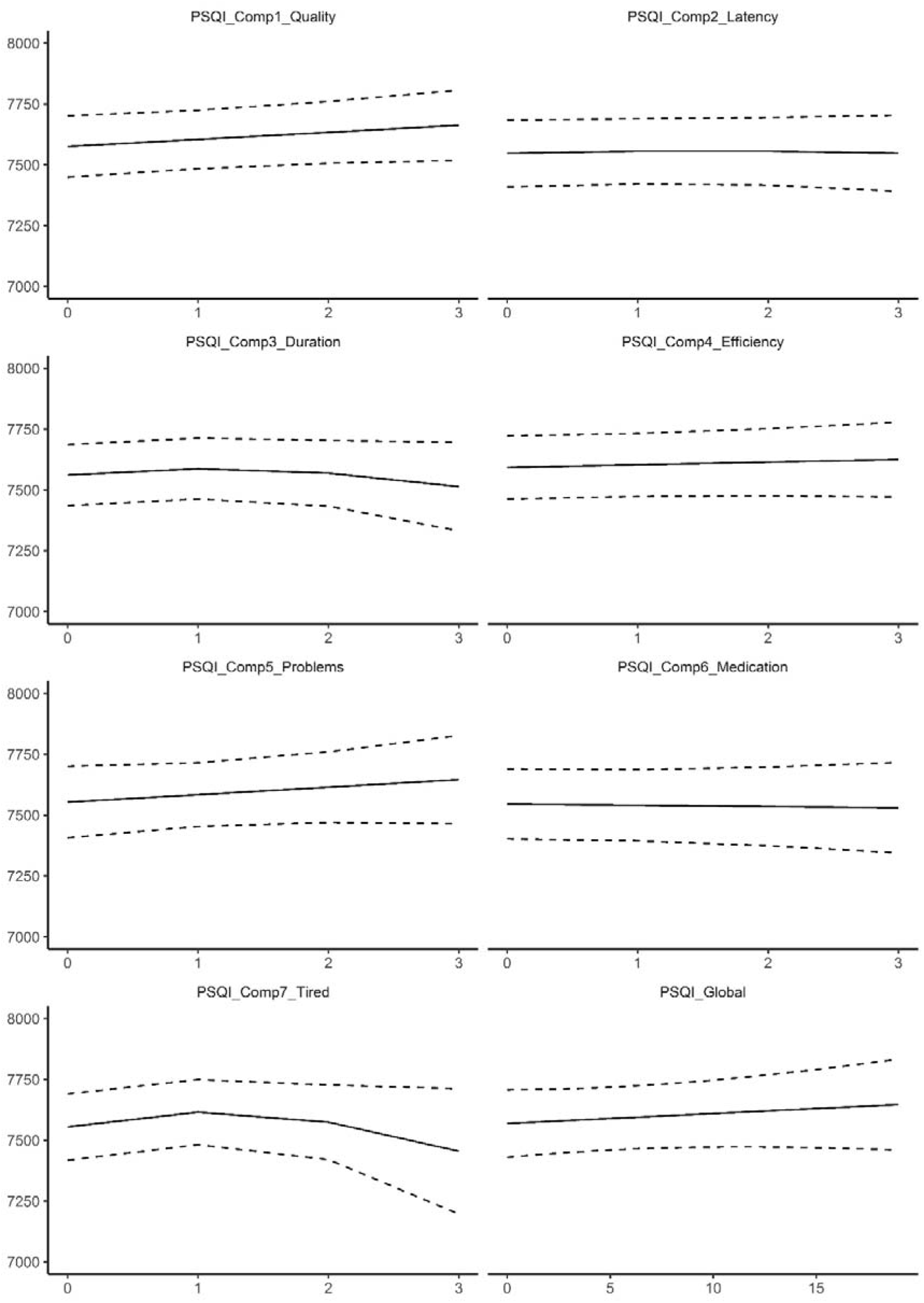
Relationships between self-reported sleep and hippocampal volume in Lifebrain. GAMs were used to test the cross-sectional relationship between self-reported sleep (x-axis) and hippocampal volume (y-axis). Sex, ICV and study were used as covariates. Dotted lines represent 95% CI. X-axis denotes sleep score, y-axis hippocampal volume in mm^3^.

### Cross-sectional analyses: Replication sample from UK Biobank

Relationships between age and each of the six sleep items from UKB, controlling for sex, are illustrated in Figure 4 (see SI for details). Similar to Lifebrain, the age-relationships for all the sleep items were highly significant. Daytime dozing, frequency of napping during the day, and sleeplessness were positively related to age, as was ease of getting up in the morning. Sleep duration also showed higher values with age, but with a negative age-relationship reported by participants below 55 years of age. Snoring showed an inverse U-shaped age-relationship, with higher prevalence of snoring during mid-life. The sleep items from UKB are not directly comparable to the PSQI items used in Lifebrain. Still, Daytime dozing and Nap during day increased with age, corresponding to the increase in Daytime tiredness in the last part of the age-span in the Lifebrain sample. Further, sleep duration increased with age among the older participants in both UKB and Lifebrain, although the negative relationship with age until the mid-fifties in UKB was not seen in Lifebrain. We tested for main effects of sex. Men reported more daytime dozing (estimate = 0.05, t = 6.98, p = 2.91e^−12^), that it was easier to get up in the morning (estimate = 0.19, t = 19.6, p < 2e^−16^), more daytime napping (estimate = 0.19, t = 23.4, p < 2e^−16^), longer sleep duration (estimate 0.07, t = 4.74, p = 2.21e^−6^), less sleeplessness (estimate = −0.22, t = 22.5, p < 2e^−16^) and more snoring (estimate = 0.16, t = 23.59, p < 2e^−16^). In sum, similar to Lifebrain, sex had a main effect on all self-reported sleep items. However, men scored worse on the UKB items dozing and snoring, which are not identical to any of the PSQI sleep scales used in Lifebrain.

**Figure 4.**
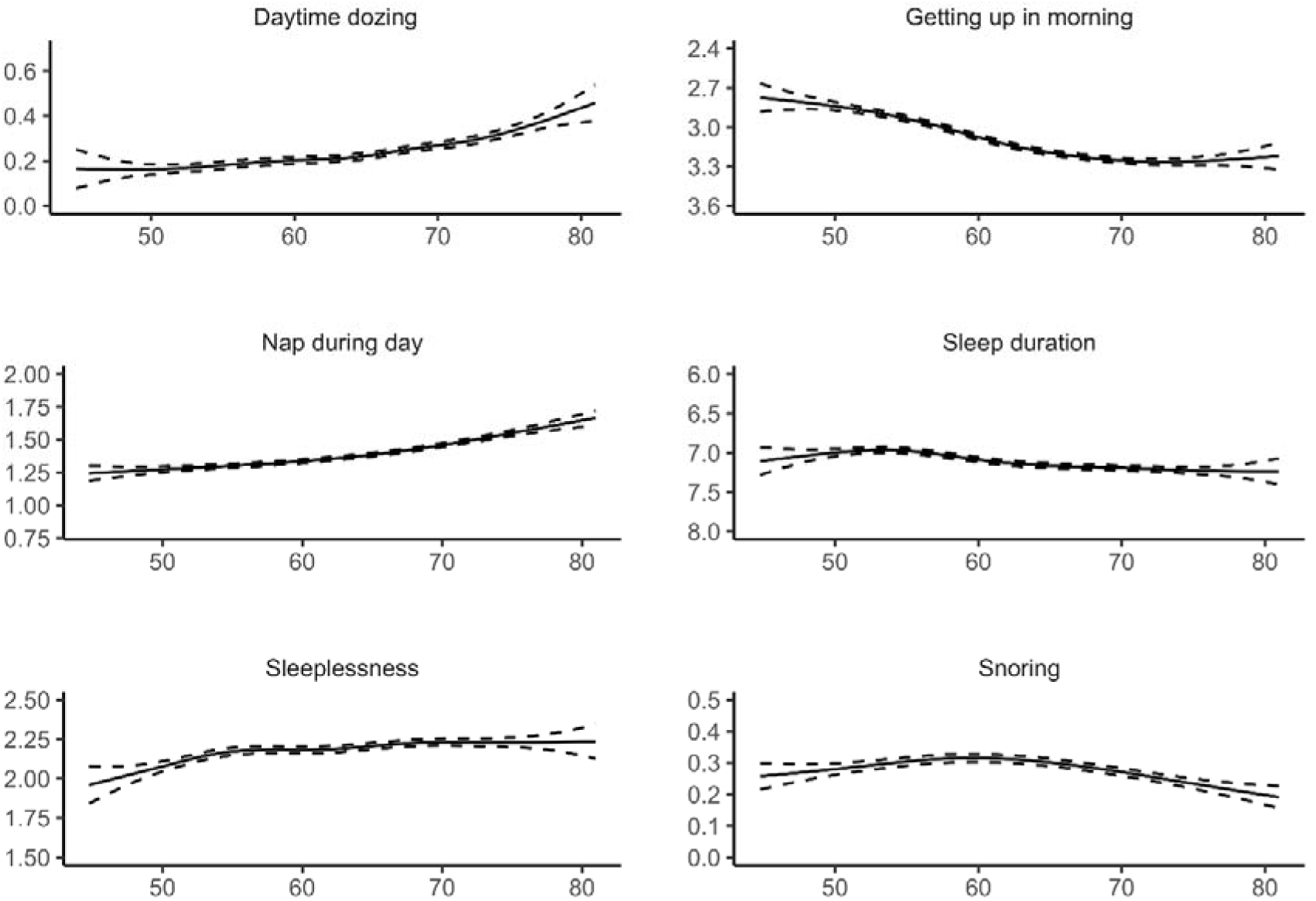
Relationships between age and self-reported sleep in UK Biobank. GAMs were used to obtain age-curves for each sleep variable. Sex was included as covariate in the analyses. Dotted lines represent 95% CI. For all items except “Sleep duration”, high scores mean poor sleep.

The relationship between age and hippocampal volume in UKB was highly significant (edf = 7.26, F = 374.5, p < 2e^−16^), see Figure 5. Relationships between the UKB sleep items and hippocampal volume, controlling for age, sex and ICV, are illustrated in Figure 6. Only sleep duration (p = 7.31e^−6^) was related to hippocampal volume. As values in the extreme ends of reported sleep seemed to be responsible for the relationship, we restricted the data to include sleep duration between 5 and 9 hours only and re-ran the analysis. Still including 20755 observations, the p-value increased to .086. No significant age-interactions were found for any of the sleep items. Thus, the UKB results were in agreement with the lack of meaningful cross-sectional sleep-hippocampal volume relationships in Lifebrain, and aligns with previous findings within a Lifebrain cohort (Cam-CAN) which found no cross-sectional associations between PSQI subcomponents and white matter microstructure (indexed by Fractional anisotropy) across ten tracts [58].

**Figure 5.**
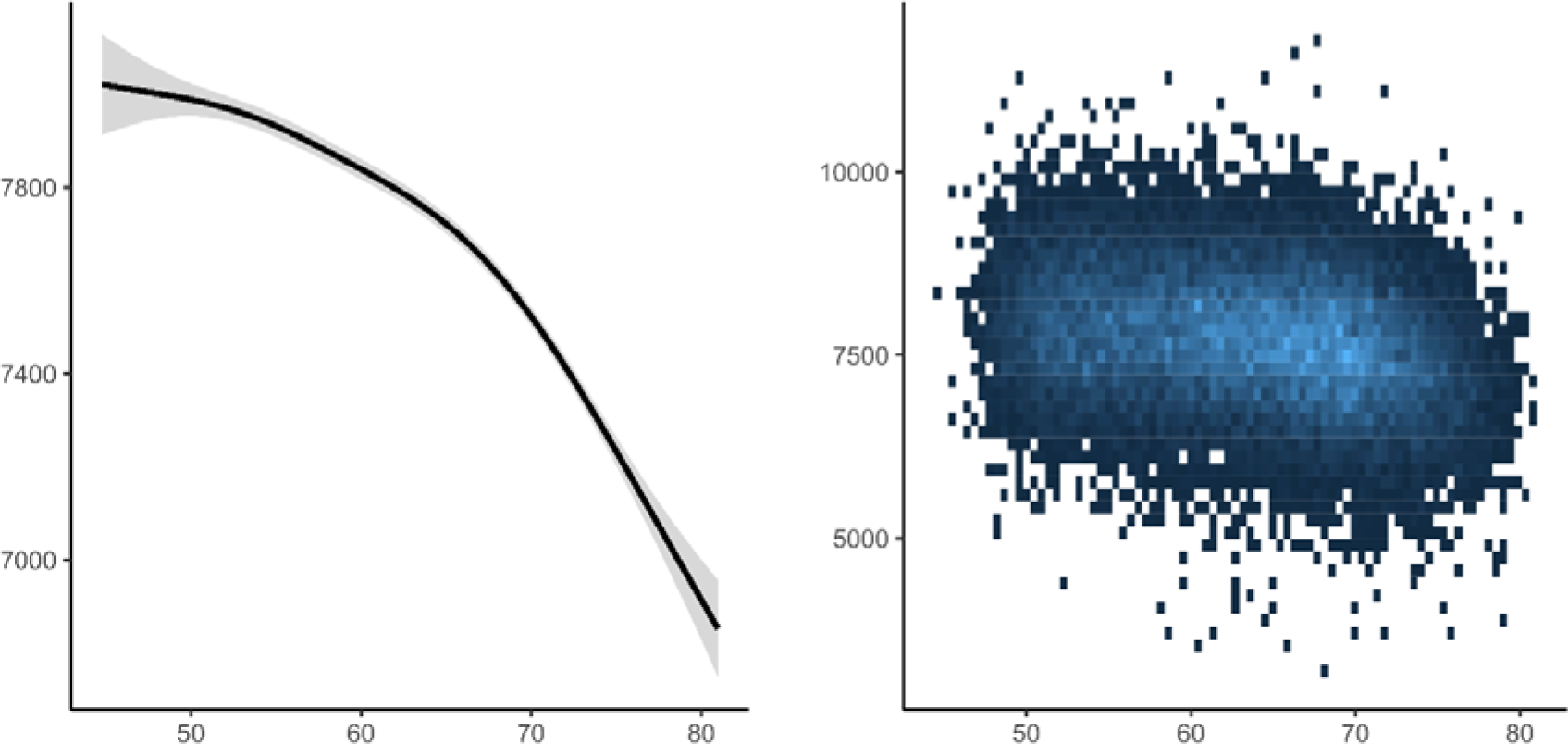
Relationships between age and hippocampal volume in UK Biobank. GAM was used to obtain the age-curve for hippocampal volume, covarying for sex and ICV.

**Figure 6.**
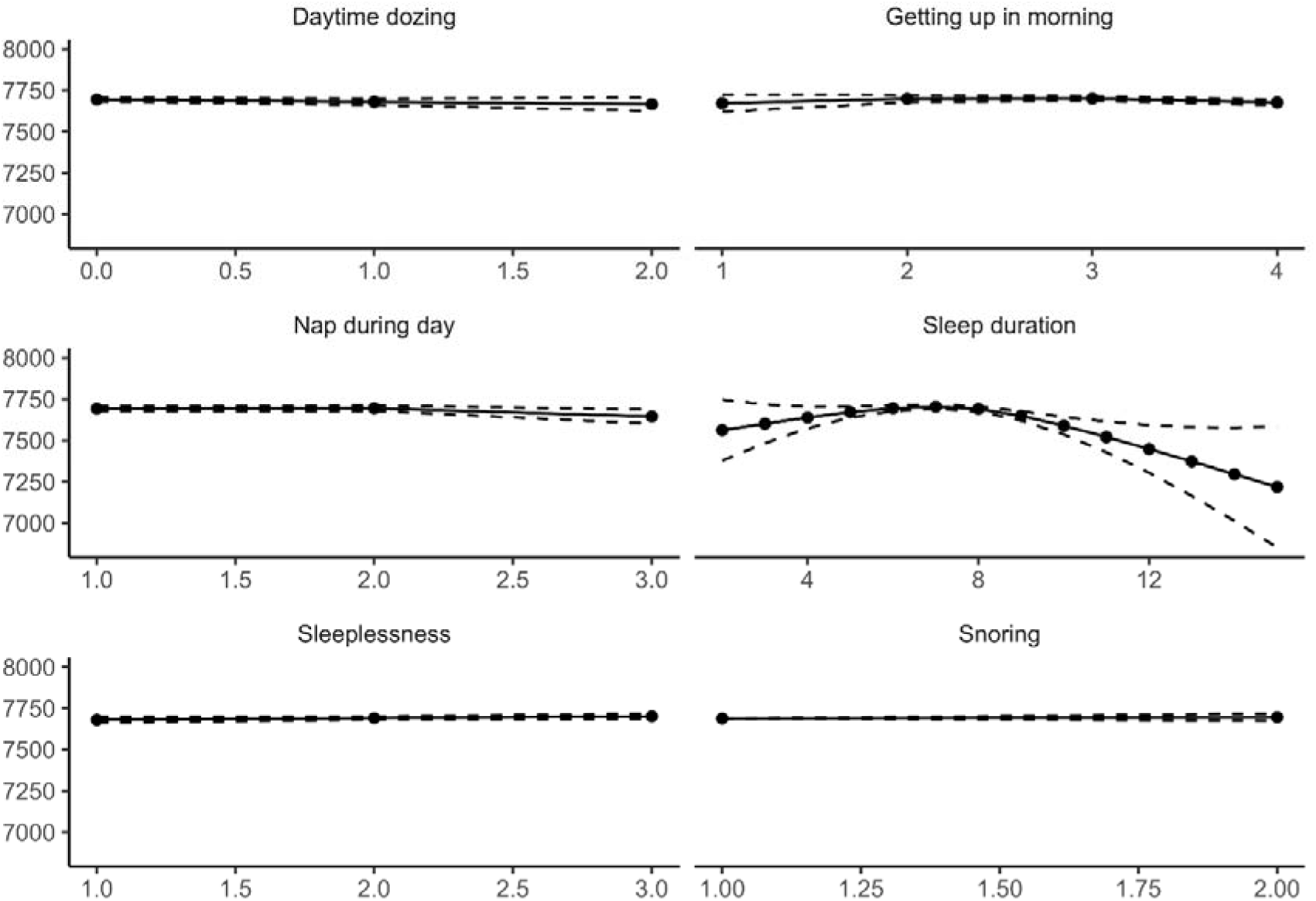
Relationships between self-reported sleep and hippocampal volume in UK Biobank. GAMs were used to test the cross-sectional relationship between self-reported sleep (x-axis) and hippocampal volume (y-axis). Sex and ICV were used as covariates. Dotted lines represent 95% CI.

### Longitudinal analyses: self-reported sleep and hippocampal volume change

We tested if sleep was related to change in hippocampal volume over time, restricting the analyses to participants with at least two MRI examinations. Sleep quality (edf = 4.80, F = 3.69, p = .0019), efficiency (edf = 6.90, F = 6.93, p = 2.36e^−08^), problems (edf = 2.22, F = 5.95, p = .0017) and daytime tiredness (edf = 4.17, F = 8.99, p = 4.13e^−07^) showed significant associations with hippocampal volume change at the α-threshold corrected for eight tests (Figure 7). These relationships were also confirmed by comparing Akaike Information Criterion (AIC) for models with and without the PSQI × time interaction term included (see Table 5). In general, the relationships reflected participants with worse sleep showing more hippocampal volume loss over time. The exception was daytime tiredness, where worse scores were non-linearly associated with volume loss after three years, exceeding those reporting no tiredness in the beginning of the interval before showing less loss in the last part. For sleep quality, only 21 participants reported the highest score. To make sure these did not unduly influence the results we repeated the analysis without these participants. Sleep quality was still significantly related to volume change at an α-threshold of .05, but not at the corrected threshold (edf = 4.03, F = 3.40, p = .008). All analyses were also run testing for interactions with age or sex, with no significant results.

**Figure 7.**
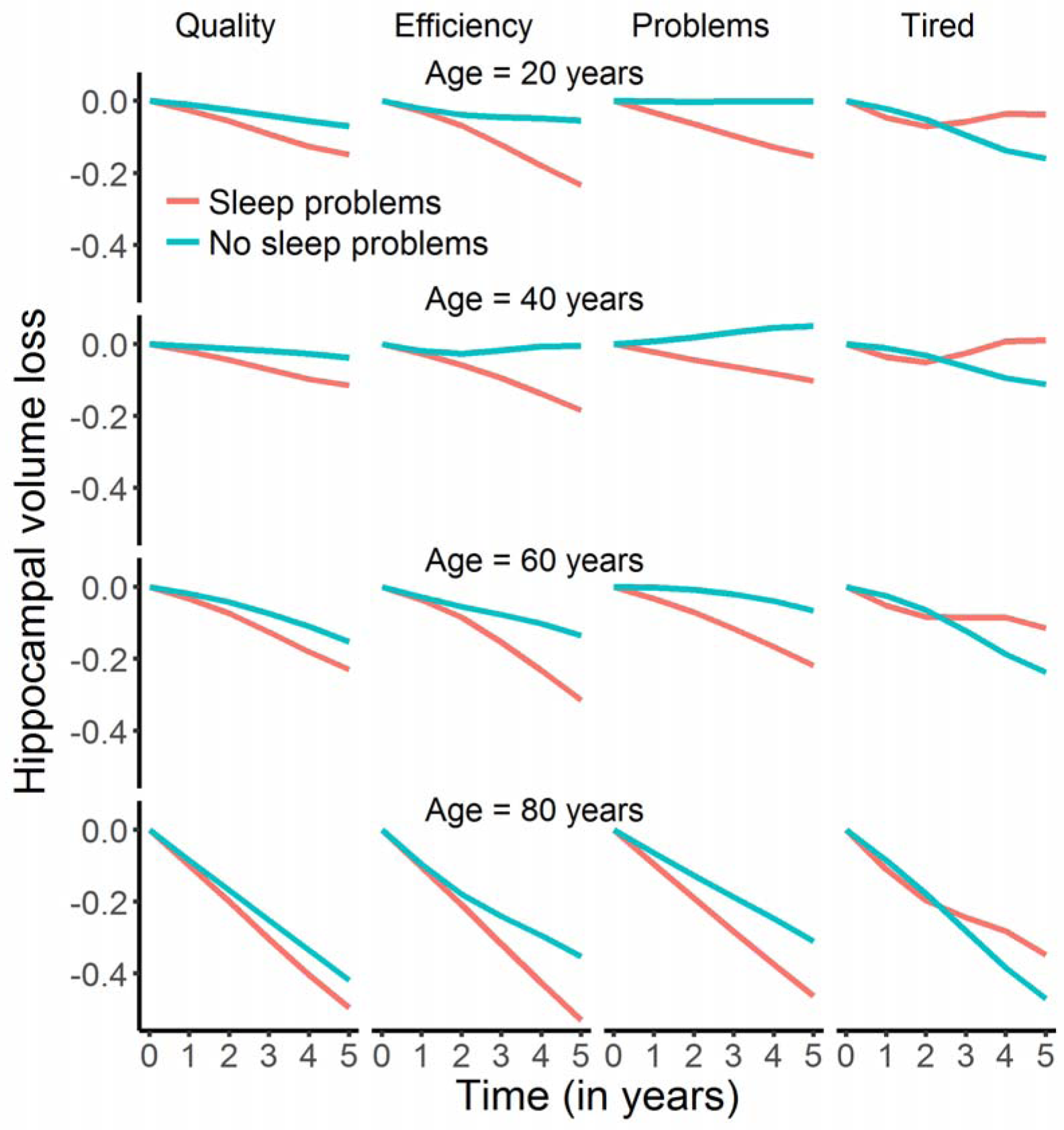
Relationships between self-reported sleep and hippocampal change in Lifebrain. The plots illustrate the relationship between self-reported sleep and hippocampal volume change over time. Only the four significant relationships are shown. The lines depict the hippocampal change trajectories over five years for those with PSQI score = 0 (no problems) or score = 2 (problems). Note that this selection was not used for the statistical analyses and is included to show the nature of the interaction with time only.

**Table 5.** Tests of sleep × time interactions in prediction of hippocampal volume change. AIC: Negative values indicate better model fit for the models including the PSQI × Time interaction term.

|  | PSQI × Time<br>(p-value) | ΔAIC |
| --- | --- | --- |
| Sleep variable |  |  |
| Quality | 0.0019 | -5.6 |
| Latency | 0.98 | 6.0 |
| Duration | 0.039 | 1.72 |
| Efficiency | 0.000000024 | -26.9 |
| Problems | 0.0017 | -6.1 |
| Medication | 0.13 | 3.20 |
| Tired | 0.00000041 | -22.6 |
| Global | 0.070 | 26.3 |

The interactions were further explored to assess effects sizes, as illustrated in Figure 8 (see Supplemental_Information_Interactions for details). We computed the expected annual change in hippocampal volume from 20, 40, 60 and 80 years, depending on whether the score on each of the four PSQI variables showing significant interactions with time was zero or two. Across these items, participants scoring zero had a mean annualized reduction of hippocampal volume of −0.41% compared to −0.63% for those scoring two. As there were no age-interactions, this difference was stable across the four tested ages, i.e. −0.16 vs −0.36% at age 20 years, −0.09 vs. −0.30% at age 40 years, −0.24 vs. −0.47% at age 60 years, and −1.14 vs. −1.39% at age 80 years for those scoring zero vs. two, respectively. These analyses also confirmed than one-year atrophy was higher for participants with high score in daytime tiredness, even though Figure 7 indicates that there is a shift to less atrophy after three years. The latter pattern is difficult to explain, and could reflect noise in the data.

**Figure 8.**
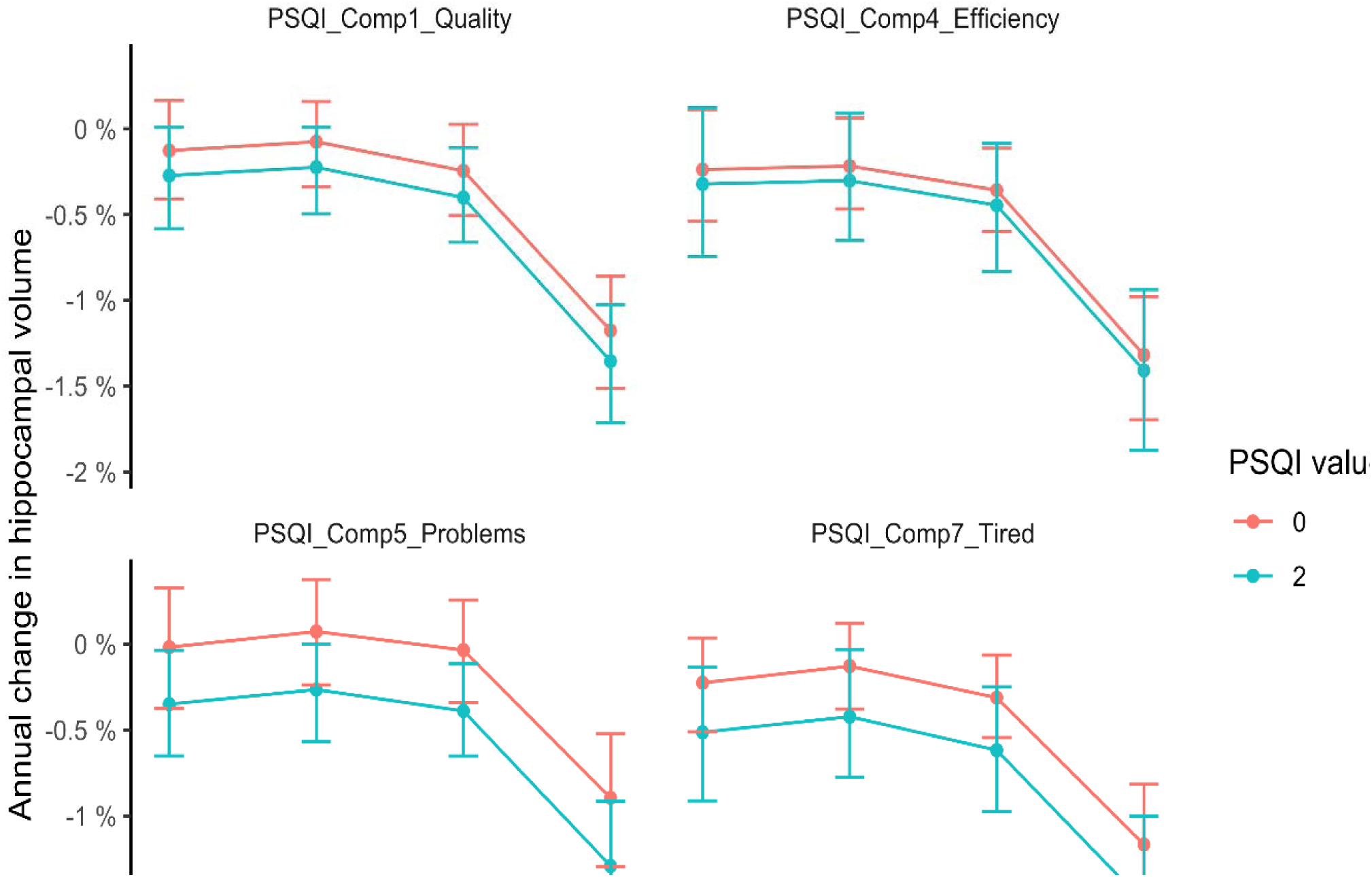
Annual percent change in volume as a function of sleep. Tested at four different ages, annual reduction in hippocampal volume was on average 0.22% greater in participants scoring two compared to zero on the PSQI items quality, efficiency, problems and daytime tiredness. Error bars denote 95% CI.

We re-analyzed the significant time-interactions using BMI and depression scores as covariates. BMI and depression did not contribute significantly and did not affect the sleep-time interactions. The most substantial effect of the additional covariates was that the p-value of sleep problems × time on hippocampal volume increased from 0.002 to 0.017, which probably was a result of lower power due to BMI and depression scores not being available for the full sample (see SI for full results).

Finally, we ran a GAMM with all sleep variables showing significant interactions with time included simultaneously as independent variables. Efficiency (edf = 7.9, F = 3.81, p = .00025), problems (edf = 5.0, F = 3.34, p = .004) and tiredness (edf = 4.6, F = 6.68, p = 1.99e^−5^) were still significantly related to hippocampal volume over time, while sleep quality was not (edf = 1.0, F = 0.15, p = .70).

### Comparison of longitudinal and cross-sectional results

We would expect the relationship between sleep and hippocampal volume change over time to be detected as an interaction between age and sleep on hippocampal volume, i.e. the relationship between sleep and hippocampal volume would be stronger in the older part of the sample. However, the lack of cross-sectional relationships could be caused by individual differences in hippocampal volume being too large compared to the limited longitudinal effects of sleep. We used the effect size from the interaction between sleep efficiency and time to simulate whether we had power to detect this as a cross-sectional age-interaction. The simulations clearly demonstrated that whereas we had excellent power (∼100%) to detect the longitudinal association in our data, the power to detect an age-interaction using the cross-sectional data was only about 10% (Figure 9). This demonstrates that the magnitude of the relationship between sleep and hippocampal change was too small to be detected in our cross-sectional dataset of more than 3000 participants (see SI for details).

**Figure 9.**
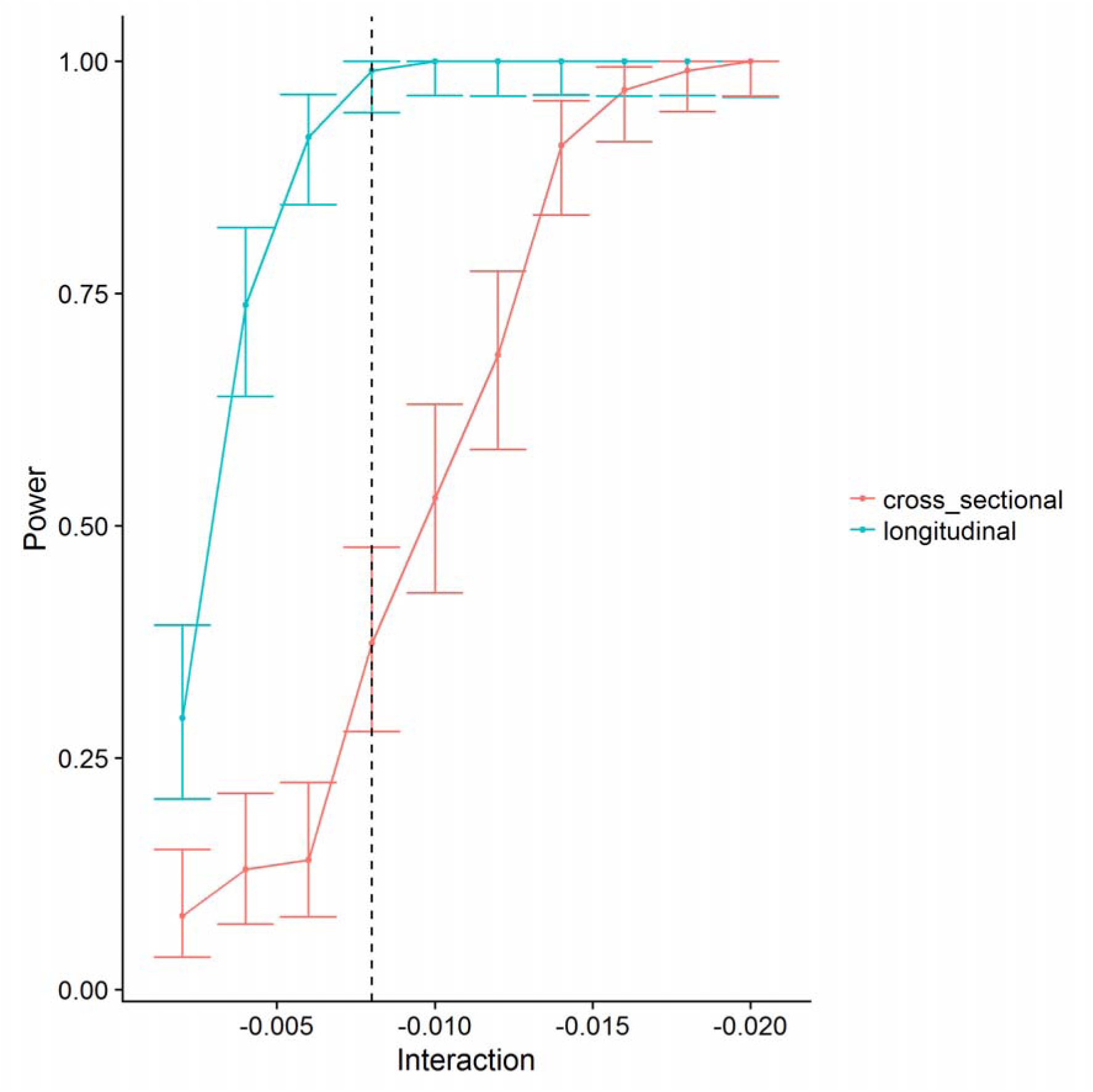
Statistical power. The figure illustrates the superior power of the longitudinal design. The x-axis represents the size of PSQI × time (longitudinal) or PSQI × age interactions (cross-sectional). The y-axis represents statistical power. The dotted vertical line represents the observed effect size of the sleep efficiency × time interaction. As shown, the power to detect this is close to 1 (100%) with the longitudinal design, and very poor with the cross-sectional design.

## Discussion

Worse self-reported sleep was related to greater hippocampal volume reduction over time. This was seen for specific aspects of sleep, i.e. quality, efficiency, problems, and daytime tiredness, where a PSQI score of two was associated with on average 0.22% more volume loss annually than a score of zero. Two caveats need to be considered. First, it is not clear how molecular mechanisms identified in the rodent experiments can be applied to the present findings, as variations in self-reported sleep are very different from experimentally induced sleep deprivation and the effects of specific molecular mechanisms on volume change are speculative. Second, the longitudinal effect sizes were small, and the relationships between sleep and volume were too weak to be identified in the cross-sectional data – both in Lifebrain and UK Biobank. Implications of the findings are discussed below.

### Self-reported sleep and relationship to age

All self-reported sleep measures, in Lifebrain and UKB, were significantly related to age. Whereas sleep duration, efficiency and problems were worse with higher age in Lifebrain, self-reported sleep quality was better with age, and sleep latency and daytime tiredness showed improvement until middle age or longer. The global PSQI score showed little change before 50 years of age, after which worse scores were seen for the rest of the age-span, suggesting PSQI sum score may not be the best measure to use in lifespan cohorts [58]. These patterns fit well with the results from a meta-analysis of polysomnography data [2]. Despite these findings, self-reported sleep quality was higher in older adults, in line with previous research [59].

Although the cross-sectional age-relationships were highly significant, the effects were relatively modest. For instance, sleep duration scores increased from about 0.3 at 20 years to 0.9 at 85. These numbers are based on the PSQI scoring system where 0 represents 7 hours of sleep or more and 1 represents 6-7 hours. Fewer hours of sleep in combination with less daytime tiredness and less sleep efficiency could suggest that older adults sleep less because their sleep needs are less. In support of this view are findings that older adults tend to sleep less despite opportunities to sleep more, they show a smaller rebound in slow wave sleep after sleep deprivation, and experience less daytime sleepiness after slow wave sleep deprivation [3]. However, arguments against this interpretation are that shorter sleep in older age may be due to desensitization to a homeostatic sleep drive, that less subjective sleepiness could be due to re-normalization of the subjective feeling of tiredness over time, and that older adults also perform worse on at least some cognitive tasks after sleep deprivation [3]. Thus, the debate on whether shorter sleep duration in aging is a result of less sleep needs or lower sleeping abilities is not settled [60].

Although the UKB data in general supported the finding of worse self-reported sleep with age, sleep duration seemed to increase through most of this older age-range, i.e. from 55 years. This is in conflict with the observation in Lifebrain. The reasons for this discrepancy are not clear, but it may be noted that the increase in sleep duration over 30 years in UKB is only about a quarter of an hour. Since data are lacking for young participants in UKB, we do not know whether this represents an old-age or a life-long pattern.

We also tested for main effects of sex, and interactions between sex and age. It has been suggested that some sleep mechanisms are differentially affected by age in men and women whereas others may remain equivalent [3, 61]. For instance, sex-specific changes in the circadian alerting signal have been proposed to account for greater daytime nap propensity in older men [3]. We found that although women in general reported worse sleep than men did, in line with previous studies [62], there were no sex-specific age-effects. The above mentioned meta-analysis concluded that the associations between sleep variables and aging were similar across sexes, but that larger effects of age were observed in women for total sleep time and sleep efficiency [2]. In Lifebrain, the lack of sex × age-interactions suggests that self-reported sleep show similar age-trajectories for men and women.

### Self-reported sleep and hippocampal volume change

Low sleep quality and efficiency, more sleep problems, and daytime tiredness were related to more hippocampal volume change. Two previous longitudinal studies with reasonably large samples of 119 [33] and 147 [32] participants reported no significant effects of self-reported sleep on hippocampal change. The statistical power in the current study allowed detection of such effects with high confidence, although the effects sizes were modest. On average, participants scoring two on these four PSQI subscales showed 0.22% more annual volume loss than those scoring zero. This does not mean that the observed relationships between self-reported sleep and hippocampal atrophy are not important. Both sleep [1] and hippocampal volume [63] are substantially affected by chronological age, and both are sensitive to age-related degenerative conditions such as AD [13, 14, 64]. Cognitively healthy older adults with greater initial levels of sleep fragmentation show more rapid rate of cognitive decline and higher risk of developing AD [65]. Still, multiple brain conditions or diseases also affect sleep [3], and it is not possible from the present observational study to infer the direction of causality. There could be a causal relationship where poor sleep contribute to increased hippocampal atrophy. However, hippocampal atrophy could also contribute to worse self-reported sleep. Hippocampal atrophy tends to increase from about 60 years of age, and in line with this, sleep efficiency and daytime tiredness also deteriorated after middle-age. Still, age was accounted for in the sleep-atrophy relationships, and no significant age-interactions were found. Thus, there are no results from the present study to suggest that increased age-related hippocampal atrophy is caused by – or causes – worsening of self-reported sleep in higher age. Rather, the relationship between worse self-reported sleep and hippocampal change seems to be stable across adult life, even in age-ranges with smaller hippocampal volume loss at group level. Future investigations with repeated measures of both brain structure and sleep quality may be able to examine whether neural changes are precursors to worsening sleep, or whether (negative) changes in sleep quality in old age are associated with accelerated grey matter aging.

Previous studies using physiological measures of sleep have found that hippocampal volume may be implicated in some but not all age-related differences in sleep architecture. In particular, it is speculated that impairments in the functional expression of sleep spindles [66] may be caused by age-related atrophy of cell bodies in hippocampus [3], whereas age differences in slow wave activity seem to be independent of hippocampal structure and rather connected to hippocampal function [11, 67]. How these features translate into aspects of self-reported sleep is unclear, and more importantly, none of these studies measured actual volumetric changes from repeated scanning.

Previous cross-sectional studies of patients with different sleep-related conditions [18–27] and older samples without specific sleep problems [29–31] yielded mixed results regarding the relationship between self-reported sleep and hippocampal volume. We found no cross-sectional relationships in Lifebrain or UKB. As shown in the simulation experiment, the longitudinal relationships were too weak to be detected as age-interactions in the cross-sectional analyses given the large inter-individual variation in hippocampal volume. If there was a strong longitudinal relationship between sleep and hippocampal volume change, this could have been detectable as an age-interaction also in our very large cross-sectional samples. This highlights the strength of the longitudinal research design when investigating brain structural changes.

### Limitations

The study has several limitations. First, we used a self-report measure for sleep. The advantage is that sleep is measured in the participants’ natural environment, increasing ecological validity. The disadvantage is that the results reflect self-reported aspects of macro-level sleep architecture, not physiological sleep, and age-related changes in these can be mechanistically distinct from micro-level changes in physiological sleep oscillations [3]. Second, although the vast majority of the participants were screened for cognitive problems, we did not screen the UK Biobank participants, and conservative screening was not performed for all sub-samples in the Lifebrain cohort. Still, the results appear robust and not driven by outliers, so we do not believe sample heterogeneity has affected the outcome. Third, since we lacked adequate longitudinal observations of the sleep variables for a substantial part of the sample, sleep was studied as a trait. This prevented us from addressing dynamic changes in sleep within individuals, which would have been very interesting to relate to hippocampal volume change. Finally, we only tested the relationship with hippocampal atrophy. There are reasons to expect relationships between sleep and other brain regions [3, 32], or white matter structure, which will be topic of later studies.

### Conclusion

The present study showed that specific aspects of self-reported sleep – quality, efficiency, problems, and daytime tiredness – are related to increased hippocampal volume loss over time. In the largest study to date combining data from multiple cohorts, we observe modest longitudinal effects and negligible to absent cross-sectional effects. Together these findings contribute to our understanding of hippocampal volume loss across the adult lifespan.

## Acknowledgement

The Lifebrain project is funded by the EU Horizon 2020 Grant: ‘Healthy minds 0–100 years: Optimising the use of European brain imaging cohorts (“Lifebrain”)’. Grant agreement number: 732592. Call: Societal challenges: Health, demographic change and well-being. In addition, the different sub-studies are supported by different sources:

LCBC: The European Research Council’s Starting/ Consolidator Grant schemes under grant agreements 283634, 725025 (to A.M.F.) and 313440 (to K.B.W.), as well as the Norwegian Research Council (to A.M.F., K.B.W.), The National Association for Public Health’s dementia research program, Norway (to A.M.F) and the Medical Student Research Program at the University of Oslo. Betula: a scholar grant from the Knut and Alice Wallenberg (KAW) foundation to L.N. Barcelona: Partially supported by a Spanish Ministry of Economy and Competitiveness (MINECO) grant to D-BF [grant number PSI2015-64227-R (AEI/FEDER, UE)]; by the Walnuts and Healthy Aging study (http://www.clinicaltrials.gov; Grant NCT01634841) funded by the California Walnut Commission, Sacramento, California. BASE-II has been supported by the German Federal Ministry of Education and Research under grant numbers 16SV5537/16SV5837/16SV5538/16SV5536K/01UW0808/01UW0706, and S.K. has received support from the European Research Council under grant agreement 677804.

Work on the Whitehall II Imaging Substudy was mainly funded by Lifelong Health and Well-being Programme Grant G1001354 from the UK Medical Research Council (“Predicting MRI Abnormalities with Longitudinal Data of the Whitehall II Substudy”) to Dr Ebmeier. The Wellcome Centre for Integrative Neuroimaging is supported by core funding from award 203139/Z/16/Z from the Wellcome Trust. Drs Suri and Zsoldos were funded by award 1117747 from the HDH Wills 1965 Charitable Trust). Dr Suri is now funded by the UK Alzheimer’s Society; CES is supported by the NIHR Oxford Health Biomedical Research Centre. Part of the research was conducted using the UK Biobank resource under application number 32048.

## References

1. Scullin, M.K. and D.L. Bliwise, Sleep, cognition, and normal aging: integrating a half century of multidisciplinary research. Perspect Psychol Sci, 2015. 10(1): p. 97–137.

2. Ohayon, M.M., et al., Meta-analysis of quantitative sleep parameters from childhood to old age in healthy individuals: developing normative sleep values across the human lifespan. Sleep, 2004. 27(7): p. 1255–73.

3. Mander, B.A., J.R. Winer, and M.P. Walker, Sleep and Human Aging. Neuron, 2017. 94(1): p. 19–36.

4. Shi, L., et al., Sleep disturbances increase the risk of dementia: A systematic review and meta-analysis. Sleep Med Rev, 2018. 40: p. 4–16.

5. Hatfield, C.F., et al., Disrupted daily activity/rest cycles in relation to daily cortisol rhythms of home-dwelling patients with early Alzheimer’s dementia. Brain, 2004. 127(Pt 5): p. 1061–74.

6. Videnovic, A., et al., ’The clocks that time us’--circadian rhythms in neurodegenerative disorders. Nat Rev Neurol, 2014. 10(12): p. 683–93.

7. Prinz, P.N., et al., Sleep, EEG and mental function changes in senile dementia of the Alzheimer’s type. Neurobiol Aging, 1982. 3(4): p. 361–70.

8. Irwin, M.R. and M.V. Vitiello, Implications of sleep disturbance and inflammation for Alzheimer’s disease dementia. Lancet Neurol, 2019.

9. Mander, B.A., et al., Sleep: A Novel Mechanistic Pathway, Biomarker, and Treatment Target in the Pathology of Alzheimer’s Disease? Trends Neurosci, 2016. 39(8): p. 552–566.

10. Krause, A.J., et al., The sleep-deprived human brain. Nat Rev Neurosci, 2017. 18(7): p. 404–418.

11. Mander, B.A., et al., Impaired prefrontal sleep spindle regulation of hippocampal-dependent learning in older adults. Cereb Cortex, 2014. 24(12): p. 3301–9.

12. Fjell, A.M., et al., Critical ages in the life course of the adult brain: nonlinear subcortical aging. Neurobiol Aging, 2013. 34(10): p. 2239–47.

13. Holland, D., et al., Subregional neuroanatomical change as a biomarker for Alzheimer’s disease. Proc Natl Acad Sci U S A, 2009. 106(49): p. 20954–9.

14. Braskie, M.N. and P.M. Thompson, A focus on structural brain imaging in the Alzheimer’s disease neuroimaging initiative. Biol Psychiatry, 2014. 75(7): p. 527–33.

15. Raven, F., et al., The role of sleep in regulating structural plasticity and synaptic strength: Implications for memory and cognitive function. Sleep Med Rev, 2018. 39: p. 3–11.

16. Kreutzmann, J.C., et al., Sleep deprivation and hippocampal vulnerability: changes in neuronal plasticity, neurogenesis and cognitive function. Neuroscience, 2015. 309: p. 173–90.

17. Novati, A., et al., Chronic sleep restriction causes a decrease in hippocampal volume in adolescent rats, which is not explained by changes in glucocorticoid levels or neurogenesis. Neuroscience, 2011. 190: p. 145–55.

18. Riemann, D., et al., Chronic insomnia and MRI-measured hippocampal volumes: a pilot study. Sleep, 2007. 30(8): p. 955–8.

19. Joo, E.Y., et al., Brain Gray Matter Deficits in Patients with Chronic Primary Insomnia. Sleep, 2013. 36(7): p. 999–1007.

20. Morrell, M.J., et al., Changes in brain morphology associated with obstructive sleep apnea. Sleep Med, 2003. 4(5): p. 451–4.

21. Dusak, A., et al., Correlation between hippocampal volume and excessive daytime sleepiness in obstructive sleep apnea syndrome. Eur Rev Med Pharmacol Sci, 2013. 17(9): p. 1198–204.

22. Joo, E.Y., et al., Hippocampal volume and memory in narcoleptics with cataplexy. Sleep Med, 2012. 13(4): p. 396–401.

23. Neylan, T.C., et al., Insomnia severity is associated with a decreased volume of the CA3/dentate gyrus hippocampal subfield. Biol Psychiatry, 2010. 68(5): p. 494–6.

24. Noh, H.J., et al., The relationship between hippocampal volume and cognition in patients with chronic primary insomnia. J Clin Neurol, 2012. 8(2): p. 130–8.

25. Spiegelhalder, K., et al., Insomnia does not appear to be associated with substantial structural brain changes. Sleep, 2013. 36(5): p. 731–7.

26. Winkelman, J.W., et al., Lack of hippocampal volume differences in primary insomnia and good sleeper controls: an MRI volumetric study at 3 Tesla. Sleep Med, 2010. 11(6): p. 576–82.

27. Morrell, M.J., et al., Changes in brain morphology in patients with obstructive sleep apnoea. Thorax, 2010. 65(10): p. 908–14.

28. Rosenzweig, I., et al., Hippocampal hypertrophy and sleep apnea: a role for the ischemic preconditioning? PLoS One, 2013. 8(12): p. e83173.

29. Alperin, N., et al., Effect of Sleep Quality on aMCI Vulnerable Brain Regions in Cognitively Normal Elderly Individuals. Sleep, 2018.

30. Carvalho, D.Z., et al., Excessive daytime sleepiness and fatigue may indicate accelerated brain aging in cognitively normal late middle-aged and older adults. Sleep Med, 2017. 32: p. 236–243.

31. Sabeti, S., et al., Sleep, hippocampal volume, and cognition in adults over 90 years old. Aging Clin Exp Res, 2018. 30(11): p. 1307–1318.

32. Sexton, C.E., et al., Poor sleep quality is associated with increased cortical atrophy in community-dwelling adults. Neurology, 2014. 83(11): p. 967–73.

33. Lo, J.C., et al., Sleep duration and age-related changes in brain structure and cognitive performance. Sleep, 2014. 37(7): p. 1171–8.

34. Keage, H.A.D., Sleep and cognitive aging: emerging bedfellows: Editorial for Carvalho, et al. Sleep Med, 2017. 32: p. 244–245.

35. Buysse, D.J., et al., The Pittsburgh Sleep Quality Index: a new instrument for psychiatric practice and research. Psychiatry Res, 1989. 28(2): p. 193–213.

36. Fischl, B., et al., Whole brain segmentation: automated labeling of neuroanatomical structures in the human brain. Neuron, 2002. 33(3): p. 341–55.

37. Walhovd, K.B., et al., Healthy minds 0-100 years: Optimising the use of European brain imaging cohorts (“Lifebrain”). Eur Psychiatry, 2018. 50: p. 47–56.

38. Bertram, L., et al., Cohort profile: The Berlin Aging Study II (BASE-II). Int J Epidemiol, 2014. 43(3): p. 703–12.

39. Gerstorf, D., et al., Editorial. Gerontology, 2016. 62(3): p. 311–5.

40. Nilsson, L.G., et al., The Betula prospective cohort study: Memory, health, and aging. Aging, Neuropsychology and Cognition, 1997. 4: p. 1–32.

41. Shafto, M.A., et al., The Cambridge Centre for Ageing and Neuroscience (Cam-CAN) study protocol: a cross-sectional, lifespan, multidisciplinary examination of healthy cognitive ageing. BMC Neurol, 2014. 14: p. 204.

42. Walhovd, K.B., et al., Neurodevelopmental origins of lifespan changes in brain and cognition. Proc Natl Acad Sci U S A, 2016. 113(33): p. 9357–62.

43. Fjell, A.M., et al., Neuroinflammation and Tau Interact with Amyloid in Predicting Sleep Problems in Aging Independently of Atrophy. Cereb Cortex, 2018. 28(8): p. 2775–2785.

44. Filippini, N., et al., Study protocol: The Whitehall II imaging sub-study. BMC Psychiatry, 2014. 14: p. 159.

45. Abellaneda-Perez, K., et al., Age-related differences in default-mode network connectivity in response to intermittent theta-burst stimulation and its relationships with maintained cognition and brain integrity in healthy aging. Neuroimage, 2019. 188: p. 794–806.

46. Rajaram, S., et al., The Walnuts and Healthy Aging Study (WAHA): Protocol for a Nutritional Intervention Trial with Walnuts on Brain Aging. Front Aging Neurosci, 2016. 8: p. 333.

47. Vidal-Pineiro, D., et al., Task-dependent activity and connectivity predict episodic memory network-based responses to brain stimulation in healthy aging. Brain Stimul, 2014. 7(2): p. 287–96.

48. Westerlund, A., et al., Using the Karolinska Sleep Questionnaire to identify obstructive sleep apnea syndrome in a sleep clinic population. Clin Respir J, 2014. 8(4): p. 444–54.

49. Nordin, M., T. Åkerstedt, and S. Nordin, Psychometric evaluation and normative data for the Karolinska Sleep Questionnaire. Sleep and Biological Rhythms, 2013. 11: p. 216–226.

50. Pillai, V., T. Roth, and C.L. Drake, The nature of stable insomnia phenotypes. Sleep, 2015. 38(1): p. 127–38.

51. Dale, A.M., B. Fischl, and M.I. Sereno, Cortical surface-based analysis. I. Segmentation and surface reconstruction. Neuroimage, 1999. 9(2): p. 179–94.

52. Reuter, M., et al., Within-subject template estimation for unbiased longitudinal image analysis. Neuroimage, 2012. 61(4): p. 1402–18.

53. Jovicich, J., et al., Brain morphometry reproducibility in multi-center 3T MRI studies: a comparison of cross-sectional and longitudinal segmentations. Neuroimage, 2013. 83: p. 472–84.

54. Miller, K.L., et al., Multimodal population brain imaging in the UK Biobank prospective epidemiological study. Nat Neurosci, 2016. 19(11): p. 1523–1536.

55. Alfaro-Almagro, F., et al., Image processing and Quality Control for the first 10,000 brain imaging datasets from UK Biobank. Neuroimage, 2018. 166: p. 400–424.

56. Team, R.C., R: A language and environment for statistical computing. R Foundation for Statistical Computing, Vienna, Austria. URL https://www.R-project.org/. 2018.

57. Wood, S.N., Generalized Additive Models: An Introduction with R. 2006: Chapman and Hall/CRC.

58. Gadie, A., et al., How are age-related differences in sleep quality associated with health outcomes? An epidemiological investigation in a UK cohort of 2406 adults. BMJ Open, 2017. 7(7): p. e014920.

59. Luca, G., et al., Age and gender variations of sleep in subjects without sleep disorders. Ann Med, 2015. 47(6): p. 482–91.

60. Scullin, M.K., Do Older Adults Need Sleep? A Review of Neuroimaging, Sleep, and Aging Studies. Curr Sleep Med Rep, 2017. 3(3): p. 204–214.

61. Redline, S., et al., The effects of age, sex, ethnicity, and sleep-disordered breathing on sleep architecture. Arch Intern Med, 2004. 164(4): p. 406–18.

62. Smagula, S.F., et al., Risk factors for sleep disturbances in older adults: Evidence from prospective studies. Sleep Med Rev, 2016. 25: p. 21–30.

63. Walhovd, K.B., et al., Consistent neuroanatomical age-related volume differences across multiple samples. Neurobiol Aging, 2011. 32(5): p. 916–32.

64. Peter-Derex, L., et al., Sleep and Alzheimer’s disease. Sleep Med Rev, 2015. 19: p. 29–38.

65. Lim, A.S., et al., Sleep Fragmentation and the Risk of Incident Alzheimer’s Disease and Cognitive Decline in Older Persons. Sleep, 2013. 36(7): p. 1027–1032.

66. Fogel, S., et al., Sleep spindles: a physiological marker of age-related changes in gray matter in brain regions supporting motor skill memory consolidation. Neurobiol Aging, 2017. 49: p. 154–164.

67. Mander, B.A., et al., Prefrontal atrophy, disrupted NREM slow waves and impaired hippocampal-dependent memory in aging. Nat Neurosci, 2013. 16(3): p. 357–64.

