## Supplementary material for "Self-reported sleep relates to hippocampal atrophy across the adult lifespan – results from the Lifebrain consortium": SI

**for**

**1.0 General Descriptions**

**2.0 Descriptive Statistics**

**2.1 Number of Observations**

**2.2 Age Distribution**

**2.3 Distribution of Hippocampal Volumes**

**2.4 Distribution of Intracranial Volumes**

**2.5 Details on Self-reported Sleep**

**2.6 Conversion from KSQ to PSQI**

**3.0 References**

**1.0 General Descriptions**

The Lifebrain sample was derived from major European brain studies. The main features of each samples, as well as key references, are provided below.

**LCBC**

*Sample source*

Center for Lifespan Changes in Brain and Cognition

*General description of study/ procedures*

Cognitively normal participants were drawn from studies coordinated by the Research Group for Lifespan Changes in Brain and Cognition (LCBC [www.oslobrains.no](http://www.oslobrains.no)), approved by a Norwegian Regional Committee for Medical and Health Research Ethics. Written informed consent was obtained from all participants.

*Recruitment*

Newspaper adds, web page adds

*Population*

The major part of the sample (n=811) consisted of normal, cognitively healthy participants across the lifespan. One sub-population (n = 103) consisted of patients scheduled for elective gynecological (genital prolapse), urological (benign prostate hyperplasia, prostate cancer, or bladder tumor/cancer) or orthopedic (knee or hip replacement) surgery in spinal anesthesia, turning 65 years or older the year of inclusion.

*Inclusion/ exclusion criteria, screening*

Adult participants were screened using a standardized health interview prior to inclusion in the study. Participants with a history of self- or parent-reported neurological or psychiatric conditions, including clinically significant stroke, serious head injury, untreated hypertension, diabetes, and use of psychoactive drugs within the last two years, were excluded. Further, participants reporting worries concerning their cognitive status, including memory function, were excluded. All participants 40-80 years of age were required to score ≥26 and participants > 80 years ≥ 25 on the Mini Mental State Examination [1] according to population norms [2]. From the sub-population of elective surgery patients, dementia, previous stroke with sequela, Parkinson's disease, and other neurodegenerative diseases likely to affect cognitive function were initial exclusion criteria. From this pool of participants, we further selected only cognitively healthy participants based on clinical examinations at Department of Geriatric Medicine at Oslo University Hospital.

*Key references*

Langnes E, Sneve MH, Sederevicius D, Amlien IK, Walhovd KB, Fjell AM. Lifespan trajectories and relationships to memory of the macro- and microstructure of the anterior and posterior hippocampus – a longitudinal multi-modal imaging study. bioRxiv. doi: <https://doi.org/10.1101/564732>

Fjell AM, Idland AV, Sala-Llonch R, Watne LO, Borza T, Brækhus A, Lona T, Zeterberg H, Blennow K, Wyller TB, Walhovd KB. Neuroinflammation and Tau interact with amyloid in predicting sleep problems in aging independently of atrophy. Cerebral Cortex, 2018, 28, 2775-2785.

**BASE-II**

*Sample source*

Berlin Study of Aging-II

*General description of study/ procedures*

The medical exam consisted of a 2-day protocol including a comprehensive anamnesis performed by a physician and involving a wide array of laboratory and functional tests (including PSQI questionnaire). Medical variables were collected about 1 year prior to cognitive testing (mean time difference in years = 1.2 years; SD = 0.80). After completion of the medical examination, participants were invited to two cognitive testing sessions scheduled 1 week apart, and were tested in small groups (e.g. about 6 participants per group) on a comprehensive cognitive battery that covers key cognitive abilities measured by 21 tasks. Each session lasted about 3.5 h. From one session to the next, participants were asked to fill out psychosocial questionnaires related to subjective health and well-being. The different elements of the study were approved by the ethics committees of the Max Planck Institute for Human Development, the Charité University ethics committee and by the ethics committees of DGPs. Participants signed written informed consent and received monetary compensation for their participation in BASE-II and the MRI study. All experiments were performed in accordance with relevant guidelines and regulations.

On average the participants had 14.01 years of education (SD = 2.89) and a body mass index of 26.70 (SD = 3.51). Most of the participants were married and still living together (63%), while 14% were divorced, 4.4% single and 4.1% widowed. None of the participants took any medication that may have affected memory function or had a history of head injuries, medical (e.g., heart attack), neurological (e.g., epilepsy), or psychiatric disorders (e.g., depression).

*Recruitment*

Baseline Sample (TP1):

Participants were community-dwelling older adults recruited from the greater Berlin

metropolitan area through advertisements in newspapers and public areas. Participants were recruited within the Berlin Aging Study II (BASE-II) (for cohort characteristics and additional details, see Bertram et al., 2014; Gerstorf et al., 2016). The baseline sample comprised 1979 participants (the original sample consists of 2200 subjects, but we reduced the sample to those of which we have sleep and/or cognitive information). Of these, 1519 were older adults aged 61–88 years (mean age 71.5, SD 3.89; 793 female), and 460 were younger adults aged 24–40years (mean age 31.1, SD 3.38; 247 female). On average, older participants had 14.59 years of education (SD 3.03), and younger participants 15.53 years (SD 2.47).

MR Sample: After completion the comprehensive cognitive examination of BASE-II, eligible participants were invited to take part in one MRI session within a time window of 2–4 weeks after cognitive testing, consisting of 341 older adults aged 61–82 years (mean age 70.1, SD = 3.89; 131 female) and 103 younger adults (mean age 31.4, SD = 3.7; 39 female).

Longitudinal data: MR scans and cognitive scores were obtained two times (a baseline (TP1): 2012-2013; and follow-up(TP2): 2015/2016 ).The follow-up sample (TP2) consisted of 325 participants (247 older adults, 68 younger adults) that were re-invited from the MR-subsample only. They were invited to one cognitive session, lasting 3,5 hours and another separate MR session consisting of identical measures of the baseline study.

*Population*

Community-dwelling older adults recruited from the greater Berlin metropolitan area

*Inclusion/ exclusion criteria, screening*

None of the participants took medication that might affect memory function, and none had neurological disorders, psychiatric disorders, or a history of head injuries. All participants reported normal or corrected to normal vision, were right-handed, and scored over 27 on the Mini-Mental Status Examination.

*Key references*

Bertram, L., Böckenhoff, A., Demuth, I., Düzel, S., Eckardt, R., Li, S.-C. C., … Steinhagen-Thiessen, E. (2014). Cohort profile: The Berlin Aging Study II (BASE-II). International journal of epidemiology,43(3), 703–12. doi:10.1093/ije/dyt018

Gerstorf, D., Bertram, L., Lindenberger, U., Pawelec, G., Demuth, I., Steinhagen-Thiessen, E., & Wagner, G. G. (2016). Editorial. Gerontology,62(3), 311–5. doi:10.1159/000441495

**BETULA**

*Sample source*

The Betula longitudinal study on aging, memory and dementia

*General description of study/ procedures*

A subset of 376 participants from the longitudinal Betula study (Nilsson et al., 1997) underwent structural and functional MRI in 2009-2010 and 232 returned for a follow-up scan in 2013-2014. The parent samples from which the scanned participants were derived from were originally recruited to the study in 1988, 1993, and 2008 respectively. The study is approved by the relevant ethical review board.

*Recruitment*

Population-based sampling was used for recruitment, detailed detailed recruitment procedures are found in Nilsson et al., 1997. Participation in the neuroimaging study was offered to all participants who had remained in the study and completed cognitive testing at the 5th Betula test wave in 2008-2009.

*Population*

Population-based, healthy middle-aged and older adults

*Inclusion/ exclusion criteria, screening*

Severe visual or auditory handicaps, intellectual or developmental disabilities, suspected dementia, having a mother tongue other than Swedish, MRI contraindications, neurological disorders, or visual/motor deficits that could interfere with fMRI data collection, MMSE <24, brain or head surgery, substantial brain anatomical deviations.

*Key references*

Nilsson, L.-G., Bäckman, L., Erngrund, K., Nyberg, L., Adolfsson, R., Bucht, G., Karlsson, S., Widing, M., Winblad, B., 1997. The Betula prospective cohort study: Memory, health, and aging. Aging, Neuropsychol. Cogn. 4, 1–32. doi:10.1080/13825589708256633

**Whitehall-II**

*Sample source*

The Whitehall II imaging sub-study

*General description of study/ procedures*

The Whitehall II study, starting in 1985, includes 10.308 British civil servants followed over time, which allows exploring factors hypothesized to affect brain health and cognitive aging. MRI was done in Phase 11 of this study, at which time the total number participants was 6035. A random sample willing and able to give informed consent to participant in the imaging sub-study of Whitehall II was included. Ethical approval was granted generically for the “Protocol for non-invasive magnetic resonance investigations in healthy volunteers” (MSD/IDREC/2010/P17.2) by the University of Oxford Central University/ Medical Science Division Interdisciplinary Research Ethics Committee (CUREC/MSD-IDREC), who also approved the specific protocol: “Predicting MRI abnormalities with longitudinal data of the Whitehall II sub-study” (MSD-IDREC-C1-2011-71).

*Recruitment*

Random selection from the Whitehall II study

*Population*

Population-representative older adults (60-85 years)

*Inclusion/ exclusion criteria, screening*

MRI contraindications, unable to travel to Oxford without assistance

*Key references*

Filippini et al., Study protocol: the Whitehall II imaging sub-study. BMC Psychiatry, 2014, 14:159. Doi:10.1186/1471-244X-14-159

**Cam-CAN**

*Sample source*

The Cambridge Centre for Ageing and Neuroscience (Cam-CAN) study

*General description of study/ procedures*

A population-based cohort of 3000 adults aged 18 was recruited to Stage 1 of the project, where they completed an interview including health and lifestyle questions, a core cognitive assessment, and a self-completed questionnaire of lifetime experiences and physical activity. Of those interviewed, ~700 participants aged 18-87 (100 per age decile) continued to Stage 2 where they undergo cognitive testing and provide measures of brain structure and function. A subset of ~250 adults returned for longitudinal follow-up data. The study is conducted in compliance with the Helsinki Declaration, and has been approved by the local ethics committee, Cambridgeshire 2 Research Ethics Committee (reference: 10/H0308/50).

*Recruitment*

Invitation letters based on the patient lists of general practitioners within the Cambridge City area

*Population*

Population-based, adult lifespan (18 years and up), cognitively healthy

*Inclusion/ exclusion criteria, screening*

General exclusion criteria: Term-time residents of colleges and universities, and participants whose Primary Care Physician feel are inappropriate to include.

Exclusion criteria for the MRI part of the study: Not cognitively normal (MMSE ≤ 24, memory defect, consent difficulties), communication difficulties (hearing problems [35db at 1000 Hz], insufficient English language, vision difficulties), medical problems by self-report of diagnosis (dementia diagnosis /Alzheimer’s Disease, Parkinson’s Disease, Motor Neurone disease, Multiple sclerosis, cancer, stroke, encephalitis, meningitis, epilepsy, head injury with serious results [coma, unconscious for >2 hours, skull fracture], recently diagnosed or uncontrolled high blood pressure, possible pregnancy, current psychiatric conditions [bipolar disorder, schizophrenia, psychosis]), mobility problems (restricted mobility which could prevent further participation, inability to walk 10 metres), substance abuse (past or current treatment for drug abuse, current drug usage), MRI/ MEG safety and comfort exclusions.

*Key references*

Shafto et al. The Cambridge Centre for Ageing and Neuroscience (Cam-CAN) study protocol: a cross-sectional, lifespan, multidisciplinary examination of healthy cognitive ageing. BMC Neurology, 2014, 14:204. doi: 10.1186/s12883-014-0204-1

**University of Barcelona**

*Sample source*

Different brain aging studies from the University of Barcelona; the WAHA cohort, CR/ iTBS cohorts, GABA cohort

*General description of study/ procedures*

Healthy middle-aged/ older adults, all gave informed consent, in accordance with the Declaration of Helsinki (1964, last revision 2013). All study procedures were approved by the local Institutional Review Board).

*Recruitment*

WAHA cohort: Eligible participants were recruited via mailing study brochures (LLU) or through the non-profit organization Institute of Aging (BCN), advertisements in the study centers, and word of mouth. Interested individuals attended an informational group meeting, completed a short medical questionnaire and signed the informed consent.

CR/ iTBS cohorts: Healthy volunteers were recruited via the Institute of Aging, Barcelona. Individuals willing to participate were gathered in an informal meeting to tell them about the investigation, which included repetitive transcranial magnetic stimulation (TMS).

GABA cohort: Participants were recruited from the Institute of Aging (Barcelona) and the University of Experience, an initiative by the University of Barcelona for students aged 55 and older, offering special one and two-year degrees.

*Population*

Middle-aged and older, cognitively normal

*Inclusion/ exclusion criteria, screening*

WAHA cohort: Participants were healthy elderly men and women with normal cognitive and visual function at the time of recruitment. Inclusion criteria were age between 63 and 79 years, apparently healthy, and equally willing to be in either of the two groups. Exclusion criteria included inability to undergo neuropsychological testing; morbid obesity (BMI ≥ 40 kg/m2); uncontrolled diabetes (HbA1c > 8%); uncontrolled hypertension (on-treatment blood pressure ≥ 150/100 mmHg); prior stroke, significant head trauma or brain surgery; relevant psychiatric illness; major depression; cognitive deterioration or dementia with a score < 24 on the Mini-Mental State Examination; other neurodegenerative disorders like Parkinson’s disease; advanced AMD or eye-related conditions precluding ophthalmological evaluation; prior chemotherapy; chronic illness with projected shortened lifespan; allergy to walnuts; customary use of fish oil and/or tree nuts (> 2 servings/week) and/or other relevant sources of ALA, such as flaxseed oil or soy lecithin.

CR/iTBS cohort: Eligible participants had a normal cognitive profile with MMSE scores≥24 and performances not below 1.5SD according to normative scores (adjusted for age and education (Peña-Casanova et al., 2009)) on a neuropsychological evaluation that covered the major cognitive domains (including: Verbal memory: Rey auditory verbal learning test; visual memory: Rey-Osterrieth complex figure; Language: Benton naming test; semantic and phonetic fluencies; Frontal/Executive functions: direct and inverse digits, symbol digits modalities test, trail making test, Stroop test, London tower test; Visuospatial: line orientation, and visuoperceptive: Popplereuter’s embedded figures test).

GABA cohort: None of the participants reported a diagnosis of a neurological or psychiatric disorder or any TMS contraindication (Rossi et al., 2009). Inclusion criteria for the older subjects included a normal cognitive profile with mini-mental state examination (MMSE; Folstein et al., 1975) scores of ≥24 and performance scores not more than 1.5 standard deviation (SD) below normative data (adjusted for age and years of education) on any of the administered neuropsychological tests (i.e., they did not fulfill the criteria for mild cognitive impairment (MCI); Petersen and Morris, 2005. The neuropsychological battery included (1) a screening test for dementia, using the MMSE, and an evaluation of: (2) premorbid cognition and intelligence quotient (IQ), using the vocabulary subtest of the Wechsler Adult Intelligence Scale-III (WAIS-III) and National Adult Reading Test (NART); (3) verbal memory, using the Free and Cued Selective Reminding Test (SRT); (4) executive functions, using the phonemic fluency task and Trail Making Test B (TMTB); (5) language, using the semantic fluency task and Boston Naming Test (BNT); and (6) speed of processing, using the Symbol Digit Modalities Test (SDMT).

*Key references*

WAHA cohort: Rajaram S, Valls-Pedret C, Cofán M, Sabaté J, Serra-Mir M, Pérez-Heras AM, Arechiga A, Casaroli-Marano RP, Alforja S, Sala-Vila A, Doménech M, Roth I, Freitas-Simoes TM, Calvo C, López-Illamola A, Haddad E, Bitok E, Kazzi N, Huey L, Fan J, Ros E. The Walnuts and Healthy Aging Study (WAHA): Protocol for a Nutritional Intervention Trial with Walnuts on Brain Aging. Front Aging Neurosci. 2017 Jan 10;8:333.

CR/ iTBS cohorts: Vidal-Piñeiro D, Martin-Trias P, Arenaza-Urquijo EM, Sala-Llonch R, Clemente IC, Mena-Sánchez I, Bargalló N, Falcón C, Pascual-Leone Á, Bartrés-Faz D. Task-dependent activity and connectivity predict episodic memory network-based responses to brain stimulation in healthy aging. Brain Stimul. 2014 Mar-Apr;7(2):287-96.

GABA cohort: Abellaneda-Pérez K, Vaqué-Alcázar L, Vidal-Piñeiro D, Jannati A, Solana E, Bargalló N, Santarnecchi E, Pascual-Leone A, Bartrés-Faz D. Age-related differences in default-mode network connectivity in response to intermitent theta-burst stimulation and its relationships with maintained cognition and brain integrity in healthy aging. Neuroimage. 2018 Nov 22.

**Replication sample: UK Biobank**

*Sample source*

UK Biobank prospective epidemiological imaging study (all available data March 2019)

*General description of study/ procedures*

UK Biobank is a prospective epidemiological resource gathering extensive questionnaires, physical and cognitive measures, and biological samples (including genotyping) in a cohort of 500,000 participants. An imaging extension to the existing UK Biobank study was funded in 2016 to scan 100,000 subjects from the existing cohort, aiming to complete by 2022. Informed consent is obtained from all UK Biobank participants; ethical procedures are controlled by a dedicated Ethics and Guidance Council (http://www.ukbiobank.ac.uk/ethics) that has developed with UK Biobank an Ethics and Governance Framework (given in full at http://www.ukbiobank.ac.uk/wp-content/uploads/2011/05/EGF20082.pdf), with IRB approval also obtained from the North West Multi-center Research Ethics Committee.

*Recruitment*

By invitation from the original UK Biobank cohort

*Population*

Population-based, middle-aged and older adults

*Inclusion/ exclusion criteria, screening*

MRI safety/quality criteria

*Key references*

Miller et al. Multimodal population brain imaging in the UK Biobank prospective epidemiological study. *Nature Neuroscience*, 2016, 19, 1523-1536.

**2.0 Descriptive statistics**

**2.1 Number of Observations**

Number of participants with varying number of repeated MRI measurements, per study.

| n | Barcelona | BASE-II | Betula | Cam-CAN | LCBC | Whitehall-II |
| --- | --- | --- | --- | --- | --- | --- |
| 1 | 106 | 2 | 122 | 384 | 420 | 773 |
| 2 | 1 | 313 | 189 | 263 | 172 | 0 |
| 3 | 38 | 0 | 0 | 0 | 165 | 0 |
| 4 | 0 | 0 | 0 | 0 | 43 | 0 |
| 5 | 0 | 0 | 0 | 0 | 33 | 0 |
| 6 | 0 | 0 | 0 | 0 | 80 | 0 |
| 7 | 0 | 0 | 0 | 0 | 1 | 0 |

**2.2 Age Distribution**

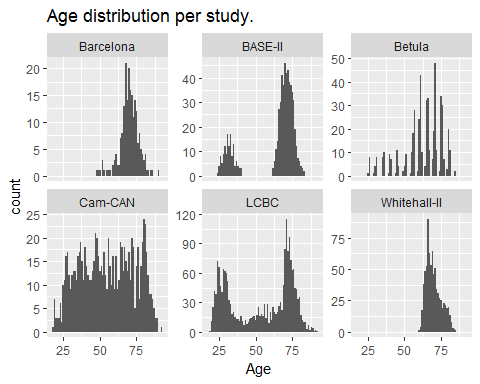

**2.3 Distribution of Hippocampal Volumes**

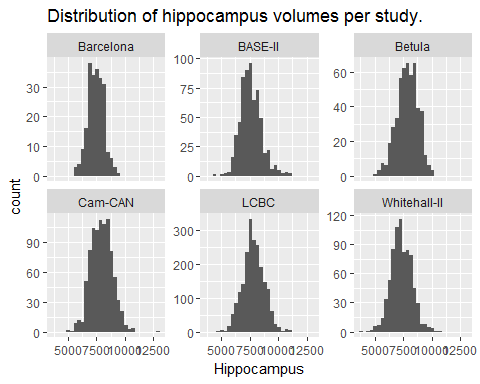

**2.4 Distribution of Intracranial Volumes**

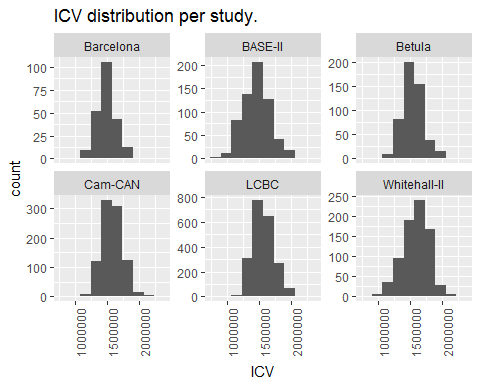

**2.5 Details on Self-reported Sleep**

- Because we average over measurements, some of the LCBC and Betula participants have decimal PSQI scores.
- Betula used Karolinska Sleep Questionnaire (KSQ) instead of PSQI. Hence, we have used the single items of KSQ to compute PSQI scales for the Betula participants, based on the conversion described below.
- Number of missing PSQI variables, after only participants with hippocampus volume is included, is shown in the table below. 100 % missing Component 6 from Betula is by design, as there is no variable in the Karolinska Sleep Questionnaire representing medication.

| key | Barcelona | BASE-II | Betula | Cam-CAN | LCBC | Whitehall-II |
| --- | --- | --- | --- | --- | --- | --- |
| PSQI_Comp1_Quality | 35.3 % | 9.6 % | 0.7 % | 0.4 % | 0.2 % | 0 % |
| PSQI_Comp2_Latency | 1.2 % | 38.5 % | 1.4 % | 4.3 % | 8.6 % | 1.4 % |
| PSQI_Comp3_Duration | 2.4 % | 6.7 % | 6.1 % | 1.8 % | 6 % | 0 % |
| PSQI_Comp4_Efficiency | 3.5 % | 8.7 % | 6.1 % | 6.9 % | 22.9 % | 18.1 % |
| PSQI_Comp5_Problems | 2.4 % | 31.7 % | 0.7 % | 1.1 % | 0.2 % | 0 % |
| PSQI_Comp6_Medication | 35.3 % | 29.8 % | 100 % | 0 % | 0.6 % | 0 % |
| PSQI_Comp7_Tired | 35.3 % | 32.7 % | 2.7 % | 0 % | 1 % | 0.3 % |
| PSQI_Global | 3.5 % | 35.6 % | 0.7 % | 10.5 % | 0.2 % | 0 % |

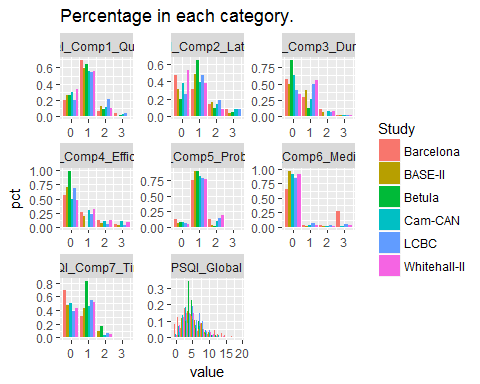

**2.6 Conversion from KSQ to PSQI**

For Betula, Karolinska Sleep Questionnaire was used, and we therefore converted the values from the single items in KSQ to PSQI [3, 4].

### PSQI Component 1

PSQI Component 1 rates the answers to the question (PSQI#6)

- During the past month, how would you rate your sleep quality overall?

The options and values are given in the table below:

| Answer | Value |
| --- | --- |
| Very good | 0 |
| Fairly good | 1 |
| Fairly bad | 2 |
| Very bad | 3 |

The question in KSQ that come closest, is KSQ#5:

- Hur tycker du att du sover på det hela taget? (English: How well do you think you sleep overall?)

The options are “mycket bra”(very well), “ganska bra” (quite well), “varken bra eller dåligt” (neither well nor poorly), “ganska dåligt” (quite poorly), and “mycket dåligt” (very poorly).

Here are the fractions in each option in the Betula study:

```
##
##    1    2    3    4    5
## 0.02 0.11 0.22 0.44 0.20
```

And here are the fractions in each option in PSQI_06 in LCBC:

```
##
##         0         1         2         3
## 0.2730711 0.4712557 0.2072617 0.0484115
```

Looking at the percentages, it seems like the numerical scale used in Betula corresponds to the following:

| Answer | Value |
| --- | --- |
| Mycket bra | 5 |
| Ganska bra | 4 |
| Varken bra eller dåligt | 3 |
| Ganska dåligt | 2 |
| Mycket dåligt | 1 |

We used the following mapping:

| KSQ5 | PSQI_Comp1 |
| --- | --- |
| Mycket bra (5) | 0 |
| Ganska bra (4) | 1 |
| Varken bra eller dåligt (3) | 1 |
| Ganska dåligt (2) | 2 |
| Mycket dåligt (1) | 3 |

Looking at the histograms above, this seems to give a distribution of answers very similar to the other studies.

### PSQI Component 2

PSQI component 2 measures sleep latency, and is computed from the answers to PSQI#2 (number of minutes it takes to fall asleep) and PSQI#5a (how often are you not able to fall asleep within 30 minutes). We used KSQ#9a:

- Har du haft känning av följande besvär de senaste tre månaderna? (…) Svårigheter att somna (Enligsh: Have you experienced the following troubles in the past three months? (…) Trouble falling asleep

The answers to this question are in the variable sleep_07a in the Betula dataset. The possible answers are “Aldrig” (Never), “Någon gång” (Sometimes, “Flera ggr/mån” (Several times/month), “1-2 ggr/vecka” (1-2 times/week), “3-4 ggr/vecka” (3-4 times/week), and “5 ggr eller mer /vecka” (5 times or more/week).

The frequencies of answers are given below:

```
##
##          0          1          2          3          4          5
## 0.16129032 0.51423150 0.15939279 0.08918406 0.04554080 0.03036053
```

The corresponding frequency for PSQI Component 2 in MOAS are given below:

```
##
##          0          1          2          3
## 0.25326170 0.49194167 0.17344589 0.08135073
```

We used the following mapping:

| KSQ9a | PSQI_Comp2 |
| --- | --- |
| Aldrig (0) | 0 |
| Sällan (1) | 1 |
| Flera ggr/mån (2) | 1 |
| 1-2 ggr/vecka (3) | 2 |
| 3-4 ggr/vecka (4) | 3 |
| 5 ggr eller mer /vecka (5) | 3 |

### PSQI Component 3

PSQI Component 3 ranks sleep duration (PSQI#4), using the following scale:

| Answer | Value |
| --- | --- |
| > 7 hours | 0 |
| 6-7 hours | 1 |
| 5-6 hours | 2 |
| < 5 hours | 3 |

This can be computed directly from the Betula data. The following assumptions are used:

- Columns sleep_03a_bedtime sleep_03b_bedtime contain timepoint at which the participant went to bed on weekdays and weekends, respectively.
- Columns sleep_03b_minbef and sleep_03b_minbef contain the number of minutes after bedtime until the participant was asleep on weekdays and weekends, respectively.
- Columns sleep_03a_minbef and sleep_03b_risetime contain the timepoint at which the participant woke up on weekdays and weekends, respectively.

A weighted average between weekdays (a) and weekends (b) is taken, with weights 5/7 and 2/7. If one is missing, we take the non-missing value.

### PSQI Component 4

This component measures sleep efficiency, and can be directly calculated from the KSQ data. In order to keep this an integer, we take the weekday efficiency if it exists, otherwise the weekend efficiency.

### PSQI Component 5

We used the following mapping between PSQI#5b-PSQI#5j and KSQ#9:

| PSQI | KSQ |
| --- | --- |
| 5b) Wake up in the middle of the night or early morning | 9c) Upprepade uppvakanden + 9i) För tidigt uppvakande (Repeated awakenings + Premature awakening) |
| 5c) Have to get up to use the bathroom | No match in KSQ. |
| 5d) Cannot breathe comfortably | 9e) Kippar efter andan, “frustar” under sömnen + 9f) Andningsuppehåll under sömnen (Gasping for breath, «snorting» during sleeping + Pauses in breathing during sleep) |
| 5e) Cough or snore loudly | 9d) Kraftiga egna snarkningar (Severe own snoring) |
| 5f) Feel too cold | No match in KSQ. |
| 5g) Feel to hot | No match in KSQ. |
| 5h) Had bad dreams | 9g) Mardrömmar. (Nightmares) |
| 5i) Have pain | No match in KSQ. |
| 5j) Other reasons | 9j) Störd/orolig sömn (Disrupted/restless sleep) |

These are a total of 7 KSQ questions, matching 9 PSQI questions. We used the same conversion as before:

| KSQ9(c-g,i-j) | PSQI_Comp2 |
| --- | --- |
| Aldrig (0) | 0 |
| Sällan (1) | 1 |
| Flera ggr/mån (2) | 1 |
| 1-2 ggr/vecka (3) | 2 |
| 3-4 ggr/vecka (4) | 3 |
| 5 ggr eller mer /vecka (5) | 3 |

The total score is summed and then multiplied by 7/9 and the component 5 score is assigned using the PSQI Component 5 calculation.

### PSQI Component 6

No question about sleep medication is found in KSQ, so this one is missing.

### PSQI Component 7

This component is based on PSQI#8 and PSQI#9:

**PSQI#8: During the past month, how often have you had trouble staying awake while driving, eating meals, or engaging in social activity?**

- Not during the past month (0)
- Less than once a week (1)
- Once or twice a week (2)
- Three or more times a week (3)

**PSQI#9: During the past month, how much of a problem has it been for you to keep up enough enthusiasm to get things done?**

- No problem at all (0)
- Only a very slight problem (1)
- Somewhat of a problem (2)
- A very big problem (3)

We used the following KSQ questions as proxies for PSQI Component 7:

**Har du haft känning av följande besvär de senaste tre månederna? (English: Have you experienced the following troubles during the last three months?)**

- 9m) Sömnig under arbete (Sleepiness during work)
- 9n) Sömnig under fritid (Sleepiness during spare time)
- 9o) Ofrivilliga sömnperioder (tillnickning) under arbetet (Involuntary sleep periods during work)
- 9p) Ofrivilliga sömnperioder (tillnickning) under fritid (Involuntary sleep periods durign spare time)
- 9q) Behov av att kämpa mot sömnen för att hålla sig vaken? (Need to fight sleepiness to stay awake)
- 9r) Trött i huvudet under dagen (Mental tiredness during the day)

We used the same ranking as before:

| KSQ#9q | PSQI Component 8 |
| --- | --- |
| Aldrig (0) | 0 |
| Sällan (1) | 1 |
| Flera ggr/mån (2) | 1 |
| 1-2 ggr/vecka (3) | 2 |
| 3-4 ggr/vecka (4) | 3 |
| 5 ggr eller mer /vecka (5) | 3 |

The answers are summed, and multiplied by 2/6 to get the same numerical scale as PSQI Component 7.

### PSQI Global

This one is computed for Betula by setting PSQI Component 6 to zero, and summing the others, then multiplying by 7/6 to compensate for the lack of Component 6.

**3. References**

1.	Folstein, M.F., S.E. Folstein, and P.R. McHugh, *"Mini-mental state". A practical method for grading the cognitive state of patients for the clinician.* J Psychiatr Res, 1975. **12**(3): p. 189-98.

2.	Crum, R.M., et al., *Population-based norms for the Mini-Mental State Examination by age and educational level.* JAMA, 1993. **269**(18): p. 2386-91.

3.	Nordin, M., T. Åkerstedt, and S. Nordin, *Psychometric evaluation and normative data for the Karolinska Sleep Questionnaire.* Sleep and Biological Rhythms, 2013. **11**: p. 216-226.

4.	Westerlund, A., et al., *Using the Karolinska Sleep Questionnaire to identify obstructive sleep apnea syndrome in a sleep clinic population.* Clin Respir J, 2014. **8**(4): p. 444-54.
