## Supplementary material for "Self-reported sleep relates to hippocampal atrophy across the adult lifespan – results from the Lifebrain consortium": SI

**for**

**Content**

**1.0 Recruitment of Travelling Brains**

**2.0 Organization of between-site scanning**

**3.0 Preprocessing**

**4.0 Volumetric differences as a function of scanner**

**5.0 Pairwise relationship between volume estimates between scanners**

**1.0 Rationale and background for the Travelling Brains**

The Travelling Brains (TB) sub-project of Lifebrain was initiated with the intent to better understand and possibly correct for the differences in MRI data that are expected to arise as a consequence of the MRI system and the sequence run at the various Lifebrain partner sites. As all the partner sites already had collected data that will be shared and combined within Lifebrain, there was consensus that there should be made an attempt to estimate how the scanners differ and explore possibilities on improving comparisons across scanners. Since MRI data in Lifebrain was derived from eight different scanners, seven participants were recruited to be scanned on seven of the scanners to allow us to estimate between-scanner differences in volumetric estimates and possible biases. The smallest sample from LCBC (n = 101, Avanto scanner) was added at a later point and was thus not included in the Travelling Brains sub-study. Also, the Verio scanner in the Whitehall-II study was not accessible for scanning at the time of the Travelling brains study. Overview of the different scanner models and sequences is provided in Table 3 in the main manuscript.

**2.0 Organization of between-site scanning**

A total of seven people volunteered to travel to each partner site, and to be scanned at every scanner at each site (some sites have multiple scanners). Participants were recruited from the staff of each partner site. Mean participant age was 41.86, (range: 24 - 73), with 4 females. All scanning happened within one month for each person, so that brain change over time would be minimized. At each scanner, participants underwent scanning with protocols either identical or equivalent to the data that site was sharing in Lifebrain, comprising structural T1 scans (in some cases also T2), diffusion weighted scans and resting state BOLD. In addition to being scanned at each scanner, each participant was also scanned twice with the same protocol at the same scanner, with a small break or reposition between the repetitions. This double-procedure was intended to help estimate the within-scanner fluctuations that might occur rapidly. One participants did not complete scanning on one of the three scanners at Oslo. In all, the 7 participants completed 61 sessions across the 11 scanners all together.

**3.0 Preprocessing**

To extract reliable volume and thickness estimates, images were automatically processed with the longitudinal stream [1] in FreeSurfer. Specifically an unbiased within-subject template space and image [2] is created using robust, inverse consistent registration [3]. Several processing steps, such as skull stripping, Talairach transforms, atlas registration as well as spherical surface maps and parcellations are then initialized with common information from the within-subject template, significantly increasing reliability and statistical power [1]. Participants followed-up on different MRI scanners were independently processed for each scanner.

**4.0 Volumetric differences as a function of scanner**

Analysis of variance (ANOVA) was run to test the effects of scanner on hippocampal volume. There was a significant main effect of scanner on hippocampal volume (F = 4.13 [2.1,30], p = .046). The chart below shows the mean hippocampal volume across left and right hemisphere for seven of the scanners used in the present study.

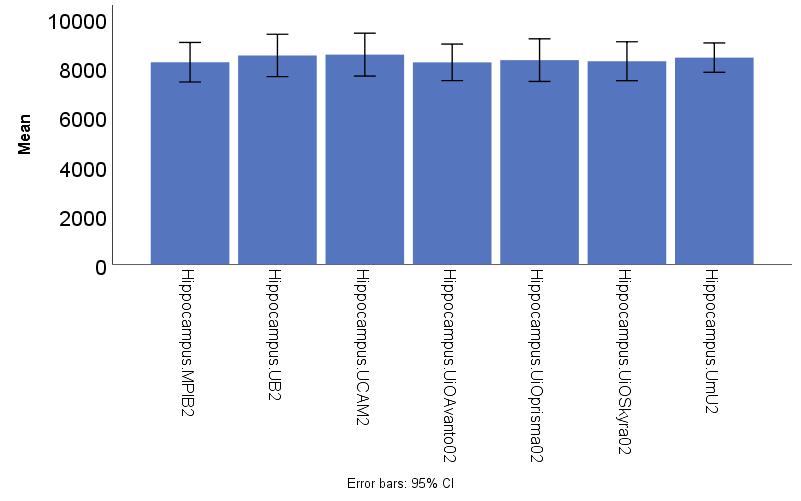

*Mean hippocampal volume across the hemispheres for each scanner. Error bars denotes 95% confidence interval.*

**5.0 Pairwise relationship between volume estimates between scanners**

Despite significant differences in absolute hippocampal volume as a function of scanner, the between-participant rank order was almost perfectly retained between scanners. The mean between-scanner Pearson correlation for bilateral hippocampal volume was r = .98 (range .94-1.00). The figure below shows the pairwise relationship between volume estimates across scanners.

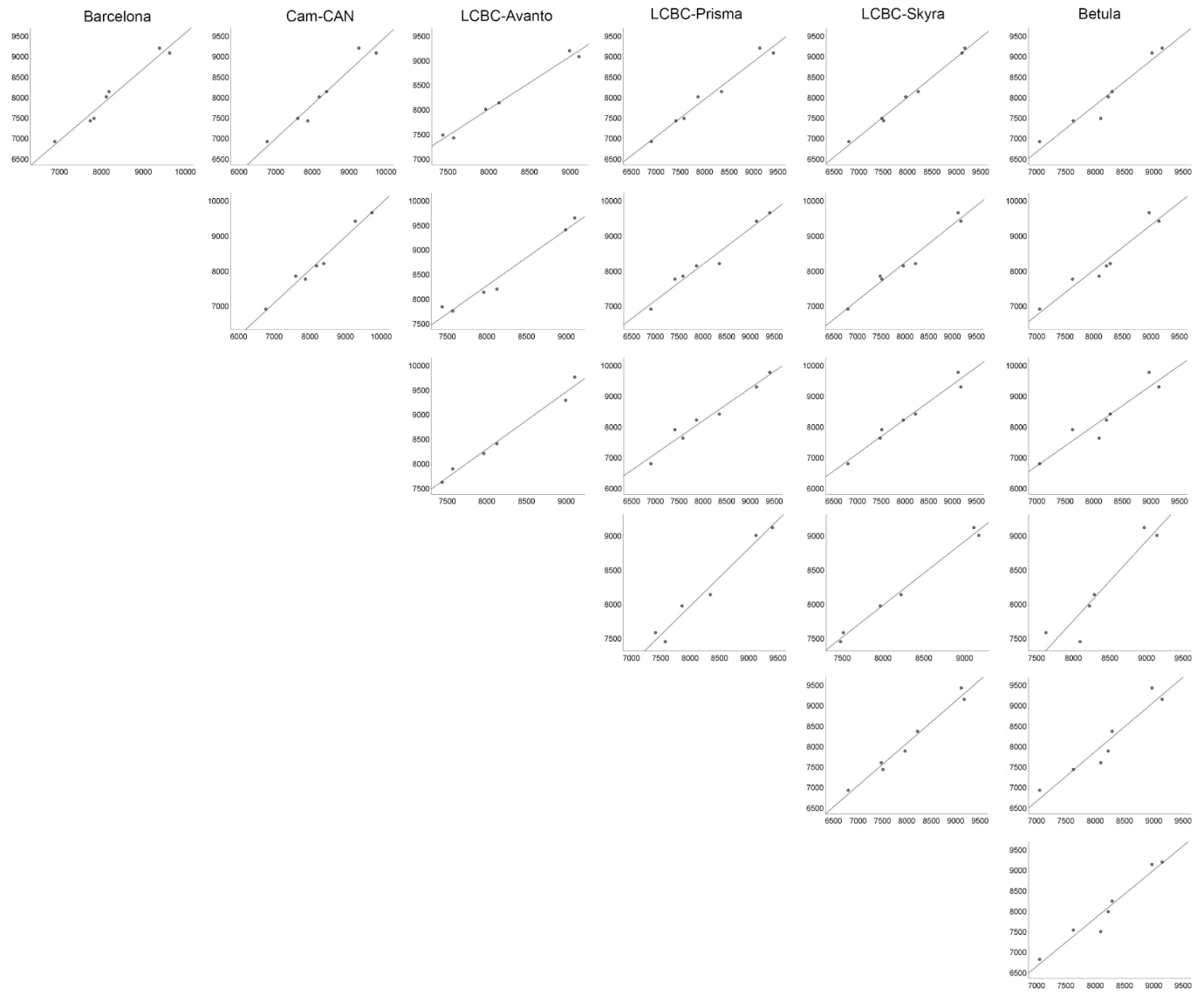

*Scatterplots of the pairwise relationship between the scanners. The first row is the relationship between the BASE-II scanner with the rest of the scanners, the second row is the relationship between the Cam-CAN scanner and the rest, etc.*

Below is the pairwise Pearson correlations between each pair of scanners. The first table shows the correlations between the first scans at each site. The second table shows the correlations between the second scan at each site.

|  | | BASE-II | Barcelona | Cam-CAN | LCBC-Avanto | LCBC-Prisma | LCBC-Skyra | Betula |
| --- | --- | --- | --- | --- | --- | --- | --- | --- |
| BASE-II |  | 1.00 | 0.98 | 0.94 | 0.96 | 0.94 | 0.99 | 0.98 |
| Barcelona |  |  | 1.00 | 0.98 | 0.98 | 0.98 | 0.98 | 0.99 |
| Cam-CAN |  |  |  | 1.00 | 0.98 | 0.98 | 0.94 | 0.96 |
| LCBC-Avanto |  |  |  |  | 1.00 | 0.98 | 0.95 | 0.96 |
| LCBC-Prisma |  |  |  |  |  | 1.00 | 0.93 | 0.95 |
| LCBC-Skyra |  |  |  |  |  |  |  | 0.99 |
| Betula |  |  |  |  |  |  |  | 1.00 |

*Pearson correlations between each pair of scanners for the first scan at each site.*

|  | | BASE-II | Barcelona | Cam-CAN | LCBC-Avanto | LCBC-Prisma | LCBC-Skyra | Betula |
| --- | --- | --- | --- | --- | --- | --- | --- | --- |
| BASE-II |  | 1.00 | 0.98 | 0.97 | 0.99 | 0.98 | 1.00 | 0.97 |
| Barcelona |  |  | 1.00 | 0.99 | 0.98 | 0.98 | 0.99 | 0.97 |
| Cam-CAN |  |  |  | 1.00 | 0.99 | 0.98 | 0.98 | 0.94 |
| LCBC-Avanto |  |  |  |  | 1.00 | 0.98 | 1.00 | 0.94 |
| LCBC-Prisma |  |  |  |  |  | 1.00 | 0.99 | 0.96 |
| LCBC-Skyra |  |  |  |  |  |  |  | 0.97 |
| Betula |  |  |  |  |  |  |  | 1.00 |

*Pearson correlations between each pair of scanners for the second scan at each site.*

**References**

1.	Reuter, M., et al., *Within-subject template estimation for unbiased longitudinal image analysis.* Neuroimage, 2012. **61**(4): p. 1402-18.

2.	Reuter, M. and B. Fischl, *Avoiding asymmetry-induced bias in longitudinal image processing.* Neuroimage, 2011. **57**(1): p. 19-21.

3.	Reuter, M., H.D. Rosas, and B. Fischl, *Highly accurate inverse consistent registration: a robust approach.* Neuroimage, 2010. **53**(4): p. 1181-96.
