## Supplementary material for "Self-reported sleep relates to hippocampal atrophy across the adult lifespan – results from the Lifebrain consortium": SI

UK Biobank Sleep Analysis

Øystein Sørensen

3/20/2019

Table of Contents

library(tidyverse)
library(readr)
library(mgcv)
library(itsadug)

### Data Preparation

Read in the data.

data <- read_csv("ukb26773_SLEEP_HC.csv")

Take a look.

```
glimpse(data)
## Observations: 21,390
## Variables: 14
## $ eid           <dbl> 5643511, 4794192, 4341865, 4237405, 2328310, 52796…
## $ sex           <dbl> 0, 1, 1, 0, 1, 0, 0, 0, 1, 0, 0, 0, 1, 1, 0, 0, 0,…
## $ mridate       <date> 2018-05-13, 2015-09-06, 2016-01-21, 2017-09-18, 2…
## $ sleepduration <dbl> 8, 7, 7, 6, 7, 9, 9, 6, 8, 7, 7, 8, 6, 6, 7, 8, 5,…
## $ sleeplessness <dbl> 2, 1, 3, 3, 2, 2, 2, 2, 2, 2, 1, 2, 1, 3, 1, 2, 3,…
## $ dozing        <dbl> 1, 1, 1, 0, 1, 0, 0, 0, 0, 0, 0, 0, 1, 0, 0, 1, 2,…
## $ napday        <dbl> 2, 1, 2, 1, 1, 1, 1, 1, 1, 1, 2, 1, 2, 1, 1, 2, 2,…
## $ gettingup     <dbl> 2, 4, 3, 3, 3, 3, 4, 4, 4, 3, 3, 4, 4, 4, 3, 4, 4,…
## $ snoring       <dbl> 1, 2, 2, 2, 1, 2, 2, 2, 1, 2, 1, 2, 1, 2, 2, 2, 1,…
## $ hc_l          <dbl> 4038, 4235, 3765, 3797, 3823, 4041, 3625, 2503, 35…
## $ hc_r          <dbl> 4366, 3831, 4398, 3917, 3680, 4325, 3976, 1473, 29…
## $ t1scaling     <dbl> 1.41738, 1.18305, 1.23224, 1.42891, 1.12407, 1.330…
## $ birthdate     <date> 1957-12-01, 1960-07-01, 1943-08-01, 1948-05-01, 1…
## $ mriage        <dbl> 60.65, 55.37, 72.72, 69.62, 68.76, 55.43, 58.70, 6…
```

The column t1scaling is *“Volumetric scaling from T1 head image to standard space”*. We use this, since we do not have an ICV variable.

For the sleep variables, -1 means *do not know* and -3 means *prefer not to answer* ([link](http://biobank.ndph.ox.ac.uk/showcase/coding.cgi?id=100291)). We encode both of these as missing.

We add left and right hippocampus and remove unnecessary variables.

```
data <- data %>%
  mutate(hippocampus = hc_l + hc_r) %>%
  mutate_at(vars(sleepduration, sleeplessness, dozing, napday, gettingup, snoring),
            list(~ if_else(. < 0, NA_integer_, as.integer(.)))) %>%
  select(-hc_l, -hc_r, -birthdate, -mridate, -eid)
```

We compare the hippocampus volume for each value of sex. It suggests that 0 = Female and 1 = Female.

```
data %>%
  group_by(sex) %>%
  summarise(
    mean_hippocampus = mean(hippocampus),
    median_hippocampus = median(hippocampus),
    num_samples = n(),
    mean_age = mean(mriage)
  )
## # A tibble: 2 x 5
##     sex mean_hippocampus median_hippocampus num_samples mean_age
##   <dbl>            <dbl>              <dbl>       <int>    <dbl>
## 1     0            7505.               7525       11237     62.7
## 2     1            7919.               7945       10153     64.1
data <- data %>%
  mutate(sex = recode(sex, `0` = "Female", `1` = "Male"))
```

Age in the sample.

```
ggplot(data, aes(mriage)) +
  geom_histogram(bins = 20)
```

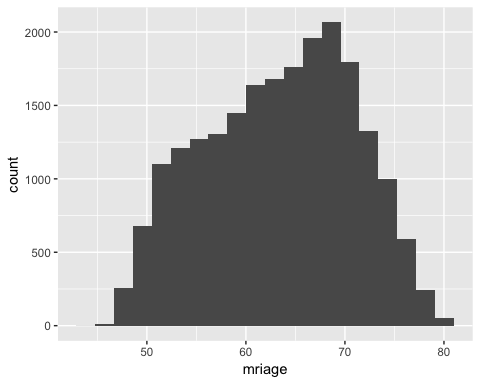

We also plot hippocampus volumes.

```
ggplot(data, aes(x = hippocampus)) +
  geom_histogram(bins = 30) +
  facet_wrap(vars(sex))
```

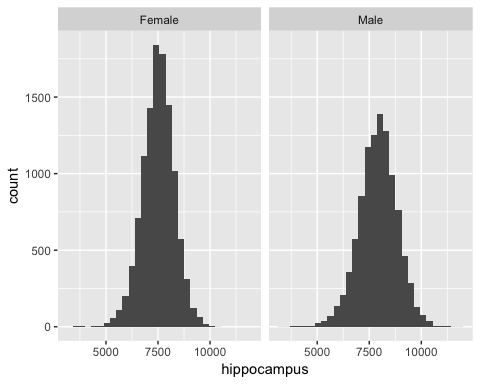

We gather the sleep measure values, for ease of filtering and plotting. We also remove missing values with the na.rm = TRUE option.

```
data <- data %>%
  gather(key = "key", value = "value", sleepduration, sleeplessness,
         dozing, napday, gettingup, snoring, na.rm = TRUE)
ggplot(data, aes(x = value)) +
  geom_histogram(binwidth = 1) +
  facet_wrap(vars(key), scales = "free_x")
```

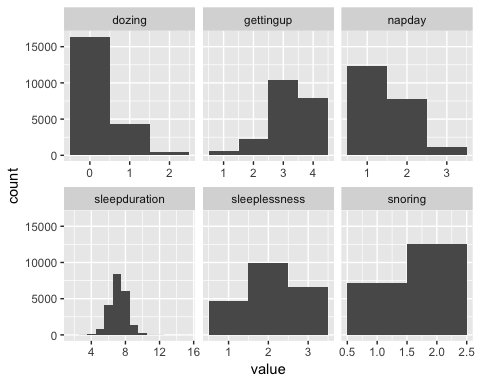

These are the number of measurements after removing missing values.

```
data %>%
  group_by(key) %>%
  count()
## # A tibble: 6 x 2
## # Groups:   key [6]
##   key               n
##   <chr>         <int>
## 1 dozing        21216
## 2 gettingup     21229
## 3 napday        21237
## 4 sleepduration 21192
## 5 sleeplessness 21229
## 6 snoring       19782
```

Function for subsetting data to the right sleep variable.

```
subset_data <- function(data, sleep_var){
  data %>%
    filter(key == !!sleep_var) %>%
    rename(!!sleep_var := value)
}
```

### Analysis

#### Dozing

sleep_var <- "dozing"

[Reference](http://biobank.ndph.ox.ac.uk/showcase/field.cgi?id=1220)

Data-Field 1220 Description: Daytime dozing / sleeping (narcolepsy)

- Never/rarely
- Sometimes
- Often
- All of the time

**Low number is good**.

```
subset_data(data, sleep_var) %>%
  group_by_at(vars(sleep_var)) %>%
  count()
## # A tibble: 3 x 2
## # Groups:   dozing [3]
##   dozing     n
##    <int> <int>
## 1      0 16378
## 2      1  4340
## 3      2   498
```

Dozing corresponds closest to PSQI Question 8.

##### Effect of Age

```
form <- reformulate("s(mriage) + sex", response = sleep_var)
m0 <- gam(form, data = subset_data(data, sleep_var))
summary(m0)
##
## Family: gaussian
## Link function: identity
##
## Formula:
## dozing ~ s(mriage) + sex
##
## Parametric coefficients:
##             Estimate Std. Error t value Pr(>|t|)
## (Intercept) 0.229437   0.004573  50.176  < 2e-16 ***
## sexMale     0.046510   0.006657   6.986 2.91e-12 ***
## ---
## Signif. codes:  0 '***' 0.001 '**' 0.01 '*' 0.05 '.' 0.1 ' ' 1
##
## Approximate significance of smooth terms:
##             edf Ref.df     F p-value
## s(mriage) 3.499  4.371 53.54  <2e-16 ***
## ---
## Signif. codes:  0 '***' 0.001 '**' 0.01 '*' 0.05 '.' 0.1 ' ' 1
##
## R-sq.(adj) =  0.0142   Deviance explained = 1.44%
## GCV = 0.23192  Scale est. = 0.23186   n = 21216
plot(m0)
```

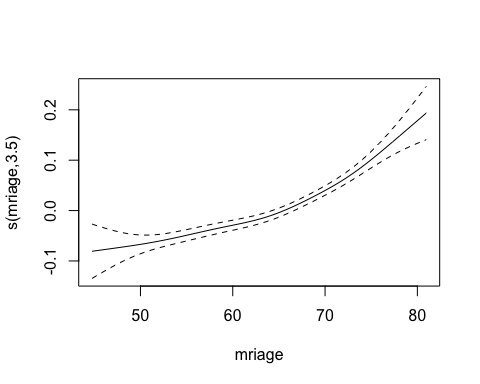

The amount of dozing increases steadily with age.

##### Effect on Hippocampus Volume

Main effect of sleep variable.

```
form <- reformulate(paste("s(mriage) + sex + t1scaling + s(", sleep_var, ", k = 3)"), response = "hippocampus")
m0 <- gam(form, data = subset_data(data, sleep_var))
summary(m0)
##
## Family: gaussian
## Link function: identity
##
## Formula:
## hippocampus ~ s(mriage) + sex + t1scaling + s(dozing, k = 3)
##
## Parametric coefficients:
##             Estimate Std. Error t value Pr(>|t|)
## (Intercept) 11460.85      74.74 153.344   <2e-16 ***
## sexMale        23.00      13.36   1.721   0.0853 .
## t1scaling   -2902.53      54.26 -53.497   <2e-16 ***
## ---
## Signif. codes:  0 '***' 0.001 '**' 0.01 '*' 0.05 '.' 0.1 ' ' 1
##
## Approximate significance of smooth terms:
##             edf Ref.df       F p-value
## s(mriage) 3.952  4.911 464.224  <2e-16 ***
## s(dozing) 1.000  1.000   1.808   0.179
## ---
## Signif. codes:  0 '***' 0.001 '**' 0.01 '*' 0.05 '.' 0.1 ' ' 1
##
## R-sq.(adj) =  0.244   Deviance explained = 24.4%
## GCV = 5.7183e+05  Scale est. = 5.7162e+05  n = 21216
plot(m0, pages = 1, seWithMean = TRUE)
```

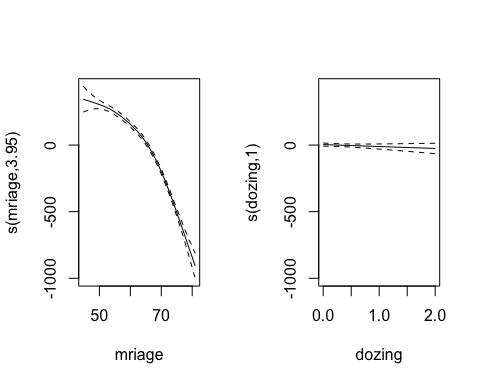

Interaction with age.

```
form <- reformulate(paste("s(mriage) + sex + t1scaling + s(",
                          sleep_var, ", k = 3) + ti(mriage, ",
                          sleep_var, ", k = c(10, 3))"),
                    response = "hippocampus")
m1 <- gam(form, data = subset_data(data, sleep_var))
summary(m1)
##
## Family: gaussian
## Link function: identity
##
## Formula:
## hippocampus ~ s(mriage) + sex + t1scaling + s(dozing, k = 3) +
##     ti(mriage, dozing, k = c(10, 3))
##
## Parametric coefficients:
##             Estimate Std. Error t value Pr(>|t|)
## (Intercept) 11459.77      74.74 153.324   <2e-16 ***
## sexMale        23.00      13.36   1.721   0.0852 .
## t1scaling   -2901.75      54.25 -53.486   <2e-16 ***
## ---
## Signif. codes:  0 '***' 0.001 '**' 0.01 '*' 0.05 '.' 0.1 ' ' 1
##
## Approximate significance of smooth terms:
##                     edf Ref.df       F p-value
## s(mriage)         3.824  4.759 464.366  <2e-16 ***
## s(dozing)         1.000  1.000   1.807   0.179
## ti(mriage,dozing) 5.727  7.195   1.670   0.110
## ---
## Signif. codes:  0 '***' 0.001 '**' 0.01 '*' 0.05 '.' 0.1 ' ' 1
##
## R-sq.(adj) =  0.244   Deviance explained = 24.5%
## GCV = 5.7174e+05  Scale est. = 5.7138e+05  n = 21216
par(mfrow = c(1, 2))
plot(m1, select = 1, seWithMean = TRUE)
plot(m1, select = 2, seWithMean = TRUE)
```

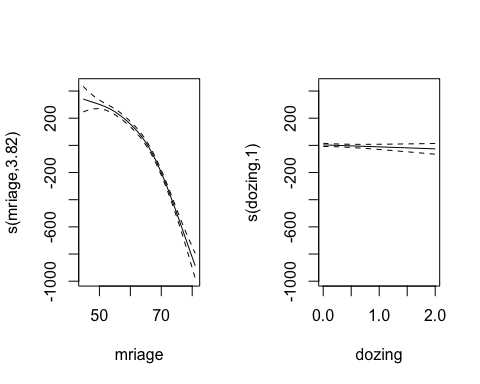

```
fvisgam(m0, view = c("mriage", sleep_var))
## Summary:
##  * sex : factor; set to the value(s): Female.
##  * t1scaling : numeric predictor; set to the value(s): 1.29561.
##  * mriage : numeric predictor; with 30 values ranging from 44.760000 to 80.970000.
##  * dozing : numeric predictor; with 30 values ranging from 0.000000 to 2.000000.
```

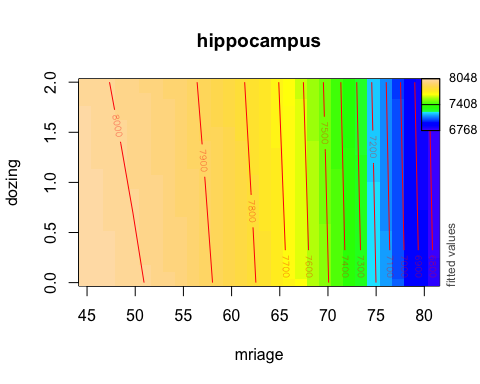

We do not find a main effect of dozing on hippocampus volume and do not find an interaction effect of dozing and age on hippocampus volume.

#### Getting Up

sleep_var <- "gettingup"

[Reference](http://biobank.ndph.ox.ac.uk/showcase/field.cgi?id=1170)

Data-Field 1170 Description: Getting up in morning

- Not at all easy
- Not very easy
- Fairly easy
- Very easy

**High number is good**.

```
subset_data(data, sleep_var) %>%
  group_by_at(vars(sleep_var)) %>%
  count()
## # A tibble: 4 x 2
## # Groups:   gettingup [4]
##   gettingup     n
##       <int> <int>
## 1         1   566
## 2         2  2298
## 3         3 10461
## 4         4  7904
```

I cannot really find a corresponding PSQI question.

##### Effect of Age

```
form <- reformulate("s(mriage) + sex", response = sleep_var)
m0 <- gam(form, data = subset_data(data, sleep_var))
summary(m0)
##
## Family: gaussian
## Link function: identity
##
## Formula:
## gettingup ~ s(mriage) + sex
##
## Parametric coefficients:
##             Estimate Std. Error t value Pr(>|t|)
## (Intercept) 3.118708   0.006785  459.66   <2e-16 ***
## sexMale     0.193921   0.009877   19.63   <2e-16 ***
## ---
## Signif. codes:  0 '***' 0.001 '**' 0.01 '*' 0.05 '.' 0.1 ' ' 1
##
## Approximate significance of smooth terms:
##             edf Ref.df     F p-value
## s(mriage) 4.408  5.442 156.7  <2e-16 ***
## ---
## Signif. codes:  0 '***' 0.001 '**' 0.01 '*' 0.05 '.' 0.1 ' ' 1
##
## R-sq.(adj) =  0.0593   Deviance explained = 5.95%
## GCV = 0.51081  Scale est. = 0.51066   n = 21229
plot(m0)
```

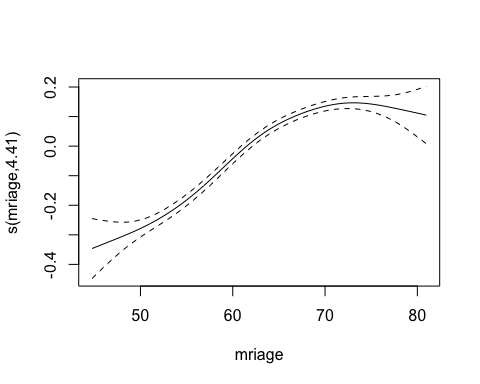

Older people have **less trouble** getting up in the morning than younger people.

##### Effect on Hippocampus Volume

Main effect of sleep variable.

```
form <- reformulate(paste("s(mriage) + sex + t1scaling + s(",
                          sleep_var, ", k = 3)"),
                    response = "hippocampus")
m0 <- gam(form, data = subset_data(data, sleep_var))
summary(m0)
##
## Family: gaussian
## Link function: identity
##
## Formula:
## hippocampus ~ s(mriage) + sex + t1scaling + s(gettingup, k = 3)
##
## Parametric coefficients:
##             Estimate Std. Error t value Pr(>|t|)
## (Intercept) 11463.83      74.72 153.428   <2e-16 ***
## sexMale        23.31      13.41   1.738   0.0822 .
## t1scaling   -2904.95      54.25 -53.550   <2e-16 ***
## ---
## Signif. codes:  0 '***' 0.001 '**' 0.01 '*' 0.05 '.' 0.1 ' ' 1
##
## Approximate significance of smooth terms:
##                edf Ref.df       F p-value
## s(mriage)    3.900  4.849 459.765  <2e-16 ***
## s(gettingup) 1.762  1.943   3.322   0.064 .
## ---
## Signif. codes:  0 '***' 0.001 '**' 0.01 '*' 0.05 '.' 0.1 ' ' 1
##
## R-sq.(adj) =  0.244   Deviance explained = 24.4%
## GCV = 5.719e+05  Scale est. = 5.7167e+05  n = 21229
plot(m0, pages = 1, seWithMean = TRUE)
```

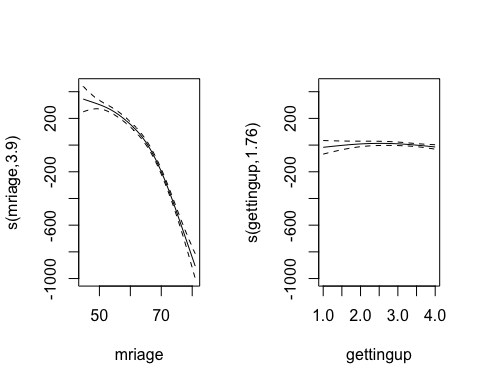

Interaction with age.

```
form <- reformulate(paste("s(mriage) + sex + t1scaling + s(",
                          sleep_var, ", k = 3) + ti(mriage, ",
                          sleep_var, ", k = c(10, 3))"),
                    response = "hippocampus")
m1 <- gam(form, data = subset_data(data, sleep_var))
summary(m1)
##
## Family: gaussian
## Link function: identity
##
## Formula:
## hippocampus ~ s(mriage) + sex + t1scaling + s(gettingup, k = 3) +
##     ti(mriage, gettingup, k = c(10, 3))
##
## Parametric coefficients:
##             Estimate Std. Error t value Pr(>|t|)
## (Intercept) 11465.46      74.73 153.417   <2e-16 ***
## sexMale        23.25      13.41   1.733   0.0831 .
## t1scaling   -2905.27      54.25 -53.555   <2e-16 ***
## ---
## Signif. codes:  0 '***' 0.001 '**' 0.01 '*' 0.05 '.' 0.1 ' ' 1
##
## Approximate significance of smooth terms:
##                        edf Ref.df       F p-value
## s(mriage)            3.979  4.944 451.031  <2e-16 ***
## s(gettingup)         1.751  1.938   3.014  0.0843 .
## ti(mriage,gettingup) 1.000  1.000   1.201  0.2731
## ---
## Signif. codes:  0 '***' 0.001 '**' 0.01 '*' 0.05 '.' 0.1 ' ' 1
##
## R-sq.(adj) =  0.244   Deviance explained = 24.4%
## GCV = 5.7193e+05  Scale est. = 5.7167e+05  n = 21229
par(mfrow = c(1, 2))
plot(m1, select = 1, seWithMean = TRUE)
plot(m1, select = 2, seWithMean = TRUE)
```

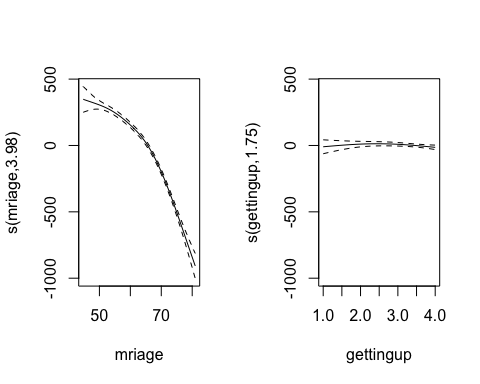

```
fvisgam(m0, view = c("mriage", sleep_var))
## Summary:
##  * sex : factor; set to the value(s): Female.
##  * t1scaling : numeric predictor; set to the value(s): 1.29554.
##  * mriage : numeric predictor; with 30 values ranging from 44.760000 to 80.970000.
##  * gettingup : numeric predictor; with 30 values ranging from 1.000000 to 4.000000.
## Warning in gradientLegend(c(min.z, max.z), n.seg = 3, pos = 0.875, color
## = pal, : Decimal set to 0 (dec=0), but the labels still don't fit in the
## margin. You may want to add the color legend to another side, or increase
## the margin of the plot.
```

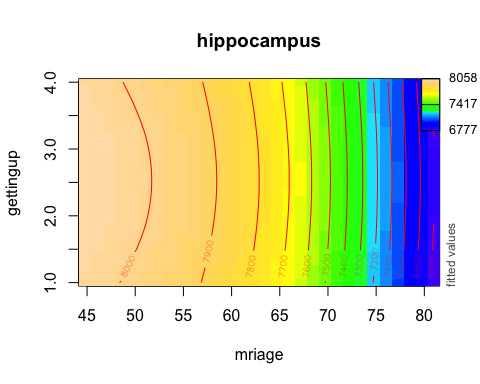

#### Nap During Day

sleep_var <- "napday"

[Reference](http://biobank.ndph.ox.ac.uk/showcase/field.cgi?id=1190)

- Never/rarely
- Sometimes
- Usually

**Low is good**.

```
subset_data(data, sleep_var) %>%
  group_by_at(vars(sleep_var)) %>%
  count()
## # A tibble: 3 x 2
## # Groups:   napday [3]
##   napday     n
##    <int> <int>
## 1      1 12284
## 2      2  7776
## 3      3  1177
```

I cannot really find a corresponding PSQI question.

##### Effect of Age

```
form <- reformulate("s(mriage) + sex", response = sleep_var)
m0 <- gam(form, data = subset_data(data, sleep_var))
summary(m0)
##
## Family: gaussian
## Link function: identity
##
## Formula:
## napday ~ s(mriage) + sex
##
## Parametric coefficients:
##             Estimate Std. Error t value Pr(>|t|)
## (Intercept) 1.387332   0.005556  249.68   <2e-16 ***
## sexMale     0.188894   0.008088   23.36   <2e-16 ***
## ---
## Signif. codes:  0 '***' 0.001 '**' 0.01 '*' 0.05 '.' 0.1 ' ' 1
##
## Approximate significance of smooth terms:
##           edf Ref.df     F p-value
## s(mriage) 2.9  3.642 120.9  <2e-16 ***
## ---
## Signif. codes:  0 '***' 0.001 '**' 0.01 '*' 0.05 '.' 0.1 ' ' 1
##
## R-sq.(adj) =  0.0491   Deviance explained = 4.93%
## GCV = 0.34272  Scale est. = 0.34264   n = 21237
plot(m0)
```

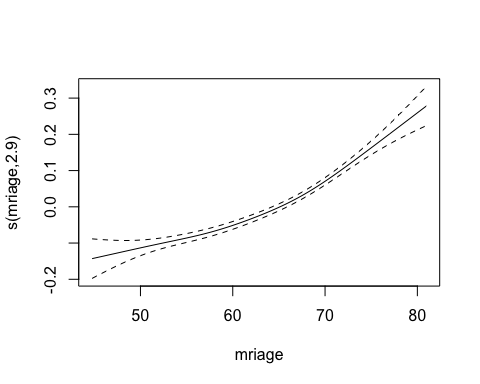

The older people are, the more likely it is that they take naps during the day.

##### Effect on Hippocampus Volume

Main effect of sleep variable.

```
form <- reformulate(paste("s(mriage) + sex + t1scaling + s(",
                          sleep_var, ", k = 3)"),
                    response = "hippocampus")
m0 <- gam(form, data = subset_data(data, sleep_var))
summary(m0)
##
## Family: gaussian
## Link function: identity
##
## Formula:
## hippocampus ~ s(mriage) + sex + t1scaling + s(napday, k = 3)
##
## Parametric coefficients:
##             Estimate Std. Error t value Pr(>|t|)
## (Intercept) 11465.40      74.72 153.453   <2e-16 ***
## sexMale        23.83      13.43   1.774   0.0761 .
## t1scaling   -2906.25      54.25 -53.571   <2e-16 ***
## ---
## Signif. codes:  0 '***' 0.001 '**' 0.01 '*' 0.05 '.' 0.1 ' ' 1
##
## Approximate significance of smooth terms:
##             edf Ref.df       F p-value
## s(mriage) 3.974  4.937 456.022  <2e-16 ***
## s(napday) 1.756  1.941   1.868   0.116
## ---
## Signif. codes:  0 '***' 0.001 '**' 0.01 '*' 0.05 '.' 0.1 ' ' 1
##
## R-sq.(adj) =  0.244   Deviance explained = 24.4%
## GCV = 5.7202e+05  Scale est. = 5.7179e+05  n = 21237
plot(m0, pages = 1, seWithMean = TRUE)
```

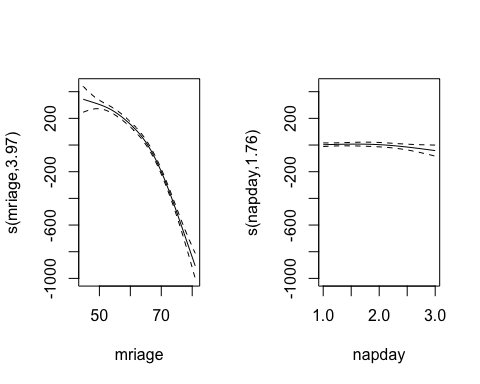

Interaction with age.

```
form <- reformulate(paste("s(mriage) + sex + t1scaling + s(",
                          sleep_var, ", k = 3) + ti(mriage, ",
                          sleep_var, ", k = c(10, 3))"),
                    response = "hippocampus")
m1 <- gam(form, data = subset_data(data, sleep_var))
summary(m1)
##
## Family: gaussian
## Link function: identity
##
## Formula:
## hippocampus ~ s(mriage) + sex + t1scaling + s(napday, k = 3) +
##     ti(mriage, napday, k = c(10, 3))
##
## Parametric coefficients:
##             Estimate Std. Error t value Pr(>|t|)
## (Intercept) 11465.75      74.71 153.471   <2e-16 ***
## sexMale        23.10      13.44   1.719   0.0856 .
## t1scaling   -2906.83      54.25 -53.585   <2e-16 ***
## ---
## Signif. codes:  0 '***' 0.001 '**' 0.01 '*' 0.05 '.' 0.1 ' ' 1
##
## Approximate significance of smooth terms:
##                     edf Ref.df       F p-value
## s(mriage)         4.024  4.996 438.608  <2e-16 ***
## s(napday)         1.758  1.940   1.945   0.105
## ti(mriage,napday) 2.863  3.939   1.112   0.339
## ---
## Signif. codes:  0 '***' 0.001 '**' 0.01 '*' 0.05 '.' 0.1 ' ' 1
##
## R-sq.(adj) =  0.244   Deviance explained = 24.4%
## GCV = 5.7198e+05  Scale est. = 5.7167e+05  n = 21237
par(mfrow = c(1, 2))
plot(m1, select = 1, seWithMean = TRUE)
plot(m1, select = 2, seWithMean = TRUE)
```

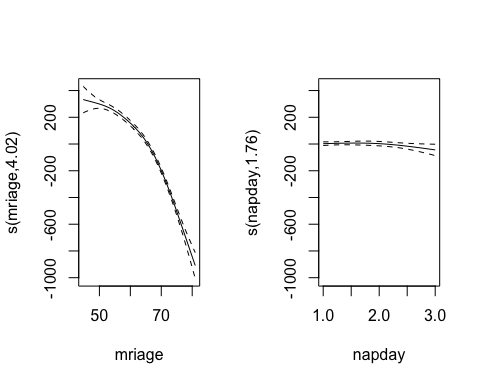

```
fvisgam(m0, view = c("mriage", sleep_var))
## Summary:
##  * sex : factor; set to the value(s): Female.
##  * t1scaling : numeric predictor; set to the value(s): 1.29556.
##  * mriage : numeric predictor; with 30 values ranging from 44.760000 to 80.970000.
##  * napday : numeric predictor; with 30 values ranging from 1.000000 to 3.000000.
## Warning in gradientLegend(c(min.z, max.z), n.seg = 3, pos = 0.875, color
## = pal, : Decimal set to 0 (dec=0), but the labels still don't fit in the
## margin. You may want to add the color legend to another side, or increase
## the margin of the plot.
```

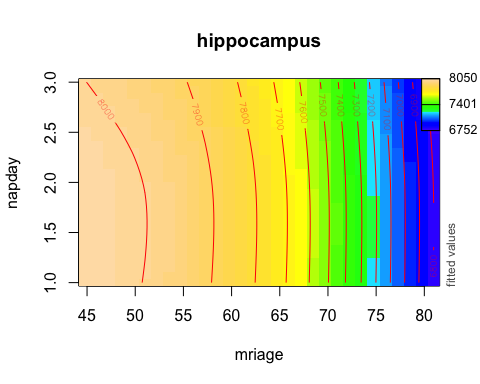

#### Sleep Duration

sleep_var <- "sleepduration"

[Reference](http://biobank.ndph.ox.ac.uk/showcase/field.cgi?id=1160)

Data-Field 1160 Description: Sleep duration

```
subset_data(data, sleep_var) %>%
  group_by_at(vars(sleep_var)) %>%
  count()
## # A tibble: 14 x 2
## # Groups:   sleepduration [14]
##    sleepduration     n
##            <int> <int>
##  1             2     2
##  2             3    23
##  3             4   143
##  4             5   867
##  5             6  4088
##  6             7  8467
##  7             8  6026
##  8             9  1307
##  9            10   225
## 10            11    21
## 11            12    18
## 12            13     1
## 13            14     1
## 14            15     3
```

This question corresponds to PSQI Question 4 (PSQI Component 3 is too coarse grained).

##### Effect of Age

```
form <- reformulate("s(mriage) + sex", response = sleep_var)
m0 <- gam(form, data = subset_data(data, sleep_var))
summary(m0)
##
## Family: gaussian
## Link function: identity
##
## Formula:
## sleepduration ~ s(mriage) + sex
##
## Parametric coefficients:
##             Estimate Std. Error t value Pr(>|t|)
## (Intercept) 7.117344   0.009891 719.572  < 2e-16 ***
## sexMale     0.068128   0.014389   4.735 2.21e-06 ***
## ---
## Signif. codes:  0 '***' 0.001 '**' 0.01 '*' 0.05 '.' 0.1 ' ' 1
##
## Approximate significance of smooth terms:
##             edf Ref.df     F p-value
## s(mriage) 4.985  6.091 25.09  <2e-16 ***
## ---
## Signif. codes:  0 '***' 0.001 '**' 0.01 '*' 0.05 '.' 0.1 ' ' 1
##
## R-sq.(adj) =  0.00862   Deviance explained = 0.89%
## GCV = 1.0824  Scale est. = 1.082     n = 21192
plot(m0)
```

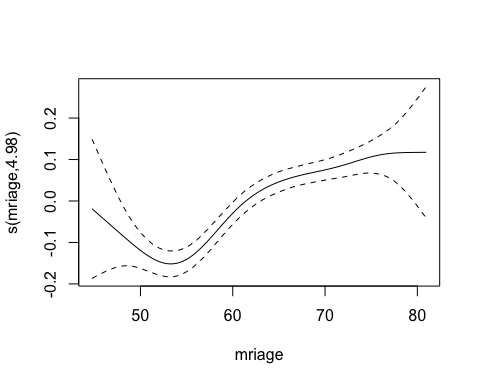

##### Effect on Hippocampus Volume

Main effect of sleep variable.

```
form <- reformulate(paste("s(mriage) + sex + t1scaling + s(",
                          sleep_var, ", k = 3)"), response = "hippocampus")
m0 <- gam(form, data = subset_data(data, sleep_var))
summary(m0)
##
## Family: gaussian
## Link function: identity
##
## Formula:
## hippocampus ~ s(mriage) + sex + t1scaling + s(sleepduration,
##     k = 3)
##
## Parametric coefficients:
##             Estimate Std. Error t value Pr(>|t|)
## (Intercept) 11458.33      74.78 153.225   <2e-16 ***
## sexMale        21.99      13.35   1.647   0.0995 .
## t1scaling   -2900.01      54.30 -53.409   <2e-16 ***
## ---
## Signif. codes:  0 '***' 0.001 '**' 0.01 '*' 0.05 '.' 0.1 ' ' 1
##
## Approximate significance of smooth terms:
##                    edf Ref.df      F  p-value
## s(mriage)        3.972  4.934 465.44  < 2e-16 ***
## s(sleepduration) 1.954  1.998  11.65 7.31e-06 ***
## ---
## Signif. codes:  0 '***' 0.001 '**' 0.01 '*' 0.05 '.' 0.1 ' ' 1
##
## R-sq.(adj) =  0.245   Deviance explained = 24.5%
## GCV = 5.7128e+05  Scale est. = 5.7104e+05  n = 21192
```

The plot suggests that sleep somewhere between 6 and 8 hours is associated with higher hippocampus volume. Note that PSQI Component 3 uses the range “< 5 hours”, “5-6 hours”, “6-7 hours”, and “> 7 hours”, which is too narrow to compare to the effects found here. We should hence try to reproduce this using the actual number of hours slept reported, in PSQI Question 4.

plot(m0, pages = 1, seWithMean = TRUE)

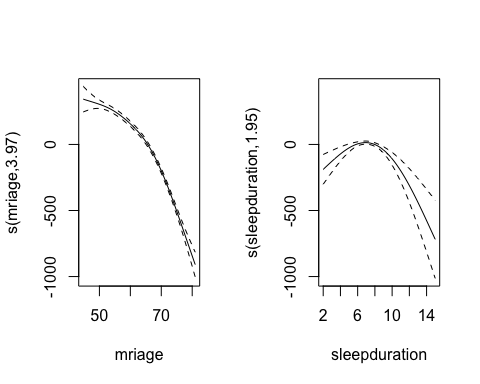

Looking at the plot above, one can hypothesize that the extreme sleep values drive the effect on hippocampus. We hence try a new analysis, restricting the number of hours slept to be between 5 and 9 hours.

```
m0b <- gam(form, data = filter(subset_data(data, sleep_var),
                              sleepduration >= 5, sleepduration <= 9))
```

For this sample, which has 20,980 participants, no effect of sleep duration on hippocampus volume is found.

```
summary(m0b)
##
## Family: gaussian
## Link function: identity
##
## Formula:
## hippocampus ~ s(mriage) + sex + t1scaling + s(sleepduration,
##     k = 3)
##
## Parametric coefficients:
##             Estimate Std. Error t value Pr(>|t|)
## (Intercept) 11471.51      75.60 151.734   <2e-16 ***
## sexMale        24.46      13.47   1.815   0.0695 .
## t1scaling   -2908.64      54.90 -52.976   <2e-16 ***
## ---
## Signif. codes:  0 '***' 0.001 '**' 0.01 '*' 0.05 '.' 0.1 ' ' 1
##
## Approximate significance of smooth terms:
##                    edf Ref.df       F p-value
## s(mriage)        4.012  4.982 456.846  <2e-16 ***
## s(sleepduration) 1.804  1.962   2.785  0.0861 .
## ---
## Signif. codes:  0 '***' 0.001 '**' 0.01 '*' 0.05 '.' 0.1 ' ' 1
##
## R-sq.(adj) =  0.246   Deviance explained = 24.7%
## GCV = 5.6987e+05  Scale est. = 5.6963e+05  n = 20755
plot(m0b, pages = 1, seWithMean = TRUE)
```

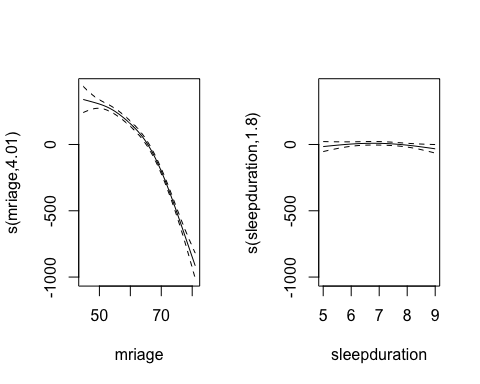

Interaction with age.

```
form <- reformulate(paste("s(mriage) + sex + t1scaling + s(",
                          sleep_var, ", k = 3) + ti(mriage, ",
                          sleep_var, ", k = c(10, 3))"),
                    response = "hippocampus")
m1 <- gam(form, data = subset_data(data, sleep_var))
summary(m1)
##
## Family: gaussian
## Link function: identity
##
## Formula:
## hippocampus ~ s(mriage) + sex + t1scaling + s(sleepduration,
##     k = 3) + ti(mriage, sleepduration, k = c(10, 3))
##
## Parametric coefficients:
##             Estimate Std. Error t value Pr(>|t|)
## (Intercept) 11456.72      74.80  153.17   <2e-16 ***
## sexMale        22.87      13.37    1.71   0.0873 .
## t1scaling   -2898.75      54.31  -53.37   <2e-16 ***
## ---
## Signif. codes:  0 '***' 0.001 '**' 0.01 '*' 0.05 '.' 0.1 ' ' 1
##
## Approximate significance of smooth terms:
##                            edf Ref.df       F  p-value
## s(mriage)                3.982  4.947 464.397  < 2e-16 ***
## s(sleepduration)         1.954  1.998  11.647 7.33e-06 ***
## ti(mriage,sleepduration) 1.000  1.000   1.106    0.293
## ---
## Signif. codes:  0 '***' 0.001 '**' 0.01 '*' 0.05 '.' 0.1 ' ' 1
##
## R-sq.(adj) =  0.245   Deviance explained = 24.5%
## GCV = 5.7131e+05  Scale est. = 5.7104e+05  n = 21192
par(mfrow = c(1, 2))
plot(m1, select = 1, seWithMean = TRUE)
plot(m1, select = 2, seWithMean = TRUE)
```

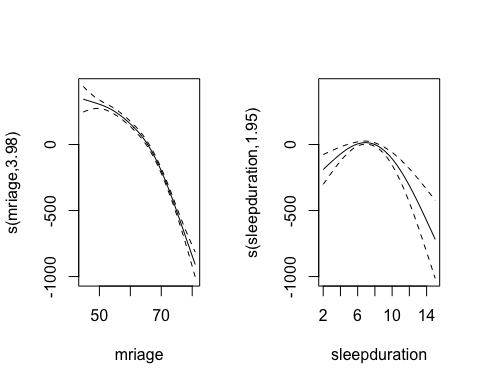

```
fvisgam(m1, view = c("mriage", sleep_var))
## Summary:
##  * sex : factor; set to the value(s): Female.
##  * t1scaling : numeric predictor; set to the value(s): 1.29537.
##  * mriage : numeric predictor; with 30 values ranging from 44.760000 to 80.970000.
##  * sleepduration : numeric predictor; with 30 values ranging from 2.000000 to 15.000000.
## Warning in gradientLegend(c(min.z, max.z), n.seg = 3, pos = 0.875, color
## = pal, : Decimal set to 0 (dec=0), but the labels still don't fit in the
## margin. You may want to add the color legend to another side, or increase
## the margin of the plot.
```

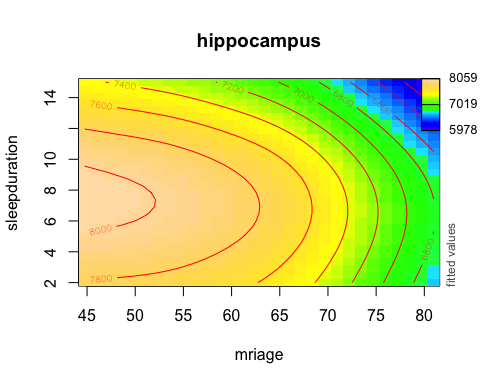

Like with the main effect, we try again, but restricting to between 5 and 9 hours of sleep. Again, no effect of duration is found in this restricted sample.

```
m1b <- gam(form,
           data = filter(subset_data(data, sleep_var),
                         sleepduration >= 5, sleepduration <= 9))
summary(m1b)
##
## Family: gaussian
## Link function: identity
##
## Formula:
## hippocampus ~ s(mriage) + sex + t1scaling + s(sleepduration,
##     k = 3) + ti(mriage, sleepduration, k = c(10, 3))
##
## Parametric coefficients:
##             Estimate Std. Error t value Pr(>|t|)
## (Intercept) 11469.45      75.61 151.692   <2e-16 ***
## sexMale        25.87      13.50   1.916   0.0553 .
## t1scaling   -2906.97      54.91 -52.939   <2e-16 ***
## ---
## Signif. codes:  0 '***' 0.001 '**' 0.01 '*' 0.05 '.' 0.1 ' ' 1
##
## Approximate significance of smooth terms:
##                            edf Ref.df       F p-value
## s(mriage)                4.034  5.008 455.056  <2e-16 ***
## s(sleepduration)         1.770  1.947   2.297  0.1378
## ti(mriage,sleepduration) 1.000  1.000   2.841  0.0919 .
## ---
## Signif. codes:  0 '***' 0.001 '**' 0.01 '*' 0.05 '.' 0.1 ' ' 1
##
## R-sq.(adj) =  0.246   Deviance explained = 24.7%
## GCV = 5.6986e+05  Scale est. = 5.6959e+05  n = 20755
```

#### Sleeplessness

sleep_var <- "sleeplessness"

[Reference](http://biobank.ndph.ox.ac.uk/showcase/field.cgi?id=1200)

Data-Field 1200 Description: Sleeplessness / insomnia

**High is bad**

```
subset_data(data, sleep_var) %>%
  group_by_at(vars(sleep_var)) %>%
  count()
## # A tibble: 3 x 2
## # Groups:   sleeplessness [3]
##   sleeplessness     n
##           <int> <int>
## 1             1  4728
## 2             2  9942
## 3             3  6559
```

This question corresponds to somewhat to PSQI Questions 5a and 5b.

##### Effect of Age

```
form <- reformulate("s(mriage) + sex", response = sleep_var)
m0 <- gam(form, data = subset_data(data, sleep_var))
summary(m0)
##
## Family: gaussian
## Link function: identity
##
## Formula:
## sleeplessness ~ s(mriage) + sex
##
## Parametric coefficients:
##              Estimate Std. Error t value Pr(>|t|)
## (Intercept)  2.191704   0.006785   323.0   <2e-16 ***
## sexMale     -0.222202   0.009878   -22.5   <2e-16 ***
## ---
## Signif. codes:  0 '***' 0.001 '**' 0.01 '*' 0.05 '.' 0.1 ' ' 1
##
## Approximate significance of smooth terms:
##             edf Ref.df     F p-value
## s(mriage) 4.873  5.968 13.53 3.1e-15 ***
## ---
## Signif. codes:  0 '***' 0.001 '**' 0.01 '*' 0.05 '.' 0.1 ' ' 1
##
## R-sq.(adj) =  0.0258   Deviance explained = 2.61%
## GCV = 0.51089  Scale est. = 0.51072   n = 21229
plot(m0)
```

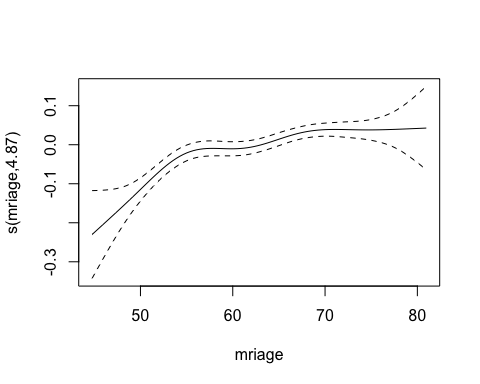

The amount of sleeplessness increases with age.

##### Effect on Hippocampus Volume

Main effect of sleep variable.

```
form <- reformulate(paste("s(mriage) + sex + t1scaling + s(",
                          sleep_var, ", k = 3)"),
                    response = "hippocampus")
m0 <- gam(form, data = subset_data(data, sleep_var))
summary(m0)
##
## Family: gaussian
## Link function: identity
##
## Formula:
## hippocampus ~ s(mriage) + sex + t1scaling + s(sleeplessness,
##     k = 3)
##
## Parametric coefficients:
##             Estimate Std. Error t value Pr(>|t|)
## (Intercept) 11463.75      74.73 153.404   <2e-16 ***
## sexMale        24.17      13.42   1.801   0.0718 .
## t1scaling   -2904.89      54.26 -53.536   <2e-16 ***
## ---
## Signif. codes:  0 '***' 0.001 '**' 0.01 '*' 0.05 '.' 0.1 ' ' 1
##
## Approximate significance of smooth terms:
##                    edf Ref.df       F p-value
## s(mriage)        3.963  4.924 468.519  <2e-16 ***
## s(sleeplessness) 1.000  1.000   2.048   0.152
## ---
## Signif. codes:  0 '***' 0.001 '**' 0.01 '*' 0.05 '.' 0.1 ' ' 1
##
## R-sq.(adj) =  0.243   Deviance explained = 24.4%
## GCV = 5.718e+05  Scale est. = 5.7159e+05  n = 21229
plot(m0, pages = 1, seWithMean = TRUE)
```

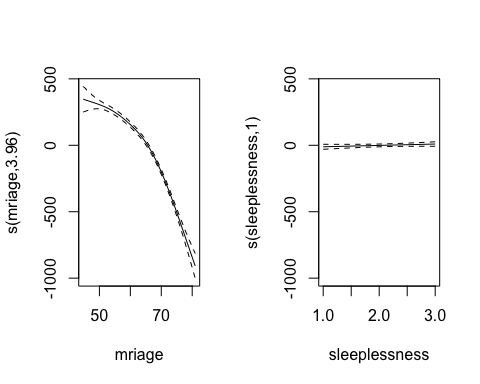

Interaction with age.

```
form <- reformulate(paste("s(mriage) + sex + t1scaling + s(",
                          sleep_var, ", k = 3) + ti(mriage, ",
                          sleep_var, ", k = c(10, 3))"),
                    response = "hippocampus")
m1 <- gam(form, data = subset_data(data, sleep_var))
summary(m1)
##
## Family: gaussian
## Link function: identity
##
## Formula:
## hippocampus ~ s(mriage) + sex + t1scaling + s(sleeplessness,
##     k = 3) + ti(mriage, sleeplessness, k = c(10, 3))
##
## Parametric coefficients:
##             Estimate Std. Error t value Pr(>|t|)
## (Intercept) 11463.32      74.73 153.398   <2e-16 ***
## sexMale        24.05      13.42   1.791   0.0733 .
## t1scaling   -2904.82      54.26 -53.534   <2e-16 ***
## ---
## Signif. codes:  0 '***' 0.001 '**' 0.01 '*' 0.05 '.' 0.1 ' ' 1
##
## Approximate significance of smooth terms:
##                            edf Ref.df       F p-value
## s(mriage)                3.962  4.923 467.579  <2e-16 ***
## s(sleeplessness)         1.000  1.000   2.240   0.134
## ti(mriage,sleeplessness) 1.523  1.772   2.158   0.211
## ---
## Signif. codes:  0 '***' 0.001 '**' 0.01 '*' 0.05 '.' 0.1 ' ' 1
##
## R-sq.(adj) =  0.243   Deviance explained = 24.4%
## GCV = 5.7178e+05  Scale est. = 5.7153e+05  n = 21229
par(mfrow = c(1, 2))
plot(m1, select = 1, seWithMean = TRUE)
plot(m1, select = 2, seWithMean = TRUE)
```

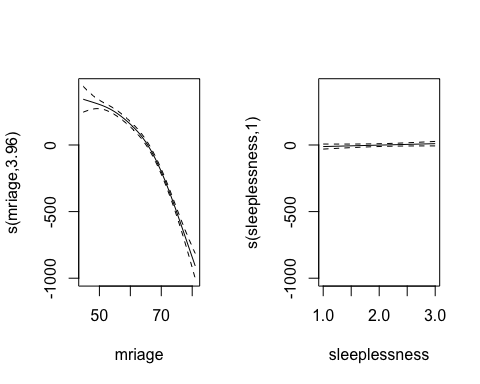

```
fvisgam(m0, view = c("mriage", sleep_var))
## Summary:
##  * sex : factor; set to the value(s): Female.
##  * t1scaling : numeric predictor; set to the value(s): 1.29557.
##  * mriage : numeric predictor; with 30 values ranging from 44.760000 to 80.970000.
##  * sleeplessness : numeric predictor; with 30 values ranging from 1.000000 to 3.000000.
## Warning in gradientLegend(c(min.z, max.z), n.seg = 3, pos = 0.875, color
## = pal, : Decimal set to 0 (dec=0), but the labels still don't fit in the
## margin. You may want to add the color legend to another side, or increase
## the margin of the plot.
```

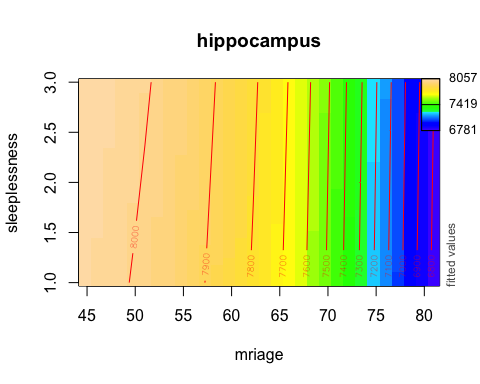

#### Snoring

sleep_var <- "snoring"

[Reference](http://biobank.ndph.ox.ac.uk/showcase/field.cgi?id=1210)

Data-Field 1210 Description: Snoring

- Yes
- No

```
subset_data(data, sleep_var) %>%
  group_by_at(vars(sleep_var)) %>%
  count()
## # A tibble: 2 x 2
## # Groups:   snoring [2]
##   snoring     n
##     <int> <int>
## 1       1  7216
## 2       2 12566
```

This question corresponds to PSQI Question 5e.

##### Effect of Age

```
form <- reformulate("s(mriage) + sex", response = sleep_var)
m0 <- gam(form, data = subset_data(data, sleep_var))
summary(m0)
##
## Family: gaussian
## Link function: identity
##
## Formula:
## snoring ~ s(mriage) + sex
##
## Parametric coefficients:
##              Estimate Std. Error t value Pr(>|t|)
## (Intercept)  1.712598   0.004704  364.06   <2e-16 ***
## sexMale     -0.160206   0.006792  -23.59   <2e-16 ***
## ---
## Signif. codes:  0 '***' 0.001 '**' 0.01 '*' 0.05 '.' 0.1 ' ' 1
##
## Approximate significance of smooth terms:
##             edf Ref.df    F  p-value
## s(mriage) 3.013   3.78 19.7 5.18e-15 ***
## ---
## Signif. codes:  0 '***' 0.001 '**' 0.01 '*' 0.05 '.' 0.1 ' ' 1
##
## R-sq.(adj) =  0.0294   Deviance explained = 2.96%
## GCV = 0.22498  Scale est. = 0.22492   n = 19782
plot(m0)
```

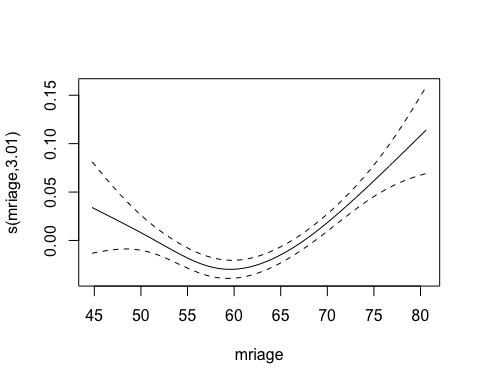

##### Effect on Hippocampus Volume

Snoring has only two values, so we code it as a factor.

```
snoring_data <- data %>%
  subset_data(sleep_var) %>%
  mutate_at(vars(sleep_var), list(snore_fac = ~ factor(.),
                                  snore_ord =~ ordered(.)))
m0 <- gam(hippocampus ~ s(mriage) + snore_fac + sex + t1scaling,
          data = snoring_data)
summary(m0)
##
## Family: gaussian
## Link function: identity
##
## Formula:
## hippocampus ~ s(mriage) + snore_fac + sex + t1scaling
##
## Parametric coefficients:
##              Estimate Std. Error t value Pr(>|t|)
## (Intercept) 11468.193     78.203 146.647   <2e-16 ***
## snore_fac2      7.674     11.361   0.675   0.4994
## sexMale        25.685     14.027   1.831   0.0671 .
## t1scaling   -2910.964     56.333 -51.674   <2e-16 ***
## ---
## Signif. codes:  0 '***' 0.001 '**' 0.01 '*' 0.05 '.' 0.1 ' ' 1
##
## Approximate significance of smooth terms:
##             edf Ref.df     F p-value
## s(mriage) 3.912  4.864 451.5  <2e-16 ***
## ---
## Signif. codes:  0 '***' 0.001 '**' 0.01 '*' 0.05 '.' 0.1 ' ' 1
##
## R-sq.(adj) =  0.245   Deviance explained = 24.6%
## GCV = 5.7425e+05  Scale est. = 5.7402e+05  n = 19782
```

Interaction with age.

```
m1 <- gam(hippocampus ~ s(mriage) + s(mriage, by = snore_ord) + sex + t1scaling,
          data = snoring_data)
summary(m1)
##
## Family: gaussian
## Link function: identity
##
## Formula:
## hippocampus ~ s(mriage) + s(mriage, by = snore_ord) + sex + t1scaling
##
## Parametric coefficients:
##             Estimate Std. Error t value Pr(>|t|)
## (Intercept) 11474.12      77.62 147.830   <2e-16 ***
## sexMale        24.50      13.89   1.764   0.0777 .
## t1scaling   -2911.21      56.32 -51.689   <2e-16 ***
## ---
## Signif. codes:  0 '***' 0.001 '**' 0.01 '*' 0.05 '.' 0.1 ' ' 1
##
## Approximate significance of smooth terms:
##                        edf Ref.df       F p-value
## s(mriage)            3.898  4.848 170.824  <2e-16 ***
## s(mriage):snore_ord2 1.000  1.000   1.889   0.169
## ---
## Signif. codes:  0 '***' 0.001 '**' 0.01 '*' 0.05 '.' 0.1 ' ' 1
##
## R-sq.(adj) =  0.245   Deviance explained = 24.6%
## GCV = 5.7421e+05  Scale est. = 5.7398e+05  n = 19782
plot(m1, pages = 1)
```

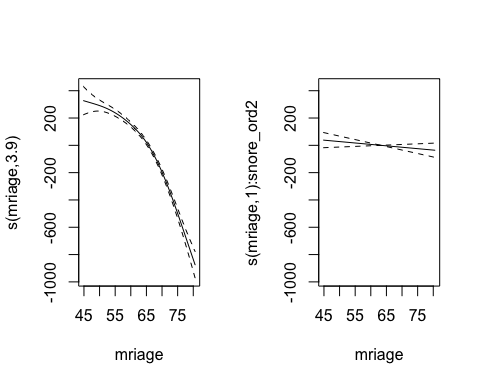
