## Supplementary material for "Self-reported sleep relates to hippocampal atrophy across the adult lifespan – results from the Lifebrain consortium": SI

Main Effect of Age on Sleep

library(dplyr)
library(ggplot2)
library(tidyr)
library(mgcv)
library(purrr)

full_data <- readRDS("./data/full_data.rds")

### Main Effect of Age on Sleep

We first investigate the main effect of Age on Sleep, including Sex as an additional covariate. These are models of the form PSQI_* ~ s(Age) + Sex, i.e., with no brain variables. Since the responses (PSQI variables) are time-invariant, we take mean age as covariate, and keep only one row per participant. This means that no random effect per participant is used, only per study. Since there are few studies, using s(Study, bs = "re") is efficient, and hence chosen rather than listing the random effects, as it allows use of gam().

We start with a data preparation for this analysis, creating a dataframe data to be used in the analyses.

```
data <- full_data %>%
  select(Age, contains("PSQI"), Sex, ID, Study) %>%
  gather(key = "key", value = "value", contains("PSQI"), na.rm = TRUE) %>%
  group_by(ID, Study, Sex, key) %>%
  summarise(
    value = mean(value),
    Age = mean(Age)
    )
```

The following defines a function for creating a plot for inspecting the data for a given PSQI question.

```
desc_plot <- function(data, psqi_var){
  data %>%
  filter(key == !! psqi_var) %>%
  ggplot(aes(x = Age, y = value)) +
  geom_jitter(aes(color = Study)) +
  ylab(psqi_var)
}
```

The following defines a function for subsetting the whole dataset, in order to obtain answers to a particular PSQI variable. It will be used repeatedly in the analyses below.

```
subset_data <- function(data, psqi_var){
  data %>%
    filter(key == psqi_var) %>%
    rename(!! psqi_var := value)
}
```

#### PSQI Component 1: Quality

We define the PSQI question to be analyzed in this section.

psqi_var <- "PSQI_Comp1_Quality"

Descriptive plot.

desc_plot(data, psqi_var)

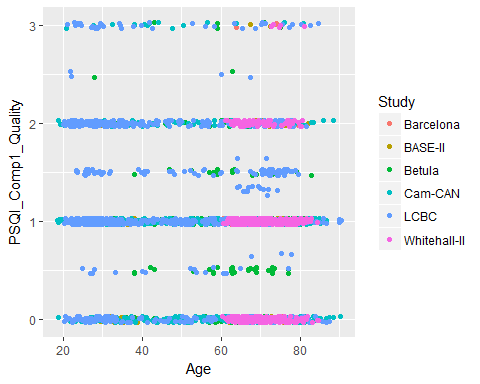

We start with a linear model with random intercept per study.

```
form <- reformulate("Age + Sex + s(Study, bs = 're')", response = psqi_var)
m0 <- gam(form, data = subset_data(data, psqi_var), method = "REML")
summary(m0)
##
## Family: gaussian
## Link function: identity
##
## Formula:
## PSQI_Comp1_Quality ~ Age + Sex + s(Study, bs = "re")
##
## Parametric coefficients:
##               Estimate Std. Error t value Pr(>|t|)
## (Intercept)  1.1176758  0.0667140  16.753  < 2e-16 ***
## Age         -0.0023842  0.0007904  -3.017  0.00258 **
## SexMale     -0.0724399  0.0272480  -2.659  0.00789 **
## ---
## Signif. codes:  0 '***' 0.001 '**' 0.01 '*' 0.05 '.' 0.1 ' ' 1
##
## Approximate significance of smooth terms:
##            edf Ref.df     F  p-value
## s(Study) 4.272      5 12.89 2.43e-14 ***
## ---
## Signif. codes:  0 '***' 0.001 '**' 0.01 '*' 0.05 '.' 0.1 ' ' 1
##
## R-sq.(adj) =  0.0464   Deviance explained = 4.84%
## -REML = 3152.3  Scale est. = 0.47664   n = 2991
```

There is a small negative association between Age and Sleep Quality. Males also seem to report slightly better sleep quality than females.

Next we try a model with a smooth term for Age.

```
form <- reformulate("s(Age) + Sex + s(Study, bs = 're')", response = psqi_var)
m1 <- gam(form, data = subset_data(data, psqi_var), method = "REML")
summary(m1)
##
## Family: gaussian
## Link function: identity
##
## Formula:
## PSQI_Comp1_Quality ~ s(Age) + Sex + s(Study, bs = "re")
##
## Parametric coefficients:
##             Estimate Std. Error t value Pr(>|t|)
## (Intercept)  0.97899    0.04670  20.964  < 2e-16 ***
## SexMale     -0.07224    0.02723  -2.653  0.00803 **
## ---
## Signif. codes:  0 '***' 0.001 '**' 0.01 '*' 0.05 '.' 0.1 ' ' 1
##
## Approximate significance of smooth terms:
##            edf Ref.df      F p-value
## s(Age)   2.266  2.839  3.955 0.00821 **
## s(Study) 4.245  5.000 12.053 1.3e-13 ***
## ---
## Signif. codes:  0 '***' 0.001 '**' 0.01 '*' 0.05 '.' 0.1 ' ' 1
##
## R-sq.(adj) =  0.0473   Deviance explained = 4.97%
## -REML = 3148.8  Scale est. = 0.47616   n = 2991
```

The estimated degrees of freedom is close to 2, suggesting that the model captures some nonlinearities.

plot(m1, seWithMean = TRUE, unconditional = TRUE, select = 1)

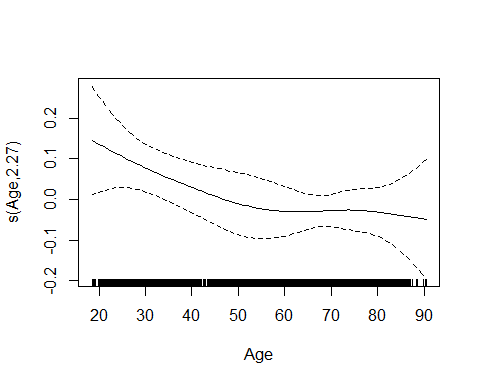

Two comments to the plotting code:

- unconditional = TRUE means that we take into account the uncertainty in smoothness estimation when computing confidence intervals.
- seWithMean = TRUE means that uncertainty about the intercept is included in the confidence bands around the curve.

We can also try random slopes per study, thus allowing the separate studies to deviate from the overall smooth fit.

```
form <- reformulate("s(Age) + Sex + s(Age, Study, bs = 're')", response = psqi_var)
m2 <- gam(form, data = subset_data(data, psqi_var), method = "REML")
summary(m2)
##
## Family: gaussian
## Link function: identity
##
## Formula:
## PSQI_Comp1_Quality ~ s(Age) + Sex + s(Age, Study, bs = "re")
##
## Parametric coefficients:
##             Estimate Std. Error t value Pr(>|t|)
## (Intercept)  0.99458    0.04300  23.132   <2e-16 ***
## SexMale     -0.07848    0.02717  -2.889   0.0039 **
## ---
## Signif. codes:  0 '***' 0.001 '**' 0.01 '*' 0.05 '.' 0.1 ' ' 1
##
## Approximate significance of smooth terms:
##                edf Ref.df      F  p-value
## s(Age)       2.843  3.561  5.296 0.000556 ***
## s(Age,Study) 4.282  5.000 10.971 2.65e-12 ***
## ---
## Signif. codes:  0 '***' 0.001 '**' 0.01 '*' 0.05 '.' 0.1 ' ' 1
##
## R-sq.(adj) =  0.0462   Deviance explained = 4.88%
## -REML = 3151.3  Scale est. = 0.47671   n = 2991
plot(m2, pages = 1, seWithMean = TRUE, unconditional = TRUE, select = 1)
```

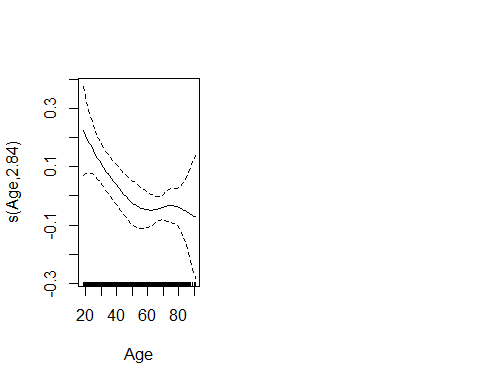

We can compare the models:

```
anova(m0, m1, m2)
## Analysis of Deviance Table
##
## Model 1: PSQI_Comp1_Quality ~ Age + Sex + s(Study, bs = "re")
## Model 2: PSQI_Comp1_Quality ~ s(Age) + Sex + s(Study, bs = "re")
## Model 3: PSQI_Comp1_Quality ~ s(Age) + Sex + s(Age, Study, bs = "re")
##   Resid. Df Resid. Dev      Df Deviance
## 1    2983.0     1422.2
## 2    2980.2     1420.1 2.80796   2.0410
## 3    2979.3     1421.5 0.88563  -1.3651
AIC(m0, m1, m2)
##           df      AIC
## m0  8.436663 6281.397
## m1 10.658999 6281.546
## m2 11.408799 6285.919
```

It seems as if a linear random intercept model is sufficient, and we conclude that there is a slight negative relationship between age and PSQI Component 1. That is, older people report slightly better sleep quality than younger people.

#### PSQI Component 2: Latency

psqi_var <- "PSQI_Comp2_Latency"

Inspection of data.

desc_plot(data, psqi_var)

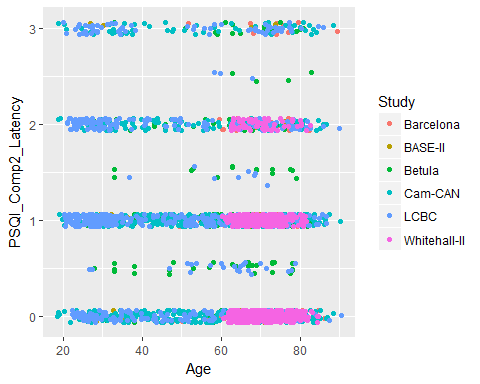

Linear model with random intercept per study.

```
form <- reformulate("Age + Sex + s(Study, bs = 're')", response = psqi_var)
m0 <- gam(form, data = subset_data(data, psqi_var), method = "REML")
summary(m0)
##
## Family: gaussian
## Link function: identity
##
## Formula:
## PSQI_Comp2_Latency ~ Age + Sex + s(Study, bs = "re")
##
## Parametric coefficients:
##               Estimate Std. Error t value Pr(>|t|)
## (Intercept)  1.1528582  0.0926373  12.445  < 2e-16 ***
## Age         -0.0017388  0.0009936  -1.750   0.0803 .
## SexMale     -0.2579115  0.0334184  -7.718 1.66e-14 ***
## ---
## Signif. codes:  0 '***' 0.001 '**' 0.01 '*' 0.05 '.' 0.1 ' ' 1
##
## Approximate significance of smooth terms:
##            edf Ref.df     F p-value
## s(Study) 4.534      5 18.02  <2e-16 ***
## ---
## Signif. codes:  0 '***' 0.001 '**' 0.01 '*' 0.05 '.' 0.1 ' ' 1
##
## R-sq.(adj) =  0.0904   Deviance explained = 9.27%
## -REML = 3236.1  Scale est. = 0.63948   n = 2693
```

Next we try a model with a smooth term for Age.

```
form <- reformulate("s(Age) + Sex + s(Study, bs = 're')", response = psqi_var)
m1 <- gam(form, data = subset_data(data, psqi_var), method = "REML")
summary(m1)
##
## Family: gaussian
## Link function: identity
##
## Formula:
## PSQI_Comp2_Latency ~ s(Age) + Sex + s(Study, bs = "re")
##
## Parametric coefficients:
##             Estimate Std. Error t value Pr(>|t|)
## (Intercept)  1.05051    0.07137  14.719  < 2e-16 ***
## SexMale     -0.25705    0.03335  -7.708 1.79e-14 ***
## ---
## Signif. codes:  0 '***' 0.001 '**' 0.01 '*' 0.05 '.' 0.1 ' ' 1
##
## Approximate significance of smooth terms:
##            edf Ref.df      F p-value
## s(Age)   2.867  3.599  4.153 0.00353 **
## s(Study) 4.528  5.000 17.191 < 2e-16 ***
## ---
## Signif. codes:  0 '***' 0.001 '**' 0.01 '*' 0.05 '.' 0.1 ' ' 1
##
## R-sq.(adj) =  0.0946   Deviance explained = 9.74%
## -REML = 3228.8  Scale est. = 0.63659   n = 2693
plot(m1, pages = 1, seWithMean = TRUE, unconditional = TRUE, select = 1)
```

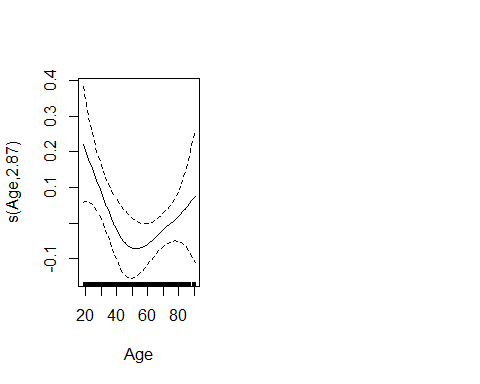

Try with a random slope per study.

```
form <- reformulate("s(Age) + Sex + s(Age, Study, bs = 're')", response = psqi_var)
m2 <- gam(form, data = subset_data(data, psqi_var), method = "REML")
summary(m2)
##
## Family: gaussian
## Link function: identity
##
## Formula:
## PSQI_Comp2_Latency ~ s(Age) + Sex + s(Age, Study, bs = "re")
##
## Parametric coefficients:
##             Estimate Std. Error t value Pr(>|t|)
## (Intercept)  1.06034    0.06380  16.619  < 2e-16 ***
## SexMale     -0.25438    0.03324  -7.652 2.73e-14 ***
## ---
## Signif. codes:  0 '***' 0.001 '**' 0.01 '*' 0.05 '.' 0.1 ' ' 1
##
## Approximate significance of smooth terms:
##                edf Ref.df      F  p-value
## s(Age)       2.947  3.695  5.422 0.000421 ***
## s(Age,Study) 4.494  5.000 18.075  < 2e-16 ***
## ---
## Signif. codes:  0 '***' 0.001 '**' 0.01 '*' 0.05 '.' 0.1 ' ' 1
##
## R-sq.(adj) =  0.0961   Deviance explained = 9.89%
## -REML = 3226.5  Scale est. = 0.63552   n = 2693
plot(m2, pages = 1, seWithMean = TRUE, unconditional = TRUE)
```

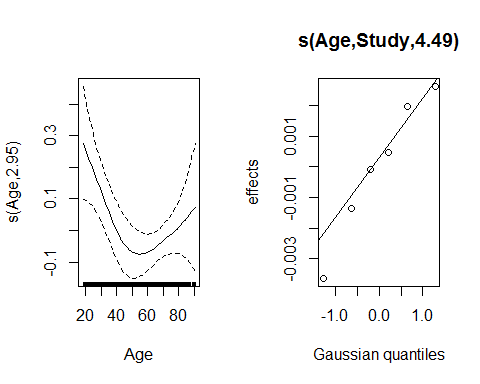

We can also try with a random smooth per study, allowing different wiggliness between studies.

```
form <- reformulate("s(Age) + s(Age, by = Study, bs = 'ts') + Sex + s(Study, bs = 're')",
                    response = psqi_var)
m3 <- gam(form, data = subset_data(data, psqi_var), method = "REML")
summary(m3)
##
## Family: gaussian
## Link function: identity
##
## Formula:
## PSQI_Comp2_Latency ~ s(Age) + s(Age, by = Study, bs = "ts") +
##     Sex + s(Study, bs = "re")
##
## Parametric coefficients:
##             Estimate Std. Error t value Pr(>|t|)
## (Intercept)  1.07575    0.05862  18.352  < 2e-16 ***
## SexMale     -0.25770    0.03330  -7.739 1.41e-14 ***
## ---
## Signif. codes:  0 '***' 0.001 '**' 0.01 '*' 0.05 '.' 0.1 ' ' 1
##
## Approximate significance of smooth terms:
##                               edf Ref.df     F  p-value
## s(Age)                   3.023184  3.788 4.857 0.000918 ***
## s(Age):StudyBarcelona    0.001598  9.000 0.000 0.872957
## s(Age):StudyBASE-II      0.004859  9.000 0.000 0.356437
## s(Age):StudyBetula       0.003128  9.000 0.000 0.340214
## s(Age):StudyCam-CAN      0.336880  9.000 0.063 0.171435
## s(Age):StudyLCBC         0.433220  9.000 0.106 0.143555
## s(Age):StudyWhitehall-II 0.852484  9.000 4.396 0.010945 *
## s(Study)                 4.119456  5.000 5.372 2.74e-06 ***
## ---
## Signif. codes:  0 '***' 0.001 '**' 0.01 '*' 0.05 '.' 0.1 ' ' 1
##
## R-sq.(adj) =  0.096   Deviance explained = 9.93%
## -REML = 3227.2  Scale est. = 0.63555   n = 2693
plot(m3, pages = 1, seWithMean = TRUE, unconditional = TRUE)
```

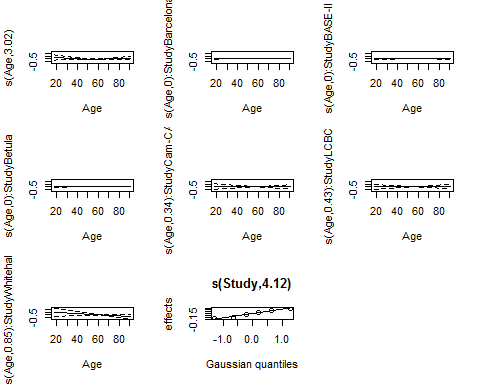

We can compare the models:

```
anova(m0, m1, m2, m3)
## Analysis of Deviance Table
##
## Model 1: PSQI_Comp2_Latency ~ Age + Sex + s(Study, bs = "re")
## Model 2: PSQI_Comp2_Latency ~ s(Age) + Sex + s(Study, bs = "re")
## Model 3: PSQI_Comp2_Latency ~ s(Age) + Sex + s(Age, Study, bs = "re")
## Model 4: PSQI_Comp2_Latency ~ s(Age) + s(Age, by = Study, bs = "ts") +
##     Sex + s(Study, bs = "re")
##   Resid. Df Resid. Dev        Df Deviance
## 1    2684.9     1717.3
## 2    2681.8     1708.3  3.131672   8.9671
## 3    2681.8     1705.5 -0.028834   2.8767
## 4    2678.4     1704.7  3.419762   0.7884
```

Random slope and smooth function of age seems justified, but different wiggliness does not seem necessary, so model m2 is chosen.

```
AIC(m0, m1, m2, m3)
##           df      AIC
## m0  8.673741 6448.179
## m1 11.071045 6438.875
## m2 11.000802 6434.196
## m3 13.687472 6438.324
```

Younger people have most trouble falling asleep, and the reported problems drop until the age around 50, after which it stabilizes and then slightly increases again.

#### PSQI Component 3: Duration

psqi_var <- "PSQI_Comp3_Duration"

Inspection of data.

desc_plot(data, psqi_var)

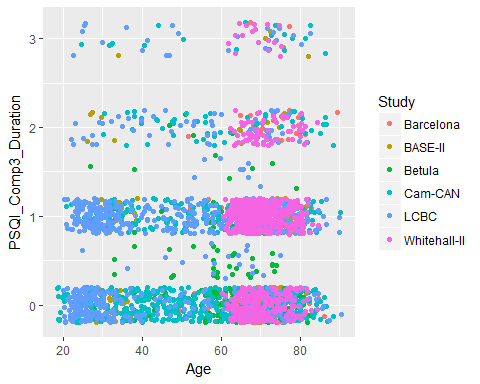

Linear model with random intercept per study.

```
form <- reformulate("Age + Sex + s(Study, bs = 're')", response = psqi_var)
m0 <- gam(form, data = subset_data(data, psqi_var), method = "REML")
summary(m0)
##
## Family: gaussian
## Link function: identity
##
## Formula:
## PSQI_Comp3_Duration ~ Age + Sex + s(Study, bs = "re")
##
## Parametric coefficients:
##               Estimate Std. Error t value Pr(>|t|)
## (Intercept)  0.3374250  0.1013599   3.329 0.000883 ***
## Age          0.0044566  0.0008305   5.366 8.68e-08 ***
## SexMale     -0.0628230  0.0279282  -2.249 0.024559 *
## ---
## Signif. codes:  0 '***' 0.001 '**' 0.01 '*' 0.05 '.' 0.1 ' ' 1
##
## Approximate significance of smooth terms:
##            edf Ref.df     F p-value
## s(Study) 4.832      5 37.77  <2e-16 ***
## ---
## Signif. codes:  0 '***' 0.001 '**' 0.01 '*' 0.05 '.' 0.1 ' ' 1
##
## R-sq.(adj) =  0.0685   Deviance explained = 7.07%
## -REML =   3065  Scale est. = 0.4817    n = 2890
```

Next we try a model with a smooth term for Age. The estimate is highly nonlinear.

```
form <- reformulate("s(Age) + Sex + s(Study, bs = 're')", response = psqi_var)
m1 <- gam(form, data = subset_data(data, psqi_var), method = "REML")
summary(m1)
##
## Family: gaussian
## Link function: identity
##
## Formula:
## PSQI_Comp3_Duration ~ s(Age) + Sex + s(Study, bs = "re")
##
## Parametric coefficients:
##             Estimate Std. Error t value Pr(>|t|)
## (Intercept)  0.60274    0.09203   6.550  6.8e-11 ***
## SexMale     -0.06248    0.02785  -2.244   0.0249 *
## ---
## Signif. codes:  0 '***' 0.001 '**' 0.01 '*' 0.05 '.' 0.1 ' ' 1
##
## Approximate significance of smooth terms:
##            edf Ref.df      F  p-value
## s(Age)   4.148   5.17  9.048 1.11e-08 ***
## s(Study) 4.842   5.00 40.829  < 2e-16 ***
## ---
## Signif. codes:  0 '***' 0.001 '**' 0.01 '*' 0.05 '.' 0.1 ' ' 1
##
## R-sq.(adj) =  0.0745   Deviance explained = 7.77%
## -REML = 3056.5  Scale est. = 0.47856   n = 2890
plot(m1, pages = 1, seWithMean = TRUE, unconditional = TRUE, select = 1)
```

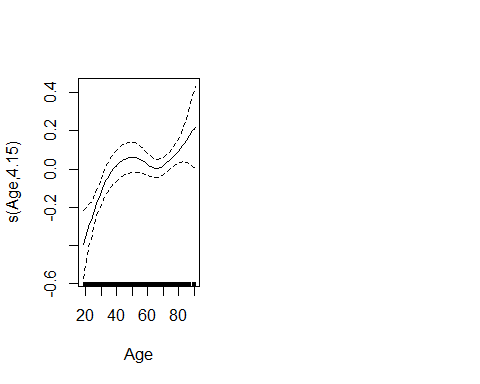

Try with a random slope per study.

```
form <- reformulate("s(Age) + Sex + s(Age, Study, bs = 're')", response = psqi_var)
m2 <- gam(form, data = subset_data(data, psqi_var), method = "REML")
summary(m2)
##
## Family: gaussian
## Link function: identity
##
## Formula:
## PSQI_Comp3_Duration ~ s(Age) + Sex + s(Age, Study, bs = "re")
##
## Parametric coefficients:
##             Estimate Std. Error t value Pr(>|t|)
## (Intercept)  0.61326    0.08695   7.053 2.19e-12 ***
## SexMale     -0.06977    0.02776  -2.513    0.012 *
## ---
## Signif. codes:  0 '***' 0.001 '**' 0.01 '*' 0.05 '.' 0.1 ' ' 1
##
## Approximate significance of smooth terms:
##                edf Ref.df      F p-value
## s(Age)       3.602  4.512  2.915  0.0164 *
## s(Age,Study) 4.855  5.000 40.130  <2e-16 ***
## ---
## Signif. codes:  0 '***' 0.001 '**' 0.01 '*' 0.05 '.' 0.1 ' ' 1
##
## R-sq.(adj) =  0.0732   Deviance explained = 7.62%
## -REML = 3057.8  Scale est. = 0.47924   n = 2890
plot(m2, select = 1, seWithMean = TRUE, unconditional = TRUE)
```

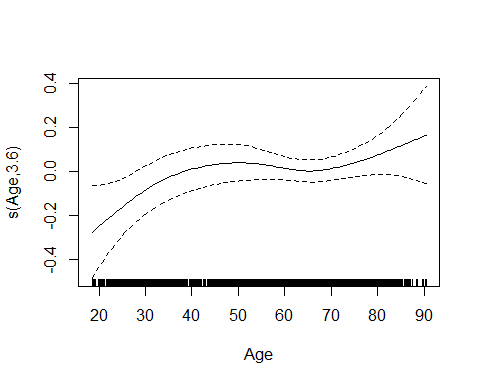

We can also try with a random smooth per study, allowing different wiggliness between studies.

```
form <- reformulate("s(Age) + s(Age, by = Study, bs = 'ts') + Sex + s(Study, bs = 're')",
                    response = psqi_var)
m3 <- gam(form, data = subset_data(data, psqi_var), method = "REML")
summary(m3)
##
## Family: gaussian
## Link function: identity
##
## Formula:
## PSQI_Comp3_Duration ~ s(Age) + s(Age, by = Study, bs = "ts") +
##     Sex + s(Study, bs = "re")
##
## Parametric coefficients:
##             Estimate Std. Error t value Pr(>|t|)
## (Intercept)  0.60538    0.08627   7.017 2.81e-12 ***
## SexMale     -0.06279    0.02779  -2.259   0.0239 *
## ---
## Signif. codes:  0 '***' 0.001 '**' 0.01 '*' 0.05 '.' 0.1 ' ' 1
##
## Approximate significance of smooth terms:
##                                edf Ref.df      F  p-value
## s(Age)                   2.919e+00  3.664  7.116 2.54e-05 ***
## s(Age):StudyBarcelona    1.248e-01  9.000  0.045 0.284171
## s(Age):StudyBASE-II      7.485e-01  9.000  0.707 0.046632 *
## s(Age):StudyBetula       9.733e-01  9.000  2.746 0.000302 ***
## s(Age):StudyCam-CAN      6.843e-05  9.000  0.000 0.983282
## s(Age):StudyLCBC         2.367e+00  9.000  1.451 0.074054 .
## s(Age):StudyWhitehall-II 1.792e-03  9.000  0.000 0.512010
## s(Study)                 4.799e+00  5.000 31.359  < 2e-16 ***
## ---
## Signif. codes:  0 '***' 0.001 '**' 0.01 '*' 0.05 '.' 0.1 ' ' 1
##
## R-sq.(adj) =  0.0801   Deviance explained = 8.42%
## -REML = 3052.5  Scale est. = 0.47568   n = 2890
plot(m3, pages = 1, seWithMean = TRUE, unconditional = TRUE)
```

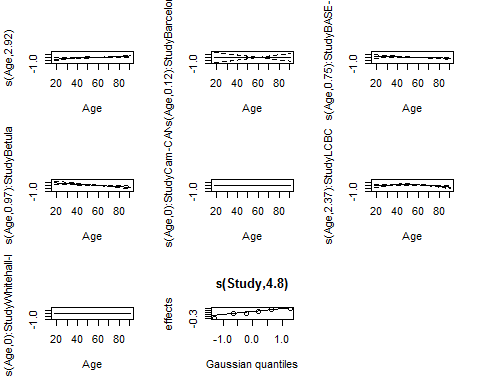

```
AIC(m0, m1, m2, m3)
##           df      AIC
## m0  8.896984 6100.460
## m1 12.585017 6085.768
## m2 12.304181 6089.863
## m3 16.861503 6073.937
```

Random smooths seem justified here, i.e., model m3 seems to fit the data best. We can extract the overall effect of age on sleep for this model. It is clear that the sleep duration is reduced with higher age (higher PSQI score means lower duration).

plot(m3, select = 1, seWithMean = TRUE, unconditional = TRUE)

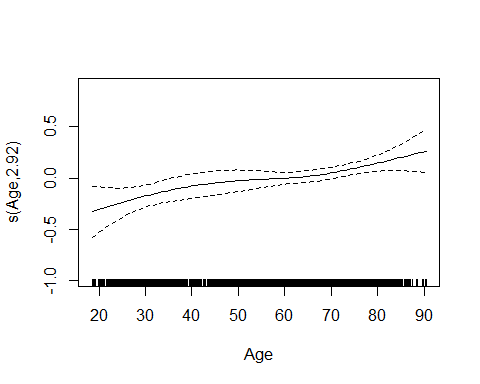

#### PSQI Component 4: Efficiency

psqi_var <- "PSQI_Comp4_Efficiency"

Inspection of data.

desc_plot(data, psqi_var)

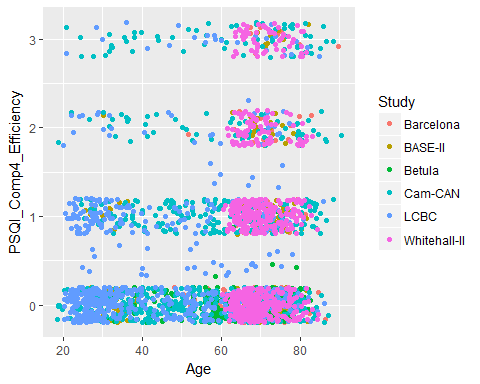

Linear model with random intercept per study.

```
form <- reformulate("Age + Sex + s(Study, bs = 're')", response = psqi_var)
m0 <- gam(form, data = subset_data(data, psqi_var), method = "REML")
summary(m0)
##
## Family: gaussian
## Link function: identity
##
## Formula:
## PSQI_Comp4_Efficiency ~ Age + Sex + s(Study, bs = "re")
##
## Parametric coefficients:
##              Estimate Std. Error t value Pr(>|t|)
## (Intercept)  0.038566   0.145090   0.266     0.79
## Age          0.009753   0.001073   9.086  < 2e-16 ***
## SexMale     -0.170415   0.034789  -4.899 1.02e-06 ***
## ---
## Signif. codes:  0 '***' 0.001 '**' 0.01 '*' 0.05 '.' 0.1 ' ' 1
##
## Approximate significance of smooth terms:
##            edf Ref.df     F p-value
## s(Study) 4.885      5 53.15  <2e-16 ***
## ---
## Signif. codes:  0 '***' 0.001 '**' 0.01 '*' 0.05 '.' 0.1 ' ' 1
##
## R-sq.(adj) =  0.122   Deviance explained = 12.4%
## -REML =   3284  Scale est. = 0.68829   n = 2649
```

Next we try a model with a smooth term for Age. The estimate is nonlinear.

```
form <- reformulate("s(Age) + Sex + s(Study, bs = 're')", response = psqi_var)
m1 <- gam(form, data = subset_data(data, psqi_var), method = "REML")
summary(m1)
##
## Family: gaussian
## Link function: identity
##
## Formula:
## PSQI_Comp4_Efficiency ~ s(Age) + Sex + s(Study, bs = "re")
##
## Parametric coefficients:
##             Estimate Std. Error t value Pr(>|t|)
## (Intercept)  0.61134    0.12867   4.751 2.13e-06 ***
## SexMale     -0.16992    0.03472  -4.894 1.05e-06 ***
## ---
## Signif. codes:  0 '***' 0.001 '**' 0.01 '*' 0.05 '.' 0.1 ' ' 1
##
## Approximate significance of smooth terms:
##            edf Ref.df     F p-value
## s(Age)   2.672  3.351 27.67  <2e-16 ***
## s(Study) 4.880  5.000 52.43  <2e-16 ***
## ---
## Signif. codes:  0 '***' 0.001 '**' 0.01 '*' 0.05 '.' 0.1 ' ' 1
##
## R-sq.(adj) =  0.125   Deviance explained = 12.8%
## -REML = 3277.5  Scale est. = 0.68564   n = 2649
plot(m1, select = 1, seWithMean = TRUE, unconditional = TRUE)
```

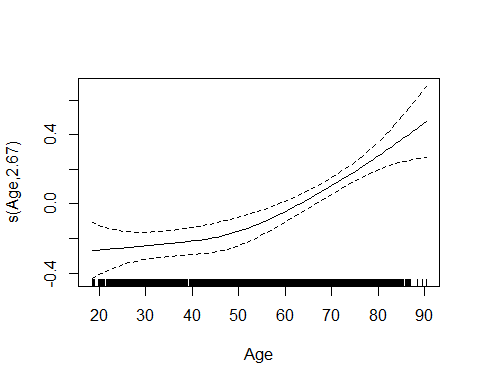

Try with a random slope per study. The overall effect of age does not change much compared to with the previous model.

```
form <- reformulate("s(Age) + Sex + s(Age, Study, bs = 're')", response = psqi_var)
m2 <- gam(form, data = subset_data(data, psqi_var), method = "REML")
summary(m2)
##
## Family: gaussian
## Link function: identity
##
## Formula:
## PSQI_Comp4_Efficiency ~ s(Age) + Sex + s(Age, Study, bs = "re")
##
## Parametric coefficients:
##             Estimate Std. Error t value Pr(>|t|)
## (Intercept)  0.61878    0.12207   5.069 4.27e-07 ***
## SexMale     -0.16965    0.03453  -4.913 9.52e-07 ***
## ---
## Signif. codes:  0 '***' 0.001 '**' 0.01 '*' 0.05 '.' 0.1 ' ' 1
##
## Approximate significance of smooth terms:
##                edf Ref.df      F  p-value
## s(Age)       2.567  3.215  6.743 0.000114 ***
## s(Age,Study) 4.893  5.000 54.840  < 2e-16 ***
## ---
## Signif. codes:  0 '***' 0.001 '**' 0.01 '*' 0.05 '.' 0.1 ' ' 1
##
## R-sq.(adj) =  0.129   Deviance explained = 13.2%
## -REML = 3272.3  Scale est. = 0.68292   n = 2649
plot(m2, select = 1, seWithMean = TRUE, unconditional = TRUE)
```

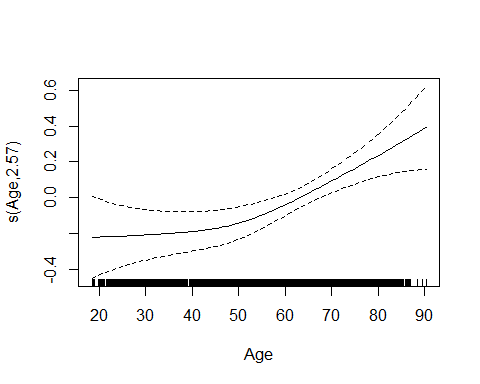

We can also try with a random smooth per study, allowing different wiggliness between studies. The overall effect now becomes linear, while many of the studies have significant deviations from the overall trend.

```
form <- reformulate("s(Age) + s(Age, by = Study, bs = 'ts') + Sex + s(Study, bs = 're')",
                    response = psqi_var)
m3 <- gam(form, data = subset_data(data, psqi_var), method = "REML")
summary(m3)
##
## Family: gaussian
## Link function: identity
##
## Formula:
## PSQI_Comp4_Efficiency ~ s(Age) + s(Age, by = Study, bs = "ts") +
##     Sex + s(Study, bs = "re")
##
## Parametric coefficients:
##             Estimate Std. Error t value Pr(>|t|)
## (Intercept)  0.61595    0.12482   4.935 8.52e-07 ***
## SexMale     -0.17141    0.03467  -4.944 8.15e-07 ***
## ---
## Signif. codes:  0 '***' 0.001 '**' 0.01 '*' 0.05 '.' 0.1 ' ' 1
##
## Approximate significance of smooth terms:
##                                edf Ref.df      F p-value
## s(Age)                   2.7170941  3.404 18.405 1.2e-12 ***
## s(Age):StudyBarcelona    0.1919164  9.000  0.138  0.2629
## s(Age):StudyBASE-II      0.0001612  9.000  0.000  0.4115
## s(Age):StudyBetula       0.9039693  9.000  2.238  0.0043 **
## s(Age):StudyCam-CAN      0.5944460  9.000  0.275  0.1167
## s(Age):StudyLCBC         0.0007512  9.000  0.000  0.8021
## s(Age):StudyWhitehall-II 0.0011725  9.000  0.000  0.7715
## s(Study)                 4.8518291  5.000 46.743 < 2e-16 ***
## ---
## Signif. codes:  0 '***' 0.001 '**' 0.01 '*' 0.05 '.' 0.1 ' ' 1
##
## R-sq.(adj) =  0.129   Deviance explained = 13.2%
## -REML =   3274  Scale est. = 0.68304   n = 2649
plot(m3, pages = 1, seWithMean = TRUE, unconditional = TRUE)
```

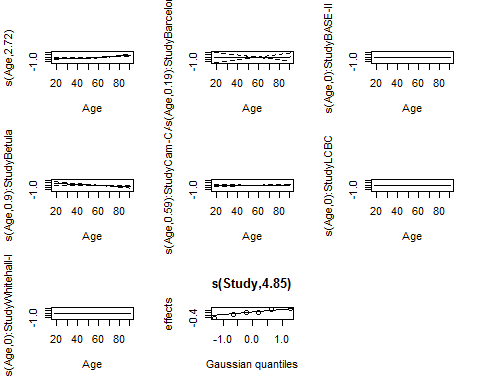

We can compare the models:

```
anova(m0, m1, m2, m3)
## Analysis of Deviance Table
##
## Model 1: PSQI_Comp4_Efficiency ~ Age + Sex + s(Study, bs = "re")
## Model 2: PSQI_Comp4_Efficiency ~ s(Age) + Sex + s(Study, bs = "re")
## Model 3: PSQI_Comp4_Efficiency ~ s(Age) + Sex + s(Age, Study, bs = "re")
## Model 4: PSQI_Comp4_Efficiency ~ s(Age) + s(Age, by = Study, bs = "ts") +
##     Sex + s(Study, bs = "re")
##   Resid. Df Resid. Dev        Df Deviance
## 1    2641.0     1817.8
## 2    2638.1     1809.7  2.821554   8.1374
## 3    2638.2     1802.6 -0.086697   7.1043
## 4    2635.1     1801.7  3.146057   0.9345
```

Random slope and smooth function of age seems justified here, but random smooths is too much, so model m2 is selected.

```
AIC(m0, m1, m2, m3)
##           df      AIC
## m0  8.929553 6537.969
## m1 11.069701 6530.365
## m2 11.024363 6519.855
## m3 13.591928 6523.616
```

We conclude that sleep efficiency decreases with age.

#### PSQI Component 5: Problems

psqi_var <- "PSQI_Comp5_Problems"

Inspection of data.

desc_plot(data, psqi_var)

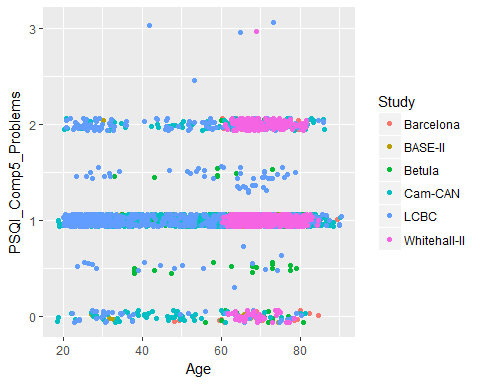

Linear model with random intercept per study.

```
form <- reformulate("Age + Sex + s(Study, bs = 're')", response = psqi_var)
m0 <- gam(form, data = subset_data(data, psqi_var), method = "REML")
summary(m0)
##
## Family: gaussian
## Link function: identity
##
## Formula:
## PSQI_Comp5_Problems ~ Age + Sex + s(Study, bs = "re")
##
## Parametric coefficients:
##               Estimate Std. Error t value Pr(>|t|)
## (Intercept)  0.9324703  0.0475052  19.629  < 2e-16 ***
## Age          0.0021195  0.0005051   4.196  2.8e-05 ***
## SexMale     -0.0446096  0.0172855  -2.581  0.00991 **
## ---
## Signif. codes:  0 '***' 0.001 '**' 0.01 '*' 0.05 '.' 0.1 ' ' 1
##
## Approximate significance of smooth terms:
##            edf Ref.df     F p-value
## s(Study) 4.536      5 16.12  <2e-16 ***
## ---
## Signif. codes:  0 '***' 0.001 '**' 0.01 '*' 0.05 '.' 0.1 ' ' 1
##
## R-sq.(adj) =  0.0315   Deviance explained = 3.37%
## -REML =   1666  Scale est. = 0.18327   n = 2887
```

Next we try a model with a smooth term for Age. The estimate is almost linear.

```
form <- reformulate("s(Age) + Sex + s(Study, bs = 're')", response = psqi_var)
m1 <- gam(form, data = subset_data(data, psqi_var), method = "REML")
summary(m1)
##
## Family: gaussian
## Link function: identity
##
## Formula:
## PSQI_Comp5_Problems ~ s(Age) + Sex + s(Study, bs = "re")
##
## Parametric coefficients:
##             Estimate Std. Error t value Pr(>|t|)
## (Intercept)  1.05573    0.03694  28.576  < 2e-16 ***
## SexMale     -0.04461    0.01729  -2.581  0.00991 **
## ---
## Signif. codes:  0 '***' 0.001 '**' 0.01 '*' 0.05 '.' 0.1 ' ' 1
##
## Approximate significance of smooth terms:
##            edf Ref.df     F  p-value
## s(Age)   1.001  1.001 17.59 2.81e-05 ***
## s(Study) 4.536  5.000 16.12  < 2e-16 ***
## ---
## Signif. codes:  0 '***' 0.001 '**' 0.01 '*' 0.05 '.' 0.1 ' ' 1
##
## R-sq.(adj) =  0.0315   Deviance explained = 3.37%
## -REML = 1663.1  Scale est. = 0.18327   n = 2887
plot(m1, select = 1, seWithMean = TRUE, unconditional = TRUE)
```

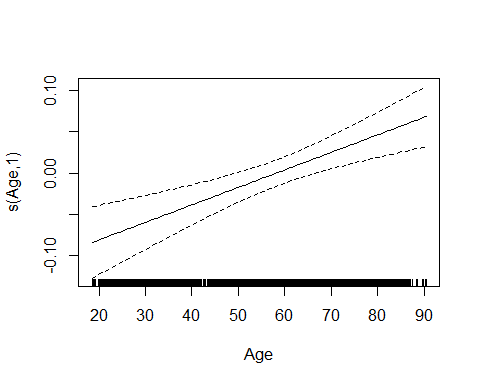

Try with a random slope per study, and linear main effect, since the smooths have all been linear thus far.

```
form <- reformulate("Age + Sex + s(Age, Study, bs = 're')", response = psqi_var)
m2 <- gam(form, data = subset_data(data, psqi_var), method = "REML")
summary(m2)
##
## Family: gaussian
## Link function: identity
##
## Formula:
## PSQI_Comp5_Problems ~ Age + Sex + s(Age, Study, bs = "re")
##
## Parametric coefficients:
##               Estimate Std. Error t value Pr(>|t|)
## (Intercept)  0.9883171  0.0281061  35.164  < 2e-16 ***
## Age          0.0013086  0.0007132   1.835  0.06662 .
## SexMale     -0.0489567  0.0172421  -2.839  0.00455 **
## ---
## Signif. codes:  0 '***' 0.001 '**' 0.01 '*' 0.05 '.' 0.1 ' ' 1
##
## Approximate significance of smooth terms:
##                edf Ref.df     F p-value
## s(Age,Study) 4.553      5 15.83  <2e-16 ***
## ---
## Signif. codes:  0 '***' 0.001 '**' 0.01 '*' 0.05 '.' 0.1 ' ' 1
##
## R-sq.(adj) =  0.031   Deviance explained = 3.32%
## -REML = 1666.8  Scale est. = 0.18336   n = 2887
```

We can compare the models:

```
anova(m0, m1, m2)
## Analysis of Deviance Table
##
## Model 1: PSQI_Comp5_Problems ~ Age + Sex + s(Study, bs = "re")
## Model 2: PSQI_Comp5_Problems ~ s(Age) + Sex + s(Study, bs = "re")
## Model 3: PSQI_Comp5_Problems ~ Age + Sex + s(Age, Study, bs = "re")
##   Resid. Df Resid. Dev        Df  Deviance
## 1    2878.9     527.71
## 2    2878.9     527.71  0.010278  0.000083
## 3    2878.9     527.97 -0.012986 -0.258244
```

We choose a linear model with random intercept here, although the differences are not big. Looking at the regression coefficients of m0, we see that reported sleep problems increase with age.

```
AIC(m0, m1, m2)
##          df      AIC
## m0 8.692983 3304.105
## m1 8.702658 3304.124
## m2 8.701990 3305.535
```

#### PSQI Component 6: Medication

psqi_var <- "PSQI_Comp6_Medication"

Inspection of data.

desc_plot(data, psqi_var)

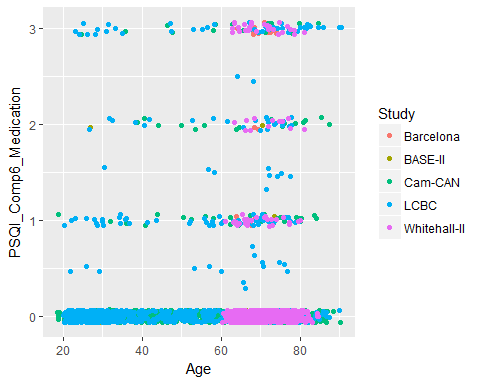

Linear model with random intercept per study.

```
form <- reformulate("Age + Sex + s(Study, bs = 're')", response = psqi_var)
m0 <- gam(form, data = subset_data(data, psqi_var), method = "REML")
summary(m0)
##
## Family: gaussian
## Link function: identity
##
## Formula:
## PSQI_Comp6_Medication ~ Age + Sex + s(Study, bs = "re")
##
## Parametric coefficients:
##              Estimate Std. Error t value Pr(>|t|)
## (Intercept) -0.090151   0.137873  -0.654 0.513255
## Age          0.007613   0.000833   9.140  < 2e-16 ***
## SexMale     -0.113619   0.029525  -3.848 0.000122 ***
## ---
## Signif. codes:  0 '***' 0.001 '**' 0.01 '*' 0.05 '.' 0.1 ' ' 1
##
## Approximate significance of smooth terms:
##            edf Ref.df     F p-value
## s(Study) 3.878      4 22.79  <2e-16 ***
## ---
## Signif. codes:  0 '***' 0.001 '**' 0.01 '*' 0.05 '.' 0.1 ' ' 1
##
## R-sq.(adj) =  0.0677   Deviance explained = 6.99%
## -REML = 2583.9  Scale est. = 0.45102   n = 2513
```

Next we try a model with a smooth term for Age.

```
form <- reformulate("s(Age) + Sex + s(Study, bs = 're')", response = psqi_var)
m1 <- gam(form, data = subset_data(data, psqi_var), method = "REML")
summary(m1)
##
## Family: gaussian
## Link function: identity
##
## Formula:
## PSQI_Comp6_Medication ~ s(Age) + Sex + s(Study, bs = "re")
##
## Parametric coefficients:
##             Estimate Std. Error t value Pr(>|t|)
## (Intercept)  0.34579    0.12893   2.682 0.007365 **
## SexMale     -0.11362    0.02953  -3.848 0.000122 ***
## ---
## Signif. codes:  0 '***' 0.001 '**' 0.01 '*' 0.05 '.' 0.1 ' ' 1
##
## Approximate significance of smooth terms:
##            edf Ref.df     F p-value
## s(Age)   1.020   1.04 79.78  <2e-16 ***
## s(Study) 3.878   4.00 22.78  <2e-16 ***
## ---
## Signif. codes:  0 '***' 0.001 '**' 0.01 '*' 0.05 '.' 0.1 ' ' 1
##
## R-sq.(adj) =  0.0677   Deviance explained = 6.99%
## -REML =   2581  Scale est. = 0.45102   n = 2513
plot(m1, select = 1, seWithMean = TRUE, unconditional = TRUE)
```

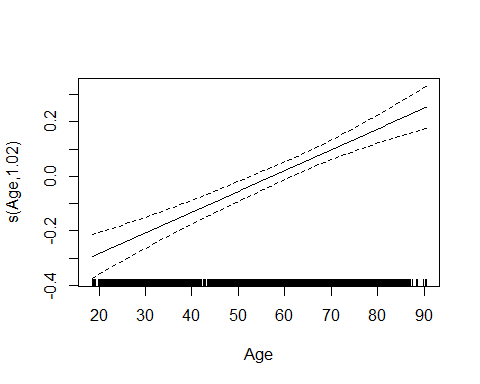

Try with a random slope per study, and linear overall effect.

```
form <- reformulate("Age + Sex + s(Age, Study, bs = 're')", response = psqi_var)
m2 <- gam(form, data = subset_data(data, psqi_var), method = "REML")
summary(m2)
##
## Family: gaussian
## Link function: identity
##
## Formula:
## PSQI_Comp6_Medication ~ Age + Sex + s(Age, Study, bs = "re")
##
## Parametric coefficients:
##              Estimate Std. Error t value Pr(>|t|)
## (Intercept) -0.102951   0.045464  -2.264 0.023630 *
## Age          0.007682   0.002096   3.666 0.000252 ***
## SexMale     -0.112485   0.029351  -3.832 0.000130 ***
## ---
## Signif. codes:  0 '***' 0.001 '**' 0.01 '*' 0.05 '.' 0.1 ' ' 1
##
## Approximate significance of smooth terms:
##                edf Ref.df     F p-value
## s(Age,Study) 3.883      4 26.34  <2e-16 ***
## ---
## Signif. codes:  0 '***' 0.001 '**' 0.01 '*' 0.05 '.' 0.1 ' ' 1
##
## R-sq.(adj) =  0.0728   Deviance explained =  7.5%
## -REML = 2577.1  Scale est. = 0.44857   n = 2513
```

We can compare the models:

```
anova(m0, m1, m2)
## Analysis of Deviance Table
##
## Model 1: PSQI_Comp6_Medication ~ Age + Sex + s(Study, bs = "re")
## Model 2: PSQI_Comp6_Medication ~ s(Age) + Sex + s(Study, bs = "re")
## Model 3: PSQI_Comp6_Medication ~ Age + Sex + s(Age, Study, bs = "re")
##   Resid. Df Resid. Dev        Df Deviance
## 1    2505.9     1130.3
## 2    2505.8     1130.3  0.059577   0.0161
## 3    2505.9     1124.2 -0.062914   6.1202
```

Linear model with random slope seems justified.

```
AIC(m0, m1, m2)
##          df      AIC
## m0 7.994003 5139.731
## m1 8.033832 5139.774
## m2 7.994830 5126.052
```

#### PSQI Component 7: Tiredness

psqi_var <- "PSQI_Comp7_Tired"

Inspection of data.

desc_plot(data, psqi_var)

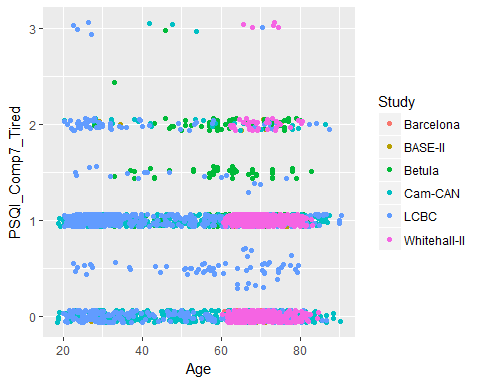

Linear model with random intercept per study.

```
form <- reformulate("Age + Sex + s(Study, bs = 're')", response = psqi_var)
m0 <- gam(form, data = subset_data(data, psqi_var), method = "REML")
summary(m0)
##
## Family: gaussian
## Link function: identity
##
## Formula:
## PSQI_Comp7_Tired ~ Age + Sex + s(Study, bs = "re")
##
## Parametric coefficients:
##               Estimate Std. Error t value Pr(>|t|)
## (Intercept)  1.0050167  0.1335618   7.525 7.07e-14 ***
## Age         -0.0052132  0.0006804  -7.662 2.50e-14 ***
## SexMale     -0.0103829  0.0234048  -0.444    0.657
## ---
## Signif. codes:  0 '***' 0.001 '**' 0.01 '*' 0.05 '.' 0.1 ' ' 1
##
## Approximate significance of smooth terms:
##            edf Ref.df     F p-value
## s(Study) 4.912      5 79.44  <2e-16 ***
## ---
## Signif. codes:  0 '***' 0.001 '**' 0.01 '*' 0.05 '.' 0.1 ' ' 1
##
## R-sq.(adj) =  0.136   Deviance explained = 13.9%
## -REML = 2433.7  Scale est. = 0.32513   n = 2813
```

Next we try a model with a smooth term for Age.

```
form <- reformulate("s(Age) + Sex + s(Study, bs = 're')", response = psqi_var)
m1 <- gam(form, data = subset_data(data, psqi_var), method = "REML")
summary(m1)
##
## Family: gaussian
## Link function: identity
##
## Formula:
## PSQI_Comp7_Tired ~ s(Age) + Sex + s(Study, bs = "re")
##
## Parametric coefficients:
##              Estimate Std. Error t value Pr(>|t|)
## (Intercept)  0.709673   0.126370   5.616 2.15e-08 ***
## SexMale     -0.008584   0.023359  -0.367    0.713
## ---
## Signif. codes:  0 '***' 0.001 '**' 0.01 '*' 0.05 '.' 0.1 ' ' 1
##
## Approximate significance of smooth terms:
##            edf Ref.df     F  p-value
## s(Age)   4.130  5.147 13.99 9.41e-14 ***
## s(Study) 4.909  5.000 80.93  < 2e-16 ***
## ---
## Signif. codes:  0 '***' 0.001 '**' 0.01 '*' 0.05 '.' 0.1 ' ' 1
##
## R-sq.(adj) =  0.141   Deviance explained = 14.4%
## -REML = 2427.6  Scale est. = 0.32358   n = 2813
plot(m1, select = 1, seWithMean = TRUE, unconditional = TRUE)
```

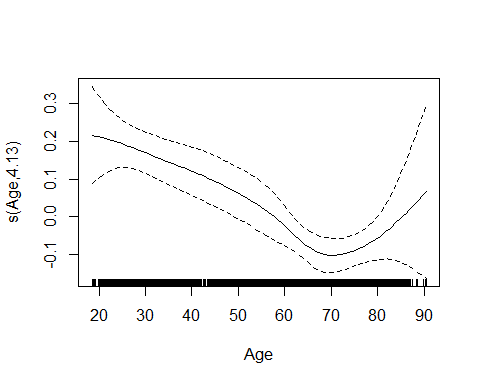

Try with a random slope per study.

```
form <- reformulate("s(Age) + Sex + s(Age, Study, bs = 're')", response = psqi_var)
m2 <- gam(form, data = subset_data(data, psqi_var), method = "REML")
summary(m2)
##
## Family: gaussian
## Link function: identity
##
## Formula:
## PSQI_Comp7_Tired ~ s(Age) + Sex + s(Age, Study, bs = "re")
##
## Parametric coefficients:
##             Estimate Std. Error t value Pr(>|t|)
## (Intercept)  0.70273    0.11122   6.318 3.07e-10 ***
## SexMale     -0.00917    0.02338  -0.392    0.695
## ---
## Signif. codes:  0 '***' 0.001 '**' 0.01 '*' 0.05 '.' 0.1 ' ' 1
##
## Approximate significance of smooth terms:
##                edf Ref.df     F p-value
## s(Age)       4.559  5.647  3.46 0.00254 **
## s(Age,Study) 4.909  5.000 76.57 < 2e-16 ***
## ---
## Signif. codes:  0 '***' 0.001 '**' 0.01 '*' 0.05 '.' 0.1 ' ' 1
##
## R-sq.(adj) =  0.135   Deviance explained = 13.8%
## -REML = 2436.9  Scale est. = 0.3256    n = 2813
plot(m2, select = 1, seWithMean = TRUE, unconditional = TRUE)
```

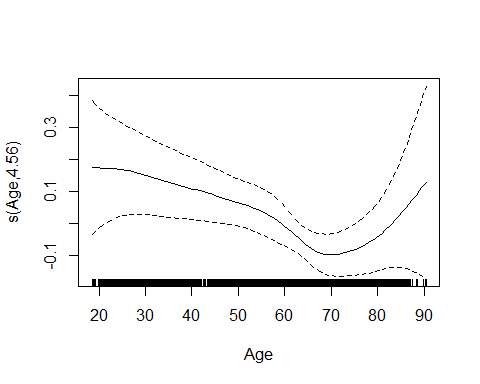

We can compare the models:

```
anova(m0, m1, m2)
## Analysis of Deviance Table
##
## Model 1: PSQI_Comp7_Tired ~ Age + Sex + s(Study, bs = "re")
## Model 2: PSQI_Comp7_Tired ~ s(Age) + Sex + s(Study, bs = "re")
## Model 3: PSQI_Comp7_Tired ~ s(Age) + Sex + s(Age, Study, bs = "re")
##   Resid. Df Resid. Dev      Df Deviance
## 1    2805.0     912.02
## 2    2799.8     906.67 5.14041   5.3466
## 3    2799.2     912.17 0.64202  -5.5053
```

The smooth model with random intercept seems sufficient.

```
AIC(m0, m1, m2)
##           df      AIC
## m0  8.952514 4832.431
## m1 13.073663 4824.134
## m2 13.644228 4842.304
```

Younger people feel more tired than older people, but the feeling of tiredness seems to increase again after the age of 70.

#### PSQI Global

psqi_var <- "PSQI_Global"

Inspection of data.

desc_plot(data, psqi_var)

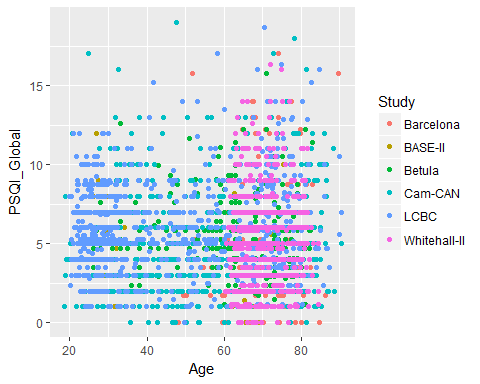

Linear model with random intercept per study.

```
form <- reformulate("Age + Sex + s(Study, bs = 're')", response = psqi_var)
m0 <- gam(form, data = subset_data(data, psqi_var), method = "REML")
summary(m0)
##
## Family: gaussian
## Link function: identity
##
## Formula:
## PSQI_Global ~ Age + Sex + s(Study, bs = "re")
##
## Parametric coefficients:
##              Estimate Std. Error t value Pr(>|t|)
## (Intercept)  4.757824   0.278703  17.071  < 2e-16 ***
## Age          0.013449   0.003354   4.010 6.23e-05 ***
## SexMale     -0.761294   0.114790  -6.632 3.95e-11 ***
## ---
## Signif. codes:  0 '***' 0.001 '**' 0.01 '*' 0.05 '.' 0.1 ' ' 1
##
## Approximate significance of smooth terms:
##            edf Ref.df     F  p-value
## s(Study) 4.251      5 10.68 6.79e-12 ***
## ---
## Signif. codes:  0 '***' 0.001 '**' 0.01 '*' 0.05 '.' 0.1 ' ' 1
##
## R-sq.(adj) =  0.0418   Deviance explained = 4.39%
## -REML = 6995.4  Scale est. = 7.9633    n = 2843
```

Next we try a model with a smooth term for Age.

```
form <- reformulate("s(Age) + Sex + s(Study, bs = 're')", response = psqi_var)
m1 <- gam(form, data = subset_data(data, psqi_var), method = "REML")
summary(m1)
##
## Family: gaussian
## Link function: identity
##
## Formula:
## PSQI_Global ~ s(Age) + Sex + s(Study, bs = "re")
##
## Parametric coefficients:
##             Estimate Std. Error t value Pr(>|t|)
## (Intercept)   5.5498     0.1902   29.17  < 2e-16 ***
## SexMale      -0.7596     0.1146   -6.63 4.02e-11 ***
## ---
## Signif. codes:  0 '***' 0.001 '**' 0.01 '*' 0.05 '.' 0.1 ' ' 1
##
## Approximate significance of smooth terms:
##            edf Ref.df     F  p-value
## s(Age)   2.593  3.249 7.741 2.61e-05 ***
## s(Study) 4.188  5.000 9.304 1.41e-10 ***
## ---
## Signif. codes:  0 '***' 0.001 '**' 0.01 '*' 0.05 '.' 0.1 ' ' 1
##
## R-sq.(adj) =  0.0448   Deviance explained = 4.74%
## -REML = 6989.3  Scale est. = 7.9388    n = 2843
plot(m1, pages = 1, seWithMean = TRUE, unconditional = TRUE)
```

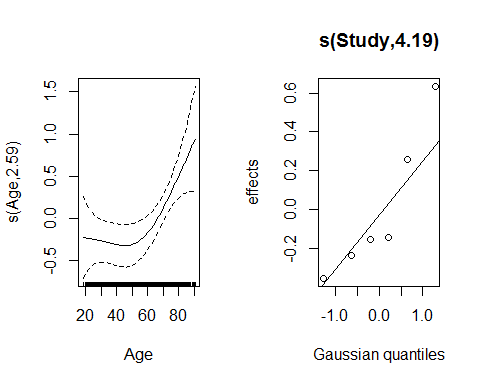

Try with a random slope per study.

```
form <- reformulate("s(Age) + Sex + s(Age, Study, bs = 're')", response = psqi_var)
m2 <- gam(form, data = subset_data(data, psqi_var), method = "REML")
summary(m2)
##
## Family: gaussian
## Link function: identity
##
## Formula:
## PSQI_Global ~ s(Age) + Sex + s(Age, Study, bs = "re")
##
## Parametric coefficients:
##             Estimate Std. Error t value Pr(>|t|)
## (Intercept)   5.5851     0.1861  30.009  < 2e-16 ***
## SexMale      -0.7679     0.1145  -6.708 2.38e-11 ***
## ---
## Signif. codes:  0 '***' 0.001 '**' 0.01 '*' 0.05 '.' 0.1 ' ' 1
##
## Approximate significance of smooth terms:
##                edf Ref.df     F  p-value
## s(Age)       2.787  3.491 4.914  0.00129 **
## s(Age,Study) 4.295  5.000 8.916 7.22e-10 ***
## ---
## Signif. codes:  0 '***' 0.001 '**' 0.01 '*' 0.05 '.' 0.1 ' ' 1
##
## R-sq.(adj) =  0.0443   Deviance explained =  4.7%
## -REML = 6990.5  Scale est. = 7.9423    n = 2843
plot(m2, select = 1)
```

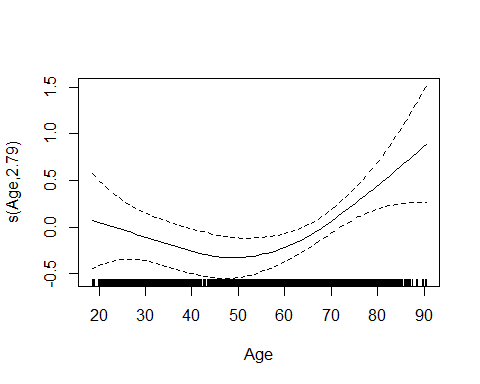

We can compare the models:

```
anova(m0, m1, m2)
## Analysis of Deviance Table
##
## Model 1: PSQI_Global ~ Age + Sex + s(Study, bs = "re")
## Model 2: PSQI_Global ~ s(Age) + Sex + s(Study, bs = "re")
## Model 3: PSQI_Global ~ s(Age) + Sex + s(Age, Study, bs = "re")
##   Resid. Df Resid. Dev      Df Deviance
## 1    2834.9      22582
## 2    2832.2      22500 2.73412   81.772
## 3    2832.0      22508 0.20074   -7.676
AIC(m0, m1, m2)
##           df      AIC
## m0  8.501515 13976.61
## m1 10.544628 13970.39
## m2 10.761798 13971.79
```

We select model m1. The amount of overall sleep trouble seems stable or slightly decreasing until the age of around 50, and then increases steadily with age.

### Grid Plots

```
grid <- tibble(
  Age = seq(from = 18, to = 90, by = 1),
  Study = "LCBC",
  Sex = "Female"
)

df <- imap_dfr(models, function(x, y){
  term <- if_else("Age" %in% attr(x$pterms, "term.labels"), "Age", "s(Age)")
  pred <- predict(x, newdata = grid, type = "iterms", se.fit = TRUE, terms = term)
  pred$fit <- pred$fit + attr(pred, "constant")
  grid %>%
    bind_cols(pred) %>%
    mutate(
      PSQI = y,
      ymin = fit - 2 * se.fit,
      ymax = fit + 2 * se.fit
    )
})


p <- ggplot(df, aes(x = Age, ymin = pmax(0, ymin - 0.5), ymax = ymax + 0.5)) +
  geom_line(aes(y = fit)) +
  geom_line(aes(y = ymin), linetype = "dashed") +
  geom_line(aes(y = ymax), linetype = "dashed") +
  facet_wrap(~PSQI, nrow = 4, ncol = 2, scales = "free_y") +
  xlab("") +
  ylab("") +
  theme_classic() +
  theme(strip.background = element_blank())

p
```

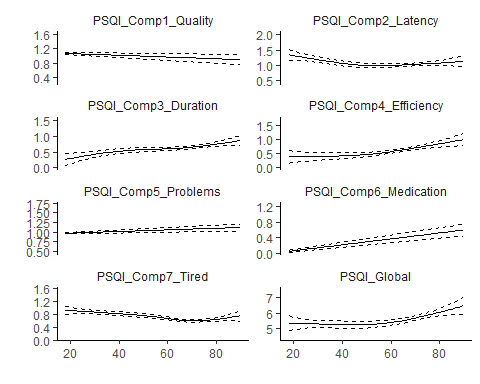

ggsave("age_grid_plot.png", plot = p, dpi = 600, height = 9, width = 4.5, units = "in")
