## Supplementary material for "Self-reported sleep relates to hippocampal atrophy across the adult lifespan – results from the Lifebrain consortium": SI

Effect of Sleep on Hippocampus Volume

```
library(dplyr)
library(ggplot2)
library(purrr)
library(tidyr)
library(mgcv)
library(cowplot)
## Warning: package 'cowplot' was built under R version 3.5.3
library(itsadug)
full_data <- readRDS("./data/full_data.rds")
```

### Effect of Sleep on Hippocampus

The data consist of (repeated) measurements of participants within multiple studies. We therefore used a nested random effect structure. The random intercept form is specified as list(Study =~ 1, ID =~ 1) with mgcv, where nesting goes from left to right. Hence, correlation among the repeated measurements of a single participant is captured by the term ID =~ 1, and correlation among all the measurements in a given study, when compared to another study, is captured by the term Study =~ 1.

The following defines a function for subsetting the whole dataset, in order to obtain answers to a particular PSQI variable. It will be used repeatedly in the analyses below.

```
subset_data <- function(data, psqi_var){
  data %>%
    filter(key == psqi_var) %>%
    rename(!! psqi_var := value) %>%
    mutate(Sex_ord = as.ordered(Sex))
}
```

We start with a data preparation for this analysis. Standardized Hippocampus volume and ICV are added, with suffix _st.

```
data <- full_data %>%
  select(Age, Age_bl, dt, contains("PSQI"), ICV, Hippocampus, Sex, ID, Study, Site_Name) %>%
  mutate_at(vars(ICV, Hippocampus), funs(st = (. - mean(.))/sd(.))) %>%
  gather(key = "key", value = "value", contains("PSQI"), na.rm = TRUE) %>%
  na.omit() %>%
  left_join(select(full_data, ID, dt, Depression, BMI, Education), by = c("ID", "dt"))
```

Function for plotting PSQI vs Hippocampus.

```
psqi_plot <- function(data, psqi_var){
  data %>%
    filter(key == !! psqi_var) %>%
    ggplot(aes(x = value, y = Hippocampus)) +
    geom_jitter(aes(color = Study)) +
    xlab(psqi_var)
}
```

As a part of the exploratory analysis, we fit separate models for each study. Keeping the PSQI variable as a linear term here, in order to compare confidence intervals across studies. When having a random effect per ID, I use the gamm() and list form, for the sake of speed. Here is a function that does this.

```
sep_fits <- function(data, psqi_var) {
  data %>%
    subset_data(psqi_var) %>%
    mutate(Study = factor(as.character(Study))) %>%
    split(.$Study) %>%
    map_dfr(function(x) {
      form <- reformulate(paste("s(Age, bs = 'cr') +", psqi_var, "+ ICV_st + Sex"),
                          response = "Hippocampus_st")
      # Check if there are repeated measurements
      rep_check <- x %>%
        group_by(ID) %>%
        count() %>%
        filter(n > 1)

      if(nrow(rep_check) > 0){
        rand = list(ID =~ 1)
      } else {
        rand = NULL
      }

      fit <- gamm(form, data = x, random = rand)$gam
      tibble(
        fit = coefficients(fit)[[psqi_var]],
        fit_se = sqrt(vcov(!! fit)[psqi_var, psqi_var])
        )
      }, .id = "Study")  %>%
    ggplot(aes(x = Study, y = fit, ymin = fit - 2 * fit_se,
               ymax = fit + 2 * fit_se)) +
    geom_point(size = 2) +
    geom_errorbar(alpha = 0.2, size = 1) +
    geom_hline(yintercept = 0, linetype = "dashed") +
    ylab(paste("Effect of", psqi_var, "on hippocampus volume")) +
    ggtitle("Point Estimates and 95 % Confidence Intervals")
  }
```

#### Preparation

##### Effect of Age on Hippocampus

```
ggplot(data, aes(x = Age, y = Hippocampus, group = Study, color = Study)) +
  geom_point(alpha = 0.1) +
  geom_line(aes(group = ID))
```

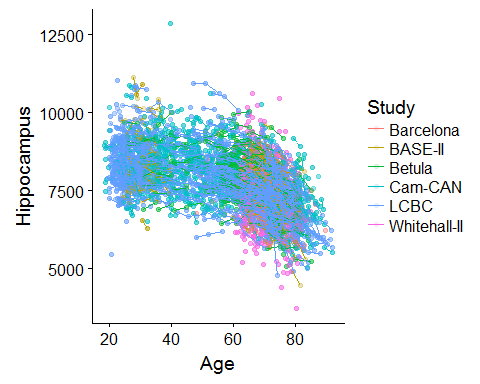

In order to avoid exploring a wide range of different models for the effect of age on hippocampus volume when fitting each PSQI component, I start by exploring various fits her.

Random intercept per study and ID:

```
m0 <- gamm(Hippocampus_st ~ s(Age, bs = "cr") + ICV_st + Sex,
           random = list(Study =~ 1, ID =~ 1),
           data = distinct(select(data, Hippocampus_st, ICV_st, Age, Sex, Study, ID)),
           method = "REML")

saveRDS(m0, "./models/age_hippo_m0.rds")
m0 <- readRDS("./models/age_hippo_m0.rds")
summary(m0$gam)
##
## Family: gaussian
## Link function: identity
##
## Formula:
## Hippocampus_st ~ s(Age, bs = "cr") + ICV_st + Sex
##
## Parametric coefficients:
##             Estimate Std. Error t value Pr(>|t|)
## (Intercept) -0.15146    0.07247  -2.090   0.0367 *
## ICV_st       0.16967    0.01609  10.546   <2e-16 ***
## SexMale      0.32994    0.03528   9.352   <2e-16 ***
## ---
## Signif. codes:  0 '***' 0.001 '**' 0.01 '*' 0.05 '.' 0.1 ' ' 1
##
## Approximate significance of smooth terms:
##          edf Ref.df     F p-value
## s(Age) 7.257  7.257 374.5  <2e-16 ***
## ---
## Signif. codes:  0 '***' 0.001 '**' 0.01 '*' 0.05 '.' 0.1 ' ' 1
##
## R-sq.(adj) =  0.411
##   Scale est. = 0.018902  n = 5075
```

Random slope for age per study, and random intercept per ID:

```
m1 <- gamm(Hippocampus_st ~ s(Age, bs = "cr") + ICV_st + Sex,
           random = list(Study =~ Age, ID =~ 1),
           data = distinct(select(data, Hippocampus_st, ICV_st, Age, Sex, Study, ID)),
           method = "REML")
saveRDS(m1, "./models/age_hippo_m1.rds")
m1 <- readRDS("./models/age_hippo_m1.rds")
summary(m1$gam)
##
## Family: gaussian
## Link function: identity
##
## Formula:
## Hippocampus_st ~ s(Age, bs = "cr") + ICV_st + Sex
##
## Parametric coefficients:
##             Estimate Std. Error t value Pr(>|t|)
## (Intercept) -0.15893    0.06075  -2.616  0.00892 **
## ICV_st       0.17073    0.01603  10.654  < 2e-16 ***
## SexMale      0.33410    0.03514   9.508  < 2e-16 ***
## ---
## Signif. codes:  0 '***' 0.001 '**' 0.01 '*' 0.05 '.' 0.1 ' ' 1
##
## Approximate significance of smooth terms:
##          edf Ref.df     F p-value
## s(Age) 7.242  7.242 98.93  <2e-16 ***
## ---
## Signif. codes:  0 '***' 0.001 '**' 0.01 '*' 0.05 '.' 0.1 ' ' 1
##
## R-sq.(adj) =  0.409
##   Scale est. = 0.018498  n = 5075
```

Including more complex random effects gave convergence problems, so we stop here. The fits from the two models are compared below. As these are almost indistinguishable, we go on with the random intercept model, m0.

```
tibble(
  p0 = as.numeric(predict(m0$gam, type = "iterms", terms = "s(Age)")),
  p1 = as.numeric(predict(m1$gam, type = "iterms", terms = "s(Age)")),
  Age = m1$gam$model$Age) %>%
  gather(key = "key", value = "value", -Age) %>%
  ggplot(aes(x = Age, y = value, group = key, color = key)) +
  geom_line()
```

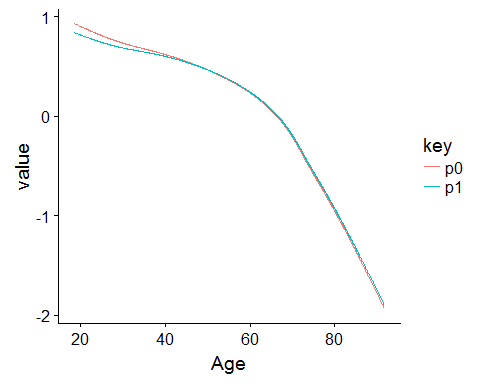

##### Effect of Baseline Age and Time on Hippocampus

For this purely longitudinal analysis, we keep only participants with at least two timepoints. Here is a function for performing this subsetting.

```
long_only <- function(data){
  data %>%
    group_by(ID) %>%
    filter(n() > 1) %>%
    ungroup()
}
```

Subsetting the data to keep only longitudinal.

```
long_data <- data %>%
  group_by(ID) %>%
  filter(key == first(key)) %>%
  long_only()
```

Random intercept of ID nested within Study.

```
m0 <- gamm(Hippocampus_st ~ te(dt, Age_bl, k = c(10, 3)) + ICV_st + Sex,
           random = list(Study =~ 1, ID =~ 1),
           data = long_data)
saveRDS(m0, "./models/long_age_hippo_m0.rds")
m0 <- readRDS("./models/long_age_hippo_m0.rds")
summary(m0$gam)
##
## Family: gaussian
## Link function: identity
##
## Formula:
## Hippocampus_st ~ te(dt, Age_bl, k = c(10, 3)) + ICV_st + Sex
##
## Parametric coefficients:
##             Estimate Std. Error t value Pr(>|t|)
## (Intercept) -0.04690    0.05224  -0.898 0.369367
## ICV_st       0.27069    0.02398  11.287  < 2e-16 ***
## SexMale      0.16805    0.04868   3.452 0.000563 ***
## ---
## Signif. codes:  0 '***' 0.001 '**' 0.01 '*' 0.05 '.' 0.1 ' ' 1
##
## Approximate significance of smooth terms:
##                 edf Ref.df     F p-value
## te(dt,Age_bl) 12.96  12.96 162.5  <2e-16 ***
## ---
## Signif. codes:  0 '***' 0.001 '**' 0.01 '*' 0.05 '.' 0.1 ' ' 1
##
## R-sq.(adj) =  0.458
##   Scale est. = 0.01873   n = 3275
fvisgam(m0$gam, view = c("Age_bl", "dt"))
## Summary:
##  * ICV_st : numeric predictor; set to the value(s): -0.046605225716453.
##  * Sex : factor; set to the value(s): Female.
##  * dt : numeric predictor; with 30 values ranging from 0.000000 to 10.976044.
##  * Age_bl : numeric predictor; with 30 values ranging from 19.333333 to 89.459274.
```

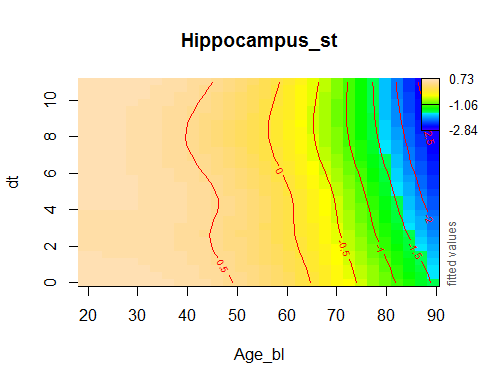

Random intercept of ID nested within Study, with random slope of baseline Age per study.

```
m1 <- gamm(Hippocampus_st ~ te(dt, Age_bl, k = c(10, 3)) + ICV_st + Sex,
           random = list(Study =~ Age_bl, ID =~ 1),
           data = long_data)

saveRDS(m1, "./models/long_age_hippo_m1.rds")
m1 <- readRDS("./models/long_age_hippo_m1.rds")

summary(m1$gam)
##
## Family: gaussian
## Link function: identity
##
## Formula:
## Hippocampus_st ~ te(dt, Age_bl, k = c(10, 3)) + ICV_st + Sex
##
## Parametric coefficients:
##             Estimate Std. Error t value Pr(>|t|)
## (Intercept) -0.05361    0.05128  -1.045 0.295906
## ICV_st       0.26712    0.02381  11.219  < 2e-16 ***
## SexMale      0.17629    0.04831   3.649 0.000268 ***
## ---
## Signif. codes:  0 '***' 0.001 '**' 0.01 '*' 0.05 '.' 0.1 ' ' 1
##
## Approximate significance of smooth terms:
##                 edf Ref.df     F p-value
## te(dt,Age_bl) 12.94  12.94 146.5  <2e-16 ***
## ---
## Signif. codes:  0 '***' 0.001 '**' 0.01 '*' 0.05 '.' 0.1 ' ' 1
##
## R-sq.(adj) =  0.459
##   Scale est. = 0.018728  n = 3275
fvisgam(m1$gam, view = c("dt", "Age_bl"))
## Summary:
##  * ICV_st : numeric predictor; set to the value(s): -0.046605225716453.
##  * Sex : factor; set to the value(s): Female.
##  * dt : numeric predictor; with 30 values ranging from 0.000000 to 10.976044.
##  * Age_bl : numeric predictor; with 30 values ranging from 19.333333 to 89.459274.
```

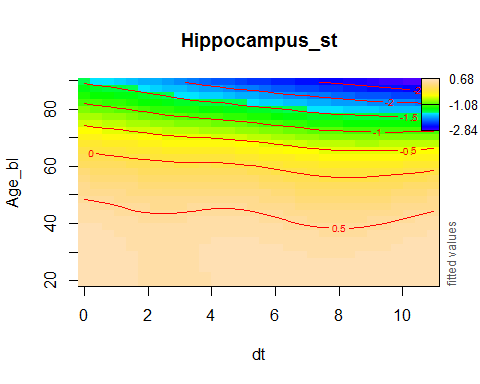

Here are effects of time, for four different ages. The fits from the two models are really indistinguishable. We hence go on with random intercept here as well.

```
crossing(Age_bl = c(20, 40, 60, 80),
         dt = seq(from = 0, to = 3, length.out = 100),
         ICV_st = 0,
         Sex = "Female",
         Study = "Betula",
         ID = "A") %>%
  bind_cols(
    m0 = predict(m0$gam, newdata = .),
    m1 = predict(m1$gam, newdata = .)
    ) %>%
  gather(key = "Model", value = "fit", m0, m1) %>%
  mutate(Age_bl = factor(Age_bl)) %>%
  group_by(Model, Age_bl) %>%
  arrange(dt, .by_group = TRUE) %>%
  mutate(fit = fit - first(fit)) %>%
  ungroup() %>%
  ggplot(aes(x = dt, y = fit, group = Model, color = Model)) +
  geom_line() +
  facet_wrap(~ Age_bl)
```

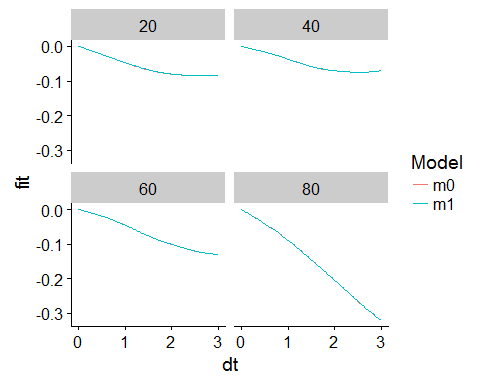

#### Function Definitions

Function for plotting interaction between PSQI and dt:

```
plot_psqi_interaction <- function(fit, psqi_var, psqi_range = 0:3, time_range = 0:3,
                                  normalize = TRUE, scales = "fixed", confint = FALSE){

  data <- crossing(dt = time_range,
         Age_bl = c(20, 50, 80),
         ICV_st = 0,
         Sex = "Female",
         Study = "Betula",
         ID = "A",
         value = psqi_range
         ) %>%
  mutate(!! psqi_var := value) %>%
  bind_cols(fit = predict(fit$gam, newdata = ., se.fit = TRUE)) %>%
  mutate_at(vars(value, Age_bl), factor) %>%
  group_by(value, Age_bl) %>%
  arrange(dt, .by_group = TRUE) %>%
  mutate(fit = fit - if_else(normalize, first(fit), 0)) %>%
  ungroup()


  p <- ggplot(data, aes(x = dt, y = fit, group = value, color = value)) +
    geom_line() +
    facet_wrap(~ paste("Age_bl =", Age_bl), scales = scales)

  if(confint){
    p + geom_ribbon(aes(ymin = fit - 2 * se.fit, ymax = fit + 2 * se.fit), alpha = 0.1)
  } else {
    p
  }

}
```

We define some functions for setting up formulas for the models, to avoid repetitive writing.

```
main_effects <- function(psqi_var, covariates = NULL){
  list(
    form = reformulate(
      paste(c("s(Age, k = 6, bs = 'cr')", paste("s(", psqi_var, ", k = 3, bs = 'cr')"),
            "ICV_st", "Sex", covariates), collapse = "+"),
      response = "Hippocampus_st"),
    rand = list(Study =~ 1, ID =~ 1)
    )
}

main_sex_interaction <- function(psqi_var, covariates = NULL){
  list(
    form = reformulate(
      paste(c("s(Age, k = 6, bs = 'cr')",
            paste("s(", psqi_var, ", k = 3, bs = 'cr')"),
            paste("s(", psqi_var, ", by = Sex_ord, k = 3, bs = 'cr')"),
            "ICV_st", "Sex_ord", covariates), collapse = "+"),
      response = "Hippocampus_st"),
    rand = list(Study =~ 1, ID =~ 1)
  )
}

age_interaction <- function(psqi_var, covariates = NULL){
  list(
    form = reformulate(
      paste(c("s(Age, k = 6, bs = 'cr')",
              paste("s(", psqi_var, ", bs = 'cr', k = 3)"),
              paste("ti(Age,", psqi_var, ", k = c(4, 3))"),
              "ICV_st", "Sex", covariates), collapse = "+"),
                    response = "Hippocampus_st"),
    rand = list(Study =~ 1, ID =~ 1)
  )
}

age_sex_interaction <- function(psqi_var, covariates = NULL){
  list(
    form = reformulate(
      paste(c("s(Age, k = 6, bs = 'cr')",
              "s(Age, by = Sex_ord, k = 3, bs = 'cr')",
              paste("s(", psqi_var, ", k = 3, bs = 'cr')"),
              paste("s(", psqi_var, ", by = Sex_ord, k = 3, bs = 'cr')"),
              paste("ti(Age,", psqi_var, ", k = c(4, 3))"),
              paste("ti(Age,", psqi_var, ", by = Sex_ord, k = c(4, 3))"),
              "ICV_st", "Sex_ord", covariates), collapse = "+"),
                    response = "Hippocampus_st"),
    rand = list(Study =~ 1, ID =~ 1)
  )
}

time_interaction <- function(psqi_var, covariates = NULL){
  list(
    form = reformulate(
      paste(c("te(dt, Age_bl, k = c(4, 10))",
              paste("ti(", psqi_var, ", k = 4)"),
              paste("ti(dt,", psqi_var, ", k = c(4, 4))"),
              "ICV_st", "Sex", covariates), collapse = "+"),
                    response = "Hippocampus_st"),
    rand = list(Study = ~ 1, ID = ~ 1)
  )
}

sex_time_interaction <- function(psqi_var, covariates = NULL){
  list(
    form = reformulate(
      paste(c("te(dt, Age_bl, k = c(4, 10))",
              "te(dt, Age_bl, by = Sex_ord, k = c(4, 10))",
              paste("ti(", psqi_var, ", k = 4)"),
              paste("ti(", psqi_var, ", by = Sex_ord, k = 4)"),
              paste("ti(dt,", psqi_var, ", k = c(4, 4))"),
              paste("ti(dt,", psqi_var, ", by = Sex_ord, k = c(4,4))"),
              "ICV_st", "Sex_ord", covariates), collapse = "+"),
                    response = "Hippocampus_st"),
    rand = list(Study = ~ 1, ID = ~ 1)
  )
}
```

#### PSQI Component 1: Sleep Quality

psqi_var <- "PSQI_Comp1_Quality"

psqi_plot(data, psqi_var)

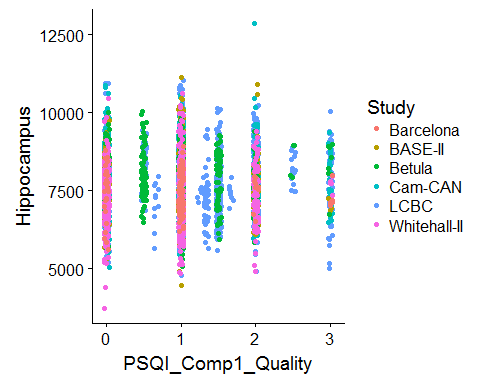

We fit one model per study first, purely for exploratory reasons.

sep_fits(data, psqi_var)

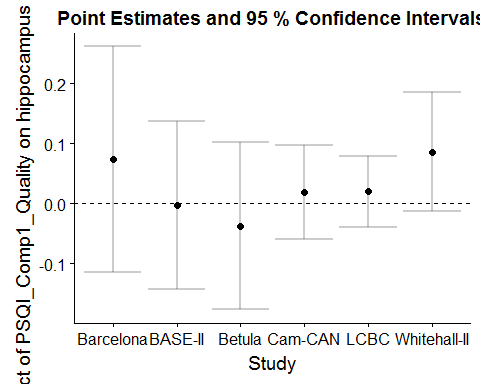

##### Main Effect of Sleep

GAMM with additive effects of Age and PSQI and random intercept of study and ID.

```
forms <- main_effects(psqi_var)
m0 <- gamm(forms[[1]], random = forms[[2]],
           data = subset_data(data, psqi_var))
saveRDS(m0, file = paste0("./models/", psqi_var, "_m0.rds"))
m0 <- readRDS(paste0("./models/", psqi_var, "_m0.rds"))
summary(m0$gam)
##
## Family: gaussian
## Link function: identity
##
## Formula:
## Hippocampus_st ~ s(Age, k = 6, bs = "cr") + s(PSQI_Comp1_Quality,
##     k = 3, bs = "cr") + ICV_st + Sex
##
## Parametric coefficients:
##             Estimate Std. Error t value Pr(>|t|)
## (Intercept) -0.14452    0.06649  -2.174   0.0298 *
## ICV_st       0.16913    0.01633  10.359   <2e-16 ***
## SexMale      0.32691    0.03599   9.084   <2e-16 ***
## ---
## Signif. codes:  0 '***' 0.001 '**' 0.01 '*' 0.05 '.' 0.1 ' ' 1
##
## Approximate significance of smooth terms:
##                         edf Ref.df       F p-value
## s(Age)                4.797  4.797 555.929  <2e-16 ***
## s(PSQI_Comp1_Quality) 1.000  1.000   2.205   0.138
## ---
## Signif. codes:  0 '***' 0.001 '**' 0.01 '*' 0.05 '.' 0.1 ' ' 1
##
## R-sq.(adj) =  0.411
##   Scale est. = 0.018901  n = 4965
plot(m0$gam, select = 2, seWithMean = TRUE)
```

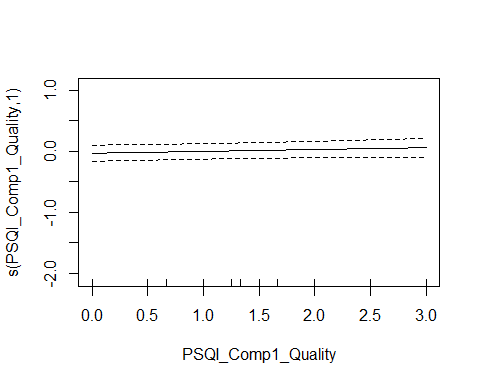

We also tried allowing for a random slope of the PSQI variable, but this led to non-convergence.

###### Controling for Covariates

```
forms <- main_effects(psqi_var, c("Depression", "BMI"))
m0_cov <- gamm(forms[[1]], random = forms[[2]],
           data = subset_data(data, psqi_var))
saveRDS(m0_cov, file = paste0("./models/", psqi_var, "_m0_cov.rds"))
m0_cov <- readRDS(paste0("./models/", psqi_var, "_m0_cov.rds"))
summary(m0_cov$gam)
##
## Family: gaussian
## Link function: identity
##
## Formula:
## Hippocampus_st ~ s(Age, k = 6, bs = "cr") + s(PSQI_Comp1_Quality,
##     k = 3, bs = "cr") + ICV_st + Sex + Depression + BMI
##
## Parametric coefficients:
##              Estimate Std. Error t value Pr(>|t|)
## (Intercept) -0.390965   0.165146  -2.367 0.017985 *
## ICV_st       0.083628   0.022259   3.757 0.000176 ***
## SexMale      0.356317   0.051226   6.956 4.39e-12 ***
## Depression   0.004108   0.021638   0.190 0.849443
## BMI          0.005425   0.005178   1.048 0.294881
## ---
## Signif. codes:  0 '***' 0.001 '**' 0.01 '*' 0.05 '.' 0.1 ' ' 1
##
## Approximate significance of smooth terms:
##                         edf Ref.df       F p-value
## s(Age)                4.507  4.507 264.621  <2e-16 ***
## s(PSQI_Comp1_Quality) 1.000  1.000   2.639   0.104
## ---
## Signif. codes:  0 '***' 0.001 '**' 0.01 '*' 0.05 '.' 0.1 ' ' 1
##
## R-sq.(adj) =  0.368
##   Scale est. = 0.023304  n = 2676
```

###### Interaction with Sex

We use Sex as an ordered factor, in order to get one smooth for Sex = "Female" and another smooth for the difference between Sex = "Female" and Sex = "Male". In this way we can test directly for the presence of an interaction.

The summary shows that no sex interaction is found.

```
forms <- main_sex_interaction(psqi_var)
m0s <- gamm(forms[[1]], random = forms[[2]],
           data = subset_data(data, psqi_var))

saveRDS(m0s, file = paste0("./models/", psqi_var, "_m0s.rds"))
m0s <- readRDS(paste0("./models/", psqi_var, "_m0s.rds"))
summary(m0s$gam)
##
## Family: gaussian
## Link function: identity
##
## Formula:
## Hippocampus_st ~ s(Age, k = 6, bs = "cr") + s(PSQI_Comp1_Quality,
##     k = 3, bs = "cr") + s(PSQI_Comp1_Quality, by = Sex_ord, k = 3,
##     bs = "cr") + ICV_st + Sex_ord
##
## Parametric coefficients:
##             Estimate Std. Error t value Pr(>|t|)
## (Intercept)  0.01889    0.06370   0.297    0.767
## ICV_st       0.16913    0.01633  10.359   <2e-16 ***
## Sex_ord.L    0.23114    0.02545   9.080   <2e-16 ***
## ---
## Signif. codes:  0 '***' 0.001 '**' 0.01 '*' 0.05 '.' 0.1 ' ' 1
##
## Approximate significance of smooth terms:
##                                     edf Ref.df       F p-value
## s(Age)                            4.797  4.797 555.768  <2e-16 ***
## s(PSQI_Comp1_Quality)             1.000  1.000   1.180   0.277
## s(PSQI_Comp1_Quality):Sex_ordMale 1.000  1.000   0.001   0.976
## ---
## Signif. codes:  0 '***' 0.001 '**' 0.01 '*' 0.05 '.' 0.1 ' ' 1
##
## R-sq.(adj) =  0.411
##   Scale est. = 0.018901  n = 4965
```

##### Interaction with Age

```
forms <- age_interaction(psqi_var)
m1 <- gamm(forms[[1]], random = forms[[2]],
           data = subset_data(data, psqi_var))

saveRDS(m1, file = paste0("./models/", psqi_var, "_m1.rds"))
m1 <- readRDS(paste0("./models/", psqi_var, "_m1.rds"))
summary(m1$gam)
##
## Family: gaussian
## Link function: identity
##
## Formula:
## Hippocampus_st ~ s(Age, k = 6, bs = "cr") + s(PSQI_Comp1_Quality,
##     bs = "cr", k = 3) + ti(Age, PSQI_Comp1_Quality, k = c(4,
##     3)) + ICV_st + Sex
##
## Parametric coefficients:
##             Estimate Std. Error t value Pr(>|t|)
## (Intercept) -0.14435    0.06646  -2.172   0.0299 *
## ICV_st       0.16922    0.01633  10.361   <2e-16 ***
## SexMale      0.32705    0.03599   9.086   <2e-16 ***
## ---
## Signif. codes:  0 '***' 0.001 '**' 0.01 '*' 0.05 '.' 0.1 ' ' 1
##
## Approximate significance of smooth terms:
##                              edf Ref.df       F p-value
## s(Age)                     4.797  4.797 556.001  <2e-16 ***
## s(PSQI_Comp1_Quality)      1.000  1.000   2.234   0.135
## ti(Age,PSQI_Comp1_Quality) 1.000  1.000   0.028   0.866
## ---
## Signif. codes:  0 '***' 0.001 '**' 0.01 '*' 0.05 '.' 0.1 ' ' 1
##
## R-sq.(adj) =  0.411
##   Scale est. = 0.018904  n = 4965
par(mfrow = c(1, 2))
plot(m1$gam, select = 2, seWithMean = TRUE)
fvisgam(m1$gam, view = c("Age", psqi_var))
## Summary:
##  * ICV_st : numeric predictor; set to the value(s): -0.017535583918877.
##  * Sex : factor; set to the value(s): Male.
##  * Age : numeric predictor; with 30 values ranging from 18.482192 to 91.885010.
##  * PSQI_Comp1_Quality : numeric predictor; with 30 values ranging from 0.000000 to 3.000000.
```

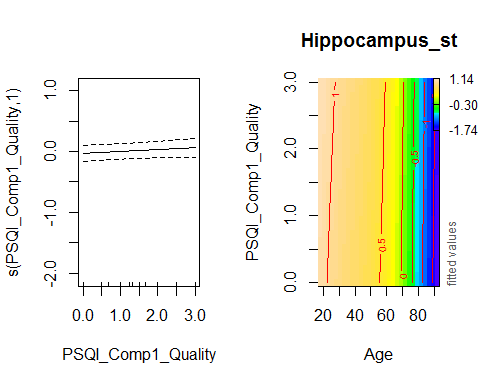

###### Controling for Covariates

```
forms <- age_interaction(psqi_var, c("Depression", "BMI"))
m1_cov <- gamm(forms[[1]], random = forms[[2]],
           data = subset_data(data, psqi_var))

saveRDS(m1_cov, file = paste0("./models/", psqi_var, "_m1_cov.rds"))
m1_cov <- readRDS(paste0("./models/", psqi_var, "_m1_cov.rds"))
summary(m1_cov$gam)
##
## Family: gaussian
## Link function: identity
##
## Formula:
## Hippocampus_st ~ s(Age, k = 6, bs = "cr") + s(PSQI_Comp1_Quality,
##     bs = "cr", k = 3) + ti(Age, PSQI_Comp1_Quality, k = c(4,
##     3)) + ICV_st + Sex + Depression + BMI
##
## Parametric coefficients:
##              Estimate Std. Error t value Pr(>|t|)
## (Intercept) -0.391626   0.165493  -2.366 0.018032 *
## ICV_st       0.083157   0.022271   3.734 0.000193 ***
## SexMale      0.355633   0.051241   6.940 4.89e-12 ***
## Depression   0.004950   0.021681   0.228 0.819429
## BMI          0.005416   0.005178   1.046 0.295636
## ---
## Signif. codes:  0 '***' 0.001 '**' 0.01 '*' 0.05 '.' 0.1 ' ' 1
##
## Approximate significance of smooth terms:
##                              edf Ref.df       F p-value
## s(Age)                     4.505  4.505 261.984  <2e-16 ***
## s(PSQI_Comp1_Quality)      1.000  1.000   2.661   0.103
## ti(Age,PSQI_Comp1_Quality) 1.000  1.000   0.377   0.539
## ---
## Signif. codes:  0 '***' 0.001 '**' 0.01 '*' 0.05 '.' 0.1 ' ' 1
##
## R-sq.(adj) =  0.367
##   Scale est. = 0.023296  n = 2676
```

###### Interaction with Age and Sex

No sex interaction in the interaction between Age and Sleep was found:

```
forms <- age_sex_interaction(psqi_var)
m1s <- gamm(forms[[1]], random = forms[[2]],
            data = subset_data(data, psqi_var), control = list(opt = "optim"))

saveRDS(m1s, file = paste0("./models/", psqi_var, "_m1s.rds"))
m1s <- readRDS(paste0("./models/", psqi_var, "_m1s.rds"))
summary(m1s$gam)
##
## Family: gaussian
## Link function: identity
##
## Formula:
## Hippocampus_st ~ s(Age, k = 6, bs = "cr") + s(Age, by = Sex_ord,
##     k = 3, bs = "cr") + s(PSQI_Comp1_Quality, k = 3, bs = "cr") +
##     s(PSQI_Comp1_Quality, by = Sex_ord, k = 3, bs = "cr") + ti(Age,
##     PSQI_Comp1_Quality, k = c(4, 3)) + ti(Age, PSQI_Comp1_Quality,
##     by = Sex_ord, k = c(4, 3)) + ICV_st + Sex_ord
##
## Parametric coefficients:
##             Estimate Std. Error t value Pr(>|t|)
## (Intercept)  0.02725    0.05935   0.459    0.646
## ICV_st       0.16951    0.01627  10.418   <2e-16 ***
## Sex_ord.L    0.22660    0.02544   8.906   <2e-16 ***
## ---
## Signif. codes:  0 '***' 0.001 '**' 0.01 '*' 0.05 '.' 0.1 ' ' 1
##
## Approximate significance of smooth terms:
##                                          edf Ref.df       F  p-value
## s(Age)                                 4.789  4.789 313.746  < 2e-16 ***
## s(Age):Sex_ordMale                     1.005  1.005  24.716 6.63e-07 ***
## s(PSQI_Comp1_Quality)                  1.014  1.014   1.337    0.250
## s(PSQI_Comp1_Quality):Sex_ordMale      1.014  1.014   0.210    0.642
## ti(Age,PSQI_Comp1_Quality)             1.313  1.313   0.244    0.775
## ti(Age,PSQI_Comp1_Quality):Sex_ordMale 1.166  1.166   0.235    0.598
## ---
## Signif. codes:  0 '***' 0.001 '**' 0.01 '*' 0.05 '.' 0.1 ' ' 1
##
## R-sq.(adj) =  0.418
##   Scale est. = 0.018898  n = 4965
```

##### Longitudinal Analysis

In this section, only participants with two timepoints or more are retained.

Interaction with time.

```
forms <- time_interaction(psqi_var)
m2 <- gamm(forms[[1]], random = forms[[2]],
           data = long_only(subset_data(data, psqi_var)))
saveRDS(m2, file = paste0("./models/", psqi_var, "_m2.rds"))
m2 <- readRDS(paste0("./models/", psqi_var, "_m2.rds"))
summary(m2$gam)
##
## Family: gaussian
## Link function: identity
##
## Formula:
## Hippocampus_st ~ te(dt, Age_bl, k = c(4, 10)) + ti(PSQI_Comp1_Quality,
##     k = 4) + ti(dt, PSQI_Comp1_Quality, k = c(4, 4)) + ICV_st +
##     Sex
##
## Parametric coefficients:
##             Estimate Std. Error t value Pr(>|t|)
## (Intercept) -0.04669    0.05327  -0.877 0.380804
## ICV_st       0.26037    0.02414  10.787  < 2e-16 ***
## SexMale      0.18024    0.04895   3.682 0.000235 ***
## ---
## Signif. codes:  0 '***' 0.001 '**' 0.01 '*' 0.05 '.' 0.1 ' ' 1
##
## Approximate significance of smooth terms:
##                              edf Ref.df       F p-value
## te(dt,Age_bl)             14.991 14.991 136.229 < 2e-16 ***
## ti(PSQI_Comp1_Quality)     1.467  1.467   0.268 0.54799
## ti(dt,PSQI_Comp1_Quality)  4.801  4.801   3.685 0.00185 **
## ---
## Signif. codes:  0 '***' 0.001 '**' 0.01 '*' 0.05 '.' 0.1 ' ' 1
##
## R-sq.(adj) =  0.459
##   Scale est. = 0.018707  n = 3235
```

The term te(Age_bl, dt) can alternatively be written as s(dt) + s(Age_bl) + ti(Age_bl, dt). This would give approximately the same model, but with more smoothing parameters to estimate.

```
fvisgam(m2$gam, view = c("dt", psqi_var))
## Summary:
##  * ICV_st : numeric predictor; set to the value(s): -0.044274334428729.
##  * Sex : factor; set to the value(s): Female.
##  * dt : numeric predictor; with 30 values ranging from 0.000000 to 10.976044.
##  * Age_bl : numeric predictor; set to the value(s): 66.
##  * PSQI_Comp1_Quality : numeric predictor; with 30 values ranging from 0.000000 to 3.000000.
```

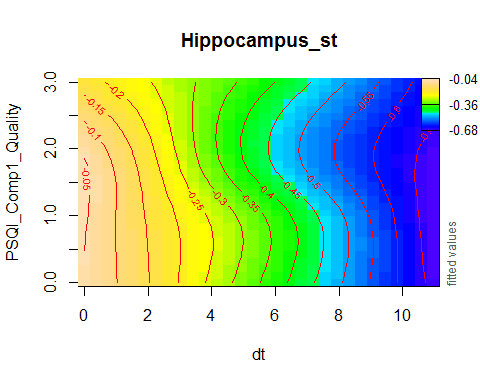

plot_psqi_interaction(m2, psqi_var, psqi_range = 0:3, time_range = 0:5)

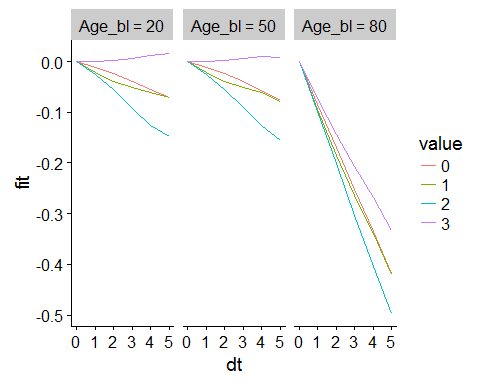

We can also include the offset. Note that those who report the worst sleep problems start out at a lower hippocampus volume.

```
plot_psqi_interaction(m2, psqi_var, psqi_range = 0:3, time_range = 0:5,
                      normalize = FALSE, scales = "free_y")
```

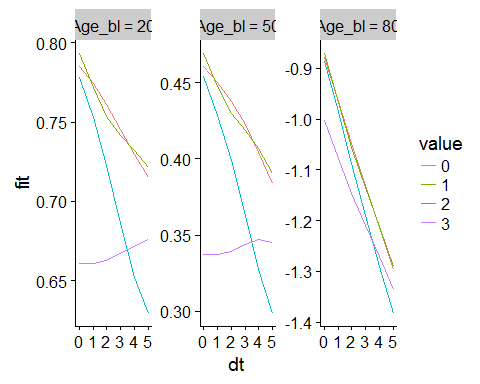

Below are the numbers of participants in each group, where we round to the nearest integer. Number shown are total observations and the number of unique participants.

```
subset_data(data, psqi_var) %>%
  long_only() %>%
  mutate_at(vars(psqi_var), funs(Comp1 = round(., 0))) %>%
  mutate(Age_bl_cat = cut(Age_bl, breaks = seq(from = 10, to = 90, by = 20))) %>%
  group_by_at(vars(Comp1, Age_bl_cat)) %>%
  summarise(Observations = paste(n(), "/", n_distinct(ID))) %>%
  spread(key = Comp1, value = Observations, sep = " = ") %>%
  knitr::kable()
```

| Age_bl_cat | Comp1 = 0 | Comp1 = 1 | Comp1 = 2 | Comp1 = 3 |
| --- | --- | --- | --- | --- |
| (10,30] | 53 / 22 | 224 / 78 | 137 / 47 | 23 / 6 |
| (30,50] | 161 / 72 | 252 / 111 | 83 / 35 | 8 / 4 |
| (50,70] | 299 / 131 | 580 / 240 | 229 / 87 | 12 / 6 |
| (70,90] | 247 / 100 | 580 / 221 | 321 / 91 | 26 / 10 |

If we remove those who report a value of 3, the effect is still significant, and the ordering of 0, 1, and 2 is like before.

```
forms <- time_interaction(psqi_var)
m2b <- gamm(forms[[1]], random = forms[[2]],
           data = filter_at(long_only(subset_data(data, psqi_var)), vars(psqi_var), all_vars(. < 3)))

saveRDS(m2b, file = paste0("./models/", psqi_var, "_m2b.rds"))
m2b <- readRDS(paste0("./models/", psqi_var, "_m2b.rds"))
summary(m2b$gam)
##
## Family: gaussian
## Link function: identity
##
## Formula:
## Hippocampus_st ~ te(dt, Age_bl, k = c(4, 10)) + ti(PSQI_Comp1_Quality,
##     k = 4) + ti(dt, PSQI_Comp1_Quality, k = c(4, 4)) + ICV_st +
##     Sex
##
## Parametric coefficients:
##             Estimate Std. Error t value Pr(>|t|)
## (Intercept) -0.05205    0.05240  -0.993    0.321
## ICV_st       0.25328    0.02436  10.397  < 2e-16 ***
## SexMale      0.19746    0.04949   3.990 6.75e-05 ***
## ---
## Signif. codes:  0 '***' 0.001 '**' 0.01 '*' 0.05 '.' 0.1 ' ' 1
##
## Approximate significance of smooth terms:
##                              edf Ref.df       F p-value
## te(dt,Age_bl)             14.799 14.799 137.051 < 2e-16 ***
## ti(PSQI_Comp1_Quality)     1.000  1.000   0.085 0.77074
## ti(dt,PSQI_Comp1_Quality)  4.025  4.025   3.395 0.00784 **
## ---
## Signif. codes:  0 '***' 0.001 '**' 0.01 '*' 0.05 '.' 0.1 ' ' 1
##
## R-sq.(adj) =  0.451
##   Scale est. = 0.018695  n = 3166
fvisgam(m2b$gam, view = c("dt", psqi_var))
## Summary:
##  * ICV_st : numeric predictor; set to the value(s): -0.0391920031697808.
##  * Sex : factor; set to the value(s): Female.
##  * dt : numeric predictor; with 30 values ranging from 0.000000 to 10.976044.
##  * Age_bl : numeric predictor; set to the value(s): 66.
##  * PSQI_Comp1_Quality : numeric predictor; with 30 values ranging from 0.000000 to 2.500000.
```

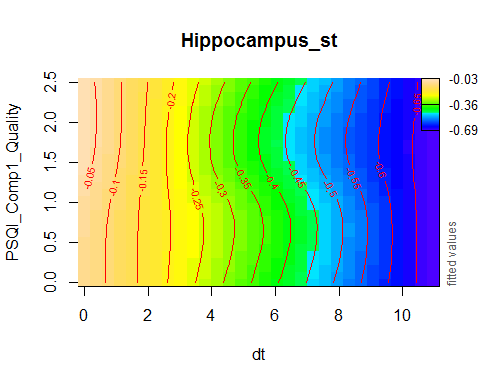

plot_psqi_interaction(m2b, psqi_var, psqi_range = 0:2, time_range = 0:5)

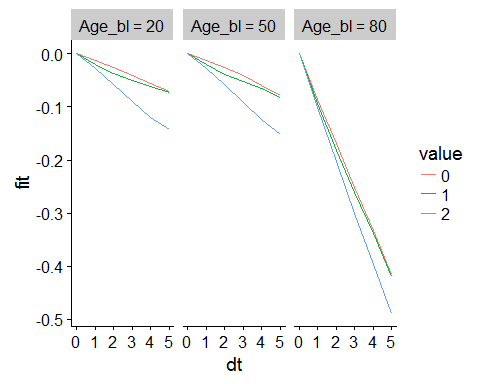

###### Controling for Covariates

```
forms <- time_interaction(psqi_var, c("Depression", "BMI"))
m2_cov <- gamm(forms[[1]], random = forms[[2]],
           data = long_only(subset_data(data, psqi_var)))
saveRDS(m2_cov, file = paste0("./models/", psqi_var, "_m2_cov.rds"))
m2_cov <- readRDS(paste0("./models/", psqi_var, "_m2_cov.rds"))
summary(m2_cov$gam)
##
## Family: gaussian
## Link function: identity
##
## Formula:
## Hippocampus_st ~ te(dt, Age_bl, k = c(4, 10)) + ti(PSQI_Comp1_Quality,
##     k = 4) + ti(dt, PSQI_Comp1_Quality, k = c(4, 4)) + ICV_st +
##     Sex + Depression + BMI
##
## Parametric coefficients:
##              Estimate Std. Error t value Pr(>|t|)
## (Intercept) -0.617568   0.200735  -3.077  0.00213 **
## ICV_st       0.322214   0.036207   8.899  < 2e-16 ***
## SexMale      0.129196   0.066478   1.943  0.05213 .
## Depression  -0.001875   0.029758  -0.063  0.94977
## BMI          0.023448   0.007326   3.200  0.00140 **
## ---
## Signif. codes:  0 '***' 0.001 '**' 0.01 '*' 0.05 '.' 0.1 ' ' 1
##
## Approximate significance of smooth terms:
##                              edf Ref.df      F  p-value
## te(dt,Age_bl)             13.859 13.859 69.779  < 2e-16 ***
## ti(PSQI_Comp1_Quality)     1.000  1.000  0.304 0.581669
## ti(dt,PSQI_Comp1_Quality)  5.333  5.333  4.009 0.000822 ***
## ---
## Signif. codes:  0 '***' 0.001 '**' 0.01 '*' 0.05 '.' 0.1 ' ' 1
##
## R-sq.(adj) =  0.493
##   Scale est. = 0.022617  n = 1695
```

###### Interaction with Sex and Time

Interaction with time and sex. We split the interaction term ti(dt, PSQI_Comp1_Quality) into a term for females and a term for the difference between males and females. We also need to include all lower-level interactions with Sex, to avoid confounding.

As the summary shows, no sex interaction was found. Note that the term te(dt,Age_bl) has a significant interaction with Sex, caused by men and women having different age trajectories for hippocmpus volume. This is not our target for testing in this case.

```
forms <- sex_time_interaction(psqi_var)
m2s <- gamm(forms[[1]], random = forms[[2]],
            data = long_only(subset_data(data, psqi_var)))

saveRDS(m2s, file = paste0("./models/", psqi_var, "_m2s.rds"))
m2s <- readRDS(paste0("./models/", psqi_var, "_m2s.rds"))
summary(m2s$gam)
##
## Family: gaussian
## Link function: identity
##
## Formula:
## Hippocampus_st ~ te(dt, Age_bl, k = c(4, 10)) + te(dt, Age_bl,
##     by = Sex_ord, k = c(4, 10)) + ti(PSQI_Comp1_Quality, k = 4) +
##     ti(PSQI_Comp1_Quality, by = Sex_ord, k = 4) + ti(dt, PSQI_Comp1_Quality,
##     k = c(4, 4)) + ti(dt, PSQI_Comp1_Quality, by = Sex_ord, k = c(4,
##     4)) + ICV_st + Sex_ord
##
## Parametric coefficients:
##             Estimate Std. Error t value Pr(>|t|)
## (Intercept)  0.04186    0.04526   0.925 0.355037
## ICV_st       0.26128    0.02395  10.909  < 2e-16 ***
## Sex_ord.L    0.12304    0.03441   3.575 0.000355 ***
## ---
## Signif. codes:  0 '***' 0.001 '**' 0.01 '*' 0.05 '.' 0.1 ' ' 1
##
## Approximate significance of smooth terms:
##                                          edf Ref.df      F p-value
## te(dt,Age_bl)                         12.046 12.046 80.811 < 2e-16 ***
## te(dt,Age_bl):Sex_ordMale              8.465  8.465  3.164 0.00111 **
## ti(PSQI_Comp1_Quality)                 1.737  1.737  1.494 0.33057
## ti(PSQI_Comp1_Quality):Sex_ordMale     1.000  1.000  3.946 0.04707 *
## ti(dt,PSQI_Comp1_Quality)              4.704  4.704  3.561 0.00244 **
## ti(dt,PSQI_Comp1_Quality):Sex_ordMale  1.000  1.000  0.215 0.64329
## ---
## Signif. codes:  0 '***' 0.001 '**' 0.01 '*' 0.05 '.' 0.1 ' ' 1
##
## R-sq.(adj) =  0.465
##   Scale est. = 0.018598  n = 3235
```

#### PSQI Component 2: Latency

psqi_var <- "PSQI_Comp2_Latency"

psqi_plot(data, psqi_var)

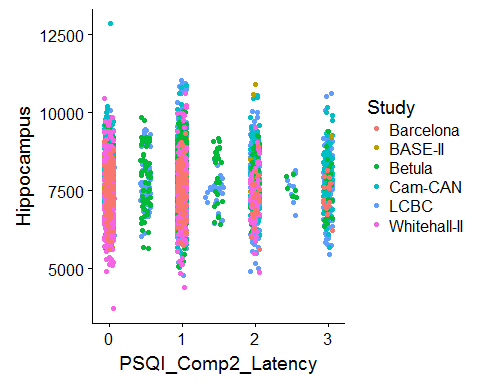

We fit one model per study first, purely for exploratory reasons.

sep_fits(data, psqi_var)

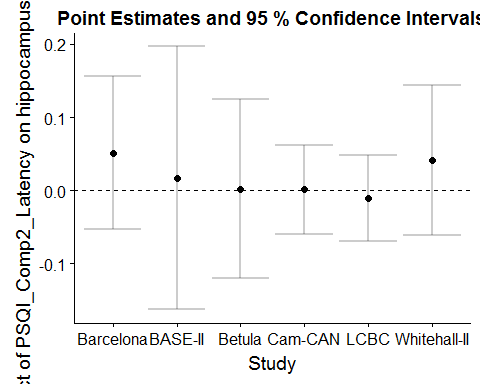

##### Main Effect of Sleep

GAMM with smooth effect of PSQI, random slope of PSQI per study, and random intercept of ID.

```
forms <- main_effects(psqi_var)
m0 <- gamm(forms[[1]], random = forms[[2]],
           data = subset_data(data, psqi_var))
saveRDS(m0, file = paste0("./models/", psqi_var, "_m0.rds"))
m0 <- readRDS(paste0("./models/", psqi_var, "_m0.rds"))
summary(m0$gam)
##
## Family: gaussian
## Link function: identity
##
## Formula:
## Hippocampus_st ~ s(Age, k = 6, bs = "cr") + s(PSQI_Comp2_Latency,
##     k = 3, bs = "cr") + ICV_st + Sex
##
## Parametric coefficients:
##             Estimate Std. Error t value Pr(>|t|)
## (Intercept) -0.19947    0.07324  -2.724  0.00648 **
## ICV_st       0.15474    0.01747   8.858  < 2e-16 ***
## SexMale      0.37797    0.03819   9.896  < 2e-16 ***
## ---
## Signif. codes:  0 '***' 0.001 '**' 0.01 '*' 0.05 '.' 0.1 ' ' 1
##
## Approximate significance of smooth terms:
##                         edf Ref.df       F p-value
## s(Age)                4.760  4.760 474.169  <2e-16 ***
## s(PSQI_Comp2_Latency) 1.201  1.201   0.108   0.841
## ---
## Signif. codes:  0 '***' 0.001 '**' 0.01 '*' 0.05 '.' 0.1 ' ' 1
##
## R-sq.(adj) =  0.391
##   Scale est. = 0.019634  n = 4369
plot(m0$gam, select = 2, seWithMean = TRUE)
```

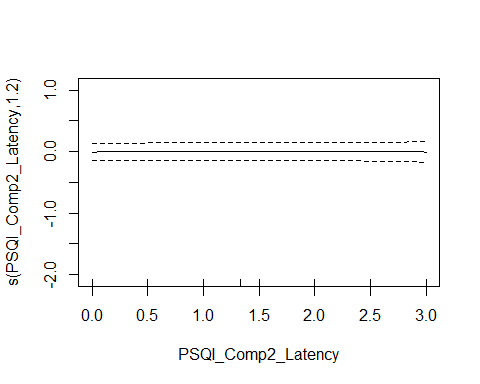

###### Controling for Covariates

```
forms <- main_effects(psqi_var, c("Depression", "BMI"))
m0_cov <- gamm(forms[[1]], random = forms[[2]],
           data = subset_data(data, psqi_var))
saveRDS(m0_cov, file = paste0("./models/", psqi_var, "_m0_cov.rds"))
m0_cov <- readRDS(paste0("./models/", psqi_var, "_m0_cov.rds"))
summary(m0_cov$gam)
##
## Family: gaussian
## Link function: identity
##
## Formula:
## Hippocampus_st ~ s(Age, k = 6, bs = "cr") + s(PSQI_Comp2_Latency,
##     k = 3, bs = "cr") + ICV_st + Sex + Depression + BMI
##
## Parametric coefficients:
##              Estimate Std. Error t value Pr(>|t|)
## (Intercept) -0.449278   0.172660  -2.602  0.00932 **
## ICV_st       0.074951   0.022862   3.278  0.00106 **
## SexMale      0.377683   0.053637   7.041 2.45e-12 ***
## Depression   0.014724   0.021870   0.673  0.50086
## BMI          0.005811   0.005380   1.080  0.28021
## ---
## Signif. codes:  0 '***' 0.001 '**' 0.01 '*' 0.05 '.' 0.1 ' ' 1
##
## Approximate significance of smooth terms:
##                         edf Ref.df       F p-value
## s(Age)                4.508  4.508 237.741  <2e-16 ***
## s(PSQI_Comp2_Latency) 1.000  1.000   0.184   0.668
## ---
## Signif. codes:  0 '***' 0.001 '**' 0.01 '*' 0.05 '.' 0.1 ' ' 1
##
## R-sq.(adj) =  0.348
##   Scale est. = 0.024422  n = 2494
```

###### Interaction with Sex

The summary shows that no sex interaction is found.

```
forms <- main_sex_interaction(psqi_var)
m0s <- gamm(forms[[1]], random = forms[[2]],
           data = subset_data(data, psqi_var), method = "REML")

saveRDS(m0s, file = paste0("./models/", psqi_var, "_m0s.rds"))
m0s <- readRDS(paste0("./models/", psqi_var, "_m0s.rds"))
summary(m0s$gam)
##
## Family: gaussian
## Link function: identity
##
## Formula:
## Hippocampus_st ~ s(Age, k = 6, bs = "cr") + s(PSQI_Comp2_Latency,
##     k = 3, bs = "cr") + s(PSQI_Comp2_Latency, by = Sex_ord, k = 3,
##     bs = "cr") + ICV_st + Sex_ord
##
## Parametric coefficients:
##              Estimate Std. Error t value Pr(>|t|)
## (Intercept) -0.009699   0.076773  -0.126    0.899
## ICV_st       0.154421   0.017483   8.833   <2e-16 ***
## Sex_ord.L    0.268214   0.027046   9.917   <2e-16 ***
## ---
## Signif. codes:  0 '***' 0.001 '**' 0.01 '*' 0.05 '.' 0.1 ' ' 1
##
## Approximate significance of smooth terms:
##                                     edf Ref.df       F p-value
## s(Age)                            4.762  4.762 470.900  <2e-16 ***
## s(PSQI_Comp2_Latency)             1.350  1.350   0.133   0.779
## s(PSQI_Comp2_Latency):Sex_ordMale 1.000  1.000   0.047   0.828
## ---
## Signif. codes:  0 '***' 0.001 '**' 0.01 '*' 0.05 '.' 0.1 ' ' 1
##
## R-sq.(adj) =  0.391
##   Scale est. = 0.019639  n = 4369
```

##### Interaction with Age

```
forms <- age_interaction(psqi_var)
m1 <- gamm(forms[[1]], random = forms[[2]],
           data = subset_data(data, psqi_var), control = list(opt = "optim"))

saveRDS(m1, file = paste0("./models/", psqi_var, "_m1.rds"))
m1 <- readRDS(paste0("./models/", psqi_var, "_m1.rds"))
summary(m1$gam)
##
## Family: gaussian
## Link function: identity
##
## Formula:
## Hippocampus_st ~ s(Age, k = 6, bs = "cr") + s(PSQI_Comp2_Latency,
##     bs = "cr", k = 3) + ti(Age, PSQI_Comp2_Latency, k = c(4,
##     3)) + ICV_st + Sex
##
## Parametric coefficients:
##             Estimate Std. Error t value Pr(>|t|)
## (Intercept) -0.19994    0.07173  -2.787  0.00534 **
## ICV_st       0.15329    0.01745   8.787  < 2e-16 ***
## SexMale      0.38799    0.03828  10.135  < 2e-16 ***
## ---
## Signif. codes:  0 '***' 0.001 '**' 0.01 '*' 0.05 '.' 0.1 ' ' 1
##
## Approximate significance of smooth terms:
##                              edf Ref.df       F p-value
## s(Age)                     4.755  4.755 477.586 < 2e-16 ***
## s(PSQI_Comp2_Latency)      1.031  1.031   0.177 0.69719
## ti(Age,PSQI_Comp2_Latency) 1.078  1.078   8.051 0.00376 **
## ---
## Signif. codes:  0 '***' 0.001 '**' 0.01 '*' 0.05 '.' 0.1 ' ' 1
##
## R-sq.(adj) =  0.394
##   Scale est. = 0.019657  n = 4369
par(mfrow = c(1, 2))
plot(m1$gam, select = 2, seWithMean = TRUE)
fvisgam(m1$gam, view = c("Age", psqi_var))
## Summary:
##  * ICV_st : numeric predictor; set to the value(s): -0.000955268254596884.
##  * Sex : factor; set to the value(s): Female.
##  * Age : numeric predictor; with 30 values ranging from 18.482192 to 91.885010.
##  * PSQI_Comp2_Latency : numeric predictor; with 30 values ranging from 0.000000 to 3.000000.
```

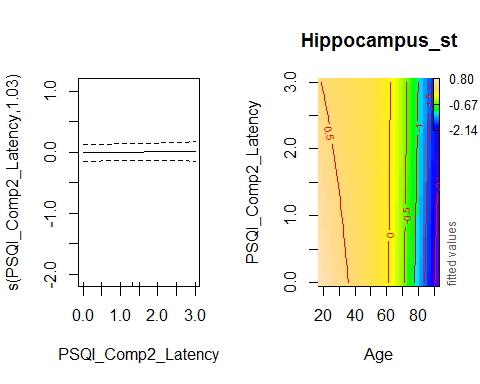

The plot illustrates the interaction.

```
crossing(
  Age = seq(from = 20, to = 80, by = 1),
  PSQI_Comp2_Latency := 0:3,
  ICV_st = 0, Sex = "Female"
) %>%
  bind_cols(fit = predict(m1$gam, newdata = .,
                          terms = c("s(Age)", "s(PSQI_Comp2_Latency)", "ti(Age,PSQI_Comp2_Latency)"))) %>%
  mutate_at(vars(PSQI_Comp2_Latency), factor) %>%
  ggplot(aes(x = Age, y = fit, group = PSQI_Comp2_Latency, color = PSQI_Comp2_Latency)) +
  geom_line()
```

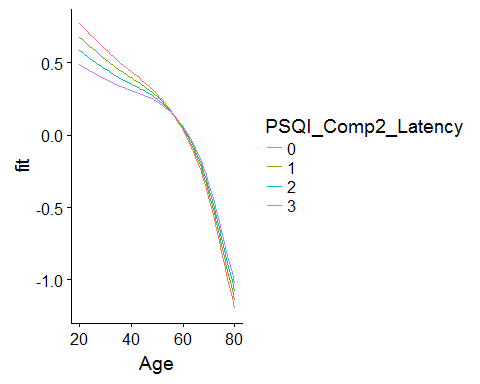

###### Controling for Covariates

```
forms <- age_interaction(psqi_var, c("Depression", "BMI"))
m1_cov <- gamm(forms[[1]], random = forms[[2]],
           data = subset_data(data, psqi_var))

saveRDS(m1_cov, file = paste0("./models/", psqi_var, "_m1_cov.rds"))
m1_cov <- readRDS(paste0("./models/", psqi_var, "_m1_cov.rds"))
summary(m1_cov$gam)
##
## Family: gaussian
## Link function: identity
##
## Formula:
## Hippocampus_st ~ s(Age, k = 6, bs = "cr") + s(PSQI_Comp2_Latency,
##     bs = "cr", k = 3) + ti(Age, PSQI_Comp2_Latency, k = c(4,
##     3)) + ICV_st + Sex + Depression + BMI
##
## Parametric coefficients:
##              Estimate Std. Error t value Pr(>|t|)
## (Intercept) -0.451104   0.171055  -2.637  0.00841 **
## ICV_st       0.074256   0.022825   3.253  0.00116 **
## SexMale      0.387870   0.053707   7.222 6.78e-13 ***
## Depression   0.014841   0.021830   0.680  0.49667
## BMI          0.005807   0.005370   1.081  0.27971
## ---
## Signif. codes:  0 '***' 0.001 '**' 0.01 '*' 0.05 '.' 0.1 ' ' 1
##
## Approximate significance of smooth terms:
##                              edf Ref.df       F p-value
## s(Age)                     4.505  4.505 239.462  <2e-16 ***
## s(PSQI_Comp2_Latency)      1.000  1.000   0.152  0.6963
## ti(Age,PSQI_Comp2_Latency) 1.000  1.000   5.932  0.0149 *
## ---
## Signif. codes:  0 '***' 0.001 '**' 0.01 '*' 0.05 '.' 0.1 ' ' 1
##
## R-sq.(adj) =  0.352
##   Scale est. = 0.024426  n = 2494
```

The interaction between age and latency is no longer significant when controling for Depression and BMI.

```
crossing(
  Age = seq(from = 20, to = 80, by = 1),
  PSQI_Comp2_Latency := 0:3,
  ICV_st = 0, Sex = "Female", BMI = 20, Depression = 0
) %>%
  bind_cols(fit = predict(m1_cov$gam, newdata = .,
                          terms = c("s(Age)", "s(PSQI_Comp2_Latency)", "ti(Age,PSQI_Comp2_Latency)"))) %>%
  mutate_at(vars(PSQI_Comp2_Latency), factor) %>%
  ggplot(aes(x = Age, y = fit, group = PSQI_Comp2_Latency, color = PSQI_Comp2_Latency)) +
  geom_line()
```

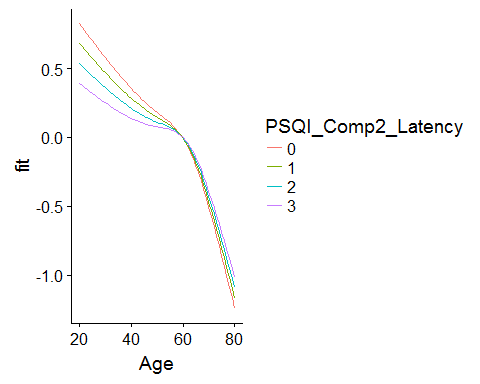

###### Interaction with Age and Sex

No sex interaction in the interaction between Age and Sleep was found:

```
forms <- age_sex_interaction(psqi_var)
m1s <- gamm(forms[[1]], random = forms[[2]],
            data = subset_data(data, psqi_var))

saveRDS(m1s, file = paste0("./models/", psqi_var, "_m1s.rds"))
m1s <- readRDS(paste0("./models/", psqi_var, "_m1s.rds"))
summary(m1s$gam)
##
## Family: gaussian
## Link function: identity
##
## Formula:
## Hippocampus_st ~ s(Age, k = 6, bs = "cr") + s(Age, by = Sex_ord,
##     k = 3, bs = "cr") + s(PSQI_Comp2_Latency, k = 3, bs = "cr") +
##     s(PSQI_Comp2_Latency, by = Sex_ord, k = 3, bs = "cr") + ti(Age,
##     PSQI_Comp2_Latency, k = c(4, 3)) + ti(Age, PSQI_Comp2_Latency,
##     by = Sex_ord, k = c(4, 3)) + ICV_st + Sex_ord
##
## Parametric coefficients:
##              Estimate Std. Error t value Pr(>|t|)
## (Intercept) -0.001367   0.066310  -0.021    0.984
## ICV_st       0.153215   0.017415   8.798   <2e-16 ***
## Sex_ord.L    0.268289   0.027110   9.896   <2e-16 ***
## ---
## Signif. codes:  0 '***' 0.001 '**' 0.01 '*' 0.05 '.' 0.1 ' ' 1
##
## Approximate significance of smooth terms:
##                                          edf Ref.df       F p-value
## s(Age)                                 4.751  4.751 272.033 < 2e-16 ***
## s(Age):Sex_ordMale                     1.000  1.000  10.444 0.00124 **
## s(PSQI_Comp2_Latency)                  1.000  1.000   0.097 0.75548
## s(PSQI_Comp2_Latency):Sex_ordMale      1.000  1.000   0.112 0.73778
## ti(Age,PSQI_Comp2_Latency)             1.000  1.000   5.114 0.02378 *
## ti(Age,PSQI_Comp2_Latency):Sex_ordMale 1.000  1.000   0.175 0.67612
## ---
## Signif. codes:  0 '***' 0.001 '**' 0.01 '*' 0.05 '.' 0.1 ' ' 1
##
## R-sq.(adj) =  0.398
##   Scale est. = 0.019677  n = 4369
```

##### Longitudinal Analysis

In this section, only participants with two timepoints or more are retained.

Interaction with time.

```
forms <- time_interaction(psqi_var)
m2 <- gamm(forms[[1]], random = forms[[2]],
           data = long_only(subset_data(data, psqi_var)))
saveRDS(m2, file = paste0("./models/", psqi_var, "_m2.rds"))
m2 <- readRDS(paste0("./models/", psqi_var, "_m2.rds"))
summary(m2$gam)
##
## Family: gaussian
## Link function: identity
##
## Formula:
## Hippocampus_st ~ te(dt, Age_bl, k = c(4, 10)) + ti(PSQI_Comp2_Latency,
##     k = 4) + ti(dt, PSQI_Comp2_Latency, k = c(4, 4)) + ICV_st +
##     Sex
##
## Parametric coefficients:
##             Estimate Std. Error t value Pr(>|t|)
## (Intercept) -0.07492    0.05809  -1.290    0.197
## ICV_st       0.28066    0.02721  10.315  < 2e-16 ***
## SexMale      0.23284    0.05379   4.329 1.55e-05 ***
## ---
## Signif. codes:  0 '***' 0.001 '**' 0.01 '*' 0.05 '.' 0.1 ' ' 1
##
## Approximate significance of smooth terms:
##                             edf Ref.df       F p-value
## te(dt,Age_bl)             14.77  14.77 110.405  <2e-16 ***
## ti(PSQI_Comp2_Latency)     1.00   1.00   1.191   0.275
## ti(dt,PSQI_Comp2_Latency)  1.00   1.00   0.001   0.979
## ---
## Signif. codes:  0 '***' 0.001 '**' 0.01 '*' 0.05 '.' 0.1 ' ' 1
##
## R-sq.(adj) =  0.457
##   Scale est. = 0.019696  n = 2697
fvisgam(m2$gam, view = c("dt", psqi_var))
## Summary:
##  * ICV_st : numeric predictor; set to the value(s): -0.00264684881092563.
##  * Sex : factor; set to the value(s): Female.
##  * dt : numeric predictor; with 30 values ranging from 0.000000 to 10.976044.
##  * Age_bl : numeric predictor; set to the value(s): 65.
##  * PSQI_Comp2_Latency : numeric predictor; with 30 values ranging from 0.000000 to 3.000000.
```

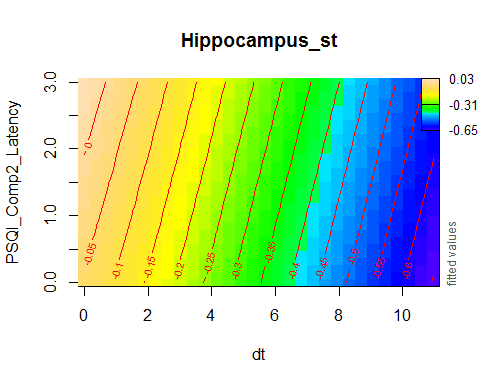

plot_psqi_interaction(m2, psqi_var, psqi_range = 0:3, time_range = 0:5)

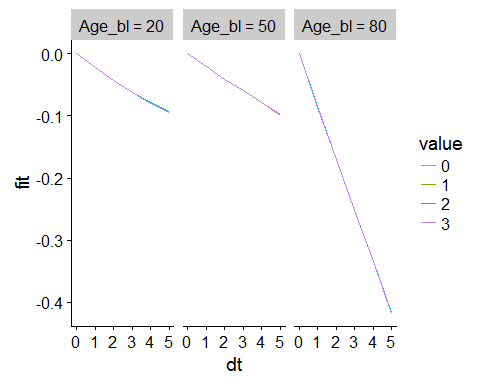

###### Controling for Covariates

```
forms <- time_interaction(psqi_var, c("Depression", "BMI"))
m2_cov <- gamm(forms[[1]], random = forms[[2]],
           data = long_only(subset_data(data, psqi_var)))
saveRDS(m2_cov, file = paste0("./models/", psqi_var, "_m2_cov.rds"))
m2_cov <- readRDS(paste0("./models/", psqi_var, "_m2_cov.rds"))
summary(m2_cov$gam)
##
## Family: gaussian
## Link function: identity
##
## Formula:
## Hippocampus_st ~ te(dt, Age_bl, k = c(4, 10)) + ti(PSQI_Comp2_Latency,
##     k = 4) + ti(dt, PSQI_Comp2_Latency, k = c(4, 4)) + ICV_st +
##     Sex + Depression + BMI
##
## Parametric coefficients:
##              Estimate Std. Error t value Pr(>|t|)
## (Intercept) -0.672249   0.214808  -3.130  0.00178 **
## ICV_st       0.311270   0.037537   8.292  2.4e-16 ***
## SexMale      0.164779   0.070442   2.339  0.01945 *
## Depression  -0.014923   0.030911  -0.483  0.62933
## BMI          0.024135   0.007852   3.074  0.00215 **
## ---
## Signif. codes:  0 '***' 0.001 '**' 0.01 '*' 0.05 '.' 0.1 ' ' 1
##
## Approximate significance of smooth terms:
##                            edf Ref.df      F p-value
## te(dt,Age_bl)             12.3   12.3 71.445  <2e-16 ***
## ti(PSQI_Comp2_Latency)     1.0    1.0  0.427   0.514
## ti(dt,PSQI_Comp2_Latency)  1.0    1.0  0.506   0.477
## ---
## Signif. codes:  0 '***' 0.001 '**' 0.01 '*' 0.05 '.' 0.1 ' ' 1
##
## R-sq.(adj) =  0.476
##   Scale est. = 0.024309  n = 1549
```

###### Interaction with Sex and Time

As the summary shows, no sex interaction was found.

```
forms <- sex_time_interaction(psqi_var)
m2s <- gamm(forms[[1]], random = forms[[2]],
            data = long_only(subset_data(data, psqi_var)), method = "REML")

saveRDS(m2s, file = paste0("./models/", psqi_var, "_m2s.rds"))
m2s <- readRDS(paste0("./models/", psqi_var, "_m2s.rds"))
summary(m2s$gam)
##
## Family: gaussian
## Link function: identity
##
## Formula:
## Hippocampus_st ~ te(dt, Age_bl, k = c(4, 10)) + te(dt, Age_bl,
##     by = Sex_ord, k = c(4, 10)) + ti(PSQI_Comp2_Latency, k = 4) +
##     ti(PSQI_Comp2_Latency, by = Sex_ord, k = 4) + ti(dt, PSQI_Comp2_Latency,
##     k = c(4, 4)) + ti(dt, PSQI_Comp2_Latency, by = Sex_ord, k = c(4,
##     4)) + ICV_st + Sex_ord
##
## Parametric coefficients:
##             Estimate Std. Error t value Pr(>|t|)
## (Intercept)  0.04110    0.05618   0.732    0.464
## ICV_st       0.28067    0.02740  10.245  < 2e-16 ***
## Sex_ord.L    0.16057    0.03827   4.196  2.8e-05 ***
## ---
## Signif. codes:  0 '***' 0.001 '**' 0.01 '*' 0.05 '.' 0.1 ' ' 1
##
## Approximate significance of smooth terms:
##                                          edf Ref.df      F p-value
## te(dt,Age_bl)                         15.265 15.265 55.934  <2e-16 ***
## te(dt,Age_bl):Sex_ordMale              6.246  6.246  1.677   0.120
## ti(PSQI_Comp2_Latency)                 1.000  1.000  1.581   0.209
## ti(PSQI_Comp2_Latency):Sex_ordMale     1.000  1.000  0.902   0.342
## ti(dt,PSQI_Comp2_Latency)              1.000  1.000  0.526   0.468
## ti(dt,PSQI_Comp2_Latency):Sex_ordMale  1.000  1.000  0.662   0.416
## ---
## Signif. codes:  0 '***' 0.001 '**' 0.01 '*' 0.05 '.' 0.1 ' ' 1
##
## R-sq.(adj) =  0.459
##   Scale est. = 0.019631  n = 2697
```

#### PSQI Component 3: Duration

psqi_var <- "PSQI_Comp3_Duration"

psqi_plot(data, psqi_var)

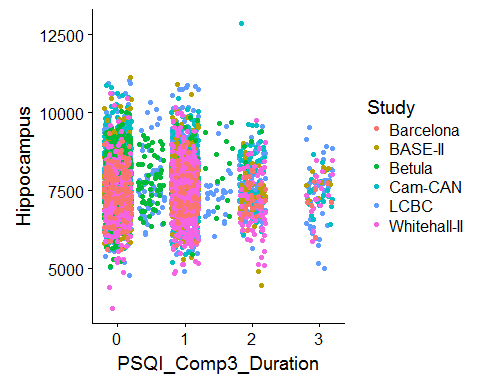

We fit one model per study first, purely for exploratory reasons.

sep_fits(data, psqi_var)

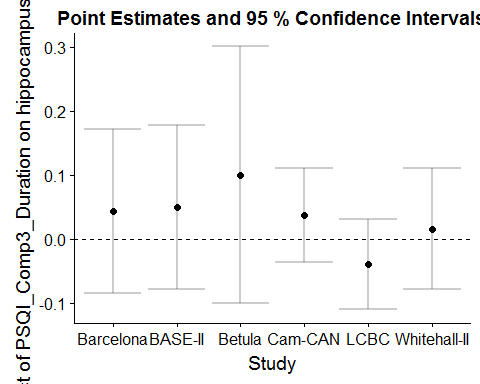

##### Main Effect of Sleep

GAMM with smooth effect of PSQI, random slope of PSQI per study, and random intercept of ID.

```
forms <- main_effects(psqi_var)
m0 <- gamm(forms[[1]], random = forms[[2]],
           data = subset_data(data, psqi_var))
saveRDS(m0, file = paste0("./models/", psqi_var, "_m0.rds"))
m0 <- readRDS(paste0("./models/", psqi_var, "_m0.rds"))
summary(m0$gam)
##
## Family: gaussian
## Link function: identity
##
## Formula:
## Hippocampus_st ~ s(Age, k = 6, bs = "cr") + s(PSQI_Comp3_Duration,
##     k = 3, bs = "cr") + ICV_st + Sex
##
## Parametric coefficients:
##             Estimate Std. Error t value Pr(>|t|)
## (Intercept) -0.17853    0.06768  -2.638  0.00837 **
## ICV_st       0.15537    0.01673   9.284  < 2e-16 ***
## SexMale      0.34143    0.03653   9.346  < 2e-16 ***
## ---
## Signif. codes:  0 '***' 0.001 '**' 0.01 '*' 0.05 '.' 0.1 ' ' 1
##
## Approximate significance of smooth terms:
##                          edf Ref.df       F p-value
## s(Age)                 4.769  4.769 533.159  <2e-16 ***
## s(PSQI_Comp3_Duration) 1.595  1.595   0.821   0.445
## ---
## Signif. codes:  0 '***' 0.001 '**' 0.01 '*' 0.05 '.' 0.1 ' ' 1
##
## R-sq.(adj) =  0.396
##   Scale est. = 0.018961  n = 4762
plot(m0$gam, select = 2, seWithMean = TRUE)
```

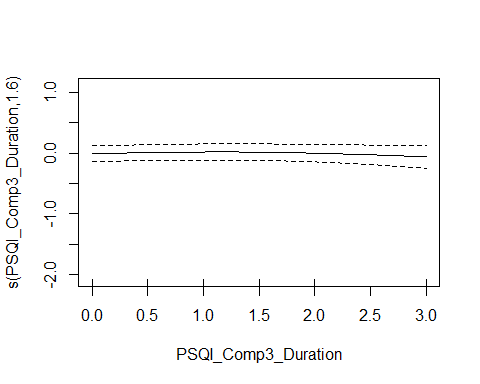

###### Controling for Covariates

```
forms <- main_effects(psqi_var, c("Depression", "BMI"))
m0_cov <- gamm(forms[[1]], random = forms[[2]],
           data = subset_data(data, psqi_var))
saveRDS(m0_cov, file = paste0("./models/", psqi_var, "_m0_cov.rds"))
m0_cov <- readRDS(paste0("./models/", psqi_var, "_m0_cov.rds"))
summary(m0_cov$gam)
##
## Family: gaussian
## Link function: identity
##
## Formula:
## Hippocampus_st ~ s(Age, k = 6, bs = "cr") + s(PSQI_Comp3_Duration,
##     k = 3, bs = "cr") + ICV_st + Sex + Depression + BMI
##
## Parametric coefficients:
##              Estimate Std. Error t value Pr(>|t|)
## (Intercept) -0.433901   0.171023  -2.537  0.01124 *
## ICV_st       0.071663   0.022836   3.138  0.00172 **
## SexMale      0.369203   0.052789   6.994 3.42e-12 ***
## Depression   0.019064   0.022042   0.865  0.38717
## BMI          0.005431   0.005383   1.009  0.31310
## ---
## Signif. codes:  0 '***' 0.001 '**' 0.01 '*' 0.05 '.' 0.1 ' ' 1
##
## Approximate significance of smooth terms:
##                          edf Ref.df   F p-value
## s(Age)                 4.486  4.486 243  <2e-16 ***
## s(PSQI_Comp3_Duration) 1.000  1.000   0   0.994
## ---
## Signif. codes:  0 '***' 0.001 '**' 0.01 '*' 0.05 '.' 0.1 ' ' 1
##
## R-sq.(adj) =  0.352
##   Scale est. = 0.024129  n = 2504
```

###### Interaction with Sex

The summary shows that no sex interaction is found.

```
forms <- main_sex_interaction(psqi_var)
m0s <- gamm(forms[[1]], random = forms[[2]],
           data = subset_data(data, psqi_var), method = "REML")

saveRDS(m0s, file = paste0("./models/", psqi_var, "_m0s.rds"))
m0s <- readRDS(paste0("./models/", psqi_var, "_m0s.rds"))
summary(m0s$gam)
##
## Family: gaussian
## Link function: identity
##
## Formula:
## Hippocampus_st ~ s(Age, k = 6, bs = "cr") + s(PSQI_Comp3_Duration,
##     k = 3, bs = "cr") + s(PSQI_Comp3_Duration, by = Sex_ord,
##     k = 3, bs = "cr") + ICV_st + Sex_ord
##
## Parametric coefficients:
##              Estimate Std. Error t value Pr(>|t|)
## (Intercept) -0.007586   0.070357  -0.108    0.914
## ICV_st       0.155094   0.016735   9.267   <2e-16 ***
## Sex_ord.L    0.240468   0.025849   9.303   <2e-16 ***
## ---
## Signif. codes:  0 '***' 0.001 '**' 0.01 '*' 0.05 '.' 0.1 ' ' 1
##
## Approximate significance of smooth terms:
##                                      edf Ref.df       F p-value
## s(Age)                             4.771  4.771 533.645  <2e-16 ***
## s(PSQI_Comp3_Duration)             1.757  1.757   3.484  0.1086
## s(PSQI_Comp3_Duration):Sex_ordMale 1.000  1.000   4.200  0.0405 *
## ---
## Signif. codes:  0 '***' 0.001 '**' 0.01 '*' 0.05 '.' 0.1 ' ' 1
##
## R-sq.(adj) =  0.397
##   Scale est. = 0.018966  n = 4762
```

##### Interaction with Age

```
forms <- age_interaction(psqi_var)
m1 <- gamm(forms[[1]], random = forms[[2]],
           data = subset_data(data, psqi_var), method = "REML")

saveRDS(m1, file = paste0("./models/", psqi_var, "_m1.rds"))
m1 <- readRDS(paste0("./models/", psqi_var, "_m1.rds"))
summary(m1$gam)
##
## Family: gaussian
## Link function: identity
##
## Formula:
## Hippocampus_st ~ s(Age, k = 6, bs = "cr") + s(PSQI_Comp3_Duration,
##     bs = "cr", k = 3) + ti(Age, PSQI_Comp3_Duration, k = c(4,
##     3)) + ICV_st + Sex
##
## Parametric coefficients:
##             Estimate Std. Error t value Pr(>|t|)
## (Intercept) -0.17860    0.07327  -2.438   0.0148 *
## ICV_st       0.15455    0.01675   9.225   <2e-16 ***
## SexMale      0.34271    0.03661   9.362   <2e-16 ***
## ---
## Signif. codes:  0 '***' 0.001 '**' 0.01 '*' 0.05 '.' 0.1 ' ' 1
##
## Approximate significance of smooth terms:
##                               edf Ref.df       F p-value
## s(Age)                      4.773  4.773 532.823  <2e-16 ***
## s(PSQI_Comp3_Duration)      1.699  1.699   1.023   0.357
## ti(Age,PSQI_Comp3_Duration) 1.734  1.734   1.290   0.292
## ---
## Signif. codes:  0 '***' 0.001 '**' 0.01 '*' 0.05 '.' 0.1 ' ' 1
##
## R-sq.(adj) =  0.396
##   Scale est. = 0.018955  n = 4762
par(mfrow = c(1, 2))
plot(m1$gam, select = 2, seWithMean = TRUE)
fvisgam(m1$gam, view = c("Age", psqi_var))
## Summary:
##  * ICV_st : numeric predictor; set to the value(s): -0.0259201445849824.
##  * Sex : factor; set to the value(s): Male.
##  * Age : numeric predictor; with 30 values ranging from 18.482192 to 91.885010.
##  * PSQI_Comp3_Duration : numeric predictor; with 30 values ranging from 0.000000 to 3.000000.
```

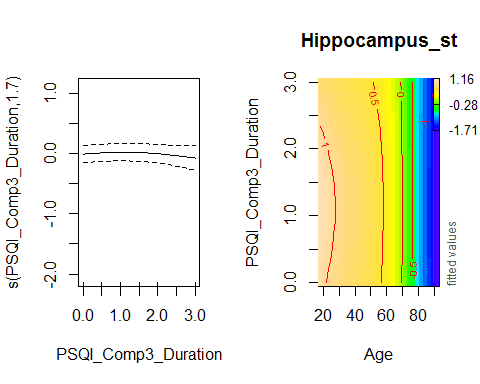

###### Controling for Covariates

```
forms <- age_interaction(psqi_var, c("Depression", "BMI"))
m1_cov <- gamm(forms[[1]], random = forms[[2]],
           data = subset_data(data, psqi_var))

saveRDS(m1_cov, file = paste0("./models/", psqi_var, "_m1_cov.rds"))
m1_cov <- readRDS(paste0("./models/", psqi_var, "_m1_cov.rds"))
summary(m1_cov$gam)
##
## Family: gaussian
## Link function: identity
##
## Formula:
## Hippocampus_st ~ s(Age, k = 6, bs = "cr") + s(PSQI_Comp3_Duration,
##     bs = "cr", k = 3) + ti(Age, PSQI_Comp3_Duration, k = c(4,
##     3)) + ICV_st + Sex + Depression + BMI
##
## Parametric coefficients:
##              Estimate Std. Error t value Pr(>|t|)
## (Intercept) -0.427067   0.171002  -2.497  0.01257 *
## ICV_st       0.071774   0.022835   3.143  0.00169 **
## SexMale      0.367848   0.052803   6.966 4.14e-12 ***
## Depression   0.019895   0.022056   0.902  0.36715
## BMI          0.005291   0.005385   0.983  0.32587
## ---
## Signif. codes:  0 '***' 0.001 '**' 0.01 '*' 0.05 '.' 0.1 ' ' 1
##
## Approximate significance of smooth terms:
##                               edf Ref.df      F p-value
## s(Age)                      4.481  4.481 243.63  <2e-16 ***
## s(PSQI_Comp3_Duration)      1.000  1.000   0.01   0.920
## ti(Age,PSQI_Comp3_Duration) 1.000  1.000   1.05   0.306
## ---
## Signif. codes:  0 '***' 0.001 '**' 0.01 '*' 0.05 '.' 0.1 ' ' 1
##
## R-sq.(adj) =  0.353
##   Scale est. = 0.024105  n = 2504
```

###### Interaction with Age and Sex

No sex interaction in the interaction between Age and Sleep was found:

```
forms <- age_sex_interaction(psqi_var)
m1s <- gamm(forms[[1]], random = forms[[2]],
            data = subset_data(data, psqi_var), control = list(opt = "optim"))

saveRDS(m1s, file = paste0("./models/", psqi_var, "_m1s.rds"))
m1s <- readRDS(paste0("./models/", psqi_var, "_m1s.rds"))
summary(m1s$gam)
##
## Family: gaussian
## Link function: identity
##
## Formula:
## Hippocampus_st ~ s(Age, k = 6, bs = "cr") + s(Age, by = Sex_ord,
##     k = 3, bs = "cr") + s(PSQI_Comp3_Duration, k = 3, bs = "cr") +
##     s(PSQI_Comp3_Duration, by = Sex_ord, k = 3, bs = "cr") +
##     ti(Age, PSQI_Comp3_Duration, k = c(4, 3)) + ti(Age, PSQI_Comp3_Duration,
##     by = Sex_ord, k = c(4, 3)) + ICV_st + Sex_ord
##
## Parametric coefficients:
##               Estimate Std. Error t value Pr(>|t|)
## (Intercept) -0.0003317  0.0599405  -0.006    0.996
## ICV_st       0.1552744  0.0166663   9.317   <2e-16 ***
## Sex_ord.L    0.2358380  0.0258081   9.138   <2e-16 ***
## ---
## Signif. codes:  0 '***' 0.001 '**' 0.01 '*' 0.05 '.' 0.1 ' ' 1
##
## Approximate significance of smooth terms:
##                                           edf Ref.df       F  p-value
## s(Age)                                  4.760  4.760 298.917  < 2e-16 ***
## s(Age):Sex_ordMale                      1.008  1.008  21.626 3.28e-06 ***
## s(PSQI_Comp3_Duration)                  1.693  1.693   2.550   0.1966
## s(PSQI_Comp3_Duration):Sex_ordMale      1.016  1.016   2.750   0.0941 .
## ti(Age,PSQI_Comp3_Duration)             1.089  1.089   0.038   0.8620
## ti(Age,PSQI_Comp3_Duration):Sex_ordMale 1.081  1.081   0.226   0.6533
## ---
## Signif. codes:  0 '***' 0.001 '**' 0.01 '*' 0.05 '.' 0.1 ' ' 1
##
## R-sq.(adj) =  0.404
##   Scale est. = 0.018968  n = 4762
```

##### Longitudinal Analysis

In this section, only participants with two timepoints or more are retained.

Interaction with time.

```
forms <- time_interaction(psqi_var)
m2 <- gamm(forms[[1]], random = forms[[2]],
           data = long_only(subset_data(data, psqi_var)))
saveRDS(m2, file = paste0("./models/", psqi_var, "_m2.rds"))
m2 <- readRDS(paste0("./models/", psqi_var, "_m2.rds"))
summary(m2$gam)
##
## Family: gaussian
## Link function: identity
##
## Formula:
## Hippocampus_st ~ te(dt, Age_bl, k = c(4, 10)) + ti(PSQI_Comp3_Duration,
##     k = 4) + ti(dt, PSQI_Comp3_Duration, k = c(4, 4)) + ICV_st +
##     Sex
##
## Parametric coefficients:
##             Estimate Std. Error t value Pr(>|t|)
## (Intercept) -0.07305    0.05138  -1.422    0.155
## ICV_st       0.25192    0.02439  10.329  < 2e-16 ***
## SexMale      0.19683    0.04962   3.967 7.44e-05 ***
## ---
## Signif. codes:  0 '***' 0.001 '**' 0.01 '*' 0.05 '.' 0.1 ' ' 1
##
## Approximate significance of smooth terms:
##                              edf Ref.df       F p-value
## te(dt,Age_bl)              14.62  14.62 130.923  <2e-16 ***
## ti(PSQI_Comp3_Duration)     1.00   1.00   0.396  0.5294
## ti(dt,PSQI_Comp3_Duration)  1.00   1.00   4.278  0.0387 *
## ---
## Signif. codes:  0 '***' 0.001 '**' 0.01 '*' 0.05 '.' 0.1 ' ' 1
##
## R-sq.(adj) =  0.448
##   Scale est. = 0.018961  n = 3092
fvisgam(m2$gam, view = c("dt", psqi_var))
## Summary:
##  * ICV_st : numeric predictor; set to the value(s): -0.0571877746249291.
##  * Sex : factor; set to the value(s): Female.
##  * dt : numeric predictor; with 30 values ranging from 0.000000 to 10.976044.
##  * Age_bl : numeric predictor; set to the value(s): 66.0260095824778.
##  * PSQI_Comp3_Duration : numeric predictor; with 30 values ranging from 0.000000 to 3.000000.
```

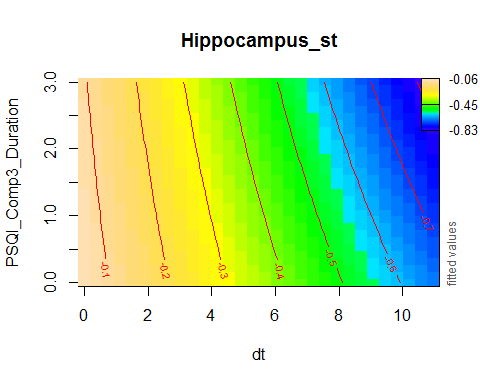

plot_psqi_interaction(m2, psqi_var, psqi_range = 0:3, time_range = 0:5)

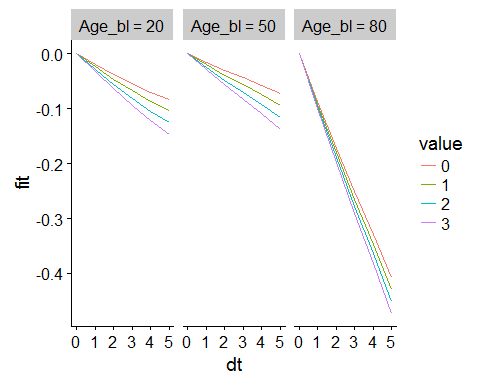

###### Controling for Covariates

```
forms <- time_interaction(psqi_var, c("Depression", "BMI"))
m2_cov <- gamm(forms[[1]], random = forms[[2]],
           data = long_only(subset_data(data, psqi_var)))
saveRDS(m2_cov, file = paste0("./models/", psqi_var, "_m2_cov.rds"))
m2_cov <- readRDS(paste0("./models/", psqi_var, "_m2_cov.rds"))
summary(m2_cov$gam)
##
## Family: gaussian
## Link function: identity
##
## Formula:
## Hippocampus_st ~ te(dt, Age_bl, k = c(4, 10)) + ti(PSQI_Comp3_Duration,
##     k = 4) + ti(dt, PSQI_Comp3_Duration, k = c(4, 4)) + ICV_st +
##     Sex + Depression + BMI
##
## Parametric coefficients:
##              Estimate Std. Error t value Pr(>|t|)
## (Intercept) -0.652319   0.210204  -3.103  0.00195 **
## ICV_st       0.316800   0.037118   8.535  < 2e-16 ***
## SexMale      0.151503   0.068431   2.214  0.02698 *
## Depression  -0.008437   0.030536  -0.276  0.78236
## BMI          0.023387   0.007759   3.014  0.00262 **
## ---
## Signif. codes:  0 '***' 0.001 '**' 0.01 '*' 0.05 '.' 0.1 ' ' 1
##
## Approximate significance of smooth terms:
##                              edf Ref.df      F p-value
## te(dt,Age_bl)              11.78  11.78 76.202  <2e-16 ***
## ti(PSQI_Comp3_Duration)     1.00   1.00  0.365   0.546
## ti(dt,PSQI_Comp3_Duration)  1.00   1.00  1.216   0.270
## ---
## Signif. codes:  0 '***' 0.001 '**' 0.01 '*' 0.05 '.' 0.1 ' ' 1
##
## R-sq.(adj) =  0.482
##   Scale est. = 0.024102  n = 1566
```

###### Interaction with Sex and Time

As the summary shows, no sex interaction was found.

```
forms <- sex_time_interaction(psqi_var)
m2s <- gamm(forms[[1]], random = forms[[2]],
            data = long_only(subset_data(data, psqi_var)), method = "REML")

saveRDS(m2s, file = paste0("./models/", psqi_var, "_m2s.rds"))
m2s <- readRDS(paste0("./models/", psqi_var, "_m2s.rds"))
summary(m2s$gam)
##
## Family: gaussian
## Link function: identity
##
## Formula:
## Hippocampus_st ~ te(dt, Age_bl, k = c(4, 10)) + te(dt, Age_bl,
##     by = Sex_ord, k = c(4, 10)) + ti(PSQI_Comp3_Duration, k = 4) +
##     ti(PSQI_Comp3_Duration, by = Sex_ord, k = 4) + ti(dt, PSQI_Comp3_Duration,
##     k = c(4, 4)) + ti(dt, PSQI_Comp3_Duration, by = Sex_ord,
##     k = c(4, 4)) + ICV_st + Sex_ord
##
## Parametric coefficients:
##             Estimate Std. Error t value Pr(>|t|)
## (Intercept)  0.02319    0.04788   0.484 0.628189
## ICV_st       0.25005    0.02441  10.245  < 2e-16 ***
## Sex_ord.L    0.13301    0.03499   3.802 0.000146 ***
## ---
## Signif. codes:  0 '***' 0.001 '**' 0.01 '*' 0.05 '.' 0.1 ' ' 1
##
## Approximate significance of smooth terms:
##                                           edf Ref.df      F p-value
## te(dt,Age_bl)                          13.600 13.600 67.458 < 2e-16 ***
## te(dt,Age_bl):Sex_ordMale               8.211  8.211  2.797 0.00393 **
## ti(PSQI_Comp3_Duration)                 1.000  1.000  0.260 0.61027
## ti(PSQI_Comp3_Duration):Sex_ordMale     1.990  1.990  3.528 0.02372 *
## ti(dt,PSQI_Comp3_Duration)              3.229  3.229  2.145 0.16402
## ti(dt,PSQI_Comp3_Duration):Sex_ordMale  1.000  1.000  0.035 0.85140
## ---
## Signif. codes:  0 '***' 0.001 '**' 0.01 '*' 0.05 '.' 0.1 ' ' 1
##
## R-sq.(adj) =  0.456
##   Scale est. = 0.018845  n = 3092
```

#### PSQI Component 4: Efficiency

psqi_var <- "PSQI_Comp4_Efficiency"

psqi_plot(data, psqi_var)

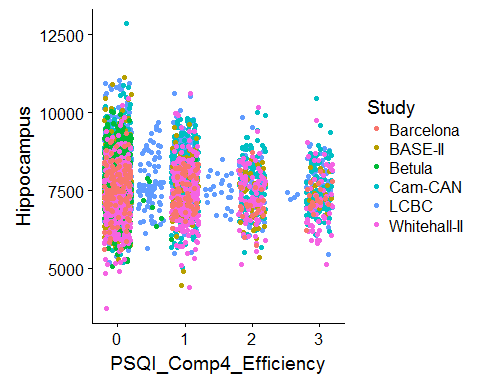

We fit one model per study first, purely for exploratory reasons.

sep_fits(data, psqi_var)

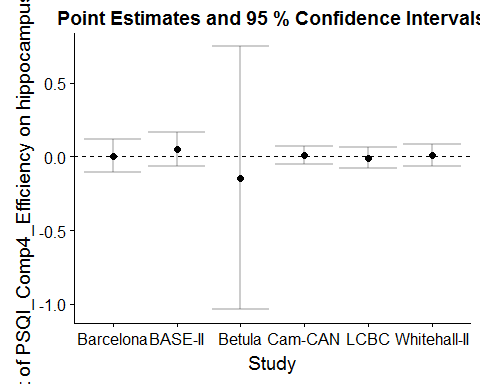

##### Main Effect of Sleep

GAMM with smooth effect of PSQI, random slope of PSQI per study, and random intercept of ID.

```
forms <- main_effects(psqi_var)
m0 <- gamm(forms[[1]], random = forms[[2]],
           data = subset_data(data, psqi_var))
saveRDS(m0, file = paste0("./models/", psqi_var, "_m0.rds"))
m0 <- readRDS(paste0("./models/", psqi_var, "_m0.rds"))
summary(m0$gam)
##
## Family: gaussian
## Link function: identity
##
## Formula:
## Hippocampus_st ~ s(Age, k = 6, bs = "cr") + s(PSQI_Comp4_Efficiency,
##     k = 3, bs = "cr") + ICV_st + Sex
##
## Parametric coefficients:
##             Estimate Std. Error t value Pr(>|t|)
## (Intercept) -0.15020    0.07084  -2.120    0.034 *
## ICV_st       0.15478    0.01749   8.851   <2e-16 ***
## SexMale      0.36059    0.03819   9.443   <2e-16 ***
## ---
## Signif. codes:  0 '***' 0.001 '**' 0.01 '*' 0.05 '.' 0.1 ' ' 1
##
## Approximate significance of smooth terms:
##                            edf Ref.df       F p-value
## s(Age)                   4.775  4.775 447.756  <2e-16 ***
## s(PSQI_Comp4_Efficiency) 1.000  1.000   0.405   0.525
## ---
## Signif. codes:  0 '***' 0.001 '**' 0.01 '*' 0.05 '.' 0.1 ' ' 1
##
## R-sq.(adj) =  0.381
##   Scale est. = 0.017883  n = 4336
plot(m0$gam, select = 2, seWithMean = TRUE)
```

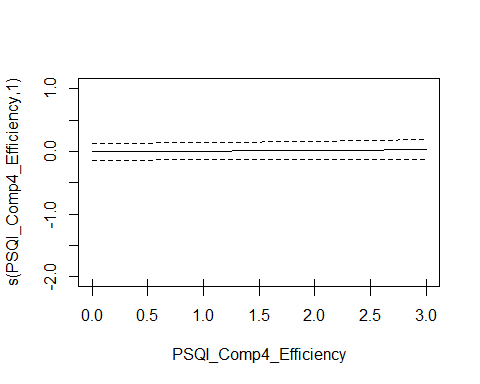

###### Controling for Covariates

```
forms <- main_effects(psqi_var, c("Depression", "BMI"))
m0_cov <- gamm(forms[[1]], random = forms[[2]],
           data = subset_data(data, psqi_var))
saveRDS(m0_cov, file = paste0("./models/", psqi_var, "_m0_cov.rds"))
m0_cov <- readRDS(paste0("./models/", psqi_var, "_m0_cov.rds"))
summary(m0_cov$gam)
##
## Family: gaussian
## Link function: identity
##
## Formula:
## Hippocampus_st ~ s(Age, k = 6, bs = "cr") + s(PSQI_Comp4_Efficiency,
##     k = 3, bs = "cr") + ICV_st + Sex + Depression + BMI
##
## Parametric coefficients:
##              Estimate Std. Error t value Pr(>|t|)
## (Intercept) -0.354705   0.184927  -1.918  0.05523 .
## ICV_st       0.064665   0.024708   2.617  0.00893 **
## SexMale      0.405165   0.057371   7.062 2.21e-12 ***
## Depression   0.011403   0.023723   0.481  0.63081
## BMI          0.004362   0.005904   0.739  0.46003
## ---
## Signif. codes:  0 '***' 0.001 '**' 0.01 '*' 0.05 '.' 0.1 ' ' 1
##
## Approximate significance of smooth terms:
##                            edf Ref.df       F p-value
## s(Age)                   4.432  4.432 189.440  <2e-16 ***
## s(PSQI_Comp4_Efficiency) 1.596  1.596   1.083    0.44
## ---
## Signif. codes:  0 '***' 0.001 '**' 0.01 '*' 0.05 '.' 0.1 ' ' 1
##
## R-sq.(adj) =  0.327
##   Scale est. = 0.022601  n = 2126
```

###### Interaction with Sex

The summary shows that no sex interaction is found.

```
forms <- main_sex_interaction(psqi_var)
m0s <- gamm(forms[[1]], random = forms[[2]],
           data = subset_data(data, psqi_var), method = "REML")

saveRDS(m0s, file = paste0("./models/", psqi_var, "_m0s.rds"))
m0s <- readRDS(paste0("./models/", psqi_var, "_m0s.rds"))
summary(m0s$gam)
##
## Family: gaussian
## Link function: identity
##
## Formula:
## Hippocampus_st ~ s(Age, k = 6, bs = "cr") + s(PSQI_Comp4_Efficiency,
##     k = 3, bs = "cr") + s(PSQI_Comp4_Efficiency, by = Sex_ord,
##     k = 3, bs = "cr") + ICV_st + Sex_ord
##
## Parametric coefficients:
##             Estimate Std. Error t value Pr(>|t|)
## (Intercept)  0.02962    0.07387   0.401    0.688
## ICV_st       0.15465    0.01750   8.838   <2e-16 ***
## Sex_ord.L    0.25613    0.02703   9.476   <2e-16 ***
## ---
## Signif. codes:  0 '***' 0.001 '**' 0.01 '*' 0.05 '.' 0.1 ' ' 1
##
## Approximate significance of smooth terms:
##                                        edf Ref.df       F p-value
## s(Age)                               4.778  4.778 447.247  <2e-16 ***
## s(PSQI_Comp4_Efficiency)             1.000  1.000   1.493   0.222
## s(PSQI_Comp4_Efficiency):Sex_ordMale 1.000  1.000   1.207   0.272
## ---
## Signif. codes:  0 '***' 0.001 '**' 0.01 '*' 0.05 '.' 0.1 ' ' 1
##
## R-sq.(adj) =  0.382
##   Scale est. = 0.017889  n = 4336
```

##### Interaction with Age

```
forms <- age_interaction(psqi_var)
m1 <- gamm(forms[[1]], random = forms[[2]],
           data = subset_data(data, psqi_var), method = "REML")

saveRDS(m1, file = paste0("./models/", psqi_var, "_m1.rds"))
m1 <- readRDS(paste0("./models/", psqi_var, "_m1.rds"))

summary(m1$gam)
##
## Family: gaussian
## Link function: identity
##
## Formula:
## Hippocampus_st ~ s(Age, k = 6, bs = "cr") + s(PSQI_Comp4_Efficiency,
##     bs = "cr", k = 3) + ti(Age, PSQI_Comp4_Efficiency, k = c(4,
##     3)) + ICV_st + Sex
##
## Parametric coefficients:
##             Estimate Std. Error t value Pr(>|t|)
## (Intercept) -0.15233    0.07701  -1.978    0.048 *
## ICV_st       0.15420    0.01750   8.811   <2e-16 ***
## SexMale      0.36261    0.03823   9.485   <2e-16 ***
## ---
## Signif. codes:  0 '***' 0.001 '**' 0.01 '*' 0.05 '.' 0.1 ' ' 1
##
## Approximate significance of smooth terms:
##                                 edf Ref.df       F p-value
## s(Age)                        4.777  4.777 445.914  <2e-16 ***
## s(PSQI_Comp4_Efficiency)      1.000  1.000   0.146   0.702
## ti(Age,PSQI_Comp4_Efficiency) 1.764  1.764   0.505   0.463
## ---
## Signif. codes:  0 '***' 0.001 '**' 0.01 '*' 0.05 '.' 0.1 ' ' 1
##
## R-sq.(adj) =  0.382
##   Scale est. = 0.017896  n = 4336
par(mfrow = c(1, 2))
plot(m1$gam, select = 2, seWithMean = TRUE)
fvisgam(m1$gam, view = c("Age", psqi_var))
## Summary:
##  * ICV_st : numeric predictor; set to the value(s): -0.045115074381007.
##  * Sex : factor; set to the value(s): Male.
##  * Age : numeric predictor; with 30 values ranging from 18.482192 to 91.666667.
##  * PSQI_Comp4_Efficiency : numeric predictor; with 30 values ranging from 0.000000 to 3.000000.
```

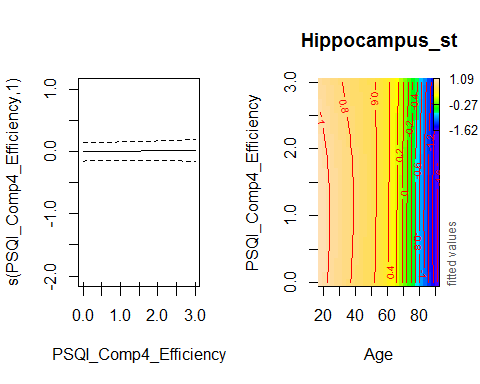

###### Controling for Covariates

```
forms <- age_interaction(psqi_var, c("Depression", "BMI"))
m1_cov <- gamm(forms[[1]], random = forms[[2]],
           data = subset_data(data, psqi_var))

saveRDS(m1_cov, file = paste0("./models/", psqi_var, "_m1_cov.rds"))
m1_cov <- readRDS(paste0("./models/", psqi_var, "_m1_cov.rds"))
summary(m1_cov$gam)
##
## Family: gaussian
## Link function: identity
##
## Formula:
## Hippocampus_st ~ s(Age, k = 6, bs = "cr") + s(PSQI_Comp4_Efficiency,
##     bs = "cr", k = 3) + ti(Age, PSQI_Comp4_Efficiency, k = c(4,
##     3)) + ICV_st + Sex + Depression + BMI
##
## Parametric coefficients:
##              Estimate Std. Error t value Pr(>|t|)
## (Intercept) -0.350622   0.184105  -1.904   0.0570 .
## ICV_st       0.063655   0.024731   2.574   0.0101 *
## SexMale      0.404841   0.057481   7.043 2.53e-12 ***
## Depression   0.012455   0.023788   0.524   0.6006
## BMI          0.004485   0.005912   0.759   0.4481
## ---
## Signif. codes:  0 '***' 0.001 '**' 0.01 '*' 0.05 '.' 0.1 ' ' 1
##
## Approximate significance of smooth terms:
##                                 edf Ref.df       F p-value
## s(Age)                        4.405  4.405 190.676  <2e-16 ***
## s(PSQI_Comp4_Efficiency)      1.673  1.673   1.800  0.2703
## ti(Age,PSQI_Comp4_Efficiency) 1.710  1.710   4.253  0.0848 .
## ---
## Signif. codes:  0 '***' 0.001 '**' 0.01 '*' 0.05 '.' 0.1 ' ' 1
##
## R-sq.(adj) =  0.326
##   Scale est. = 0.0223    n = 2126
```

###### Interaction with Age and Sex

No sex interaction in the interaction between Age and Sleep was found:

```
forms <- age_sex_interaction(psqi_var)
m1s <- gamm(forms[[1]], random = forms[[2]],
            data = subset_data(data, psqi_var), control = list(opt = "optim"))

saveRDS(m1s, file = paste0("./models/", psqi_var, "_m1s.rds"))
m1s <- readRDS(paste0("./models/", psqi_var, "_m1s.rds"))
summary(m1s$gam)
##
## Family: gaussian
## Link function: identity
##
## Formula:
## Hippocampus_st ~ s(Age, k = 6, bs = "cr") + s(Age, by = Sex_ord,
##     k = 3, bs = "cr") + s(PSQI_Comp4_Efficiency, k = 3, bs = "cr") +
##     s(PSQI_Comp4_Efficiency, by = Sex_ord, k = 3, bs = "cr") +
##     ti(Age, PSQI_Comp4_Efficiency, k = c(4, 3)) + ti(Age, PSQI_Comp4_Efficiency,
##     by = Sex_ord, k = c(4, 3)) + ICV_st + Sex_ord
##
## Parametric coefficients:
##             Estimate Std. Error t value Pr(>|t|)
## (Intercept)  0.03782    0.07870   0.481    0.631
## ICV_st       0.15457    0.01746   8.851   <2e-16 ***
## Sex_ord.L    0.25728    0.02732   9.418   <2e-16 ***
## ---
## Signif. codes:  0 '***' 0.001 '**' 0.01 '*' 0.05 '.' 0.1 ' ' 1
##
## Approximate significance of smooth terms:
##                                             edf Ref.df       F  p-value
## s(Age)                                    4.760  4.760 253.914  < 2e-16
## s(Age):Sex_ordMale                        1.016  1.016  16.769 4.05e-05
## s(PSQI_Comp4_Efficiency)                  1.013  1.013   0.393    0.536
## s(PSQI_Comp4_Efficiency):Sex_ordMale      1.014  1.014   0.107    0.742
## ti(Age,PSQI_Comp4_Efficiency)             1.150  1.150   0.827    0.324
## ti(Age,PSQI_Comp4_Efficiency):Sex_ordMale 2.048  2.048   1.900    0.133
##
## s(Age)                                    ***
## s(Age):Sex_ordMale                        ***
## s(PSQI_Comp4_Efficiency)
## s(PSQI_Comp4_Efficiency):Sex_ordMale
## ti(Age,PSQI_Comp4_Efficiency)
## ti(Age,PSQI_Comp4_Efficiency):Sex_ordMale
## ---
## Signif. codes:  0 '***' 0.001 '**' 0.01 '*' 0.05 '.' 0.1 ' ' 1
##
## R-sq.(adj) =  0.388
##   Scale est. = 0.01777   n = 4336
```

##### Longitudinal Analysis

In this section, only participants with two timepoints or more are retained.

Interaction with time.

```
forms <- time_interaction(psqi_var)
m2 <- gamm(forms[[1]], random = forms[[2]],
           data = long_only(subset_data(data, psqi_var)))
saveRDS(m2, file = paste0("./models/", psqi_var, "_m2.rds"))
m2 <- readRDS(paste0("./models/", psqi_var, "_m2.rds"))
summary(m2$gam)
##
## Family: gaussian
## Link function: identity
##
## Formula:
## Hippocampus_st ~ te(dt, Age_bl, k = c(4, 10)) + ti(PSQI_Comp4_Efficiency,
##     k = 4) + ti(dt, PSQI_Comp4_Efficiency, k = c(4, 4)) + ICV_st +
##     Sex
##
## Parametric coefficients:
##             Estimate Std. Error t value Pr(>|t|)
## (Intercept) -0.03035    0.04416  -0.687    0.492
## ICV_st       0.24898    0.02517   9.892  < 2e-16 ***
## SexMale      0.21275    0.05161   4.122 3.86e-05 ***
## ---
## Signif. codes:  0 '***' 0.001 '**' 0.01 '*' 0.05 '.' 0.1 ' ' 1
##
## Approximate significance of smooth terms:
##                                 edf Ref.df       F  p-value
## te(dt,Age_bl)                16.308 16.308 103.343  < 2e-16 ***
## ti(PSQI_Comp4_Efficiency)     1.000  1.000   0.292    0.589
## ti(dt,PSQI_Comp4_Efficiency)  6.899  6.899   6.934 2.36e-08 ***
## ---
## Signif. codes:  0 '***' 0.001 '**' 0.01 '*' 0.05 '.' 0.1 ' ' 1
##
## R-sq.(adj) =  0.426
##   Scale est. = 0.017305  n = 2793
fvisgam(m2$gam, view = c("dt", psqi_var))
## Summary:
##  * ICV_st : numeric predictor; set to the value(s): -0.0720019148228849.
##  * Sex : factor; set to the value(s): Female.
##  * dt : numeric predictor; with 30 values ranging from 0.000000 to 10.976044.
##  * Age_bl : numeric predictor; set to the value(s): 64.7643835616438.
##  * PSQI_Comp4_Efficiency : numeric predictor; with 30 values ranging from 0.000000 to 3.000000.
```

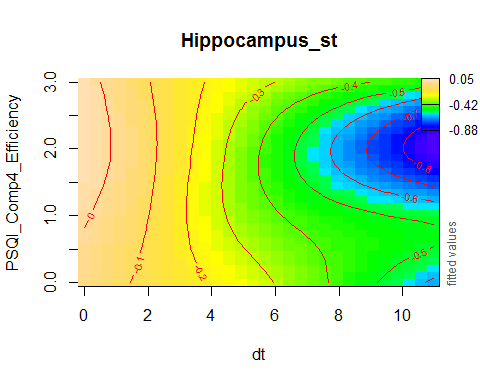

plot_psqi_interaction(m2, psqi_var, psqi_range = 0:3, time_range = 0:5)

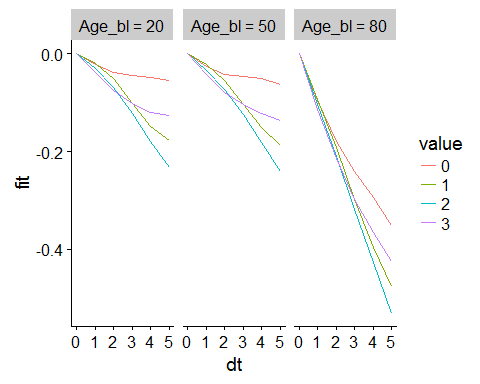

###### Controling for Covariates

```
forms <- time_interaction(psqi_var, c("Depression", "BMI"))
m2_cov <- gamm(forms[[1]], random = forms[[2]],
           data = long_only(subset_data(data, psqi_var)), optim = list(opt = "optim"))
saveRDS(m2_cov, file = paste0("./models/", psqi_var, "_m2_cov.rds"))
m2_cov <- readRDS(paste0("./models/", psqi_var, "_m2_cov.rds"))
summary(m2_cov$gam)
##
## Family: gaussian
## Link function: identity
##
## Formula:
## Hippocampus_st ~ te(dt, Age_bl, k = c(4, 10)) + ti(PSQI_Comp4_Efficiency,
##     k = 4) + ti(dt, PSQI_Comp4_Efficiency, k = c(4, 4)) + ICV_st +
##     Sex + Depression + BMI
##
## Parametric coefficients:
##              Estimate Std. Error t value Pr(>|t|)
## (Intercept) -0.415594   0.218144  -1.905   0.0570 .
## ICV_st       0.333173   0.041540   8.021 2.36e-15 ***
## SexMale      0.170875   0.075405   2.266   0.0236 *
## Depression  -0.007943   0.033458  -0.237   0.8124
## BMI          0.018741   0.008370   2.239   0.0253 *
## ---
## Signif. codes:  0 '***' 0.001 '**' 0.01 '*' 0.05 '.' 0.1 ' ' 1
##
## Approximate significance of smooth terms:
##                                 edf Ref.df      F  p-value
## te(dt,Age_bl)                15.895 15.895 37.190  < 2e-16 ***
## ti(PSQI_Comp4_Efficiency)     1.000  1.000  0.102     0.75
## ti(dt,PSQI_Comp4_Efficiency)  7.482  7.482  9.260 1.05e-10 ***
## ---
## Signif. codes:  0 '***' 0.001 '**' 0.01 '*' 0.05 '.' 0.1 ' ' 1
##
## R-sq.(adj) =  0.438
##   Scale est. = 0.020438  n = 1305
```

###### Interaction with Sex and Time

```
forms <- sex_time_interaction(psqi_var)
m2s <- gamm(forms[[1]], random = forms[[2]],
            data = long_only(subset_data(data, psqi_var)), method = "REML")

saveRDS(m2s, file = paste0("./models/", psqi_var, "_m2s.rds"))
m2s <- readRDS(paste0("./models/", psqi_var, "_m2s.rds"))
summary(m2s$gam)
##
## Family: gaussian
## Link function: identity
##
## Formula:
## Hippocampus_st ~ te(dt, Age_bl, k = c(4, 10)) + te(dt, Age_bl,
##     by = Sex_ord, k = c(4, 10)) + ti(PSQI_Comp4_Efficiency, k = 4) +
##     ti(PSQI_Comp4_Efficiency, by = Sex_ord, k = 4) + ti(dt, PSQI_Comp4_Efficiency,
##     k = c(4, 4)) + ti(dt, PSQI_Comp4_Efficiency, by = Sex_ord,
##     k = c(4, 4)) + ICV_st + Sex_ord
##
## Parametric coefficients:
##             Estimate Std. Error t value Pr(>|t|)
## (Intercept)  0.07681    0.03740   2.054 0.040096 *
## ICV_st       0.25340    0.02520  10.057  < 2e-16 ***
## Sex_ord.L    0.14017    0.03648   3.842 0.000125 ***
## ---
## Signif. codes:  0 '***' 0.001 '**' 0.01 '*' 0.05 '.' 0.1 ' ' 1
##
## Approximate significance of smooth terms:
##                                             edf Ref.df      F  p-value
## te(dt,Age_bl)                            16.599 16.599 49.231  < 2e-16 ***
## te(dt,Age_bl):Sex_ordMale                 5.723  5.723  3.636  0.00192 **
## ti(PSQI_Comp4_Efficiency)                 1.000  1.000  1.563  0.21135
## ti(PSQI_Comp4_Efficiency):Sex_ordMale     1.000  1.000  2.725  0.09890 .
## ti(dt,PSQI_Comp4_Efficiency)              4.331  4.331  3.132  0.00942 **
## ti(dt,PSQI_Comp4_Efficiency):Sex_ordMale  6.350  6.350  6.551 6.78e-06 ***
## ---
## Signif. codes:  0 '***' 0.001 '**' 0.01 '*' 0.05 '.' 0.1 ' ' 1
##
## R-sq.(adj) =  0.434
##   Scale est. = 0.017028  n = 2793
par(mfrow = c(1, 2))
plot(m2s$gam, select = 5, scheme = 2)
plot(m2s$gam, select = 6, scheme = 2)
```

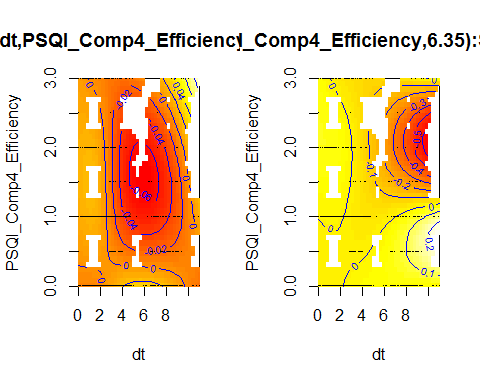

```
crossing(dt = 0:10,
         Age_bl = c(20, 50, 80),
         ICV_st = 0,
         Sex_ord = c("Female", "Male"),
         Study = "Betula",
         ID = "A",
         value = 0:3
         ) %>%
  mutate(!! psqi_var := value) %>%
  bind_cols(fit = predict(m2s$gam, newdata = .)) %>%
  mutate_at(vars(value, Age_bl, Sex_ord), factor) %>%
  group_by(value, Age_bl, Sex_ord) %>%
  arrange(dt, .by_group = TRUE) %>%
  mutate(fit = fit - first(fit)) %>%
  ungroup() %>%
  ggplot(aes(x = dt, y = fit, group = value, color = value)) +
  geom_line() +
  facet_wrap(~ Sex_ord + Age_bl)
```

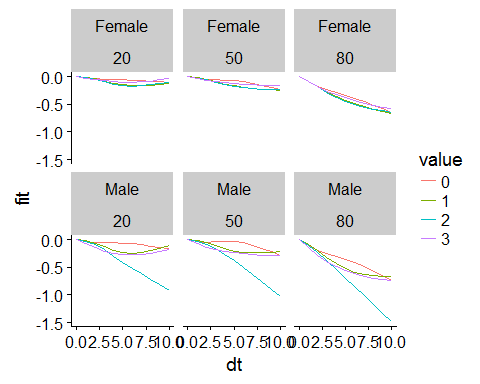

*An outlier analysis follows. This might have to be removed later, due to privacy concerns about showing a single individual’s data.*

The difference starts after about 4 years. How many males have answered 2 on the question and have a follow-up of more than 4 years?

```
long_only(subset_data(data, psqi_var)) %>%
  group_by(ID) %>%
  filter(max(dt, na.rm = TRUE) > 4) %>%
  mutate_at(vars(psqi_var), as.integer) %>%
  group_by_at(vars(Sex, !!psqi_var)) %>%
  summarise(Participants = n_distinct(ID))
## # A tibble: 8 x 3
## # Groups:   Sex [?]
##   Sex    PSQI_Comp4_Efficiency Participants
##   <chr>                  <int>        <int>
## 1 Female                     0           87
## 2 Female                     1           30
## 3 Female                     2            5
## 4 Female                     3            6
## 5 Male                       0           63
## 6 Male                       1           18
## 7 Male                       2            1
## 8 Male                       3            3
```

There are only two participants in this group!! Looking further into one of them, we see an extreme reduction in hippocampus volume from age 68 to age 74. We hence try again removing this single participant.

```
long_only(subset_data(data, psqi_var)) %>%
  group_by(ID) %>%
  filter(max(dt, na.rm = TRUE) > 4) %>%
  mutate_at(vars(psqi_var), as.integer) %>%
  filter(Sex == "Male") %>%
  filter_at(vars(!!psqi_var), all_vars(. == 2L))
## # A tibble: 4 x 17
## # Groups:   ID [1]
##     Age Age_bl    dt    ICV Hippocampus Sex   ID    Study Site_Name ICV_st
##   <dbl>  <dbl> <dbl>  <dbl>       <dbl> <chr> <fct> <fct> <chr>      <dbl>
## 1  65.3   65.3  0.   1.71e6       7457. Male  LCBC~ LCBC  ousAvanto  0.872
## 2  68.3   65.3  3.03 1.71e6       7421. Male  LCBC~ LCBC  ousAvanto  0.872
## 3  72.4   65.3  7.15 1.71e6       6666. Male  LCBC~ LCBC  ousAvanto  0.872
## 4  74.3   65.3  9.03 1.66e6       6033. Male  LCBC~ LCBC  ousSkyra   0.586
## # ... with 7 more variables: Hippocampus_st <dbl>, key <chr>,
## #   PSQI_Comp4_Efficiency <int>, Depression <dbl>, BMI <dbl>,
## #   Education <dbl>, Sex_ord <ord>
```

We fit the model again, removing the single outlying participant. Now there is no significant Sex interaction, with a p-value of 0.09.

```
forms <- sex_time_interaction(psqi_var)
m2sb <- gamm(forms[[1]], random = forms[[2]],
            data = filter(long_only(subset_data(data, psqi_var)), ID != "LCBC1100558"),
            method = "REML", control = list(opt = "optim"))
saveRDS(m2sb, file = paste0("./models/", psqi_var, "_m2sb.rds"))
m2sb <- readRDS(paste0("./models/", psqi_var, "_m2sb.rds"))
summary(m2sb$gam)
##
## Family: gaussian
## Link function: identity
##
## Formula:
## Hippocampus_st ~ te(dt, Age_bl, k = c(4, 10)) + te(dt, Age_bl,
##     by = Sex_ord, k = c(4, 10)) + ti(PSQI_Comp4_Efficiency, k = 4) +
##     ti(PSQI_Comp4_Efficiency, by = Sex_ord, k = 4) + ti(dt, PSQI_Comp4_Efficiency,
##     k = c(4, 4)) + ti(dt, PSQI_Comp4_Efficiency, by = Sex_ord,
##     k = c(4, 4)) + ICV_st + Sex_ord
##
## Parametric coefficients:
##             Estimate Std. Error t value Pr(>|t|)
## (Intercept)  0.07939    0.03749   2.118   0.0343 *
## ICV_st       0.25281    0.02524  10.016  < 2e-16 ***
## Sex_ord.L    0.14437    0.03651   3.954 7.86e-05 ***
## ---
## Signif. codes:  0 '***' 0.001 '**' 0.01 '*' 0.05 '.' 0.1 ' ' 1
##
## Approximate significance of smooth terms:
##                                             edf Ref.df      F p-value
## te(dt,Age_bl)                            17.951 17.951 47.226 < 2e-16 ***
## te(dt,Age_bl):Sex_ordMale                 3.360  3.360  4.087 0.00524 **
## ti(PSQI_Comp4_Efficiency)                 1.011  1.011  1.547 0.21258
## ti(PSQI_Comp4_Efficiency):Sex_ordMale     1.014  1.014  2.198 0.13643
## ti(dt,PSQI_Comp4_Efficiency)              4.977  4.977  4.677 0.00031 ***
## ti(dt,PSQI_Comp4_Efficiency):Sex_ordMale  1.912  1.912  3.525 0.06430 .
## ---
## Signif. codes:  0 '***' 0.001 '**' 0.01 '*' 0.05 '.' 0.1 ' ' 1
##
## R-sq.(adj) =  0.435
##   Scale est. = 0.016992  n = 2789
```

#### PSQI Component 5: Problems

psqi_var <- "PSQI_Comp5_Problems"

psqi_plot(data, psqi_var)

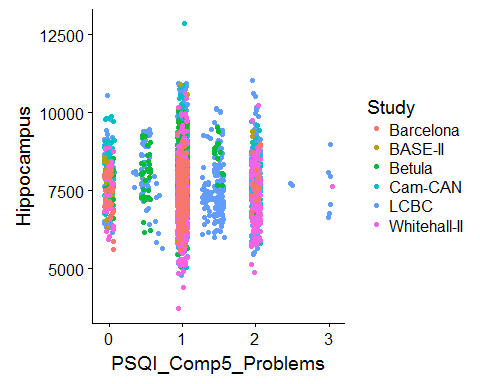

We fit one model per study first, purely for exploratory reasons.

sep_fits(data, psqi_var)

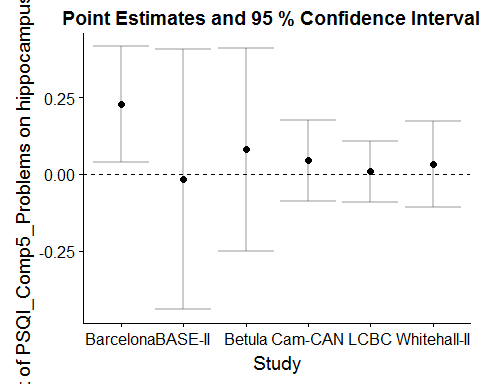

##### Main Effect of Sleep

GAMM with smooth effect of PSQI, random slope of PSQI per study, and random intercept of ID.

```
forms <- main_effects(psqi_var)
m0 <- gamm(forms[[1]], random = forms[[2]],
           data = subset_data(data, psqi_var))
saveRDS(m0, file = paste0("./models/", psqi_var, "_m0.rds"))
m0 <- readRDS(paste0("./models/", psqi_var, "_m0.rds"))
summary(m0$gam)
##
## Family: gaussian
## Link function: identity
##
## Formula:
## Hippocampus_st ~ s(Age, k = 6, bs = "cr") + s(PSQI_Comp5_Problems,
##     k = 3, bs = "cr") + ICV_st + Sex
##
## Parametric coefficients:
##             Estimate Std. Error t value Pr(>|t|)
## (Intercept) -0.16277    0.07208  -2.258    0.024 *
## ICV_st       0.16836    0.01677  10.040   <2e-16 ***
## SexMale      0.35872    0.03649   9.830   <2e-16 ***
## ---
## Signif. codes:  0 '***' 0.001 '**' 0.01 '*' 0.05 '.' 0.1 ' ' 1
##
## Approximate significance of smooth terms:
##                          edf Ref.df       F p-value
## s(Age)                 4.791  4.791 514.656  <2e-16 ***
## s(PSQI_Comp5_Problems) 1.000  1.000   0.927   0.336
## ---
## Signif. codes:  0 '***' 0.001 '**' 0.01 '*' 0.05 '.' 0.1 ' ' 1
##
## R-sq.(adj) =  0.409
##   Scale est. = 0.019224  n = 4690
plot(m0$gam, select = 2, seWithMean = TRUE)
```

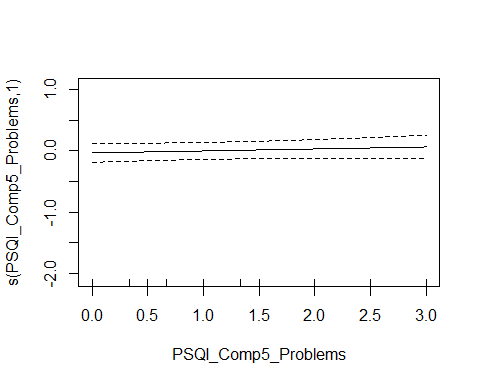

###### Controling for Covariates

```
forms <- main_effects(psqi_var, c("Depression", "BMI"))
m0_cov <- gamm(forms[[1]], random = forms[[2]],
           data = subset_data(data, psqi_var))
saveRDS(m0_cov, file = paste0("./models/", psqi_var, "_m0_cov.rds"))
m0_cov <- readRDS(paste0("./models/", psqi_var, "_m0_cov.rds"))
summary(m0_cov$gam)
##
## Family: gaussian
## Link function: identity
##
## Formula:
## Hippocampus_st ~ s(Age, k = 6, bs = "cr") + s(PSQI_Comp5_Problems,
##     k = 3, bs = "cr") + ICV_st + Sex + Depression + BMI
##
## Parametric coefficients:
##              Estimate Std. Error t value Pr(>|t|)
## (Intercept) -0.398448   0.165489  -2.408 0.016121 *
## ICV_st       0.082855   0.022292   3.717 0.000206 ***
## SexMale      0.352399   0.051305   6.869 8.03e-12 ***
## Depression   0.011431   0.021468   0.532 0.594435
## BMI          0.005690   0.005205   1.093 0.274404
## ---
## Signif. codes:  0 '***' 0.001 '**' 0.01 '*' 0.05 '.' 0.1 ' ' 1
##
## Approximate significance of smooth terms:
##                          edf Ref.df       F p-value
## s(Age)                 4.503  4.503 264.463  <2e-16 ***
## s(PSQI_Comp5_Problems) 1.000  1.000   0.008   0.928
## ---
## Signif. codes:  0 '***' 0.001 '**' 0.01 '*' 0.05 '.' 0.1 ' ' 1
##
## R-sq.(adj) =  0.367
##   Scale est. = 0.023372  n = 2670
```

###### Interaction with Sex

The summary shows that no sex interaction is found.

```
forms <- main_sex_interaction(psqi_var)
m0s <- gamm(forms[[1]], random = forms[[2]],
           data = subset_data(data, psqi_var), method = "REML")

saveRDS(m0s, file = paste0("./models/", psqi_var, "_m0s.rds"))
m0s <- readRDS(paste0("./models/", psqi_var, "_m0s.rds"))
summary(m0s$gam)
##
## Family: gaussian
## Link function: identity
##
## Formula:
## Hippocampus_st ~ s(Age, k = 6, bs = "cr") + s(PSQI_Comp5_Problems,
##     k = 3, bs = "cr") + s(PSQI_Comp5_Problems, by = Sex_ord,
##     k = 3, bs = "cr") + ICV_st + Sex_ord
##
## Parametric coefficients:
##             Estimate Std. Error t value Pr(>|t|)
## (Intercept)  0.01712    0.07643   0.224    0.823
## ICV_st       0.16774    0.01678   9.996   <2e-16 ***
## Sex_ord.L    0.25576    0.02584   9.897   <2e-16 ***
## ---
## Signif. codes:  0 '***' 0.001 '**' 0.01 '*' 0.05 '.' 0.1 ' ' 1
##
## Approximate significance of smooth terms:
##                                      edf Ref.df       F p-value
## s(Age)                             4.794  4.794 514.372  <2e-16 ***
## s(PSQI_Comp5_Problems)             1.000  1.000   0.055   0.814
## s(PSQI_Comp5_Problems):Sex_ordMale 1.000  1.000   1.852   0.174
## ---
## Signif. codes:  0 '***' 0.001 '**' 0.01 '*' 0.05 '.' 0.1 ' ' 1
##
## R-sq.(adj) =  0.409
##   Scale est. = 0.019229  n = 4690
```

##### Interaction with Age

```
forms <- age_interaction(psqi_var)
m1 <- gamm(forms[[1]], random = forms[[2]],
           data = subset_data(data, psqi_var), method = "REML")

saveRDS(m1, file = paste0("./models/", psqi_var, "_m1.rds"))
m1 <- readRDS(paste0("./models/", psqi_var, "_m1.rds"))
summary(m1$gam)
##
## Family: gaussian
## Link function: identity
##
## Formula:
## Hippocampus_st ~ s(Age, k = 6, bs = "cr") + s(PSQI_Comp5_Problems,
##     bs = "cr", k = 3) + ti(Age, PSQI_Comp5_Problems, k = c(4,
##     3)) + ICV_st + Sex
##
## Parametric coefficients:
##             Estimate Std. Error t value Pr(>|t|)
## (Intercept) -0.16327    0.07820  -2.088   0.0369 *
## ICV_st       0.16787    0.01678  10.002   <2e-16 ***
## SexMale      0.36047    0.03654   9.866   <2e-16 ***
## ---
## Signif. codes:  0 '***' 0.001 '**' 0.01 '*' 0.05 '.' 0.1 ' ' 1
##
## Approximate significance of smooth terms:
##                               edf Ref.df       F p-value
## s(Age)                      4.792  4.792 512.387  <2e-16 ***
## s(PSQI_Comp5_Problems)      1.000  1.000   0.851   0.356
## ti(Age,PSQI_Comp5_Problems) 1.587  1.587   0.963   0.496
## ---
## Signif. codes:  0 '***' 0.001 '**' 0.01 '*' 0.05 '.' 0.1 ' ' 1
##
## R-sq.(adj) =  0.409
##   Scale est. = 0.019208  n = 4690
par(mfrow = c(1, 2))
plot(m1$gam, select = 2, seWithMean = TRUE)
fvisgam(m1$gam, view = c("Age", psqi_var))
## Summary:
##  * ICV_st : numeric predictor; set to the value(s): -0.000955268254596884.
##  * Sex : factor; set to the value(s): Female.
##  * Age : numeric predictor; with 30 values ranging from 18.482192 to 91.885010.
##  * PSQI_Comp5_Problems : numeric predictor; with 30 values ranging from 0.000000 to 3.000000.
```

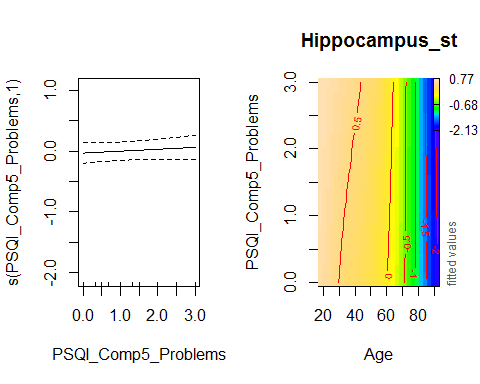

###### Controling for Covariates

```
forms <- age_interaction(psqi_var, c("Depression", "BMI"))
m1_cov <- gamm(forms[[1]], random = forms[[2]],
           data = subset_data(data, psqi_var))

saveRDS(m1_cov, file = paste0("./models/", psqi_var, "_m1_cov.rds"))
m1_cov <- readRDS(paste0("./models/", psqi_var, "_m1_cov.rds"))
summary(m1_cov$gam)
##
## Family: gaussian
## Link function: identity
##
## Formula:
## Hippocampus_st ~ s(Age, k = 6, bs = "cr") + s(PSQI_Comp5_Problems,
##     bs = "cr", k = 3) + ti(Age, PSQI_Comp5_Problems, k = c(4,
##     3)) + ICV_st + Sex + Depression + BMI
##
## Parametric coefficients:
##              Estimate Std. Error t value Pr(>|t|)
## (Intercept) -0.396452   0.165625  -2.394  0.01675 *
## ICV_st       0.082283   0.022309   3.688  0.00023 ***
## SexMale      0.354632   0.051379   6.902 6.37e-12 ***
## Depression   0.011614   0.021482   0.541  0.58881
## BMI          0.005651   0.005208   1.085  0.27801
## ---
## Signif. codes:  0 '***' 0.001 '**' 0.01 '*' 0.05 '.' 0.1 ' ' 1
##
## Approximate significance of smooth terms:
##                               edf Ref.df       F p-value
## s(Age)                      4.493  4.493 265.152  <2e-16 ***
## s(PSQI_Comp5_Problems)      1.000  1.000   0.095   0.758
## ti(Age,PSQI_Comp5_Problems) 1.000  1.000   1.083   0.298
## ---
## Signif. codes:  0 '***' 0.001 '**' 0.01 '*' 0.05 '.' 0.1 ' ' 1
##
## R-sq.(adj) =  0.366
##   Scale est. = 0.023299  n = 2670
```

###### Interaction with Age and Sex

No sex interaction in the interaction between Age and Sleep was found:

```
forms <- age_sex_interaction(psqi_var)
m1s <- gamm(forms[[1]], random = forms[[2]],
            data = subset_data(data, psqi_var))

saveRDS(m1s, file = paste0("./models/", psqi_var, "_m1s.rds"))
m1s <- readRDS(paste0("./models/", psqi_var, "_m1s.rds"))
summary(m1s$gam)
##
## Family: gaussian
## Link function: identity
##
## Formula:
## Hippocampus_st ~ s(Age, k = 6, bs = "cr") + s(Age, by = Sex_ord,
##     k = 3, bs = "cr") + s(PSQI_Comp5_Problems, k = 3, bs = "cr") +
##     s(PSQI_Comp5_Problems, by = Sex_ord, k = 3, bs = "cr") +
##     ti(Age, PSQI_Comp5_Problems, k = c(4, 3)) + ti(Age, PSQI_Comp5_Problems,
##     by = Sex_ord, k = c(4, 3)) + ICV_st + Sex_ord
##
## Parametric coefficients:
##             Estimate Std. Error t value Pr(>|t|)
## (Intercept)  0.02697    0.06594   0.409    0.683
## ICV_st       0.16671    0.01674   9.962   <2e-16 ***
## Sex_ord.L    0.25508    0.02583   9.876   <2e-16 ***
## ---
## Signif. codes:  0 '***' 0.001 '**' 0.01 '*' 0.05 '.' 0.1 ' ' 1
##
## Approximate significance of smooth terms:
##                                           edf Ref.df       F  p-value
## s(Age)                                  4.785  4.785 307.214  < 2e-16 ***
## s(Age):Sex_ordMale                      1.000  1.000  15.807 7.12e-05 ***
## s(PSQI_Comp5_Problems)                  1.000  1.000   0.106   0.7452
## s(PSQI_Comp5_Problems):Sex_ordMale      1.000  1.000   3.186   0.0743 .
## ti(Age,PSQI_Comp5_Problems)             1.000  1.000   0.012   0.9125
## ti(Age,PSQI_Comp5_Problems):Sex_ordMale 1.000  1.000   2.360   0.1246
## ---
## Signif. codes:  0 '***' 0.001 '**' 0.01 '*' 0.05 '.' 0.1 ' ' 1
##
## R-sq.(adj) =  0.414
##   Scale est. = 0.019193  n = 4690
```

##### Longitudinal Analysis

In this section, only participants with two timepoints or more are retained.

Interaction with time.

```
forms <- time_interaction(psqi_var)
m2 <- gamm(forms[[1]], random = forms[[2]],
           data = long_only(subset_data(data, psqi_var)))
saveRDS(m2, file = paste0("./models/", psqi_var, "_m2.rds"))
m2 <- readRDS(paste0("./models/", psqi_var, "_m2.rds"))
summary(m2$gam)
##
## Family: gaussian
## Link function: identity
##
## Formula:
## Hippocampus_st ~ te(dt, Age_bl, k = c(4, 10)) + ti(PSQI_Comp5_Problems,
##     k = 4) + ti(dt, PSQI_Comp5_Problems, k = c(4, 4)) + ICV_st +
##     Sex
##
## Parametric coefficients:
##             Estimate Std. Error t value Pr(>|t|)
## (Intercept) -0.05906    0.05745  -1.028    0.304
## ICV_st       0.27556    0.02632  10.470  < 2e-16 ***
## SexMale      0.22001    0.05179   4.248 2.22e-05 ***
## ---
## Signif. codes:  0 '***' 0.001 '**' 0.01 '*' 0.05 '.' 0.1 ' ' 1
##
## Approximate significance of smooth terms:
##                               edf Ref.df       F p-value
## te(dt,Age_bl)              15.961 15.961 110.884 < 2e-16 ***
## ti(PSQI_Comp5_Problems)     1.000  1.000   0.024 0.87758
## ti(dt,PSQI_Comp5_Problems)  2.215  2.215   5.947 0.00173 **
## ---
## Signif. codes:  0 '***' 0.001 '**' 0.01 '*' 0.05 '.' 0.1 ' ' 1
##
## R-sq.(adj) =  0.466
##   Scale est. = 0.019056  n = 2893
fvisgam(m2$gam, view = c("dt", psqi_var))
## Summary:
##  * ICV_st : numeric predictor; set to the value(s): -0.00764243395840525.
##  * Sex : factor; set to the value(s): Female.
##  * dt : numeric predictor; with 30 values ranging from 0.000000 to 10.976044.
##  * Age_bl : numeric predictor; set to the value(s): 64.8876712328767.
##  * PSQI_Comp5_Problems : numeric predictor; with 30 values ranging from 0.000000 to 3.000000.
```

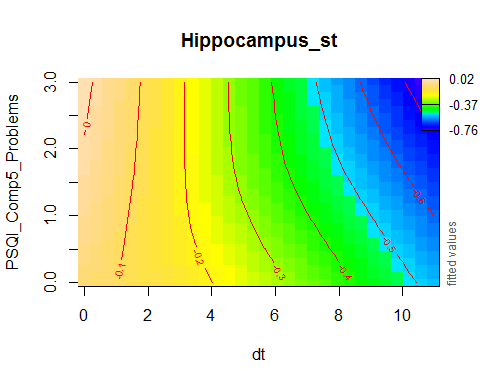

plot_psqi_interaction(m2, psqi_var, psqi_range = 0:3, time_range = 0:5)

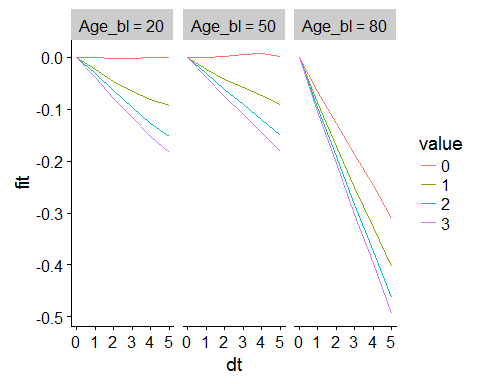

###### Controling for Covariates

```
forms <- time_interaction(psqi_var, c("Depression", "BMI"))
m2_cov <- gamm(forms[[1]], random = forms[[2]],
           data = long_only(subset_data(data, psqi_var)))
saveRDS(m2_cov, file = paste0("./models/", psqi_var, "_m2_cov.rds"))
m2_cov <- readRDS(paste0("./models/", psqi_var, "_m2_cov.rds"))
summary(m2_cov$gam)
##
## Family: gaussian
## Link function: identity
##
## Formula:
## Hippocampus_st ~ te(dt, Age_bl, k = c(4, 10)) + ti(PSQI_Comp5_Problems,
##     k = 4) + ti(dt, PSQI_Comp5_Problems, k = c(4, 4)) + ICV_st +
##     Sex + Depression + BMI
##
## Parametric coefficients:
##              Estimate Std. Error t value Pr(>|t|)
## (Intercept) -0.611317   0.202843  -3.014  0.00262 **
## ICV_st       0.315922   0.035898   8.800  < 2e-16 ***
## SexMale      0.139945   0.066606   2.101  0.03578 *
## Depression  -0.010581   0.029538  -0.358  0.72023
## BMI          0.023199   0.007353   3.155  0.00163 **
## ---
## Signif. codes:  0 '***' 0.001 '**' 0.01 '*' 0.05 '.' 0.1 ' ' 1
##
## Approximate significance of smooth terms:
##                              edf Ref.df      F p-value
## te(dt,Age_bl)              12.18  12.18 79.096 < 2e-16 ***
## ti(PSQI_Comp5_Problems)     1.00   1.00  0.052 0.82049
## ti(dt,PSQI_Comp5_Problems)  1.00   1.00 10.345 0.00132 **
## ---
## Signif. codes:  0 '***' 0.001 '**' 0.01 '*' 0.05 '.' 0.1 ' ' 1
##
## R-sq.(adj) =   0.49
##   Scale est. = 0.02308   n = 1689
```

###### Interaction with Sex and Time

As the summary shows, no sex interaction was found.

```
forms <- sex_time_interaction(psqi_var)
m2s <- gamm(forms[[1]], random = forms[[2]],
            data = long_only(subset_data(data, psqi_var)))

saveRDS(m2s, file = paste0("./models/", psqi_var, "_m2s.rds"))
m2s <- readRDS(paste0("./models/", psqi_var, "_m2s.rds"))
summary(m2s$gam)
##
## Family: gaussian
## Link function: identity
##
## Formula:
## Hippocampus_st ~ te(dt, Age_bl, k = c(4, 10)) + te(dt, Age_bl,
##     by = Sex_ord, k = c(4, 10)) + ti(PSQI_Comp5_Problems, k = 4) +
##     ti(PSQI_Comp5_Problems, by = Sex_ord, k = 4) + ti(dt, PSQI_Comp5_Problems,
##     k = c(4, 4)) + ti(dt, PSQI_Comp5_Problems, by = Sex_ord,
##     k = c(4, 4)) + ICV_st + Sex_ord
##
## Parametric coefficients:
##             Estimate Std. Error t value Pr(>|t|)
## (Intercept)  0.05552    0.04994   1.112    0.266
## ICV_st       0.27495    0.02629  10.460  < 2e-16 ***
## Sex_ord.L    0.15953    0.03668   4.350 1.41e-05 ***
## ---
## Signif. codes:  0 '***' 0.001 '**' 0.01 '*' 0.05 '.' 0.1 ' ' 1
##
## Approximate significance of smooth terms:
##                                           edf Ref.df      F p-value
## te(dt,Age_bl)                          15.837 15.837 58.982  <2e-16 ***
## te(dt,Age_bl):Sex_ordMale               4.843  4.843  1.835  0.0939 .
## ti(PSQI_Comp5_Problems)                 1.000  1.000  0.945  0.3312
## ti(PSQI_Comp5_Problems):Sex_ordMale     1.000  1.000  3.005  0.0831 .
## ti(dt,PSQI_Comp5_Problems)              2.372  2.372  3.473  0.0210 *
## ti(dt,PSQI_Comp5_Problems):Sex_ordMale  1.000  1.000  0.013  0.9106
## ---
## Signif. codes:  0 '***' 0.001 '**' 0.01 '*' 0.05 '.' 0.1 ' ' 1
##
## R-sq.(adj) =  0.469
##   Scale est. = 0.018981  n = 2893
```

#### PSQI Component 6: Medication

psqi_var <- "PSQI_Comp6_Medication"

Note that Betula does not have this component.

psqi_plot(data, psqi_var)

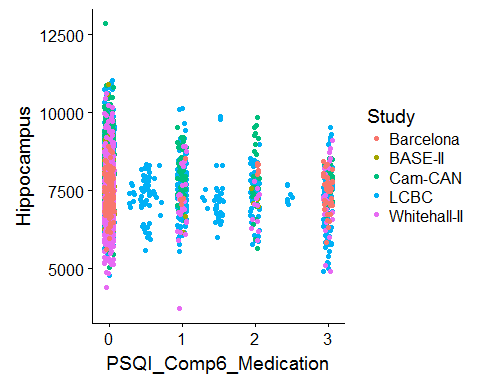

We fit one model per study first, purely for exploratory reasons.

sep_fits(data, psqi_var)

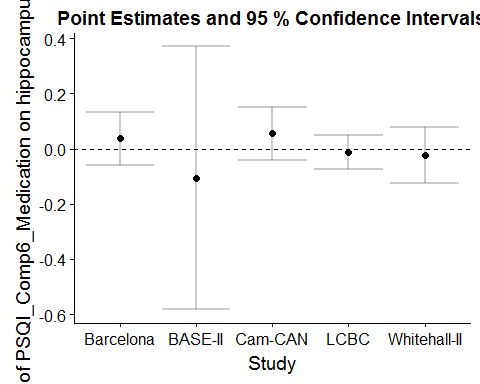

##### Main Effect of Sleep

GAMM with smooth effect of PSQI, random slope of PSQI per study, and random intercept of ID.

```
forms <- main_effects(psqi_var)
m0 <- gamm(forms[[1]], random = forms[[2]],
           data = subset_data(data, psqi_var))
saveRDS(m0, file = paste0("./models/", psqi_var, "_m0.rds"))
m0 <- readRDS(paste0("./models/", psqi_var, "_m0.rds"))
summary(m0$gam)
##
## Family: gaussian
## Link function: identity
##
## Formula:
## Hippocampus_st ~ s(Age, k = 6, bs = "cr") + s(PSQI_Comp6_Medication,
##     k = 3, bs = "cr") + ICV_st + Sex
##
## Parametric coefficients:
##             Estimate Std. Error t value Pr(>|t|)
## (Intercept) -0.20764    0.07872  -2.638  0.00838 **
## ICV_st       0.13994    0.01767   7.921 3.02e-15 ***
## SexMale      0.39993    0.04000   9.999  < 2e-16 ***
## ---
## Signif. codes:  0 '***' 0.001 '**' 0.01 '*' 0.05 '.' 0.1 ' ' 1
##
## Approximate significance of smooth terms:
##                            edf Ref.df       F p-value
## s(Age)                   4.759  4.759 477.891  <2e-16 ***
## s(PSQI_Comp6_Medication) 1.000  1.000   0.051   0.821
## ---
## Signif. codes:  0 '***' 0.001 '**' 0.01 '*' 0.05 '.' 0.1 ' ' 1
##
## R-sq.(adj) =  0.407
##   Scale est. = 0.018273  n = 4128
plot(m0$gam, select = 2, seWithMean = TRUE)
```

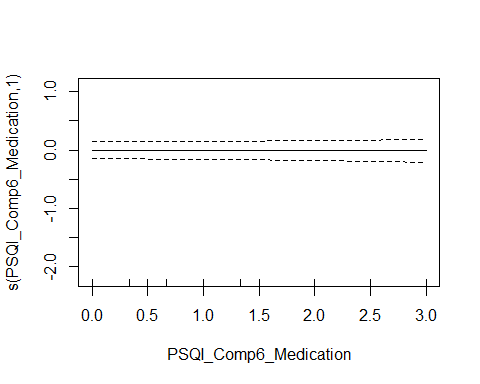

###### Controling for Covariates

```
forms <- main_effects(psqi_var, c("Depression", "BMI"))
m0_cov <- gamm(forms[[1]], random = forms[[2]],
           data = subset_data(data, psqi_var))
saveRDS(m0_cov, file = paste0("./models/", psqi_var, "_m0_cov.rds"))
m0_cov <- readRDS(paste0("./models/", psqi_var, "_m0_cov.rds"))
summary(m0_cov$gam)
##
## Family: gaussian
## Link function: identity
##
## Formula:
## Hippocampus_st ~ s(Age, k = 6, bs = "cr") + s(PSQI_Comp6_Medication,
##     k = 3, bs = "cr") + ICV_st + Sex + Depression + BMI
##
## Parametric coefficients:
##              Estimate Std. Error t value Pr(>|t|)
## (Intercept) -0.531492   0.179991  -2.953  0.00318 **
## ICV_st       0.039777   0.024255   1.640  0.10116
## SexMale      0.398689   0.059360   6.716 2.37e-11 ***
## Depression   0.037296   0.023896   1.561  0.11872
## BMI          0.007550   0.005774   1.307  0.19121
## ---
## Signif. codes:  0 '***' 0.001 '**' 0.01 '*' 0.05 '.' 0.1 ' ' 1
##
## Approximate significance of smooth terms:
##                            edf Ref.df       F p-value
## s(Age)                   4.374  4.374 237.486  <2e-16 ***
## s(PSQI_Comp6_Medication) 1.000  1.000   0.268   0.605
## ---
## Signif. codes:  0 '***' 0.001 '**' 0.01 '*' 0.05 '.' 0.1 ' ' 1
##
## R-sq.(adj) =  0.367
##   Scale est. = 0.022431  n = 2175
```

###### Interaction with Sex

The summary shows that no sex interaction is found.

```
forms <- main_sex_interaction(psqi_var)
m0s <- gamm(forms[[1]], random = forms[[2]],
           data = subset_data(data, psqi_var), method = "REML")

saveRDS(m0s, file = paste0("./models/", psqi_var, "_m0s.rds"))
m0s <- readRDS(paste0("./models/", psqi_var, "_m0s.rds"))
summary(m0s$gam)
##
## Family: gaussian
## Link function: identity
##
## Formula:
## Hippocampus_st ~ s(Age, k = 6, bs = "cr") + s(PSQI_Comp6_Medication,
##     k = 3, bs = "cr") + s(PSQI_Comp6_Medication, by = Sex_ord,
##     k = 3, bs = "cr") + ICV_st + Sex_ord
##
## Parametric coefficients:
##             Estimate Std. Error t value Pr(>|t|)
## (Intercept) -0.01050    0.08419  -0.125    0.901
## ICV_st       0.14017    0.01768   7.926 2.88e-15 ***
## Sex_ord.L    0.28140    0.02839   9.913  < 2e-16 ***
## ---
## Signif. codes:  0 '***' 0.001 '**' 0.01 '*' 0.05 '.' 0.1 ' ' 1
##
## Approximate significance of smooth terms:
##                                        edf Ref.df       F p-value
## s(Age)                               4.762  4.762 477.390  <2e-16 ***
## s(PSQI_Comp6_Medication)             1.000  1.000   0.278   0.598
## s(PSQI_Comp6_Medication):Sex_ordMale 1.000  1.000   1.274   0.259
## ---
## Signif. codes:  0 '***' 0.001 '**' 0.01 '*' 0.05 '.' 0.1 ' ' 1
##
## R-sq.(adj) =  0.407
##   Scale est. = 0.01828   n = 4128
```

##### Interaction with Age

```
forms <- age_interaction(psqi_var)
m1 <- gamm(forms[[1]], random = forms[[2]],
           data = subset_data(data, psqi_var), method = "REML")

saveRDS(m1, file = paste0("./models/", psqi_var, "_m1.rds"))
m1 <- readRDS(paste0("./models/", psqi_var, "_m1.rds"))
summary(m1$gam)
##
## Family: gaussian
## Link function: identity
##
## Formula:
## Hippocampus_st ~ s(Age, k = 6, bs = "cr") + s(PSQI_Comp6_Medication,
##     bs = "cr", k = 3) + ti(Age, PSQI_Comp6_Medication, k = c(4,
##     3)) + ICV_st + Sex
##
## Parametric coefficients:
##             Estimate Std. Error t value Pr(>|t|)
## (Intercept) -0.21033    0.08679  -2.423   0.0154 *
## ICV_st       0.13930    0.01769   7.876 4.29e-15 ***
## SexMale      0.40186    0.04005  10.035  < 2e-16 ***
## ---
## Signif. codes:  0 '***' 0.001 '**' 0.01 '*' 0.05 '.' 0.1 ' ' 1
##
## Approximate significance of smooth terms:
##                                 edf Ref.df       F p-value
## s(Age)                        4.764  4.764 476.785  <2e-16 ***
## s(PSQI_Comp6_Medication)      1.000  1.000   0.246   0.620
## ti(Age,PSQI_Comp6_Medication) 1.000  1.000   0.775   0.379
## ---
## Signif. codes:  0 '***' 0.001 '**' 0.01 '*' 0.05 '.' 0.1 ' ' 1
##
## R-sq.(adj) =  0.407
##   Scale est. = 0.018281  n = 4128
par(mfrow = c(1, 2))
plot(m1$gam, select = 2, seWithMean = TRUE)
fvisgam(m1$gam, view = c("Age", psqi_var))
## Summary:
##  * ICV_st : numeric predictor; set to the value(s): 0.0283618430856087.
##  * Sex : factor; set to the value(s): Female.
##  * Age : numeric predictor; with 30 values ranging from 18.482192 to 91.885010.
##  * PSQI_Comp6_Medication : numeric predictor; with 30 values ranging from 0.000000 to 3.000000.
```

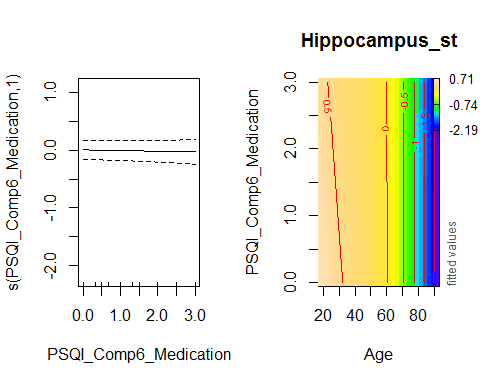

###### Controling for Covariates

```
forms <- age_interaction(psqi_var, c("Depression", "BMI"))
m1_cov <- gamm(forms[[1]], random = forms[[2]],
           data = subset_data(data, psqi_var))

saveRDS(m1_cov, file = paste0("./models/", psqi_var, "_m1_cov.rds"))
m1_cov <- readRDS(paste0("./models/", psqi_var, "_m1_cov.rds"))
summary(m1_cov$gam)
##
## Family: gaussian
## Link function: identity
##
## Formula:
## Hippocampus_st ~ s(Age, k = 6, bs = "cr") + s(PSQI_Comp6_Medication,
##     bs = "cr", k = 3) + ti(Age, PSQI_Comp6_Medication, k = c(4,
##     3)) + ICV_st + Sex + Depression + BMI
##
## Parametric coefficients:
##              Estimate Std. Error t value Pr(>|t|)
## (Intercept) -0.529762   0.180076  -2.942   0.0033 **
## ICV_st       0.039845   0.024256   1.643   0.1006
## SexMale      0.398139   0.059396   6.703 2.59e-11 ***
## Depression   0.037189   0.023899   1.556   0.1198
## BMI          0.007549   0.005775   1.307   0.1912
## ---
## Signif. codes:  0 '***' 0.001 '**' 0.01 '*' 0.05 '.' 0.1 ' ' 1
##
## Approximate significance of smooth terms:
##                                 edf Ref.df       F p-value
## s(Age)                        4.368  4.368 237.819  <2e-16 ***
## s(PSQI_Comp6_Medication)      1.000  1.000   0.088   0.767
## ti(Age,PSQI_Comp6_Medication) 1.000  1.000   0.074   0.786
## ---
## Signif. codes:  0 '***' 0.001 '**' 0.01 '*' 0.05 '.' 0.1 ' ' 1
##
## R-sq.(adj) =  0.367
##   Scale est. = 0.022428  n = 2175
```

###### Interaction with Age and Sex

No sex interaction in the interaction between Age and Sleep was found:

```
forms <- age_sex_interaction(psqi_var)
m1s <- gamm(forms[[1]], random = forms[[2]],
            data = subset_data(data, psqi_var))

saveRDS(m1s, file = paste0("./models/", psqi_var, "_m1s.rds"))
m1s <- readRDS(paste0("./models/", psqi_var, "_m1s.rds"))
summary(m1s$gam)
##
## Family: gaussian
## Link function: identity
##
## Formula:
## Hippocampus_st ~ s(Age, k = 6, bs = "cr") + s(Age, by = Sex_ord,
##     k = 3, bs = "cr") + s(PSQI_Comp6_Medication, k = 3, bs = "cr") +
##     s(PSQI_Comp6_Medication, by = Sex_ord, k = 3, bs = "cr") +
##     ti(Age, PSQI_Comp6_Medication, k = c(4, 3)) + ti(Age, PSQI_Comp6_Medication,
##     by = Sex_ord, k = c(4, 3)) + ICV_st + Sex_ord
##
## Parametric coefficients:
##               Estimate Std. Error t value Pr(>|t|)
## (Intercept) -0.0005794  0.0695099  -0.008    0.993
## ICV_st       0.1398202  0.0176267   7.932 2.75e-15 ***
## Sex_ord.L    0.2723162  0.0286227   9.514  < 2e-16 ***
## ---
## Signif. codes:  0 '***' 0.001 '**' 0.01 '*' 0.05 '.' 0.1 ' ' 1
##
## Approximate significance of smooth terms:
##                                             edf Ref.df       F  p-value
## s(Age)                                    4.753  4.753 272.828  < 2e-16
## s(Age):Sex_ordMale                        1.000  1.000  17.233 3.37e-05
## s(PSQI_Comp6_Medication)                  1.000  1.000   0.000    0.992
## s(PSQI_Comp6_Medication):Sex_ordMale      1.000  1.000   0.667    0.414
## ti(Age,PSQI_Comp6_Medication)             1.605  1.605   0.048    0.881
## ti(Age,PSQI_Comp6_Medication):Sex_ordMale 1.000  1.000   0.229    0.632
##
## s(Age)                                    ***
## s(Age):Sex_ordMale                        ***
## s(PSQI_Comp6_Medication)
## s(PSQI_Comp6_Medication):Sex_ordMale
## ti(Age,PSQI_Comp6_Medication)
## ti(Age,PSQI_Comp6_Medication):Sex_ordMale
## ---
## Signif. codes:  0 '***' 0.001 '**' 0.01 '*' 0.05 '.' 0.1 ' ' 1
##
## R-sq.(adj) =  0.413
##   Scale est. = 0.018186  n = 4128
```

##### Longitudinal Analysis

In this section, only participants with two timepoints or more are retained.

Interaction with time.

```
forms <- time_interaction(psqi_var)
m2 <- gamm(forms[[1]], random = forms[[2]],
           data = long_only(subset_data(data, psqi_var)))
saveRDS(m2, file = paste0("./models/", psqi_var, "_m2.rds"))
m2 <- readRDS(paste0("./models/", psqi_var, "_m2.rds"))
summary(m2$gam)
##
## Family: gaussian
## Link function: identity
##
## Formula:
## Hippocampus_st ~ te(dt, Age_bl, k = c(4, 10)) + ti(PSQI_Comp6_Medication,
##     k = 4) + ti(dt, PSQI_Comp6_Medication, k = c(4, 4)) + ICV_st +
##     Sex
##
## Parametric coefficients:
##             Estimate Std. Error t value Pr(>|t|)
## (Intercept) -0.12819    0.04701  -2.727  0.00644 **
## ICV_st       0.26404    0.02744   9.622  < 2e-16 ***
## SexMale      0.25857    0.05558   4.652 3.46e-06 ***
## ---
## Signif. codes:  0 '***' 0.001 '**' 0.01 '*' 0.05 '.' 0.1 ' ' 1
##
## Approximate significance of smooth terms:
##                                 edf Ref.df       F p-value
## te(dt,Age_bl)                17.468 17.468 101.683  <2e-16 ***
## ti(PSQI_Comp6_Medication)     1.000  1.000   0.075   0.784
## ti(dt,PSQI_Comp6_Medication)  1.901  1.901   1.754   0.128
## ---
## Signif. codes:  0 '***' 0.001 '**' 0.01 '*' 0.05 '.' 0.1 ' ' 1
##
## R-sq.(adj) =  0.482
##   Scale est. = 0.017968  n = 2519
fvisgam(m2$gam, view = c("dt", psqi_var))
## Summary:
##  * ICV_st : numeric predictor; set to the value(s): -0.000955268254596884.
##  * Sex : factor; set to the value(s): Female.
##  * dt : numeric predictor; with 30 values ranging from 0.000000 to 10.976044.
##  * Age_bl : numeric predictor; set to the value(s): 65.2867898699521.
##  * PSQI_Comp6_Medication : numeric predictor; with 30 values ranging from 0.000000 to 3.000000.
```

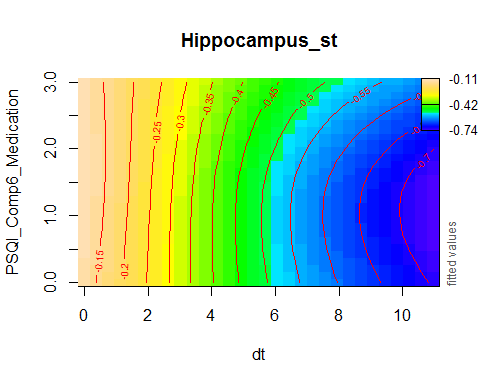

plot_psqi_interaction(m2, psqi_var, psqi_range = 0:3, time_range = 0:5)

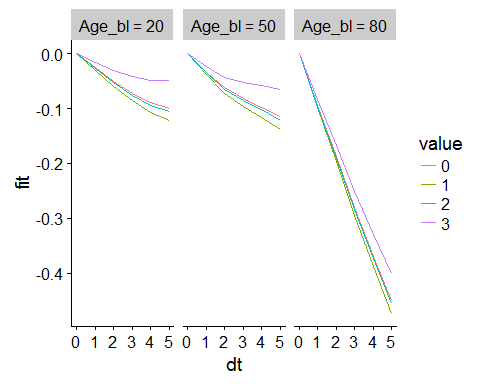

###### Controling for Covariates

```
forms <- time_interaction(psqi_var, c("Depression", "BMI"))
m2_cov <- gamm(forms[[1]], random = forms[[2]],
           data = long_only(subset_data(data, psqi_var)), optim = list(opt = "optim"))
saveRDS(m2_cov, file = paste0("./models/", psqi_var, "_m2_cov.rds"))
m2_cov <- readRDS(paste0("./models/", psqi_var, "_m2_cov.rds"))
summary(m2_cov$gam)
##
## Family: gaussian
## Link function: identity
##
## Formula:
## Hippocampus_st ~ te(dt, Age_bl, k = c(4, 10)) + ti(PSQI_Comp6_Medication,
##     k = 4) + ti(dt, PSQI_Comp6_Medication, k = c(4, 4)) + ICV_st +
##     Sex + Depression + BMI
##
## Parametric coefficients:
##             Estimate Std. Error t value Pr(>|t|)
## (Intercept) -0.75277    0.22016  -3.419 0.000648 ***
## ICV_st       0.29203    0.04146   7.044 3.02e-12 ***
## SexMale      0.19734    0.07760   2.543 0.011106 *
## Depression   0.01087    0.03325   0.327 0.743867
## BMI          0.02499    0.00843   2.965 0.003085 **
## ---
## Signif. codes:  0 '***' 0.001 '**' 0.01 '*' 0.05 '.' 0.1 ' ' 1
##
## Approximate significance of smooth terms:
##                                 edf Ref.df      F p-value
## te(dt,Age_bl)                11.521 11.521 78.634  <2e-16 ***
## ti(PSQI_Comp6_Medication)     1.000  1.000  0.215   0.643
## ti(dt,PSQI_Comp6_Medication)  1.905  1.905  0.985   0.363
## ---
## Signif. codes:  0 '***' 0.001 '**' 0.01 '*' 0.05 '.' 0.1 ' ' 1
##
## R-sq.(adj) =  0.524
##   Scale est. = 0.022229  n = 1311
```

###### Interaction with Sex and Time

As the summary shows, no sex interaction was found.

```
forms <- sex_time_interaction(psqi_var)
m2s <- gamm(forms[[1]], random = forms[[2]],
            data = long_only(subset_data(data, psqi_var)),
            control = list(opt = "optim"))

saveRDS(m2s, file = paste0("./models/", psqi_var, "_m2s.rds"))
m2s <- readRDS(paste0("./models/", psqi_var, "_m2s.rds"))
summary(m2s$gam)
##
## Family: gaussian
## Link function: identity
##
## Formula:
## Hippocampus_st ~ te(dt, Age_bl, k = c(4, 10)) + te(dt, Age_bl,
##     by = Sex_ord, k = c(4, 10)) + ti(PSQI_Comp6_Medication, k = 4) +
##     ti(PSQI_Comp6_Medication, by = Sex_ord, k = 4) + ti(dt, PSQI_Comp6_Medication,
##     k = c(4, 4)) + ti(dt, PSQI_Comp6_Medication, by = Sex_ord,
##     k = c(4, 4)) + ICV_st + Sex_ord
##
## Parametric coefficients:
##              Estimate Std. Error t value Pr(>|t|)
## (Intercept) 0.0007457  0.0380440   0.020    0.984
## ICV_st      0.2664381  0.0274015   9.723  < 2e-16 ***
## Sex_ord.L   0.1764624  0.0393964   4.479 7.83e-06 ***
## ---
## Signif. codes:  0 '***' 0.001 '**' 0.01 '*' 0.05 '.' 0.1 ' ' 1
##
## Approximate significance of smooth terms:
##                                             edf Ref.df      F p-value
## te(dt,Age_bl)                            17.859 17.859 49.714 < 2e-16 ***
## te(dt,Age_bl):Sex_ordMale                 3.000  3.000  4.123 0.00629 **
## ti(PSQI_Comp6_Medication)                 1.000  1.000  0.077 0.78103
## ti(PSQI_Comp6_Medication):Sex_ordMale     1.000  1.000  0.115 0.73459
## ti(dt,PSQI_Comp6_Medication)              1.967  1.967  2.778 0.07782 .
## ti(dt,PSQI_Comp6_Medication):Sex_ordMale  2.126  2.126  4.898 0.00676 **
## ---
## Signif. codes:  0 '***' 0.001 '**' 0.01 '*' 0.05 '.' 0.1 ' ' 1
##
## R-sq.(adj) =  0.483
##   Scale est. = 0.01774   n = 2519
```

#### PSQI Component 7: Tiredness

psqi_var <- "PSQI_Comp7_Tired"

psqi_plot(data, psqi_var)

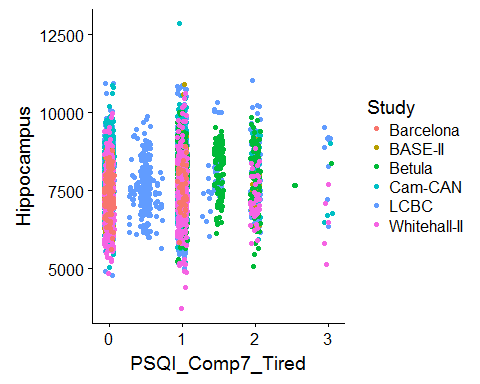

We fit one model per study first, purely for exploratory reasons.

sep_fits(data, psqi_var)

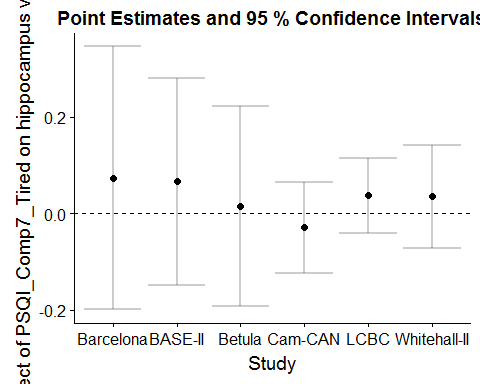

##### Main Effect of Sleep

GAMM with smooth effect of PSQI, random slope of PSQI per study, and random intercept of ID.

```
forms <- main_effects(psqi_var)
m0 <- gamm(forms[[1]], random = forms[[2]],
           data = subset_data(data, psqi_var))
saveRDS(m0, file = paste0("./models/", psqi_var, "_m0.rds"))
m0 <- readRDS(paste0("./models/", psqi_var, "_m0.rds"))
summary(m0$gam)
##
## Family: gaussian
## Link function: identity
##
## Formula:
## Hippocampus_st ~ s(Age, k = 6, bs = "cr") + s(PSQI_Comp7_Tired,
##     k = 3, bs = "cr") + ICV_st + Sex
##
## Parametric coefficients:
##             Estimate Std. Error t value Pr(>|t|)
## (Intercept) -0.16068    0.07224  -2.224   0.0262 *
## ICV_st       0.16260    0.01690   9.618   <2e-16 ***
## SexMale      0.35827    0.03704   9.672   <2e-16 ***
## ---
## Signif. codes:  0 '***' 0.001 '**' 0.01 '*' 0.05 '.' 0.1 ' ' 1
##
## Approximate significance of smooth terms:
##                       edf Ref.df       F p-value
## s(Age)              4.790  4.790 501.792  <2e-16 ***
## s(PSQI_Comp7_Tired) 1.777  1.777   3.327   0.106
## ---
## Signif. codes:  0 '***' 0.001 '**' 0.01 '*' 0.05 '.' 0.1 ' ' 1
##
## R-sq.(adj) =  0.408
##   Scale est. = 0.01911   n = 4615
plot(m0$gam, select = 2, seWithMean = TRUE)
```

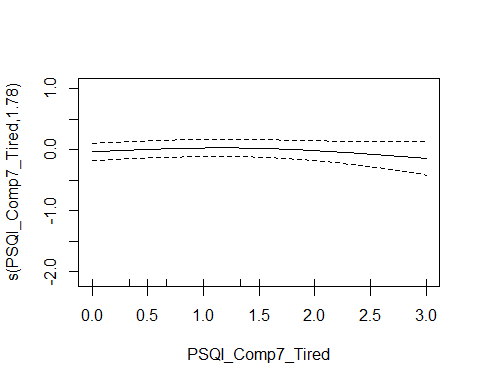

###### Controling for Covariates

```
forms <- main_effects(psqi_var, c("Depression", "BMI"))
m0_cov <- gamm(forms[[1]], random = forms[[2]],
           data = subset_data(data, psqi_var))
saveRDS(m0_cov, file = paste0("./models/", psqi_var, "_m0_cov.rds"))
m0_cov <- readRDS(paste0("./models/", psqi_var, "_m0_cov.rds"))
summary(m0_cov$gam)
##
## Family: gaussian
## Link function: identity
##
## Formula:
## Hippocampus_st ~ s(Age, k = 6, bs = "cr") + s(PSQI_Comp7_Tired,
##     k = 3, bs = "cr") + ICV_st + Sex + Depression + BMI
##
## Parametric coefficients:
##               Estimate Std. Error t value Pr(>|t|)
## (Intercept) -0.4073157  0.1643057  -2.479 0.013236 *
## ICV_st       0.0782120  0.0222560   3.514 0.000448 ***
## SexMale      0.3568795  0.0511759   6.974 3.89e-12 ***
## Depression  -0.0003101  0.0227201  -0.014 0.989113
## BMI          0.0061114  0.0051802   1.180 0.238204
## ---
## Signif. codes:  0 '***' 0.001 '**' 0.01 '*' 0.05 '.' 0.1 ' ' 1
##
## Approximate significance of smooth terms:
##                       edf Ref.df       F p-value
## s(Age)              4.515  4.515 262.014  <2e-16 ***
## s(PSQI_Comp7_Tired) 1.874  1.874   5.432  0.0207 *
## ---
## Signif. codes:  0 '***' 0.001 '**' 0.01 '*' 0.05 '.' 0.1 ' ' 1
##
## R-sq.(adj) =  0.372
##   Scale est. = 0.02315   n = 2665
```

###### Interaction with Sex

The summary shows that no sex interaction is found.

```
forms <- main_sex_interaction(psqi_var)
m0s <- gamm(forms[[1]], random = forms[[2]],
           data = subset_data(data, psqi_var), method = "REML")

saveRDS(m0s, file = paste0("./models/", psqi_var, "_m0s.rds"))
m0s <- readRDS(paste0("./models/", psqi_var, "_m0s.rds"))
summary(m0s$gam)
##
## Family: gaussian
## Link function: identity
##
## Formula:
## Hippocampus_st ~ s(Age, k = 6, bs = "cr") + s(PSQI_Comp7_Tired,
##     k = 3, bs = "cr") + s(PSQI_Comp7_Tired, by = Sex_ord, k = 3,
##     bs = "cr") + ICV_st + Sex_ord
##
## Parametric coefficients:
##             Estimate Std. Error t value Pr(>|t|)
## (Intercept)  0.01832    0.07557   0.242    0.808
## ICV_st       0.16237    0.01691   9.601   <2e-16 ***
## Sex_ord.L    0.25378    0.02622   9.679   <2e-16 ***
## ---
## Signif. codes:  0 '***' 0.001 '**' 0.01 '*' 0.05 '.' 0.1 ' ' 1
##
## Approximate significance of smooth terms:
##                                   edf Ref.df       F p-value
## s(Age)                          4.793  4.793 501.357  <2e-16 ***
## s(PSQI_Comp7_Tired)             1.233  1.233   0.244   0.762
## s(PSQI_Comp7_Tired):Sex_ordMale 1.773  1.773   2.133   0.190
## ---
## Signif. codes:  0 '***' 0.001 '**' 0.01 '*' 0.05 '.' 0.1 ' ' 1
##
## R-sq.(adj) =  0.408
##   Scale est. = 0.019115  n = 4615
```

##### Interaction with Age

```
forms <- age_interaction(psqi_var)
m1 <- gamm(forms[[1]], random = forms[[2]],
           data = subset_data(data, psqi_var), control = list(opt = "optim"))

saveRDS(m1, file = paste0("./models/", psqi_var, "_m1.rds"))
m1 <- readRDS(paste0("./models/", psqi_var, "_m1.rds"))
summary(m1$gam)
##
## Family: gaussian
## Link function: identity
##
## Formula:
## Hippocampus_st ~ s(Age, k = 6, bs = "cr") + s(PSQI_Comp7_Tired,
##     bs = "cr", k = 3) + ti(Age, PSQI_Comp7_Tired, k = c(4, 3)) +
##     ICV_st + Sex
##
## Parametric coefficients:
##             Estimate Std. Error t value Pr(>|t|)
## (Intercept) -0.16257    0.07245  -2.244   0.0249 *
## ICV_st       0.16273    0.01690   9.629   <2e-16 ***
## SexMale      0.35822    0.03703   9.674   <2e-16 ***
## ---
## Signif. codes:  0 '***' 0.001 '**' 0.01 '*' 0.05 '.' 0.1 ' ' 1
##
## Approximate significance of smooth terms:
##                            edf Ref.df       F p-value
## s(Age)                   4.791  4.791 501.237  <2e-16 ***
## s(PSQI_Comp7_Tired)      1.798  1.798   3.478  0.0863 .
## ti(Age,PSQI_Comp7_Tired) 1.098  1.098   0.814  0.3517
## ---
## Signif. codes:  0 '***' 0.001 '**' 0.01 '*' 0.05 '.' 0.1 ' ' 1
##
## R-sq.(adj) =  0.408
##   Scale est. = 0.019122  n = 4615
par(mfrow = c(1, 2))
plot(m1$gam, select = 2, seWithMean = TRUE)
fvisgam(m1$gam, view = c("Age", psqi_var))
## Summary:
##  * ICV_st : numeric predictor; set to the value(s): 0.0114354964127359.
##  * Sex : factor; set to the value(s): Female.
##  * Age : numeric predictor; with 30 values ranging from 18.482192 to 91.885010.
##  * PSQI_Comp7_Tired : numeric predictor; with 30 values ranging from 0.000000 to 3.000000.
```

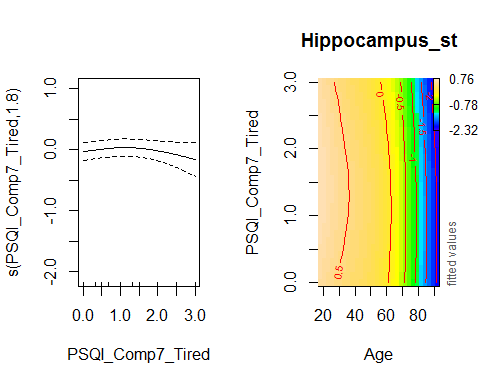

###### Controling for Covariates

```
forms <- age_interaction(psqi_var, c("Depression", "BMI"))
m1_cov <- gamm(forms[[1]], random = forms[[2]],
           data = subset_data(data, psqi_var))

saveRDS(m1_cov, file = paste0("./models/", psqi_var, "_m1_cov.rds"))
m1_cov <- readRDS(paste0("./models/", psqi_var, "_m1_cov.rds"))
summary(m1_cov$gam)
##
## Family: gaussian
## Link function: identity
##
## Formula:
## Hippocampus_st ~ s(Age, k = 6, bs = "cr") + s(PSQI_Comp7_Tired,
##     bs = "cr", k = 3) + ti(Age, PSQI_Comp7_Tired, k = c(4, 3)) +
##     ICV_st + Sex + Depression + BMI
##
## Parametric coefficients:
##               Estimate Std. Error t value Pr(>|t|)
## (Intercept) -0.4056776  0.1640027  -2.474 0.013438 *
## ICV_st       0.0767771  0.0222531   3.450 0.000569 ***
## SexMale      0.3598989  0.0511700   7.033 2.55e-12 ***
## Depression   0.0004246  0.0227181   0.019 0.985091
## BMI          0.0060501  0.0051786   1.168 0.242795
## ---
## Signif. codes:  0 '***' 0.001 '**' 0.01 '*' 0.05 '.' 0.1 ' ' 1
##
## Approximate significance of smooth terms:
##                            edf Ref.df       F p-value
## s(Age)                   4.514  4.514 263.828  <2e-16 ***
## s(PSQI_Comp7_Tired)      1.878  1.878   5.031  0.0231 *
## ti(Age,PSQI_Comp7_Tired) 1.825  1.825   2.878  0.0885 .
## ---
## Signif. codes:  0 '***' 0.001 '**' 0.01 '*' 0.05 '.' 0.1 ' ' 1
##
## R-sq.(adj) =  0.372
##   Scale est. = 0.023051  n = 2665
```

###### Interaction with Age and Sex

No sex interaction in the interaction between Age and Sleep was found:

```
forms <- age_sex_interaction(psqi_var)
m1s <- gamm(forms[[1]], random = forms[[2]],
            data = subset_data(data, psqi_var), control = list(opt = "optim"))

saveRDS(m1s, file = paste0("./models/", psqi_var, "_m1s.rds"))
m1s <- readRDS(paste0("./models/", psqi_var, "_m1s.rds"))
summary(m1s$gam)
##
## Family: gaussian
## Link function: identity
##
## Formula:
## Hippocampus_st ~ s(Age, k = 6, bs = "cr") + s(Age, by = Sex_ord,
##     k = 3, bs = "cr") + s(PSQI_Comp7_Tired, k = 3, bs = "cr") +
##     s(PSQI_Comp7_Tired, by = Sex_ord, k = 3, bs = "cr") + ti(Age,
##     PSQI_Comp7_Tired, k = c(4, 3)) + ti(Age, PSQI_Comp7_Tired,
##     by = Sex_ord, k = c(4, 3)) + ICV_st + Sex_ord
##
## Parametric coefficients:
##             Estimate Std. Error t value Pr(>|t|)
## (Intercept)  0.02458    0.06684   0.368    0.713
## ICV_st       0.16274    0.01687   9.650   <2e-16 ***
## Sex_ord.L    0.24869    0.02618   9.498   <2e-16 ***
## ---
## Signif. codes:  0 '***' 0.001 '**' 0.01 '*' 0.05 '.' 0.1 ' ' 1
##
## Approximate significance of smooth terms:
##                                        edf Ref.df       F  p-value
## s(Age)                               4.794  4.794 287.418  < 2e-16 ***
## s(Age):Sex_ordMale                   1.104  1.104  14.098 0.000108 ***
## s(PSQI_Comp7_Tired)                  1.799  1.799   2.493 0.122437
## s(PSQI_Comp7_Tired):Sex_ordMale      1.013  1.013   0.207 0.661308
## ti(Age,PSQI_Comp7_Tired)             1.086  1.086   0.625 0.417263
## ti(Age,PSQI_Comp7_Tired):Sex_ordMale 1.186  1.186   0.060 0.883596
## ---
## Signif. codes:  0 '***' 0.001 '**' 0.01 '*' 0.05 '.' 0.1 ' ' 1
##
## R-sq.(adj) =  0.413
##   Scale est. = 0.019099  n = 4615
```

##### Longitudinal Analysis

In this section, only participants with two timepoints or more are retained.

Interaction with time.

```
forms <- time_interaction(psqi_var)
m2 <- gamm(forms[[1]], random = forms[[2]],
           data = long_only(subset_data(data, psqi_var)))
saveRDS(m2, file = paste0("./models/", psqi_var, "_m2.rds"))
m2 <- readRDS(paste0("./models/", psqi_var, "_m2.rds"))
summary(m2$gam)
##
## Family: gaussian
## Link function: identity
##
## Formula:
## Hippocampus_st ~ te(dt, Age_bl, k = c(4, 10)) + ti(PSQI_Comp7_Tired,
##     k = 4) + ti(dt, PSQI_Comp7_Tired, k = c(4, 4)) + ICV_st +
##     Sex
##
## Parametric coefficients:
##             Estimate Std. Error t value Pr(>|t|)
## (Intercept) -0.06058    0.05241  -1.156    0.248
## ICV_st       0.28283    0.02611  10.832  < 2e-16 ***
## SexMale      0.20962    0.05138   4.080 4.62e-05 ***
## ---
## Signif. codes:  0 '***' 0.001 '**' 0.01 '*' 0.05 '.' 0.1 ' ' 1
##
## Approximate significance of smooth terms:
##                            edf Ref.df       F  p-value
## te(dt,Age_bl)           15.478 15.478 117.772  < 2e-16 ***
## ti(PSQI_Comp7_Tired)     1.000  1.000   2.063    0.151
## ti(dt,PSQI_Comp7_Tired)  4.174  4.174   8.989 4.13e-07 ***
## ---
## Signif. codes:  0 '***' 0.001 '**' 0.01 '*' 0.05 '.' 0.1 ' ' 1
##
## R-sq.(adj) =  0.468
##   Scale est. = 0.018761  n = 2893
fvisgam(m2$gam, view = c("dt", psqi_var))
## Summary:
##  * ICV_st : numeric predictor; set to the value(s): -0.00764243395840525.
##  * Sex : factor; set to the value(s): Female.
##  * dt : numeric predictor; with 30 values ranging from 0.000000 to 10.976044.
##  * Age_bl : numeric predictor; set to the value(s): 64.8876712328767.
##  * PSQI_Comp7_Tired : numeric predictor; with 30 values ranging from 0.000000 to 3.000000.
```

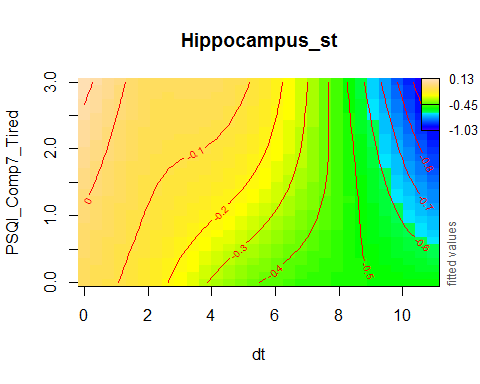

plot_psqi_interaction(m2, psqi_var, psqi_range = 0:3, time_range = 0:5)

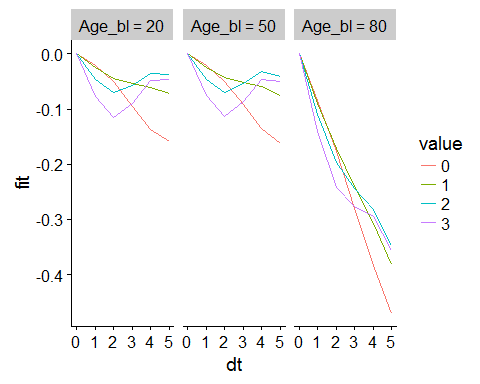

###### Controling for Covariates

```
forms <- time_interaction(psqi_var, c("Depression", "BMI"))
m2_cov <- gamm(forms[[1]], random = forms[[2]],
           data = long_only(subset_data(data, psqi_var)))
saveRDS(m2_cov, file = paste0("./models/", psqi_var, "_m2_cov.rds"))
m2_cov <- readRDS(paste0("./models/", psqi_var, "_m2_cov.rds"))
summary(m2_cov$gam)
##
## Family: gaussian
## Link function: identity
##
## Formula:
## Hippocampus_st ~ te(dt, Age_bl, k = c(4, 10)) + ti(PSQI_Comp7_Tired,
##     k = 4) + ti(dt, PSQI_Comp7_Tired, k = c(4, 4)) + ICV_st +
##     Sex + Depression + BMI
##
## Parametric coefficients:
##              Estimate Std. Error t value Pr(>|t|)
## (Intercept) -0.612819   0.196465  -3.119  0.00184 **
## ICV_st       0.326338   0.035738   9.131  < 2e-16 ***
## SexMale      0.123534   0.066186   1.866  0.06215 .
## Depression  -0.031872   0.031016  -1.028  0.30429
## BMI          0.023089   0.007315   3.156  0.00163 **
## ---
## Signif. codes:  0 '***' 0.001 '**' 0.01 '*' 0.05 '.' 0.1 ' ' 1
##
## Approximate significance of smooth terms:
##                            edf Ref.df      F  p-value
## te(dt,Age_bl)           11.713 11.713 84.619  < 2e-16 ***
## ti(PSQI_Comp7_Tired)     1.000  1.000  6.177    0.013 *
## ti(dt,PSQI_Comp7_Tired)  4.424  4.424  7.662 1.97e-06 ***
## ---
## Signif. codes:  0 '***' 0.001 '**' 0.01 '*' 0.05 '.' 0.1 ' ' 1
##
## R-sq.(adj) =  0.498
##   Scale est. = 0.022379  n = 1689
```

###### Interaction with Sex and Time

As the summary shows, no sex interaction was found.

```
forms <- sex_time_interaction(psqi_var)
m2s <- gamm(forms[[1]], random = forms[[2]],
            data = long_only(subset_data(data, psqi_var)), method = "REML")

saveRDS(m2s, file = paste0("./models/", psqi_var, "_m2s.rds"))
m2s <- readRDS(paste0("./models/", psqi_var, "_m2s.rds"))
summary(m2s$gam)
##
## Family: gaussian
## Link function: identity
##
## Formula:
## Hippocampus_st ~ te(dt, Age_bl, k = c(4, 10)) + te(dt, Age_bl,
##     by = Sex_ord, k = c(4, 10)) + ti(PSQI_Comp7_Tired, k = 4) +
##     ti(PSQI_Comp7_Tired, by = Sex_ord, k = 4) + ti(dt, PSQI_Comp7_Tired,
##     k = c(4, 4)) + ti(dt, PSQI_Comp7_Tired, by = Sex_ord, k = c(4,
##     4)) + ICV_st + Sex_ord
##
## Parametric coefficients:
##             Estimate Std. Error t value Pr(>|t|)
## (Intercept)  0.04224    0.04695   0.900    0.368
## ICV_st       0.28300    0.02622  10.795  < 2e-16 ***
## Sex_ord.L    0.14226    0.03644   3.904  9.7e-05 ***
## ---
## Signif. codes:  0 '***' 0.001 '**' 0.01 '*' 0.05 '.' 0.1 ' ' 1
##
## Approximate significance of smooth terms:
##                                        edf Ref.df      F p-value
## te(dt,Age_bl)                       16.052 16.052 59.256  <2e-16 ***
## te(dt,Age_bl):Sex_ordMale            5.306  5.306  1.969  0.0763 .
## ti(PSQI_Comp7_Tired)                 2.495  2.495  0.545  0.4466
## ti(PSQI_Comp7_Tired):Sex_ordMale     1.000  1.000  1.074  0.3002
## ti(dt,PSQI_Comp7_Tired)              3.047  3.047  3.017  0.0235 *
## ti(dt,PSQI_Comp7_Tired):Sex_ordMale  4.746  4.746  3.582  0.0117 *
## ---
## Signif. codes:  0 '***' 0.001 '**' 0.01 '*' 0.05 '.' 0.1 ' ' 1
##
## R-sq.(adj) =  0.472
##   Scale est. = 0.018579  n = 2893
```

The plot shows the difference between Males and Females.

plot(m2s$gam, select = 6, scheme = 2)

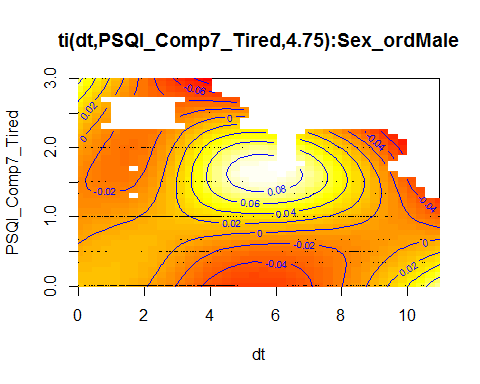

The bottom row of the plots below illustrates the three-way interaction: Males who answer 3 on the PSQI question have a higher reduction with time than females.

```
crossing(dt = 0:10,
         Age_bl = c(20, 50, 80),
         ICV_st = 0,
         Sex_ord = c("Female", "Male"),
         Study = "Betula",
         ID = "A",
         value = 0:3
         ) %>%
  mutate(!! psqi_var := value) %>%
  bind_cols(fit = predict(m2s$gam, newdata = .)) %>%
  mutate_at(vars(value, Age_bl, Sex_ord), factor) %>%
  group_by(value, Age_bl, Sex_ord) %>%
  arrange(dt, .by_group = TRUE) %>%
  mutate(fit = fit - first(fit)) %>%
  ungroup() %>%
  ggplot(aes(x = dt, y = fit, group = Sex_ord, color = Sex_ord)) +
  geom_line() +
  facet_wrap(~ paste("PSQI =", value) + Age_bl, ncol = 3,
             strip.position = "left")
```

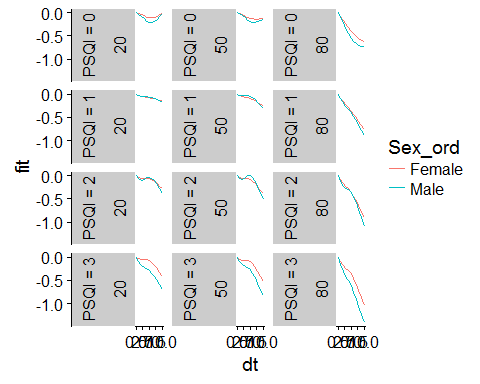

#### PSQI Global

psqi_var <- "PSQI_Global"

psqi_plot(data, psqi_var)

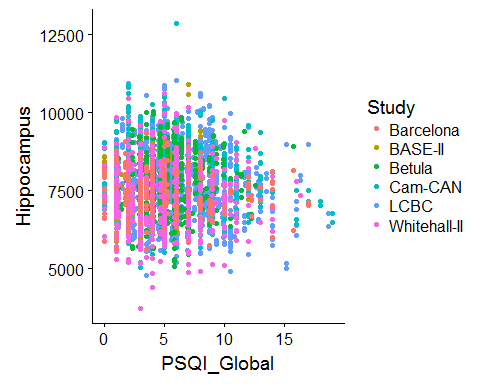

We fit one model per study first, purely for exploratory reasons.

sep_fits(data, psqi_var)

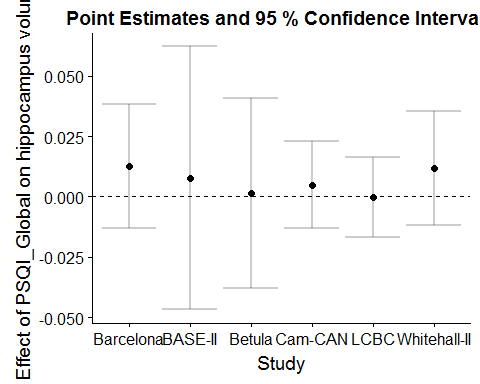

##### Main Effect of Sleep

GAMM with smooth effect of PSQI, random slope of PSQI per study, and random intercept of ID.

```
forms <- main_effects(psqi_var)
m0 <- gamm(forms[[1]], random = forms[[2]],
           data = subset_data(data, psqi_var))
saveRDS(m0, file = paste0("./models/", psqi_var, "_m0.rds"))
m0 <- readRDS(paste0("./models/", psqi_var, "_m0.rds"))
summary(m0$gam)
##
## Family: gaussian
## Link function: identity
##
## Formula:
## Hippocampus_st ~ s(Age, k = 6, bs = "cr") + s(PSQI_Global, k = 3,
##     bs = "cr") + ICV_st + Sex
##
## Parametric coefficients:
##             Estimate Std. Error t value Pr(>|t|)
## (Intercept) -0.15817    0.07110  -2.225   0.0262 *
## ICV_st       0.16867    0.01692   9.971   <2e-16 ***
## SexMale      0.36210    0.03697   9.793   <2e-16 ***
## ---
## Signif. codes:  0 '***' 0.001 '**' 0.01 '*' 0.05 '.' 0.1 ' ' 1
##
## Approximate significance of smooth terms:
##                  edf Ref.df       F p-value
## s(Age)         4.791  4.791 508.628  <2e-16 ***
## s(PSQI_Global) 1.000  1.000   0.701   0.402
## ---
## Signif. codes:  0 '***' 0.001 '**' 0.01 '*' 0.05 '.' 0.1 ' ' 1
##
## R-sq.(adj) =   0.41
##   Scale est. = 0.019255  n = 4621
plot(m0$gam, select = 2, seWithMean = TRUE)
```

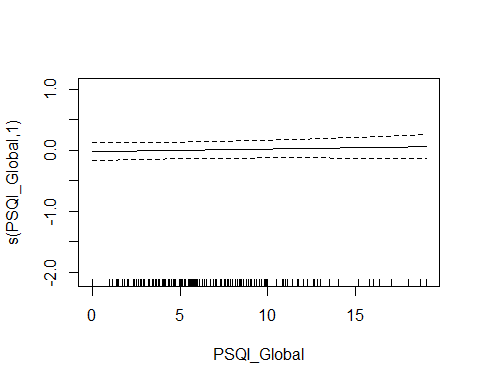

###### Controling for Covariates

```
forms <- main_effects(psqi_var, c("Depression", "BMI"))
m0_cov <- gamm(forms[[1]], random = forms[[2]],
           data = subset_data(data, psqi_var))
saveRDS(m0_cov, file = paste0("./models/", psqi_var, "_m0_cov.rds"))
m0_cov <- readRDS(paste0("./models/", psqi_var, "_m0_cov.rds"))
summary(m0_cov$gam)
##
## Family: gaussian
## Link function: identity
##
## Formula:
## Hippocampus_st ~ s(Age, k = 6, bs = "cr") + s(PSQI_Global, k = 3,
##     bs = "cr") + ICV_st + Sex + Depression + BMI
##
## Parametric coefficients:
##              Estimate Std. Error t value Pr(>|t|)
## (Intercept) -0.384289   0.165590  -2.321 0.020377 *
## ICV_st       0.084525   0.022367   3.779 0.000161 ***
## SexMale      0.357923   0.051832   6.905 6.25e-12 ***
## Depression   0.006017   0.022334   0.269 0.787647
## BMI          0.004950   0.005229   0.947 0.343925
## ---
## Signif. codes:  0 '***' 0.001 '**' 0.01 '*' 0.05 '.' 0.1 ' ' 1
##
## Approximate significance of smooth terms:
##                  edf Ref.df       F p-value
## s(Age)         4.495  4.495 264.022  <2e-16 ***
## s(PSQI_Global) 1.000  1.000   1.138   0.286
## ---
## Signif. codes:  0 '***' 0.001 '**' 0.01 '*' 0.05 '.' 0.1 ' ' 1
##
## R-sq.(adj) =  0.369
##   Scale est. = 0.023514  n = 2650
```

###### Interaction with Sex

The summary shows that no sex interaction is found.

```
forms <- main_sex_interaction(psqi_var)
m0s <- gamm(forms[[1]], random = forms[[2]],
           data = subset_data(data, psqi_var), method = "REML")

saveRDS(m0s, file = paste0("./models/", psqi_var, "_m0s.rds"))
m0s <- readRDS(paste0("./models/", psqi_var, "_m0s.rds"))
summary(m0s$gam)
##
## Family: gaussian
## Link function: identity
##
## Formula:
## Hippocampus_st ~ s(Age, k = 6, bs = "cr") + s(PSQI_Global, k = 3,
##     bs = "cr") + s(PSQI_Global, by = Sex_ord, k = 3, bs = "cr") +
##     ICV_st + Sex_ord
##
## Parametric coefficients:
##             Estimate Std. Error t value Pr(>|t|)
## (Intercept)  0.02271    0.07485   0.303    0.762
## ICV_st       0.16778    0.01693   9.910   <2e-16 ***
## Sex_ord.L    0.25535    0.02619   9.751   <2e-16 ***
## ---
## Signif. codes:  0 '***' 0.001 '**' 0.01 '*' 0.05 '.' 0.1 ' ' 1
##
## Approximate significance of smooth terms:
##                              edf Ref.df       F p-value
## s(Age)                     4.793  4.793 506.139  <2e-16 ***
## s(PSQI_Global)             1.733  1.733   2.608   0.180
## s(PSQI_Global):Sex_ordMale 1.000  1.000   0.901   0.343
## ---
## Signif. codes:  0 '***' 0.001 '**' 0.01 '*' 0.05 '.' 0.1 ' ' 1
##
## R-sq.(adj) =  0.411
##   Scale est. = 0.01926   n = 4621
```

##### Interaction with Age

```
forms <- age_interaction(psqi_var)
m1 <- gamm(forms[[1]], random = forms[[2]],
           data = subset_data(data, psqi_var), control = list(opt = "optim"))

saveRDS(m1, file = paste0("./models/", psqi_var, "_m1.rds"))
m1 <- readRDS(paste0("./models/", psqi_var, "_m1.rds"))
summary(m1$gam)
##
## Family: gaussian
## Link function: identity
##
## Formula:
## Hippocampus_st ~ s(Age, k = 6, bs = "cr") + s(PSQI_Global, bs = "cr",
##     k = 3) + ti(Age, PSQI_Global, k = c(4, 3)) + ICV_st + Sex
##
## Parametric coefficients:
##             Estimate Std. Error t value Pr(>|t|)
## (Intercept) -0.15714    0.07083  -2.218   0.0266 *
## ICV_st       0.16800    0.01692   9.930   <2e-16 ***
## SexMale      0.36264    0.03701   9.798   <2e-16 ***
## ---
## Signif. codes:  0 '***' 0.001 '**' 0.01 '*' 0.05 '.' 0.1 ' ' 1
##
## Approximate significance of smooth terms:
##                       edf Ref.df       F p-value
## s(Age)              4.791  4.791 506.175  <2e-16 ***
## s(PSQI_Global)      1.521  1.521   1.421   0.386
## ti(Age,PSQI_Global) 1.077  1.077   0.159   0.687
## ---
## Signif. codes:  0 '***' 0.001 '**' 0.01 '*' 0.05 '.' 0.1 ' ' 1
##
## R-sq.(adj) =  0.411
##   Scale est. = 0.019266  n = 4621
par(mfrow = c(1, 2))
plot(m1$gam, select = 2, seWithMean = TRUE)
fvisgam(m1$gam, view = c("Age", psqi_var))
## Summary:
##  * ICV_st : numeric predictor; set to the value(s): 0.00824076166163113.
##  * Sex : factor; set to the value(s): Female.
##  * Age : numeric predictor; with 30 values ranging from 18.482192 to 91.885010.
##  * PSQI_Global : numeric predictor; with 30 values ranging from 0.000000 to 19.000000.
```

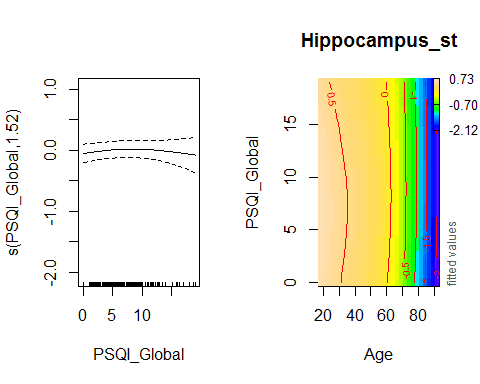

###### Controling for Covariates

```
forms <- age_interaction(psqi_var, c("Depression", "BMI"))
m1_cov <- gamm(forms[[1]], random = forms[[2]],
           data = subset_data(data, psqi_var))

saveRDS(m1_cov, file = paste0("./models/", psqi_var, "_m1_cov.rds"))
m1_cov <- readRDS(paste0("./models/", psqi_var, "_m1_cov.rds"))
summary(m1_cov$gam)
##
## Family: gaussian
## Link function: identity
##
## Formula:
## Hippocampus_st ~ s(Age, k = 6, bs = "cr") + s(PSQI_Global, bs = "cr",
##     k = 3) + ti(Age, PSQI_Global, k = c(4, 3)) + ICV_st + Sex +
##     Depression + BMI
##
## Parametric coefficients:
##              Estimate Std. Error t value Pr(>|t|)
## (Intercept) -0.381505   0.165912  -2.299 0.021557 *
## ICV_st       0.084395   0.022370   3.773 0.000165 ***
## SexMale      0.356801   0.051887   6.876 7.63e-12 ***
## Depression   0.006019   0.022336   0.269 0.787587
## BMI          0.004889   0.005231   0.935 0.350075
## ---
## Signif. codes:  0 '***' 0.001 '**' 0.01 '*' 0.05 '.' 0.1 ' ' 1
##
## Approximate significance of smooth terms:
##                       edf Ref.df       F p-value
## s(Age)              4.494  4.494 262.507  <2e-16 ***
## s(PSQI_Global)      1.000  1.000   1.331   0.249
## ti(Age,PSQI_Global) 1.000  1.000   0.274   0.601
## ---
## Signif. codes:  0 '***' 0.001 '**' 0.01 '*' 0.05 '.' 0.1 ' ' 1
##
## R-sq.(adj) =  0.368
##   Scale est. = 0.023499  n = 2650
```

###### Interaction with Age and Sex

No sex interaction in the interaction between Age and Sleep was found:

```
forms <- age_sex_interaction(psqi_var)
m1s <- gamm(forms[[1]], random = forms[[2]],
            data = subset_data(data, psqi_var))

saveRDS(m1s, file = paste0("./models/", psqi_var, "_m1s.rds"))
m1s <- readRDS(paste0("./models/", psqi_var, "_m1s.rds"))
summary(m1s$gam)
##
## Family: gaussian
## Link function: identity
##
## Formula:
## Hippocampus_st ~ s(Age, k = 6, bs = "cr") + s(Age, by = Sex_ord,
##     k = 3, bs = "cr") + s(PSQI_Global, k = 3, bs = "cr") + s(PSQI_Global,
##     by = Sex_ord, k = 3, bs = "cr") + ti(Age, PSQI_Global, k = c(4,
##     3)) + ti(Age, PSQI_Global, by = Sex_ord, k = c(4, 3)) + ICV_st +
##     Sex_ord
##
## Parametric coefficients:
##             Estimate Std. Error t value Pr(>|t|)
## (Intercept)  0.03003    0.06492   0.463    0.644
## ICV_st       0.16761    0.01689   9.924   <2e-16 ***
## Sex_ord.L    0.25150    0.02617   9.611   <2e-16 ***
## ---
## Signif. codes:  0 '***' 0.001 '**' 0.01 '*' 0.05 '.' 0.1 ' ' 1
##
## Approximate significance of smooth terms:
##                                   edf Ref.df       F  p-value
## s(Age)                          4.787  4.787 297.270  < 2e-16 ***
## s(Age):Sex_ordMale              1.000  1.000  12.632 0.000383 ***
## s(PSQI_Global)                  1.635  1.635   2.095 0.261225
## s(PSQI_Global):Sex_ordMale      1.000  1.000   0.666 0.414563
## ti(Age,PSQI_Global)             1.000  1.000   0.154 0.694470
## ti(Age,PSQI_Global):Sex_ordMale 1.000  1.000   0.128 0.720278
## ---
## Signif. codes:  0 '***' 0.001 '**' 0.01 '*' 0.05 '.' 0.1 ' ' 1
##
## R-sq.(adj) =  0.415
##   Scale est. = 0.019264  n = 4621
```

##### Longitudinal Analysis

In this section, only participants with two timepoints or more are retained.

Interaction with time.

```
forms <- time_interaction(psqi_var)
m2 <- gamm(forms[[1]], random = forms[[2]],
           data = long_only(subset_data(data, psqi_var)),
           method = "REML")
saveRDS(m2, file = paste0("./models/", psqi_var, "_m2.rds"))
m2 <- readRDS(paste0("./models/", psqi_var, "_m2.rds"))
summary(m2$gam)
##
## Family: gaussian
## Link function: identity
##
## Formula:
## Hippocampus_st ~ te(dt, Age_bl, k = c(4, 10)) + ti(PSQI_Global,
##     k = 4) + ti(dt, PSQI_Global, k = c(4, 4)) + ICV_st + Sex
##
## Parametric coefficients:
##             Estimate Std. Error t value Pr(>|t|)
## (Intercept) -0.04320    0.06192  -0.698    0.485
## ICV_st       0.28908    0.02670  10.829  < 2e-16 ***
## SexMale      0.20910    0.05244   3.988 6.84e-05 ***
## ---
## Signif. codes:  0 '***' 0.001 '**' 0.01 '*' 0.05 '.' 0.1 ' ' 1
##
## Approximate significance of smooth terms:
##                       edf Ref.df       F p-value
## te(dt,Age_bl)      16.056 16.056 110.217  <2e-16 ***
## ti(PSQI_Global)     2.106  2.106   1.113  0.2557
## ti(dt,PSQI_Global)  1.731  1.731   2.264  0.0696 .
## ---
## Signif. codes:  0 '***' 0.001 '**' 0.01 '*' 0.05 '.' 0.1 ' ' 1
##
## R-sq.(adj) =  0.471
##   Scale est. = 0.019247  n = 2843
fvisgam(m2$gam, view = c("dt", psqi_var))
## Summary:
##  * ICV_st : numeric predictor; set to the value(s): -0.00264684881092563.
##  * Sex : factor; set to the value(s): Female.
##  * dt : numeric predictor; with 30 values ranging from 0.000000 to 10.976044.
##  * Age_bl : numeric predictor; set to the value(s): 64.8213552361396.
##  * PSQI_Global : numeric predictor; with 30 values ranging from 0.000000 to 19.000000.
```

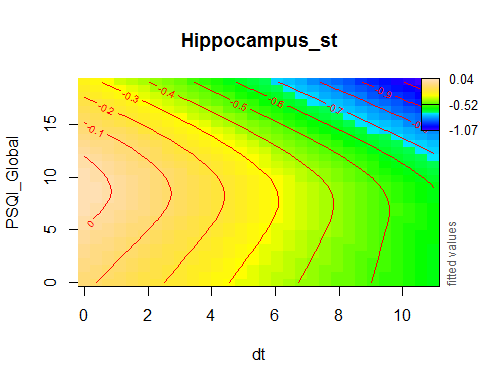

plot_psqi_interaction(m2, psqi_var, psqi_range = 0:3, time_range = 0:5)

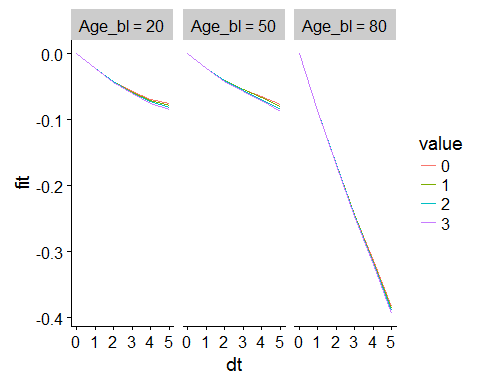

###### Controling for Covariates

```
forms <- time_interaction(psqi_var, c("Depression", "BMI"))
m2_cov <- gamm(forms[[1]], random = forms[[2]],
           data = long_only(subset_data(data, psqi_var)),
           control = list(opt = "optim"))
saveRDS(m2_cov, file = paste0("./models/", psqi_var, "_m2_cov.rds"))
m2_cov <- readRDS(paste0("./models/", psqi_var, "_m2_cov.rds"))
summary(m2_cov$gam)
##
## Family: gaussian
## Link function: identity
##
## Formula:
## Hippocampus_st ~ te(dt, Age_bl, k = c(4, 10)) + ti(PSQI_Global,
##     k = 4) + ti(dt, PSQI_Global, k = c(4, 4)) + ICV_st + Sex +
##     Depression + BMI
##
## Parametric coefficients:
##              Estimate Std. Error t value Pr(>|t|)
## (Intercept) -0.579490   0.205526  -2.820  0.00487 **
## ICV_st       0.328990   0.036169   9.096  < 2e-16 ***
## SexMale      0.128827   0.067408   1.911  0.05616 .
## Depression  -0.010692   0.031400  -0.341  0.73351
## BMI          0.021934   0.007435   2.950  0.00322 **
## ---
## Signif. codes:  0 '***' 0.001 '**' 0.01 '*' 0.05 '.' 0.1 ' ' 1
##
## Approximate significance of smooth terms:
##                       edf Ref.df      F p-value
## te(dt,Age_bl)      12.185 12.185 80.735  <2e-16 ***
## ti(PSQI_Global)     1.007  1.007  0.338   0.558
## ti(dt,PSQI_Global)  1.001  1.001  1.779   0.183
## ---
## Signif. codes:  0 '***' 0.001 '**' 0.01 '*' 0.05 '.' 0.1 ' ' 1
##
## R-sq.(adj) =  0.494
##   Scale est. = 0.023419  n = 1669
```

###### Interaction with Sex and Time

As the summary shows, no sex interaction was found.

```
forms <- sex_time_interaction(psqi_var)
m2s <- gamm(forms[[1]], random = forms[[2]],
            data = long_only(subset_data(data, psqi_var)), method = "REML")

saveRDS(m2s, file = paste0("./models/", psqi_var, "_m2s.rds"))
m2s <- readRDS(paste0("./models/", psqi_var, "_m2s.rds"))
summary(m2s$gam)
##
## Family: gaussian
## Link function: identity
##
## Formula:
## Hippocampus_st ~ te(dt, Age_bl, k = c(4, 10)) + te(dt, Age_bl,
##     by = Sex_ord, k = c(4, 10)) + ti(PSQI_Global, k = 4) + ti(PSQI_Global,
##     by = Sex_ord, k = 4) + ti(dt, PSQI_Global, k = c(4, 4)) +
##     ti(dt, PSQI_Global, by = Sex_ord, k = c(4, 4)) + ICV_st +
##     Sex_ord
##
## Parametric coefficients:
##             Estimate Std. Error t value Pr(>|t|)
## (Intercept)  0.05436    0.05504   0.988 0.323347
## ICV_st       0.28871    0.02665  10.833  < 2e-16 ***
## Sex_ord.L    0.14083    0.03712   3.794 0.000151 ***
## ---
## Signif. codes:  0 '***' 0.001 '**' 0.01 '*' 0.05 '.' 0.1 ' ' 1
##
## Approximate significance of smooth terms:
##                                   edf Ref.df      F p-value
## te(dt,Age_bl)                  15.413 15.413 58.854  <2e-16 ***
## te(dt,Age_bl):Sex_ordMale       6.487  6.487  1.725  0.1048
## ti(PSQI_Global)                 2.321  2.321  2.262  0.0648 .
## ti(PSQI_Global):Sex_ordMale     1.000  1.000  4.358  0.0369 *
## ti(dt,PSQI_Global)              1.000  1.000  5.705  0.0170 *
## ti(dt,PSQI_Global):Sex_ordMale  1.960  1.960  1.172  0.3557
## ---
## Signif. codes:  0 '***' 0.001 '**' 0.01 '*' 0.05 '.' 0.1 ' ' 1
##
## R-sq.(adj) =  0.474
##   Scale est. = 0.019159  n = 2843
```

### Plotting

```
df <- imap_dfr(main_models, function(x, y){
  grid <- tibble(
  Age = 0, ICV_st = 0, Sex = "Female",
  !!y := unique(as.integer(x$gam$model[[y]]))
  )

  pred <- predict(x$gam, newdata = grid, se.fit = TRUE,
                  terms = paste0("s(", y, ")"), type = "iterms")

pred$fit <- (pred$fit + attr(pred, "constant")) * sd(full_data$Hippocampus) + mean(full_data$Hippocampus)
pred$se.fit <- pred$se.fit * sd(full_data$Hippocampus)

grid %>%
  bind_cols(pred) %>%
  rename_at(vars(matches(y)), funs(function(x) "xvar")) %>%
  select(xvar, fit, se.fit) %>%
  mutate(PSQI = y)
})

p <- ggplot(df, aes(x = xvar)) +
  geom_line(aes(y = fit)) +
  geom_line(aes(y = fit - 2 * se.fit), linetype = "dashed") +
  geom_line(aes(y = fit + 2 * se.fit), linetype = "dashed") +
  facet_wrap(~ PSQI, scales = "free_x", ncol = 2) +
  xlab("") +
  ylab("") +
  ylim(7000, 8000) +
  theme_classic() +
  theme(strip.background = element_blank())


p
ggsave("psqi_hippocampus_grid.png", plot = p, dpi = 900,
       height = 28, width = 20, units = "cm")
```
