## Supplementary material for "Self-reported sleep relates to hippocampal atrophy across the adult lifespan – results from the Lifebrain consortium": SI

Interpreting Interactions

The plot below considers the following set-up:

- We start at a baseline age of 20, 40, 60, or 80 years. Then we compute the expected change in hippocampus volume over the next year (dt = 1) when the corresponding PSQI score is 0 vs. 2, for each of the four PSQI variables that showed a significant interaction with age.
- The points show the expected value, and the errorbars show 95 % confidence intervals.
- Note that variables that only affect the offset (Sex, ICV) do not influence the change in these plots.

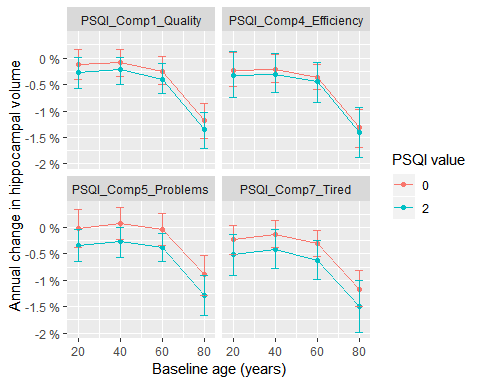

\## Saving 5 x 4 in image

The table below summarizes the mean annual change values from the previous plot.

Expected annual change for high and low PSQI score and different baseline ages.

| Age_bl | PSQI_value | PSQI_Comp1_Quality | PSQI_Comp4_Efficiency | PSQI_Comp5_Problems | PSQI_Comp7_Tired |
| --- | --- | --- | --- | --- | --- |
| 20 | 0 | -0.13 % | -0.24 % | -0.02 % | -0.23 % |
| 20 | 2 | -0.27 % | -0.32 % | -0.35 % | -0.51 % |
| 40 | 0 | -0.08 % | -0.22 % | 0.07 % | -0.13 % |
| 40 | 2 | -0.22 % | -0.3 % | -0.26 % | -0.42 % |
| 60 | 0 | -0.25 % | -0.36 % | -0.03 % | -0.31 % |
| 60 | 2 | -0.4 % | -0.45 % | -0.39 % | -0.62 % |
| 80 | 0 | -1.18 % | -1.32 % | -0.9 % | -1.17 % |
| 80 | 2 | -1.36 % | -1.41 % | -1.29 % | -1.5 % |

The next table shows the probability that someone giving a PSQI score of 2 experiences larger decline over the next year than someone giving a PSQI score of 0.

Probability that someone given a high PSQI score has a larger decline over the next year than someone giving a low PSQI score.

| Age_bl | PSQI_Comp1_Quality | PSQI_Comp4_Efficiency | PSQI_Comp5_Problems | PSQI_Comp7_Tired |
| --- | --- | --- | --- | --- |
| 20 | 0.888 | 0.698 | 0.978 | 0.920 |
| 40 | 0.888 | 0.700 | 0.976 | 0.922 |
| 60 | 0.886 | 0.696 | 0.978 | 0.922 |
| 80 | 0.886 | 0.680 | 0.974 | 0.916 |
