## Supplementary material for "Self-reported sleep relates to hippocampal atrophy across the adult lifespan – results from the Lifebrain consortium": SI

**for**

### Introduction

This document describes power simulations performed to understand why some of the change interactions ti(dt, PSQI_*) were significant in the longitudinal models, while the corresponding age interactions ti(Age, PSQI_*) were not significant in the corresponding model with both longitudinal and cross-sectional observations. As change interactions in principle should induce age interactions in the data, one might suspect that this is a power issue. We therefore start with the models for hippocampus volume as a function of Age, ICV, and Sex, and add change interactions of various strength.

library(tidyverse)
library(mgcv)
full_data <- readRDS("./data/full_data.rds")

We load the models fitted for hippocampus volume as a function of age.

m0_long <- readRDS("./models/long_age_hippo_m0.rds")
m0_cross <- readRDS("./models/age_hippo_m0.rds")

### PSQI Component 4: Efficiency

We start with PSQI_Comp4_Efficiency because this component has a significant change interaction and a non-significant age interaction.

psqi_var <- "PSQI_Comp4_Efficiency"

We load the models and show the outputs here for ease of reference.

m1 <- readRDS(paste0("./models/", psqi_var, "_m1.rds"))
knitr::kable(summary(m1$gam)$s.table)

|  | edf | Ref.df | F | p-value |
| --- | --- | --- | --- | --- |
| s(Age) | 4.781455 | 4.781455 | 438.2652937 | 0.0000000 |
| s(PSQI_Comp4_Efficiency) | 1.000006 | 1.000006 | 0.1014040 | 0.7501702 |
| ti(Age,PSQI_Comp4_Efficiency) | 1.586845 | 1.586845 | 0.4478028 | 0.4381368 |

m2 <- readRDS(paste0("./models/", psqi_var, "_m2.rds"))
knitr::kable(summary(m2$gam)$s.table)

|  | edf | Ref.df | F | p-value |
| --- | --- | --- | --- | --- |
| te(dt,Age_bl) | 16.332151 | 16.332151 | 99.8232682 | 0.0000000 |
| ti(PSQI_Comp4_Efficiency) | 1.000002 | 1.000002 | 0.4930246 | 0.4826408 |
| ti(dt,PSQI_Comp4_Efficiency) | 6.304656 | 6.304656 | 6.7466878 | 0.0000001 |

We subset the data and drop rows with missing values.

```
data <- full_data %>%
  select_at(vars(Age, Age_bl, dt, psqi_var, ICV, Hippocampus, Sex, ID, Study, Site_Name)) %>%
  mutate_at(vars(ICV, Hippocampus), funs(st = (. - mean(.))/sd(.))) %>%
  na.omit()
```

We see from the lme part of the fitted models that the random intercept has standard deviation about 0.7-0.8 within ID and about 0.1 within Study. The residual is around 0.13-0.14. The table below summarizes.

```
map_dfr(list(change_model = m0_long, age_model = m0_cross,
             age_interaction = m1, change_interaction = m2), function(x) {
  corrs <- c(
    as.numeric(VarCorr(x$lme)[grep("Intercept", rownames(VarCorr(x$lme))), "StdDev"]),
    x$lme$sigma)
  corrs <- as_tibble(matrix(corrs, nrow = 1))
  colnames(corrs) <- c("Within-study corr.", "Within-ID corr.", "Residual s.d.")
  corrs
}, .id = "Model") %>%
  knitr::kable()
```

| Model | Within-study corr. | Within-ID corr. | Residual s.d. |
| --- | --- | --- | --- |
| change_model | 0.0860546 | 0.7046787 | 0.1373831 |
| age_model | 0.1653622 | 0.7658401 | 0.1379021 |
| age_interaction | 0.1755314 | 0.7740073 | 0.1341669 |
| change_interaction | 0.0602776 | 0.7087345 | 0.1319974 |

We use these estimates in the following function, which simulates hippocampus volumes based on the longitudinal model.

```
simulate_random <- function(data, psqi_var){
  # Random effects per unique study
  studies <- unique(as.character(data$Study))
  study_effect <- rnorm(length(studies), sd = 0.1)
  names(study_effect) <- studies

  # Random effects per unique ID
  IDs <- unique(as.character(data$ID))
  ID_effect <- rnorm(length(IDs), sd = 0.7)
  names(ID_effect) <- IDs

  # Sample new hippocampus volumes, and then center dt and PSQI
  tmp <- data %>%
    mutate(
      study_effect = !!study_effect[as.character(.$Study)],
      ID_effect = !!ID_effect[as.character(.$ID)],
      residual = rnorm(nrow(.), sd = 0.14),
      Hippocampus_rnd = as.numeric(predict(m0_long$gam, newdata = .) + study_effect + ID_effect + residual)
    ) %>%
    mutate_at(vars(dt, psqi_var), funs(c = (. - mean(.))))
}
```

The plots below shows one realized random sample (Hippocampus_rnd, left) and the real dataset (Hippocampus_st, right). The values are quite similar. The color scale shows the value given on PSQI Component 4.

```
data %>%
  simulate_random(psqi_var) %>%
  gather(key = "key", value = "value", Hippocampus_rnd, Hippocampus_st) %>%
  ggplot(aes(x = Age, y = value, group = ID, color = PSQI_Comp4_Efficiency)) +
  geom_point() +
  geom_line() +
  facet_wrap(~ key)
```

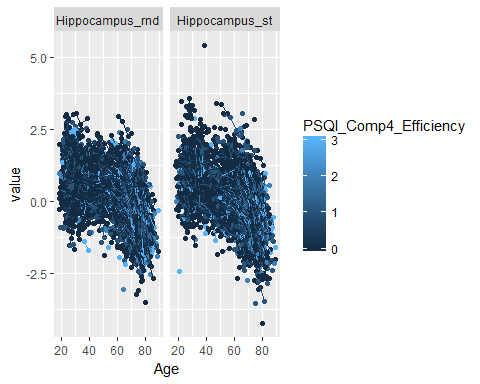

We now proceed be adding an interaction between PSQI_Comp4_Efficiency and change, dt. This will serve to change the individual slopes in the plots above, depending on the PSQI value. From the investigations in the document describing the effect of sleep on hippocampus, we know that an approximate effect size of the linear interaction dt:PSQI_Comp4_Efficiency is -0.008 when both variables are centered, while the offset effect of PSQI_Comp4_Efficiency is about 0.02, but only the interaction is significant. We would expect such an interaction to also yield a significant age interaction, since participants grow old when dt is increase, so our lack of significance for this term might be a power issues. We hence simulate adding interactions of different magnitudes, and see when we have power to detect ti(Age, PSQI_Comp4_Efficiency) and ti(dt, PSQI_Comp4_Efficiency).

Before we start simulating, we illustrate the interaction effect with a strong magnitude of -0.02, in the plow below. The dashed lines show the simulated hippocampus volumes before the interaction is added, while the solid lines show the volumes after the interaction is added. Note that the negative interaction serves to increase the atrophy for those with sleep problems as measured by PSQI Component 4, while it reduces the atrophy for those who score 0 on this component. A subset of the data was extracted for this plot, in order to increase readability.

```
data %>%
  simulate_random(psqi_var) %>%
  mutate(
    PSQI_integer = factor(round(PSQI_Comp4_Efficiency, 0)),
    Hippocampus_intrx = Hippocampus_rnd + -0.02 * PSQI_Comp4_Efficiency_c * dt_c
    ) %>%
  group_by(PSQI_integer) %>%
  filter(row_number() < 200) %>%
  gather(key = "key", value = "value", Hippocampus_rnd, Hippocampus_intrx) %>%
  ggplot(aes(x = Age, y = value, group = paste(ID, key), linetype = key)) +
  geom_point() +
  geom_line() +
  facet_wrap(~ paste("PSQI_Comp4_Efficiency =", PSQI_integer))
```

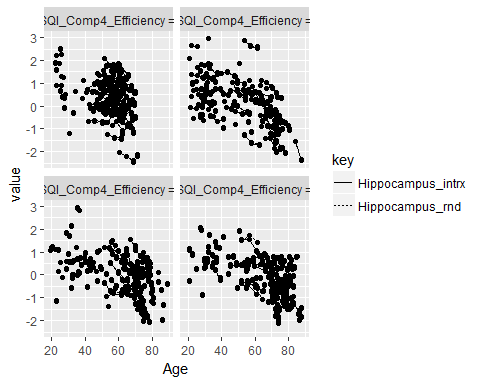

We first set the simulation parameters:

```
interaction_size <- c(0, -0.002, -0.004, -0.006, -.008,
                      -0.01, -0.012, -0.014, -0.016, -0.018, -0.02)

nsim <- 10
```

Next, we perform the actual simulations. The use of tryCatch() in the code is there because there might be some of the simulated datasets for which gamm() fails to converge. Rather than failing, the p-values in these cases are recorded as missing values.

```
sims <- map_dfr(interaction_size, function(x){
  map_dfr(1:nsim, function(i){
    df <- data %>%
      simulate_random(psqi_var) %>%
      mutate(Hippocampus_rnd = Hippocampus_rnd + x * PSQI_Comp4_Efficiency_c * dt_c)

    pval_long <- tryCatch({
       mod_long <- gamm(Hippocampus_rnd ~ te(dt_c, Age_bl, k = c(4, 10)) +
                          ti(PSQI_Comp4_Efficiency_c, k = 4) + ICV_st + Sex +
                       ti(dt_c, PSQI_Comp4_Efficiency_c, k = c(4, 4)),
                     random = list(Study =~ 1, ID =~ 1), data = df,
                     control = list(opt = "optim"))
       summary(mod_long$gam)$s.table["ti(dt_c,PSQI_Comp4_Efficiency_c)", "p-value"]
       },
       error = function(e) NA_real_
       )

    pval_cross <- tryCatch({
      mod_cross <- gamm(Hippocampus_rnd ~ s(Age, k = 10, bs = "cr") +
                          s(PSQI_Comp4_Efficiency, bs = "cr", k = 3) +
                          ti(Age, PSQI_Comp4_Efficiency, k = c(10, 3)) + ICV_st + Sex,
                        random = list(Study =~ 1, ID =~ 1), data = df,
                        control = list(opt = "optim"))
      pval_cross <- summary(mod_cross$gam)$s.table["ti(Age,PSQI_Comp4_Efficiency)", "p-value"]
      },
      error = function(e) NA_real_)

    tibble(
      interaction = x,
      simulation = i,
      pval_long = pval_long,
      pval_cross = pval_cross
    )
    })
  })


saveRDS(sims, "./simulations/psqi_comp4_interaction.rds")
```

The figure below shows the simulated power curves. The vertical dashed line is approximately the interaction strength estimated for ti(dt, PSQI_Comp4_Efficiency), and as we can see, we have power to detect this interaction longitudinally but not cross-sectionally.

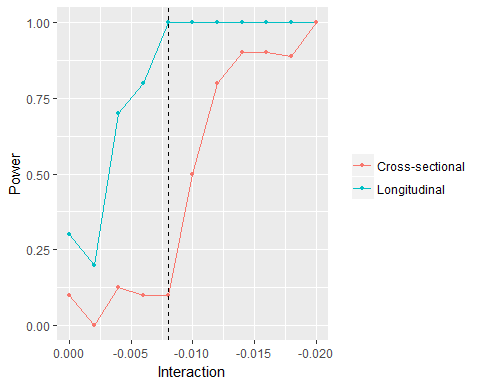
